# Translation in the cell under fierce competition for shared resources: a mathematical model

**DOI:** 10.1101/2022.07.24.501278

**Authors:** Rami Katz, Elad Attias, Tamir Tuller, Michael Margaliot

## Abstract

During the process of translation the mRNAs in the cell “compete” for shared resources like tRNA molecules and ribosomes. This creates an indirect and intricate coupling between the mRNAs. For example, if ribosomal “traffic jams” evolve on some mRNA then the abundance of free ribosomes may decrease leading to lower initiation rates in the other mRNAs. When the shared resources are abundant the coupling between mRNAs due to this competition is weak. However, when the resources are scarce, e.g., when the pool of free ribosomes is starved, the competition may have a dramatic effect on the dynamics of translation in the cell. This scenario may be relevant for example under stress conditions or during a high yield viral infection, where the viral mRNAs “hijack” components of the translation machinery. Fierce competition for shared resources may also take place in synthetic or engineered systems such as cell free systems or in the case of high-throughput heteroglougs gene expression.

We study this scenario using a mathematical model that includes a network of *m* ribosome flow models (RFMs) interconnected via a pool of free ribosomes. Each RFM is a non-linear dynamical model for ribosome flow along a single mRNA molecule, and the interconnection via the pool encapsulates the competition for shared resources. We analyze the case where *m* is large, i.e., a there is a large number of mRNAs. This implies that many ribosomes are attached to the mRNAs and thus the pool is starved.

Our model allows quantitative and qualitative analysis of the network steady state when the pool is starved. Our analysis results show that adding an mRNA to the network always decreases the steady state pool density. This makes sense, as every new mRNA “consumes” ribosomes. We also show that adding an mRNA has an intricate effect on the total protein production in the network: on the one-hand, the new mRNA produces new proteins. On the other-hand, the other mRNAs produce less proteins, as the pool that feeds these mRNAs now has a smaller abundance of ribosomes. Our analysis yields an explicit bound for the total production rate of the network when the number of RFMIOs is very large. In particular, we analyze how the total density of ribosomes in the network bounds the total production rate. This bound demonstrates that when the number of mRNAs increases, the marginal utility of adding another mRNA diminishes, and the total protein production rate saturates to a limiting value. We demonstrate our analysis approach using an example of producing insulin in a cell free system.

## I. Introduction

mRNA translation is a fundamental process in gene expression, i.e., the transformation of genetic information into a functional protein [2]. During translation, the ribosomes (complex macro-molecules) scan the mRNA in a sequential manner, “read” it codon by codon, and build a corresponding chain of amino-acids. When the ribosome reaches a stop codon, it detaches from the mRNA and releases the chain that, after additional processing, becomes a functional protein.

Many ribosomes may attach to the same mRNA and translate it in parallel. This pipelining increases the protein production rate. The speed of the ribosome movement along the mRNA is determined by the mRNA sequence and structure, pauses due to collisions with other ribosomes, and by various translation factors, e.g., the abundance of cognate tRNA molecules in the vicinity of the mRNA. Understanding the dynamics of translation is of considerable importance, as it plays a major role in determining the protein production rate [17], [45]. Furthermore, the dynamics of mRNA translation and specifically the evolution of ribosomal “traffic jams” [12], [16] along an mRNA have been implicated to various diseases [43], [39]. Computational models for ribosome flow along an mRNA and the resulting protein production rate can be used to integrate and explain the growing number of experimental findings (e.g., via methods like ribosome profiling [19] that can be performed for single cells [20], and methods for imaging the translation of a single mRNA molecule [51]). Such models can also predict the effect of various manipulations or regulation of the genetic machinery on the protein production rate. Such manipulations are common both in biotechnology and also during viral infection, where the virus “hijacks” and potentially shuts down parts of the host cell translation machinery. For example, SARS-CoV-2 uses a multipronged strategy to impede host protein synthesis and affect translation in a global manner [13]. In addition, heterologous genes tend to overload the translation machinery and also have a global effect on translation [37], [9].

A popular model for ribosome flow, and many other natural and artificial systems and processes, is the totally asymmetric simple exclusion process (TASEP) (see, e.g., [40], [6]). This is a stochastic discrete-time process that includes a one dimensional chain of sites and particles that hop stochastically along the chain in a unidirectional manner. Simple exclusion refers to the fact that two particles cannot be in the same site at the same time, i.e., a particle can only hop to an empty site. In the context of translation, the particles are ribosomes and the sites are consecutive (groups of) codons along the mRNA [50], [52], [15]. Simple exclusion corresponds to the fact that a ribosome cannot “overtake” a ribosome in front of it, as the information on the mRNA must be decoded in a sequential manner. TASEP has become a phenomenological model in statistical mechanics, yet rigorous analysis of this model is difficult, except for some very special cases, e.g., when all the internal hopping rates are assumed to be equal [6].

The ribosome flow model (RFM) is the dynamic mean-field approximation of TASEP. This yields a continuous-time, deterministic, non-linear model for the flow of ribosomes along the mRNA [36]. The RFM is highly amenable to analysis using tools from systems and control theory, and it has been used to model and analyze many important aspects of translation including: entrainment of the protein production rate to periodic initiation and elongation rates with a common period [26], sensitivity analysis of the steady-state production rate [33], optimizing the protein production rate subject to convex constraints on the rates [32], the effect of ribosome recycling [28], [34], determining the ribosome density that maximizes protein production [48], stochastic variability in translation [29], maximizing the average protein production [1], and more [5].

The cell is a factory for producing proteins that includes a large number of mRNA molecules and ribosomes. For example, an *S. cerevisiae* cell includes about 60,000 mRNA molecules and 200,000 ribosomes. About 85% of the ribosomes are associated with mRNAs [30], [49], [53], and the rest form the “pool of free ribosomes”. The mRNAs thus indirectly “compete” for the available ribosomes. This generates a network with intricate indirect coupling between the mRNAs. For example, ribosomal “traffic jams” on mRNAs may deplete the pool leading to lower initiation rates in the other mRNAs. When the shared resources are abundant the coupling between mRNAs due to this competition is weak, and the network can potentially be analyzed using models of translation along a single, isolated RNA. However, when the resources are scarce, e.g., when the pool of free ribosomes is starved, the competition may have a strong effect on the global dynamics of translation in the cell.

The latter scenario is relevant both in synthetic systems, where the goal is to optimize the production rate, and under physiological conditions. For example, under stress conditions or during a high yield viral infection, where the viral mRNAs “hijack” the translation machinery, and consume many of the shared resources to produce viral proteins. It is also relevant in heterologous gene expression (e.g., when the heterologous gene is overexpressed and consumes most of the ribosomes in the cell) and in cell free systems where the number of mRNA molecules may be relatively large in comparison to the number of ribosomes [22].

We study this scenario using a mathematical model that includes a network of *m* ribosome flow models (RFMs) interconnected via a pool of free ribosomes. The RFM belongs to the class of compartmental models that have been used in various domains of systems biology including genetics, physiology, and pharmacology [38]. Each RFM describes the dynamics of translation along one mRNA molecule, and the interconnection via the pool encapsulates the competition for shared resources. This model was first suggested in [35]. Mathematically, it is a cooperative dynamical system [41], [3] that admits a first integral: the total density of ribosomes in the network is conserved. Such systems have a well-ordered asymptotic behaviour [31], [4]. It was shown in [35] that any solution of the network converges to a steady state, where the flow of ribosomes into and out of any site along any mRNA is equal. Also, the flows in and out of the pool are equal. Sensitivity analysis of this steady state [35] with respect to modifying one of the translation rates in a specific mRNA demonstrated that the steady-state production rates in all the other mRNAs either all increase or all decrease.

A generalization of this network, that includes the possibility of ribosome drop-off and attachment at each site along the RFM, was recently suggested in [21]. Simulations of this model showed that ribosome drop-off from an isolated mRNA always decreases the protein production rate, yet in the network ribosome drop-off from a jammed mRNA may increase the total production rate of all the mRNAs, as the drop-off frees ribosomes that enter the pool, and this improves the production rate in the other mRNAs. This illustrates how the network perspective provides new biological insights.

In this paper, we analyze this network of *m* RFMs interconnected via a pool from a new, structural perspective, that is, we study how adding new RFMs to the network affects the dynamics. The main contributions of this paper include the following:

1. We prove that adding an RFM to the network always decreases the steady state pool density. This makes sense, as every new RFM “consumes” ribosomes;
2. We show that adding an RFM has an intricate effect on the network steady state: on the one-hand, the additional RFM produces new proteins. On the other-hand, the other RFMs produce less proteins, as the pool that feeds these RFMs now has a smaller abundance of ribosomes;
3. We provide a detailed asymptotic analysis of the network steady state when the pool is starved. In particular, we show that in this case the initiation rates in every RFM become the bottleneck rates, and provide a closed-form expression for the total steady state density and total production rate on any subset of RFMs in the network, relative to the steady state pool density.
4. We provide an explicit bound for the total production rate of the network when the number of RFMs is very large. In particular, we show how the total density of ribosomes in the networks bounds the total production rate. This bound demonstrates that when the number of RFMs increases the marginal utility of adding another RFM diminishes, and the total protein production saturates to a limiting value.
5. We demonstrate our analysis approach for the case of producing insulin proteins in a cell free system, and show how it can provide useful guidelines for setting the parameters in such a system.

The remainder of this paper is organized as follows. The next section reviews the network model. Before going into the mathematical analysis, Section III illustrates several structural questions that can be studied using simulations of the model. Section IV details the main mathematical results. Section V reports analyses of a biological system (gene expression in a cell free system) with our models. For the sake of readability, all the proofs are placed in the Appendix. Throughout, we try to describe the biological implications of every analysis result.

## II. The mathematical model

We use a model that includes *m* RFMs interconnected via a pool of free ribosomes. Each RFM models the dynamics of ribosome flow along one mRNA molecule and, in particular, each RFM may have a different length and different parameters (i.e., different codon decoding rates and initiation rates). The pool of free ribosomes represents ribosomes in the cell that are not attached to any mRNA. We begin by reviewing the various components of this networks.

### 1) Ribosome flow model

The RFM includes *n* state-variables *x*_1_, …, *x*_*n*_ representing the normalized ribosome density in *n* sites along the mRNA, where each site corresponds to a group of consecutive codons. The density is normalized such that *x*_*i*_(*t*) ∈ [0, 1] for all *t*, where *x*_*i*_(*t*) = 0 represents that the site is empty, and *x*_*i*_(*t*) = 1 represents that the site is completely full. Thus, *x*_*i*_(*t*) may also be interpreted as the probability that site *i* is occupied at time *t*. The RFM also includes *n* + 1 positive parameters *λ*_0_, …, *λ*_*n*_, where *λ*_*i*_ controls the transition rate from site *i* to site *i* + 1. In particular *λ*_0_ controls the initiation rate, and *λ*_*n*_ controls the exit rate.

The dynamics of the RFM is described by *n* balance equations:

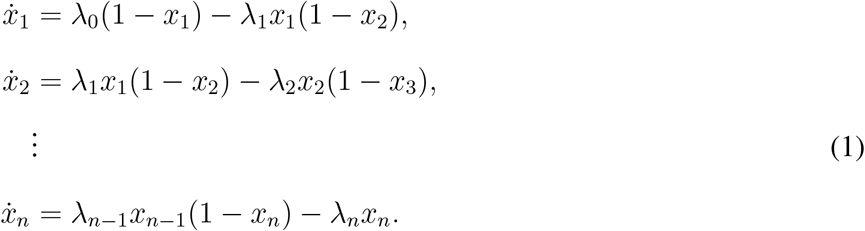

To explain this, consider the equation for the change in density in the second site, namely,

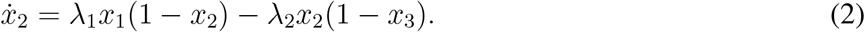

The term *λ*_1_*x*_1_(1 *− x*_2_) represents the flow of ribosomes from site 1 to site 2. This is proportional to the transition rate *λ*_1_, to the density of ribosomes *x*_1_ in site 1 and to the “free space” (1 *− x*_2_) in site 2. In particular, if site 2 fills up, i.e., *x*_2_ is close to one, then the flow into site 2 decreases to zero. This is a “soft” version of the simple exclusion principle, i.e., the notion that two particles cannot be in the same place at the same time. Similarly, the second term on the right-hand side of (2) is the flow from site 2 to site 3. Thus, Eq. (2) states that the change in density in site 2 is the flow from site 1 to site 2 minus the flow from site 2 to site 3. The exit rate from the last site is *R*(*t*) := *λ*_*n*_*x*_*n*_(*t*), and this is also the protein production rate at time *t* (see Fig. 1). Note that *x*_*i*_ is dimensionless, and that *λ*_*i*_ has units of 1*/*time. In all the biological simulations below, *λ*_*i*_ is in units of 1*/*sec.

**Fig. 1:**
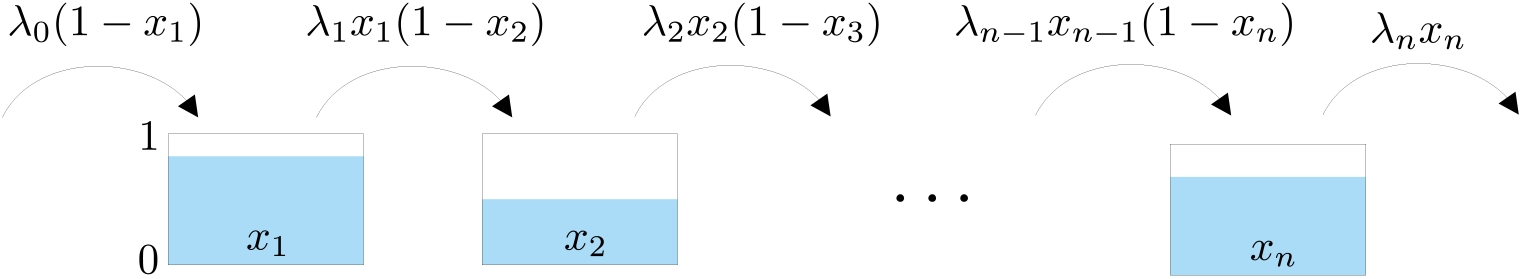
Ribosome flow model.

The state space of the RFM is the *n*-dimensional cube [0, 1]^*n*^. Since this invariant set is compact and convex, the RFM admits an equilibrium point *e* ∈ [0, 1]^*n*^. At an equilibrium, the flows into each site and out of each site are equal, so all the site densities are constant. Analysis of the equations describing the equilibrium point shows that there is a single equilibrium point *e* ∈ [0, 1]^*n*^ [27].

The RFM has been extensively used for studying the translation of a single, isolated mRNA. The model is highly amenable to analysis using tools from systems and control theory. It was shown in [27] that the RFM is a totally positive differential system [25] and this implies that any solution of the RFM converges to the unique equilibrium *e*. In particular, the protein production rate *R*(*t*) = *λ*_*n*_*x*_*n*_(*t*) converges to the steady-state production rate *R* := *λ*_*n*_*e*_*n*_. In other words, the positive transition rates *λ*_0_, …, *λ*_*n*_ determine a unique steady-state density *x*_1_ = *e*_1_, …, *x*_*n*_ = *e*_*n*_ along the mRNA, and for any initial density the dynamics converges to this profile.

Ref. [32] derived a useful *spectral representation* for the mapping from the rates *λ*_0_, …, *λ*_*n*_ to the steady state *e*. Given the RFM, consider the (*n* + 2) *×* (*n* + 2) tri-diagonal matrix

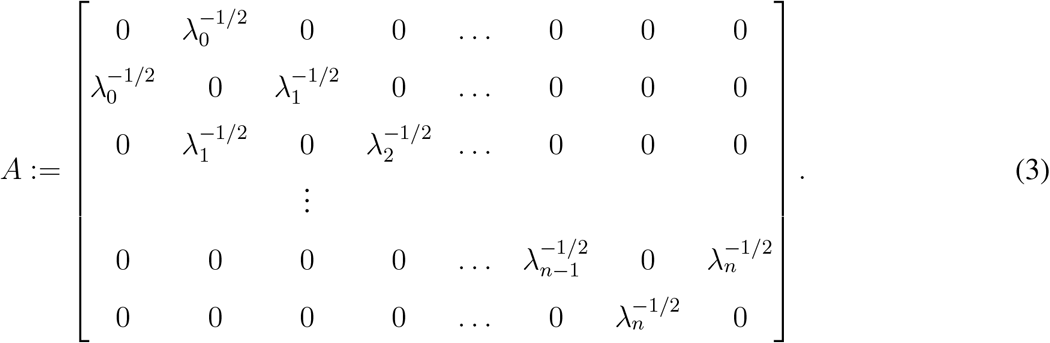

Since *A* is symmetric, all its eigenvalues are real. Since *A* is an irreducible matrix, with all entries non-negative, the Perron-Frobenius theorem [18] implies that *A* admits a simple maximal eigenvalue *σ >* 0, and the corresponding eigenvector *ζ* ∈ ℝ^*n*+2^ is unique (up to scaling) and satisfies *ζ*_*i*_ *>* 0 for all *i* ∈ {1, …, *n* + 2}. Then, the entries of *e* satisfy [5]:

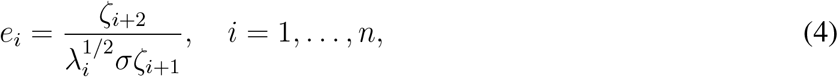

and the steady-state production rate satisfies

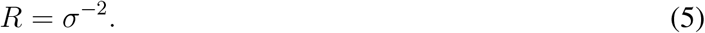

In other words, the Perron eigenvalue and eigenvector of *A* provide all the information needed to determine the steady state profile *e*, and the steady state production rate *R* in the RFM.

This spectral representation has several implications. For example, it implies that it is possible to numerically calculate efficiently the steady state even for very large RFMs using algorithms for computing the Perron eigenvalue and eigenvector of tri-diagonal matrices. Also, it follows from (5) that the function *R* = *R*(*λ*_0_, …, *λ*_*n*_) is strictly concave [32], thus allowing to show that general steady-state protein production optimization problems are convex optimization problems [32]. It also reduces the sensitivity analysis of *R* with respect to any rate *λ*_*i*_ to an eigenvalue sensitivity problem for the matrix *A* [33].

### 2) Ribosome flow model with input and output (RFMIO)

As noted above, a cell typically includes a large number of mRNA molecules and ribosomes, that compete for shared resources, and studying translation on a single, isolated mRNA may thus provide limited insight on large-sale translation in the cell. To model translation in the cell requires a network of interconnected RFMs. The first step in building such a network is adding an input and output to the RFM. This yields the *ribosome flow model with input and output (RFMIO)* [35]. The RFMIO dynamics is

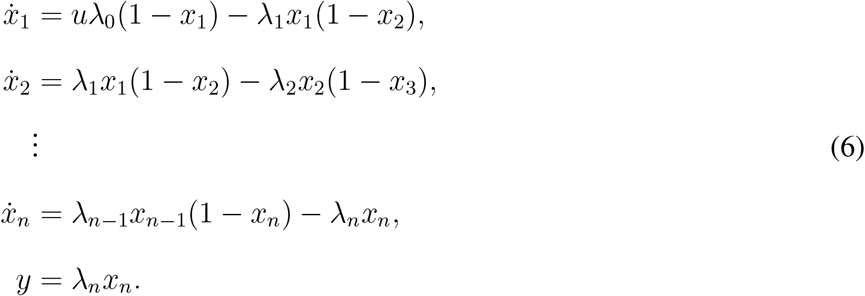

The scalar input *u* : ℝ_+_ → ℝ_+_ represents the density of ribosomes in the vicinity of the initiation site. Thus, if *u*(*t*) is large then the effective initiation rate at time *t*, given by *u*(*t*)*λ*_0_, increases. The scalar output *y*(*t*) = *λ*_*n*_*x*_*n*_(*t*) is the rate of ribosomes exiting the mRNA at time *t*. The additional input and output allow to connect RFMIOs in a network. Note that for *u*(*t*) *≡* 1, Eq. (6) reduces to the RFM.

### 3) The network

Ref. [35] introduced a model composed of *m* RFMIOs interconnected via a pool of free ribosomes (see Fig. 2). Let *n*_*i*_, *i* = 1, …, *m*, denote the length of RFMIO #*i*. The state-variables in RFMIO #*i* are denoted by 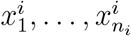. The density in the pool at time *t* is modeled by the scalar function *z*(*t*). The ribosomes that initiate translation in RFMIO #*i* are supplied from the pool through the pool output function *G*_*i*_(*z*(*t*)). Thus, the input to RFMIO #*i* is *u*_*i*_(*t*) = *G*_*i*_(*z*(*t*)), so the effective initiation rate in RFMIO #*i* is 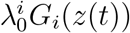.

**Fig. 2:**
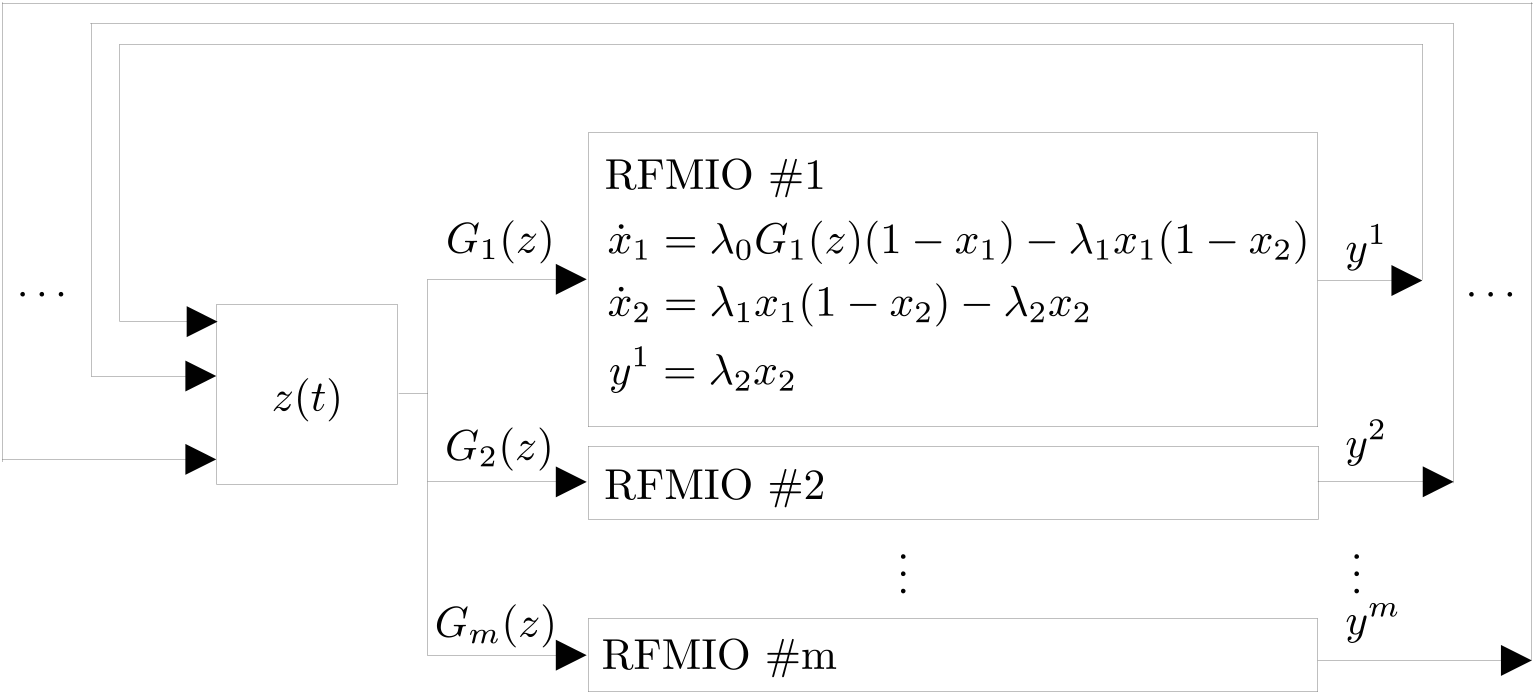
The network includes *m* RFMIOs interconnected via a pool of free ribosomes. For illustration only, we assume that RFMIO #1 has length *n*_1_ = 2 and write its equations explicitly.

The functions *G*_*i*_ : ℝ_+_ → ℝ_+_ satisfy *G*_*i*_(0) = 0 (i.e., when the pool is empty the initiation rate in the RFMIO is zero), and *G*_*i*_(*z*) is continuous and strictly increasing in *z* (i.e., an increase in the pool density yields an increase in the initiation rates). Many possible functions satisfy these constraints, e.g., *G*_*i*_(*z*) = *cz*, with *c >* 0, and the uniformly bounded function *G*_*i*_(*z*) = *α* tanh(*βz*), with *α, β >* 0.

The pool feeds all the RFMIOs, and is fed by the ribosomes exiting all the RFMIOs, so the balance equation for the change in *z*(*t*) is

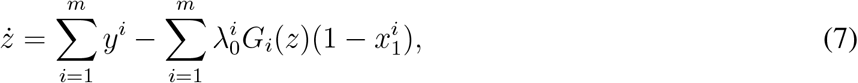

where *y*^*i*^ is the ribosome exit rate from RFMIO #*i*.

Let

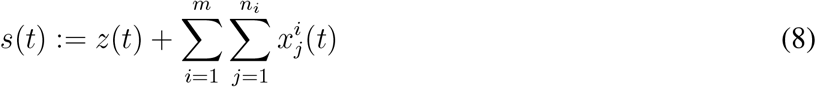

denote the total density of ribosomes in the network at time *t*. Since ribosomes cannot leave nor enter the network,

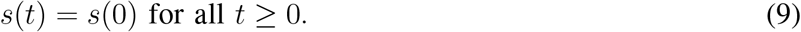

In other words, *s*(*t*) is a first integral of the dynamics.

Ref. [35] used results on cooperative systems with a first integral (see. e.g., [31], [4]) to prove that any solution of the network converges to a steady-state. More precisely, for any *p* ≥ 0, let *L*_*p*_ denote the *p* level set of the first integral, that is, *L*_*p*_ includes all the initial conditions of pool and RFMIO densities with total density *s*(0) = *p*. Then *L*_*p*_ includes a single equilibrium point and any trajectory emanating from an initial condition in *L*_*p*_ converges to this equilibrium point. In other words, any two initial conditions of the network with the same total ribosome density will converge to the same equilibrium state. At this state, the total density is distributed along the mRNAs and the pool such that the flow into and out of each site are equal.

Let *e*_*z*_ ∈ [0, *s*(0)] denote the steady-state pool density, and let 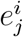 denote the steady-state density in site *j* in RFMIO #*i*. Also, let

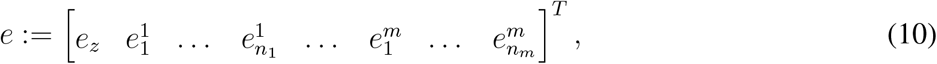

i.e., *e* collects all the steady state values in the network.

## III. Simulation Results

We begin with several synthetic simulation results that demonstrate the wealth of biological questions that can be addressed using the network model when allowing the number of RFMIOs *m* to vary, that is, when mRNAs are added or removed from the network. This also illustrates our general analysis approach that combines the spectral representation of the steady state in every RFMIO with the equation describing the first integral of the network.

We begin by considering a network that includes *m* identical RFMIOs, where each RFMIO has length *n*_*i*_ = 2, *i* = 1, …, *m*. We also assume that every RFMIO is homogeneous, with *λ*_0_ = *λ*_1_ = *λ*_2_ = 1. (Note, however, that all the theoretical results in Section IV below hold for general lengths and rates.) We also assume that *G*_*i*_(*z*) = *z* for all *i* (i.e., the effective initiation rate is proportional to the number of free ribosomes in the pool).

To apply the spectral approach to each RFMIO in the network, let

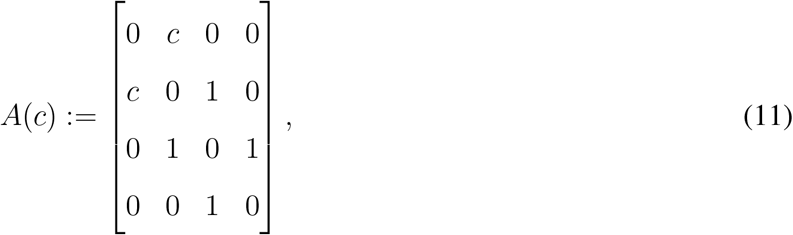

where 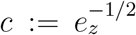. Note that this is exactly the matrix (3) with *n* = 2, *λ*_0_*G*(*e*_*z*_) = *e*_*z*_, and *λ*_*i*_ = 1 for *i* = 1, 2. The Perron root of *A*(*c*) is

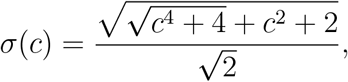

and the corresponding Perron eigenvector is

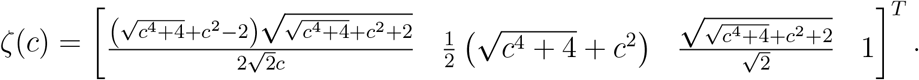

It follows from (4) that the steady-state densities in every RFMIO are

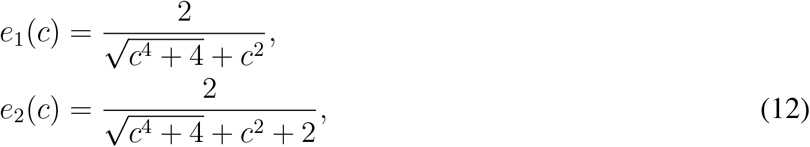

and since *λ*_2_ = 1, the steady-state production rate of each RFMIO is

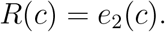

The equation for the total density of ribosomes *s* in the network is

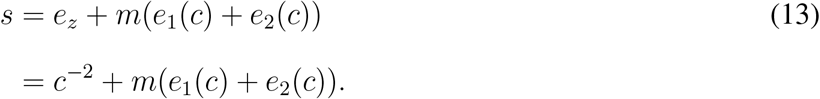

Combining this with (12) provides an explicit expression for *s* as a function of *c*. This can be inverted (at least numerically) to conclude for every total density *s* the corresponding *c* (and thus *e*_*z*_), and then the spectral approach allows to obtain all the steady state profiles in all the RFMIOs.

The network allows to study how important steady state quantities depend on the number of RFMIOs in the network. We first define several such quantities. The ratio between the density of ribosomes in the pool and the total density of ribosomes in the network is

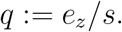

The total protein production at steady state, denoted TPR, is the production rate of all the RFMIOs in the network. Since there are *m* identical RFMIOs,

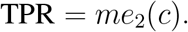

As *m* is increased, more ribosomes are attached to mRNAs and thus we can expect the steady state pool density *e*_*z*_ to go to zero. Then the initiation rate *λ*_0_*G*(*e*_*z*_) = *e*_*z*_ in each RFMIO becomes the bottleneck rate, and thus *e*_*i*_ ≈ *e*_*z*_ for *i* = 1, 2, in every RFMIO. Substituting this in (13) gives *e*_*z*_ ≈ *s/*(1 + 2*m*), and the total production rate is then

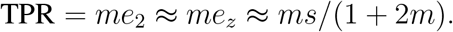

Thus, for a large *m* we expect the total steady state production rate in the network to converge to *s/*2.

Figure 3 depicts the exact network steady state values for *s* = 50 as a function of the number of RFMIOs *m*. It may be seen that: (1) the steady state densities *e*_1_ and *e*_2_ in every RFMIO decrease monotonically with *m*. The same holds for *q* = *e*_*z*_*/s*; and (2) the total production rate *me*_2_ increases monotonically with *m*, and converges to a saturation value of *s/*2 = 25.

**Fig. 3:**
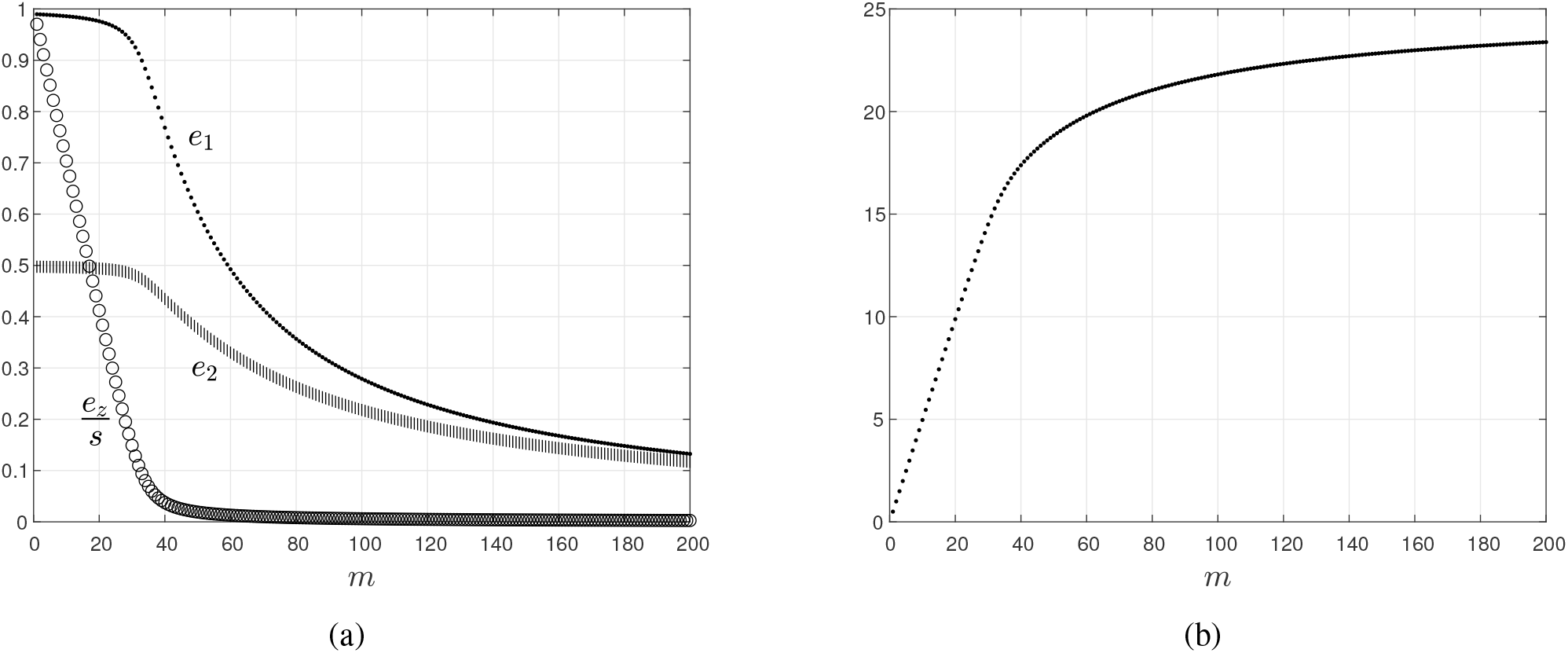
(a) steady-states *e*_1_, *e*_2_ and *e*_*z*_*/s* as a function of the number of RFMIOs *m* when *s* = 50; (b) total production rate *me*_2_ in the network as a function of the number of RFMIOs *m*.

Summarizing, the addition of mRNAs increases the total protein production, as ribosomes now also translate the new mRNAs. However, it decreases the translation rate of the other mRNAs by depleting the pool of free ribosomes. As the number of mRNAs increases, the marginal utility in adding another mRNA decreases to zero. The total production rate in the network is bounded, and an important factor in this bound is the total density of ribosomes in the network.

These simulation results suggest that when optimizing the production rate of a synthetic system (e.g., in high-throughput heterologous gene expression) it nay be useful to use mRNA levels below the regime where the marginal utility in adding another mRNA become negligible.

The next section provides a rigorous treatment of these topics.

## IV. Main results

We begin by considering the effect of adding an RFMIO to the network. We introduce some notation. Recall that RFMIO #*i* is characterized by the tuple

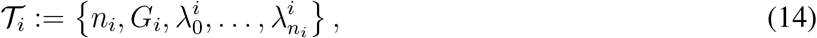

where *n*_*i*_ is the length of RFMIO #*i, G*_*i*_ : ℝ_+_ → ℝ_+_ is the *i*th pool output function, and 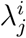 are the rates along RFMIO #*i*.

Consider a network with *m* + 1 RFMIOs obtained by adding an RFMIO to a network of *m* RFMIOs. Let *e*_*z*_(*m*) [*e*_*z*_(*m* + 1)] denote the pool density in the network with *m* [*m* + 1] RFMIOs. What is the relation between the steady state pool densities before and after adding RFMIO #(*m* + 1)? The next result shows that *e*_*z*_(*m* + 1) is always smaller than *e*_*z*_(*m*).

### Proposition 1.

*Fix s >* 0, *and a collection of RFMIOs* 𝒯_1_, 𝒯_2_, …. *For any m, consider a network of m RFMIOs* 𝒯_1_, …, 𝒯_*m*_ *interconnected via a pool of free ribosomes, with total ribosome density s. Let e*(*m*) *denote the corresponding network steady-state (see* (10)*), where the coordinates depend on m. Then*

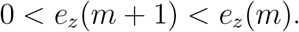

The proof is placed in the Appendix.

In other words, adding an RFMIO to the network, while keeping the total ribosome density constant, always decreases the steady state pool density. This makes sense, as the new mRNA “consumes” ribosomes from the pool.

Proposition 1 implies in particular that in a network built by repeatedly adding new RFMIOs the sequence of steady-state pool densities *e*_*z*_(1), *e*_*z*_(2), … is monotonically decreasing. Since *e*_*z*_(*m*) ≥ 0 for all *m*, this implies that the limit

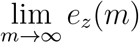

exists. The next result shows that this limit is zero. Since we take *m* → *∞*, we need to impose some technical conditions on the RFMIOs.

### Assumption 1.

*From here on we always assume that the following properties hold*.

1. *There exists λ*_*∗*_ *>* 0 *such that*

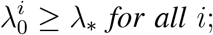

*i*.*e*., *all the initiation rates are bounded from below by λ*_*∗*_;
2. *There exists λ*^*∗*^ *>* 0 *such that*

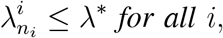

*i*.*e*., *all the exit rates are bounded from above by λ*^*∗*^;
3. *There exist p >* 0 *and g*_*∗*_ *>* 0 *such that for any z* ∈ [0, *p*], *we have*

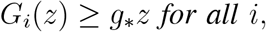

*i*.*e*., *all the pool output functions G*_*i*_(*z*) *are bounded from below by the linear function g*_*∗*_*z on the interval* [0, *p*].

These three conditions are clearly reasonable.

### Proposition 2.

*Fix s >* 0, *and a collection of RFMIOs* 𝒯_1_, 𝒯_2_, …. *For any m, consider a network of m RFMIOs* 𝒯_1_, …, 𝒯_*m*_ *interconnected via a pool of free ribosomes, with total ribosome density s. Let e*(*m*) *denote the corresponding network steady-state (see* (10)*), with coordinates that depend on m. Then*

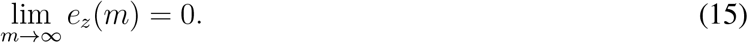

The proof is placed in the Appendix.

From a biological point of view, the case *m* → ∞ may seem unreasonable. However, Proposition 2 implies that for a large *m, e*_*z*_(*m*) will be small. This scenario is the focus of this paper. Indeed, when the pool includes a large number of free ribosomes there is little competition, and thus the indirect coupling between the mRNAs is weak. The interesting case is thus when the pool is close to being depleted, that is, when *e*_*z*_ is small. In the biological context, this may represent a cell under stress conditions, or under a high yield viral infection, or when there are many ribosomal “traffic jams” along the mRNAs, so the pool is depleted.

The next result uses the spectral representation to analyze the steady state densities and production rate in every RFMIO when the pool is starved.

### Proposition 3.

*Consider a network of RFMIOs* 𝒯_1_, 𝒯_2_, …, *interconnected via a pool. Suppose that either the total ribosome density s goes to zero, or that the number of RFMIOs tends to infinity, so that e*_*z*_ → 0.

*Then for any i, RFMIO* #*i satisifes*

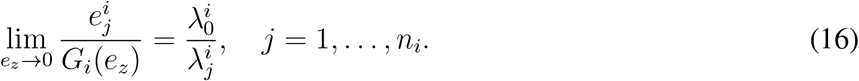

*In particular, if the pool output function G*_*i*_ *is differentiable at zero, then*

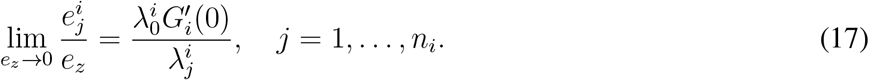

The proof is placed in the Appendix.

In other words, when the pool becomes depleted (e.g., because *m* is large or the total ribosome density *s* is small) every density along the *i*th mRNA behaves asymptotically like the pool output function *G*_*i*_(*e*_*z*_). This makes sense, as the effective initiation rate 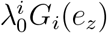 becomes the bottleneck rate in the mRNA. Note that (16) implies that 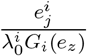 is inversely proportional to 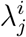. This is reasonable, as 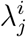 controls the flow *out* of site *j*.

### Example 1.

*To illustrate Proposition 3, we use a network consisting of m identical RFMIOs. Each RFMIO has length* 5 *and rates*

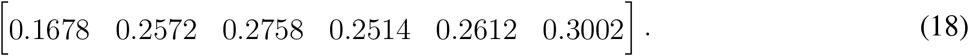

*These values are taken from [47] and correspond to the S. cerevisiae gene YBL025W that encodes the protein RRN10, which is related to regulation of RNA polymerase I. This gene has 145 codons (excluding the stop codon), and was divided into 6 consecutive groups of codons: the first group includes the first 24 codons (that are also related to later stages of initiation). The other groups include 25 non-overlapping codons each, except for the last one that includes 21 codons. This partitioning was found to optimize the correlation between the RFM prediction and biological data measurements (see [47] for more details). We increased m from* 1 *to* 1000 *while keeping the total ribosome density fixed at s* = 20, *thus depleting the steady-state pool density. The input functions are G*_*i*_(*x*) = *x for all i. Since all the RFMIOs are identical, it is sufficient to consider the steady state density in a single RFMIO. Figure 4 depicts e*_*j*_*/e*_*z*_, *j* = 1, …, 5, *as a function of* (1*/e*_*z*_). *It may be seen that as e*_*z*_ *decreases, every ratio e*_*j*_*/e*_*z*_ *converges to the asymptotic value given in Proposition 3*.

**Fig. 4:**
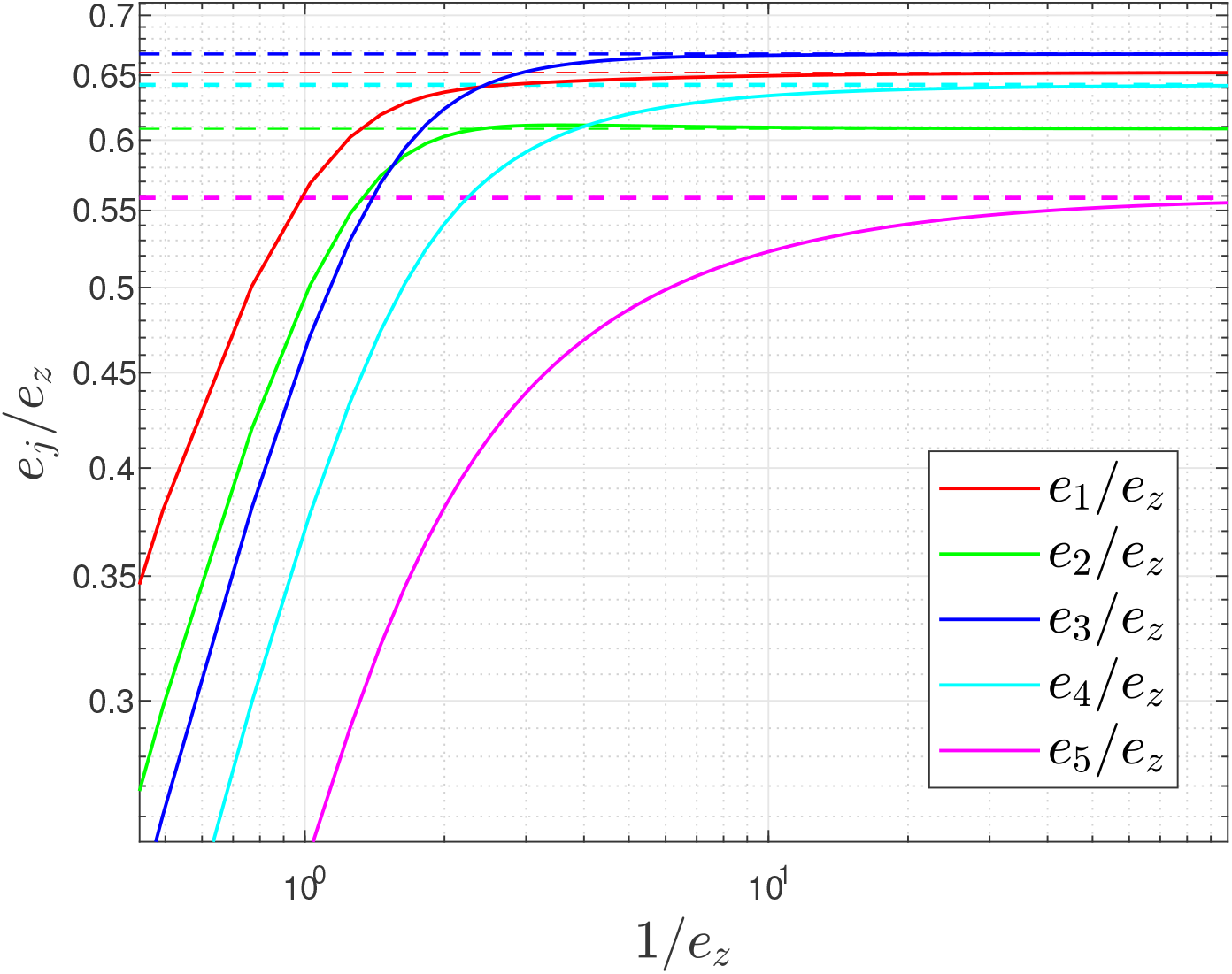
Ratios *e*_*j*_*/e*_*z*_ for *j* = 1, …, 5, as a function of 1*/e*_*z*_. Dashed lines are the asymptotic values in (17).

The following result is an immediate corollary of Proposition 3. Recall that the constant total ribosome density in the network is

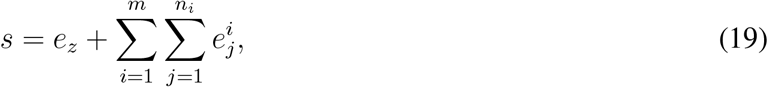

and the total steady state production rate is

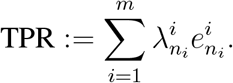

Also, let *q* := *e*_*z*_*/s* denote the ratio between the free ribosomes and the total number of ribosomes in the network.

### Corollary 1.

*Suppose that the pool output functions G*_*i*_ *are differentiable at zero for all i* = 1, 2, …. *Then*,

1. *The total production rate satisfies*

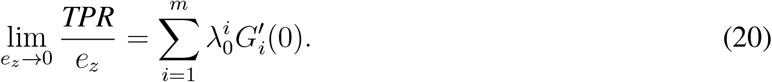
2. *The ratio between the free ribosomes to the total number of ribosomes in the network satisfies*

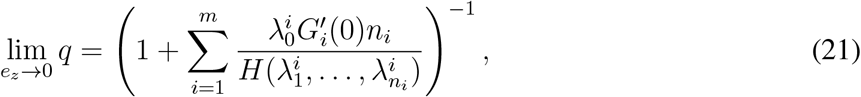

*where* 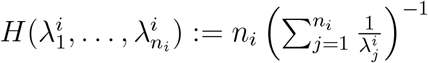 *is the harmonic mean of the rates* 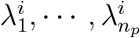, *that is, all the rates except for the initiaion rate*.

The proof is placed in the Appendix.

These results provide closed-form asymptotic expressions for important biological quantities when the pool is starved. Note that as 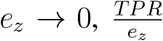 depends on all the initiation rates 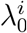, but not on any of the other rates (since the number of ribosome along any mRNA is low, there are no traffic jams). However, the ratio between the density of ribosomes in the pool and the total number of ribosomes in the network does depend on the harmonic mean of all the rates in the network.

We note that a similar closed-form expression can be obtained for the total steady-state density and total production rate on any subset of RFMIOs in the network.

Corollary 1 imlies that when the pool is starved we can replace an entire network of *m* identical RFMIOs by a much simpler network, while keeping the asymptotic steady state properties unchanged.

### Proposition 4.

*Let s >* 0 *denote the total number of ribosomes. Consider the following two networks of RFMIOs:*

1. *A network of m identical RFMIOs, each of length n, rates λ*_0_, …, *λ*_*n*_, *and output functions G*(*z*) = *gz, with g >* 0. *Let TPR [q] denote the total production rate [ratio between free ribosomes and s] in this network*.
2. *A network consisting of a single RFMIO of length* 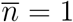, *rates* 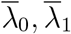, *and* 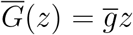, *with* 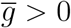. *Let* 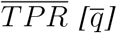 *denote the total production rate [ratio between free ribosomes and s] in this network*. *If the parameters of the second network are chosen such that*

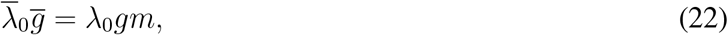

*and*

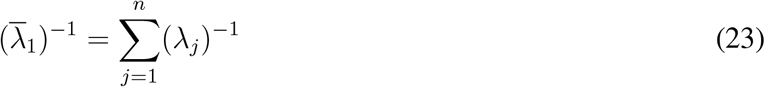

*then, as s* → 0

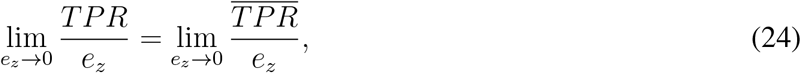

*and*

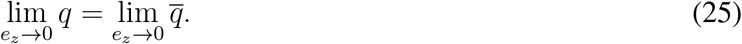

In other words, we can use a network with a pool and a single RFMIO, with a single site, to simulate and analyze the first network. Note that conditions (22) and (23) are quite intuitive. Roughly speaking, the first implies that the initiation rates in the two networks are equal (taking into account that in the first network there are *m* RFMIOs and in the second a single RFMIO), whereas the second condition requires that some mean of the other rates along the RFMIO is also equal.

*Proof*. Using (20), we have 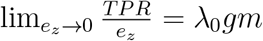 and 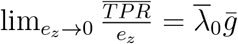, and combining this with (22) gives (24). Similarly, Eq. (21) gives

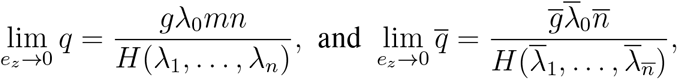

and using (22) and (23) implies that these expressions are equal. □

### Example 2.

*Consider again the network in Example 1. Recall that this has m identical RFMIOs of length n* = 5 *and the rates given in* (18). *In this case, g* = 1, *λ*_0_ = 0.1678, *and*

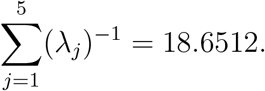

*Proposition 4 implies that we can replace this network of m RFMIOs by a network consisting of a single RFMIO of length one, with* 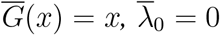.1678*m, and* 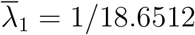, *and the asymptotic behaviour of the two networks when the pool is starved will be identical*.

## V. A biological example: production of insulin in a cell free system

In this section, we apply our model to analyze the production of insulin protein in a cell free system. A cell free system is an in vitro based approach to study and/or generate biological reactions that take place within cells via the isolation of relevant cellular components (e.g. ribosomes, RNA polymerases, tRNA molecules). This approach reduces the complex interactions typically found when working with whole cells. Cell free systems are often used in biotechnology for heterologous gene expression [44].

Data on the coding region of insulin fitted to *S. cerevisiae* was taken from [23]. The codon decoding rates, that are based on the analysis of typical decoding rates in-vivo from ribo-seq data [10], were taken from [11]. Groups of 10 consecutive codons were coarse grained into one RFMIO site, as was done in previous studies (see, e.g., [53]). This yields 12 sites. The transition rate *λ*_*i*_ of site *i* is the inverse of the sum of the decoding times along the 10 related codons. These rates were then normalized such that their average value is 10 codons per second (the typical decoding rate in eukaryotes [8]).^1^

Since we study the translation of insulin in a cell free system, we assume that all the mRNAs in the system encode the protein insulin. Thus, the network includes *m* identical RFMIOs interconnected via a pool.

We set *s* = 250, 000, *m* = 39, 500 (see the data in [46], [24], [53]), and assume that all the pool output functions are identical: *G*_*i*_(*x*) = *cx* for all *i* = 1, …, *m*. To calibrate the constant *c*, we assume that *c* maximizes the effective initiation rate while keeping it lower than all the other transition rates. Mathematically, this yields the equation

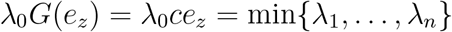

and this gives *c* = 7.4758 · 10^*−*5^.

To study the scenario where the pool is starved, we consider two cases: in the first we fix *m* and decrease *s*, and in the second we fix *s* and increase *m*. From a biological perspective, both cases correspond to the fact that ribosomes may be “more expensive” then mRNAs, and the goal is to optimize production while using a minimal number of ribosomes.

### A. Varying the total density of ribosomes

Consider the case where *m* = 39, 500 is fixed, and *s* varies. Figure 5 depicts TPR*/e*_*z*_ (that is, the ratio between the steady-state total production rate and the steady-state pool density) as a function of *m/s* (that is, the number of mRNA molecules divided by the total number of ribosomes in the network). As expected, TPR*/e*_*z*_ increases with *m/s*, and converges, as *s* → 0, to the asymptotic value 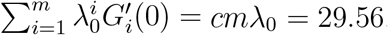.

**Fig. 5:**
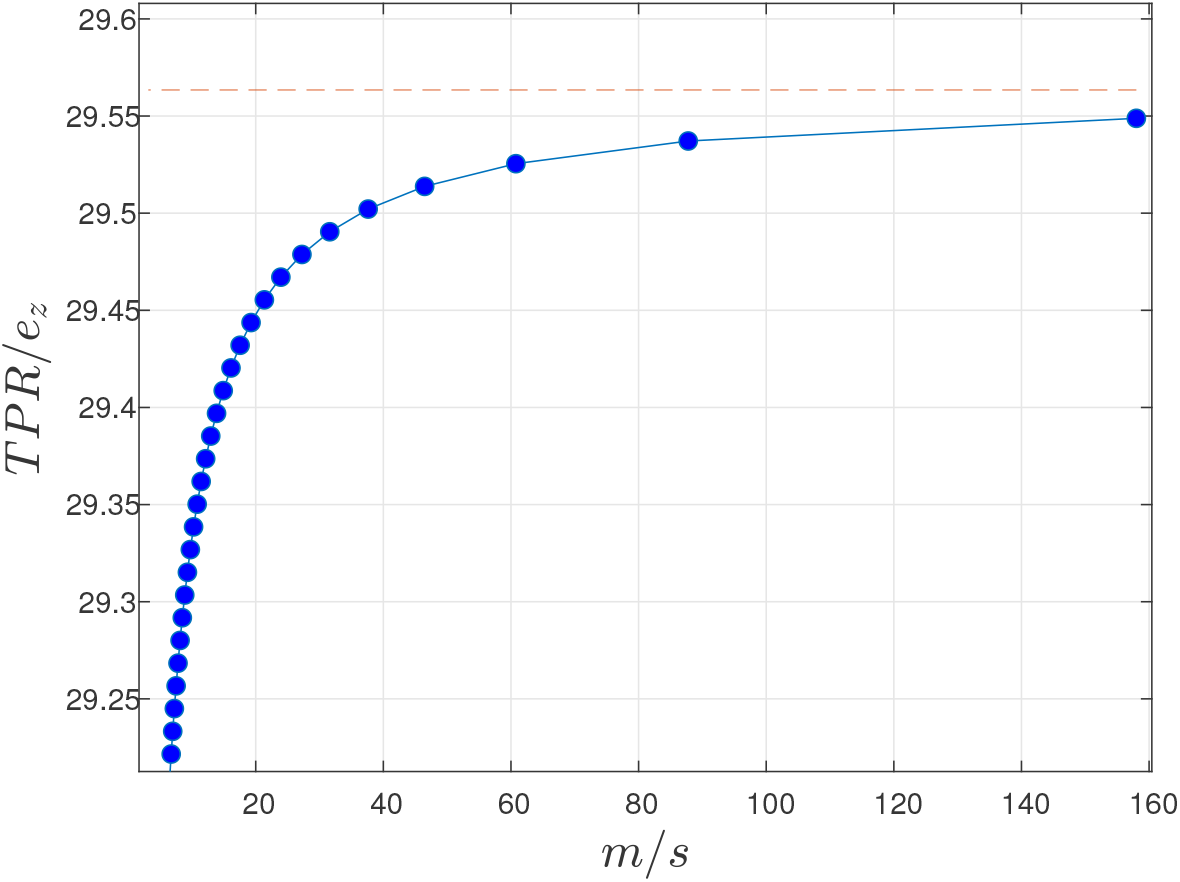
The ratio TPR*/e*_*z*_ as a function of *m/s* for *m* = 39, 500, and *s* varies in the range [250, 12, 500] (blue dots). The predicted asymptotic value 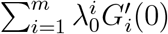 is marked with a dashed red line.

### B. Varying the number of mRNAs

Consider the case where *s* = 250, 000 is fixed, and *m* is varied. Figure 6 depicts the number of free ribosomes *e*_*z*_ as a function of *m/s*. As *m* increases, *e*_*z*_ sharply decreases, as more ribosomes attach to the additional mRNAs, and thus the pool is starved. In particular, *e*_*z*_ goes to zero as *m* → *∞* (see Proposition 2). In practice, when *m/s* = 80 we already get a very low value of *e*_*z*_, and then the asymptotic results described in our analysis can already be used.

**Fig. 6:**
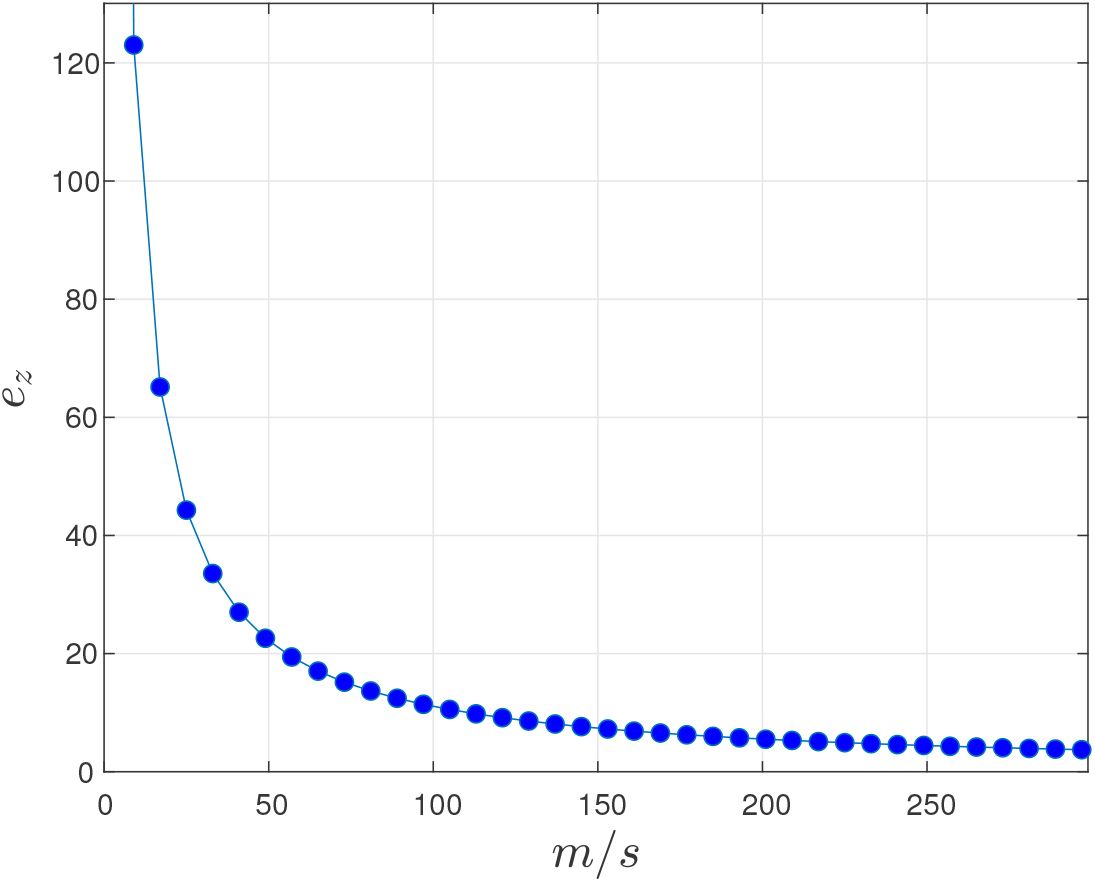
*e*_*z*_ as a function of *m/s* for *s* = 250, 000, and *m* in the range [2.5, 750] · 10^5^.

Note that these results can provide important guidelines for setting the cell free system parameters. For example, Figure 5 shows that to achieve a steady state production rate that is 99% of the maximal possible production rate we should set *m/s* = 10, that is, 10 mRNA molecules for each ribosome in the system.

## VI. Discussion

We considered a network of mRNAs fed by a pool of free ribosomes in the scenario when the pool is starved. This scenario is relevant for example in biological networks under stress conditions or viral attack, and for synthetic networks where the goal is to optimize the total production rate. Furthermore, we believe that studying systems under extreme conditions may provide intuition for how the system was designed and how it will operate even under normal conditions.

We used a mathematical model of a network of RFMIOs connected via a pool of free ribosomes. Using the spectral representation of the RFMIO steady state we derived closed-form expressions for several relevant biological quantities in the regime when the pool is starved. These include the total protein production rate in the network, and the ratio between the number of ribosomes in the pool and the total number of ribosomes in the network.

We demonstrated the analytical results using both synthetic examples and an example based on biological data. The results reported here can be used both in systems biology studies of natural systems and for the design of synthetic systems. We believe that an interesting direction for further research is to use our results in the biological context of a cell attacked by viruses. This will allow to study both qualitatively and quantitatively important questions. For example, can the virus effectively shut down the host protein production (and in particular immune-related proteins) by depleting the pool of free ribosomes or are other mechanisms needed?

## Supporting information

Latex source files

## Appendix: Proofs

For the sake of readability, we begin with a short and general description of our analysis approach. Consider without loss of generality the steady-state densities 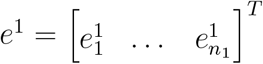 along RFMIO #1 in the network. The spectral representation implies that we can retrieve *e*^1^ from the Perron root and Perron eigenvector of the matrix

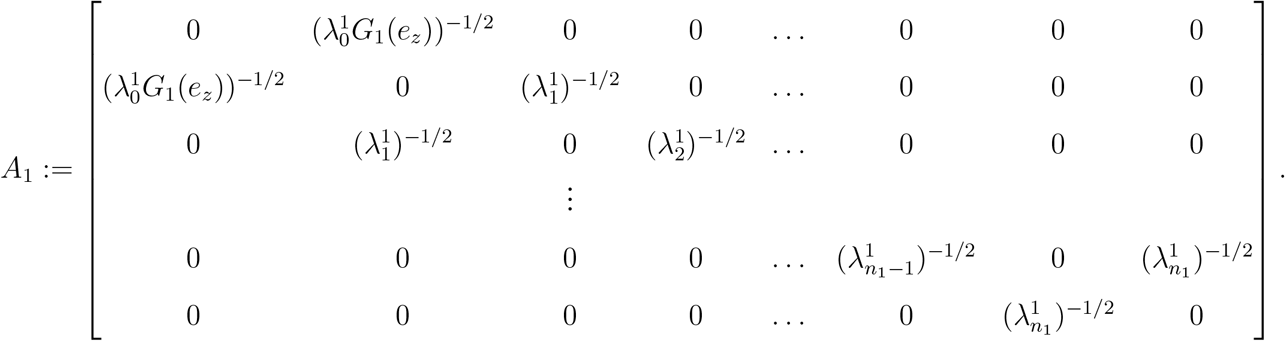

Indeed, at steady state the pool density is *z*(*t*) = *e*_*z*_, so the initiation rate at RFMIO #1 is 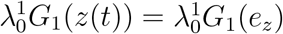.

When *e*_*z*_ is close to zero, so is *G*_1_(*e*_*z*_) and this implies that entries (1, 2) and (2, 1) in *A*_1_ are very large. We use an asymptotic analysis of the spectral properties of *A*_1_ to derive an approximate expression for *e*^1^ and, similarly, for any *e*^*i*^, *i* = 1, …, *m*. Now by (8) and (9),

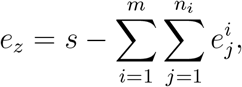

and thus we obtain the entire steady state of the network.

We begin with several auxiliary results that describe the asymptotic spectral properties of a specific tri-diagonal matrix. We use 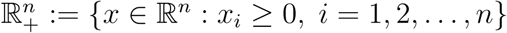 to denote the non-negative orthant in ℝ^*n*^, and 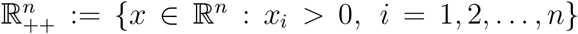 to denote the positive orthant in R^*n*^. For a Hermitian matrix *S* ∈ **C**^*N×N*^ we denote its eigenvalues by

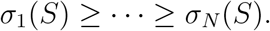

### Theorem 1.

*Given c >* 0 *and a vector α* =[*α*_1_, …, *α*_*N−*2_]^*T*^, *with α*_*i*_ *>* 0, *consider the N×N tri-diagonal and symmetric matrix*

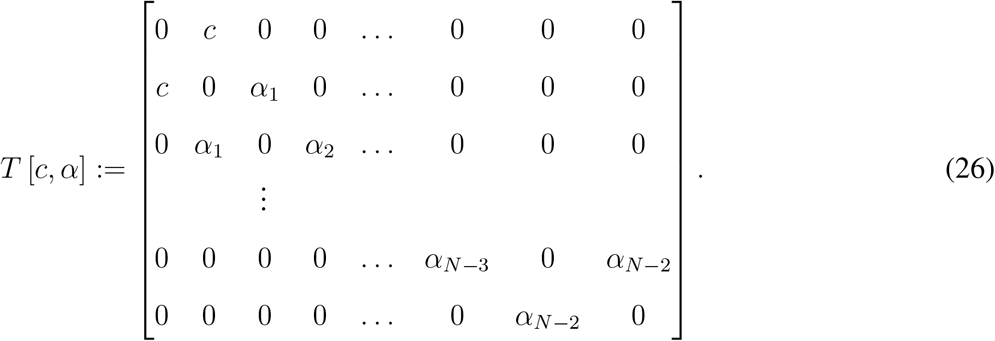

*Let* 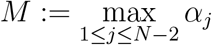. *Then for any c* ≥ 2*M, we have*

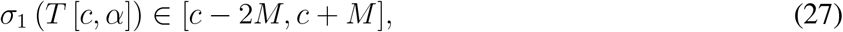

*and*

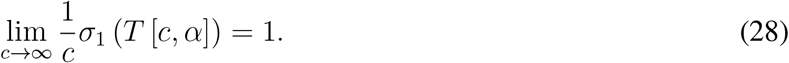

*Furthermore, let* 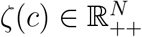 *be the eigenvector of T* [*c, α*] *corresponding to σ*_1_ (*T* [*c, α*]), *normalized such that its first entry is ζ*_1_(*c*) = 1. *Then*

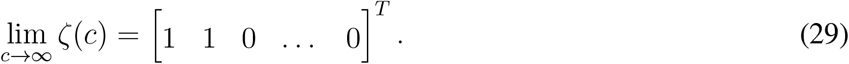

*More precisely, the entries of this vector satisfy*

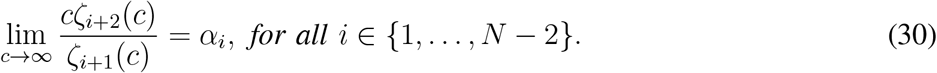

**Proof**. *Recall that a theorem of Weyl [18, Section 4*.*3] asserts that if A, B* ∈ **C**^*N×N*^ *are Hermitian then for any i, j* ∈ {1, …, *N*}, *we have*

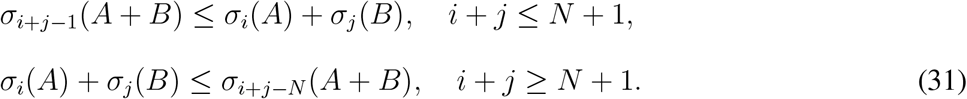

*Recall also that if* | · | : **C**^*N*^ → ℝ_+_ *is a vector norm, and* || · || : **C**^*N×N*^ → ℝ_+_ *is the induced matrix norm, then* |*σ*_*i*_(*S*)| *≤* ||*S*|| *for any i*.

*We can now prove Theorem 1. First note that for any c ≥ M, we have*

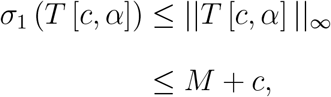

*and this proves the upper bound in* (27).

*To prove the lower bound, fix c >* 0. *Define A* := *T* [*c*, 0, …, 0], *B* := *T* [0, *α*]. *Note that A* + *B* = *T* [*c, α*]. *Applying* (31) *with i* = 1 *and j* = *N gives*

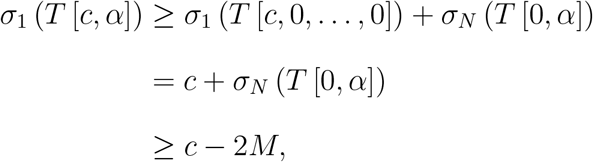

*where the last inequality follows from the fact that*

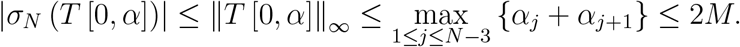

*This completes the proof of* (27). *Taking c* → *∞ in* (27) *proves* (28).

*To prove* (29), *let* 0 *< c*_1_ *< c*_2_ *<* … *be such that* lim_*k*→*∞*_ *c*_*k*_ = *∞. To simplify the notation*,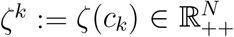. *We may assume that every ζ*^*k*^ *has norm one, and thus we can extract a subsequence*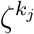, *j* = 1, 2, …, *that converges to a limit vector* 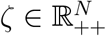, *that also has norm one. Then*

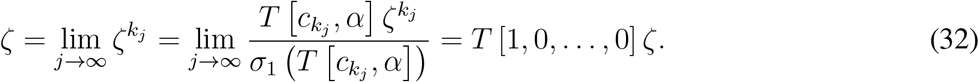

*Thus, ζ is a normalized eigenvector of T* [1, 0, …, 0] *corresponding to σ*_1_ (*T* [1, 0, …, 0]) = 1, *and it is straightforward to verify that*

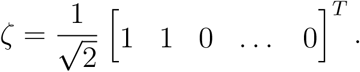

*We next claim that* lim_*k*→*∞*_ *ζ*^*k*^ = *ζ. Indeed, assume the contrary. Then there is a subsequence* 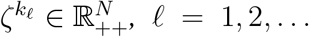, *ℓ* = 1, 2, …, *and ε >* 0 *such that* 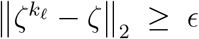 *for all ℓ. Extracting a convergent subsequence yields a normalized eigenvector* 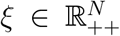 *of T* [1, 0, …, 0] *corresponding to σ*_1_ (*T* [1, 0, …, 0]) = 1, *and satisfying* ∥*ξ − ζ*∥_2_ *≥ ϵ. Hence, σ*_1_ (*T* [1, 0, …, 0]) *is not a simple eigenvalue of T* [1, 0, …, 0]. *This contradiction completes the proof of* (29).

*We now analyze the ratios between consecutive entries of the Perron eigenvector of T* [*c, α*]. *Fix c >* 0. *For simplicity, we will denote T* [*c, α*] *and σ*_1_ (*T* [*c, α*]) *by T* (*c*), *and σ*(*c*), *respectively. Define the N × N diagonal scaling matrix*

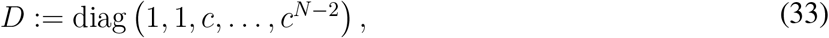

*A direct calculation gives*

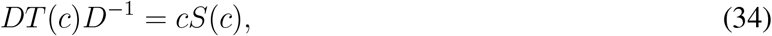

*where*

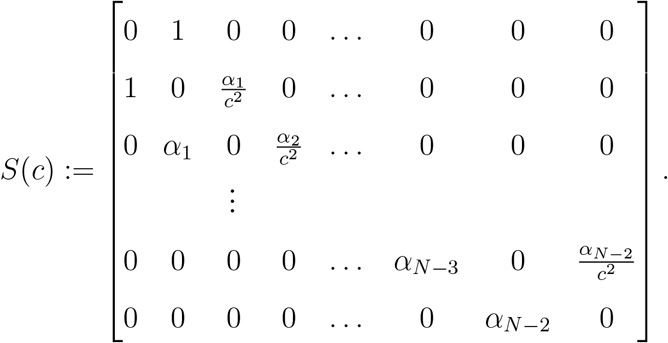

*Let*

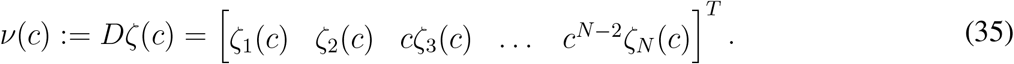

*Then*

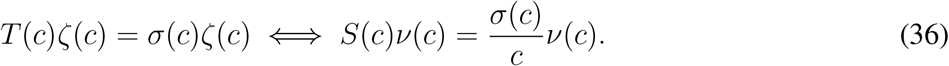

*In particular*

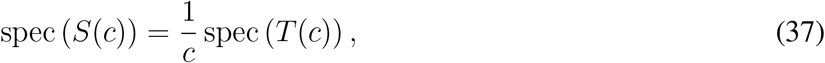

*where* spec(*A*) *is the spectrum of A. Define*

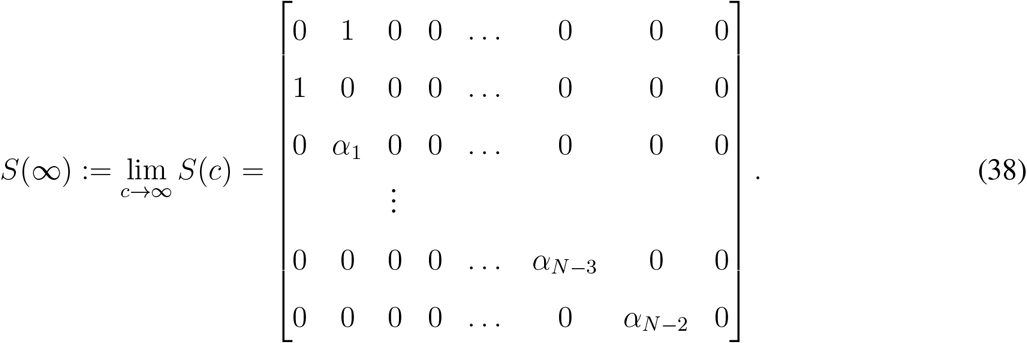

*The characteristic polynomial of S*(*∞*) *is q*(*z*) := det(*zI*_*N*_ *−S*(*∞*)) = *z*^*N−*2^(*z−*1)(*z* +1), *so σ*_1_ (*S*(*∞*)) = 1 *is a simple eigenvalue with corresponding eigenvector ν*(*∞*), *which is given by*

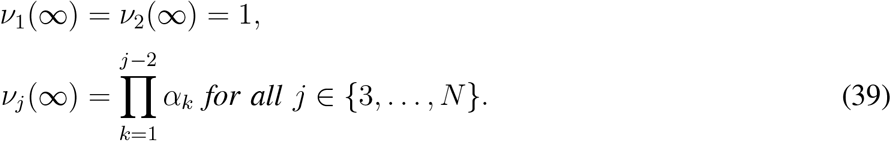

*Using arguments similar to* (32) *and taking the limit c* → *∞ in* (36), *and using the fact that* lim 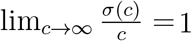 *yields S*(*∞*)*ν*(*∞*) = *ν*(*∞*). *By* (35), *for any i* = 1, …, *N−*2 *we have* 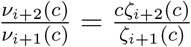. *This proves* (30). ▪

In practice *e*_*z*_ is positive (even if small), so it is useful to derive *explicit* error bounds in the asymptotic expressions. Note that since we are interested in ratios between entries of the Perron vector, we do not necessarily require it to be normalized. For two functions *f, g* : ℝ_+_ → ℝ_+_, we write *f* = O(*g*) if there exist *c, y >* 0 such that

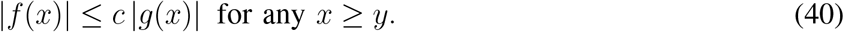

The next result provides more explicit information on the Perron eigenvector of *T* [*c, α*].

### Theorem 2.

*Assume that the assumptions of Theorem 1 hold. Then as c* → *∞, T* [*c, α*] *admits a Perron eigenvector* 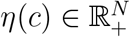 *satisfying*

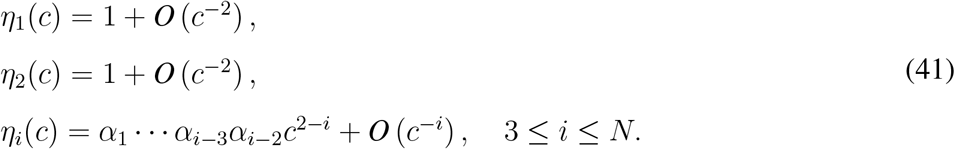

In the proof of this result we actually provide explicit upper bounds for the asymptotic terms in (41) (see Eq. (54) below).

**Proof**. *For a vector q* ∈ ℝ^*N*^, *let sp*(*q*) := {*rq* | *r* ∈ ℝ} *denote the span of q. Define an N × N matrix R by*

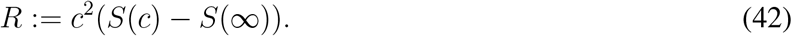

*Note that this implies that R does not depend on c, and that*

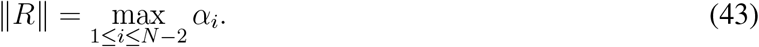

*Let ν*(*c*) *be the Perron eigenvector of S*(*c*) *defined in* (35), *and let*

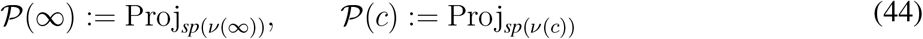

*be the projection operators on sp* (*ν*(*∞*)) *and sp* (*ν*(*c*)), *respectively. Let ξ*(*c*) := *𝒫*(*c*)*ν*(*∞*). *Since we project on sp* (*ν*(*c*)), *this implies that ξ*(*c*) *is a Perron eigenvector of S*(*c*) *corresponding to* 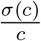. *Let η*(*c*) := *D*^*−*1^*ξ*(*c*). *Then* (36) *implies that η*(*c*) *is a Perron eigenvector of T* (*c*) *corresponding to σ*(*c*). *We will show that η*(*c*) *satisfies* (41).

*We begin by analyzing ξ*(*c*). *Let* Γ *be the circle in the complex plane parametrized by* 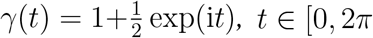 *(see Figure 7)*.

**Fig. 7:**
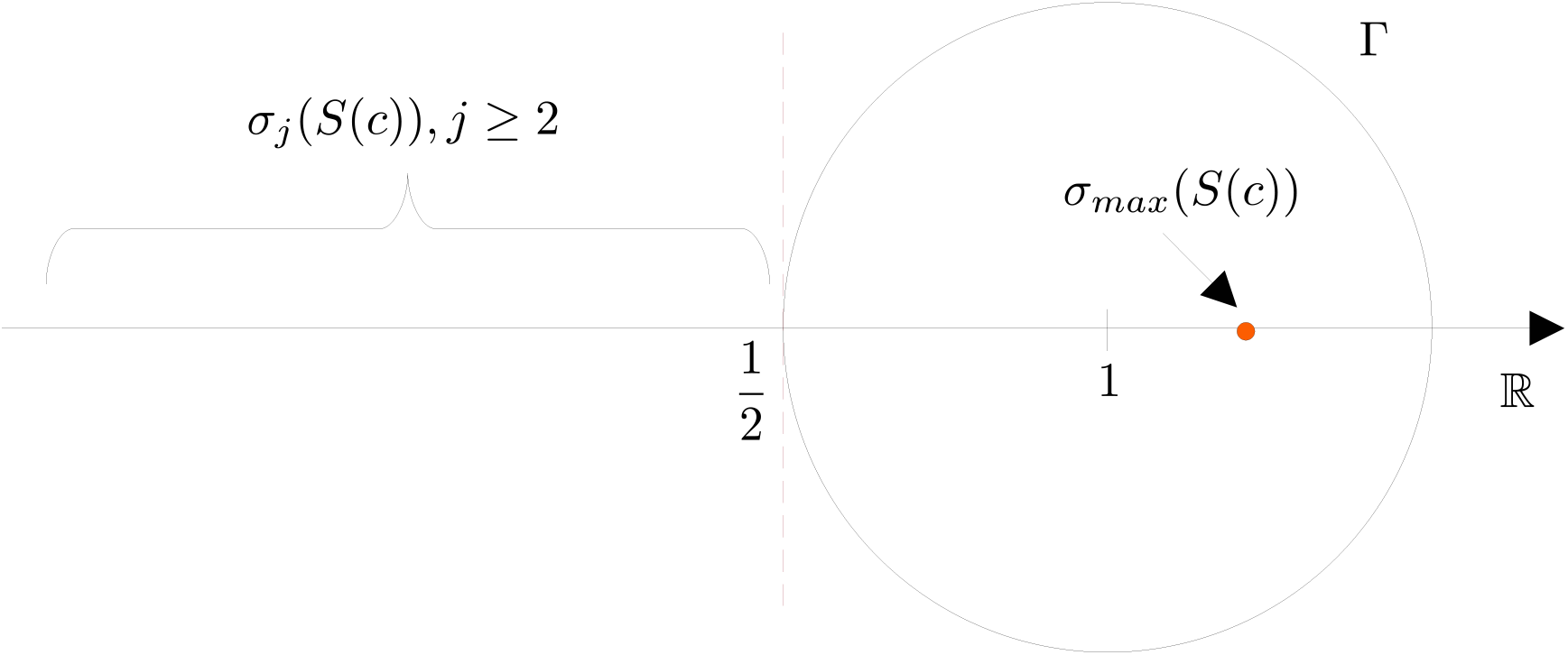
The curve Γ in the complex plane. The eigenvalues of *S*(*∞*) are *−* 1, 0, 1, implying that for any *c* large enough all the eigenvalues of *S*(*c*), except for *σ*_1_(*S*(*c*)), are located outside of Γ. For presentation purposes, *σ*_1_ (*S*(*c*)) is assumed to be on the right of the point [1 0]^*T*^.

### Proposition 5.

*For any c >* 4 *∥R∥ we have that σ*_1_ (*S*(*c*)) *and σ*_1_ (*S*(*∞*)) = 1 *are the only eigenvalues of S*(*c*) *and S*(*∞*), *respectively, located in the interior of* Γ. *All other (real) eigenvalues are to the left of the line* {*s* ∈ ℂ | *ℜ𝔢*(*s*) = 1*/*2}, *where ℜ𝔢*(*s*) *is the real part of s*.

**Proof**. *Recall that* spec (*S*(*∞*)) = {*−*1, 0, 1}. *Hence, the claim is true for S*(*∞*). *For S*(*c*), *consider first the corresponding T* (*c*), *satisfying* (34). *Recall that all eigenvalues of T* (*c*) *are real. By Theorem 1 and* (43), *we have*

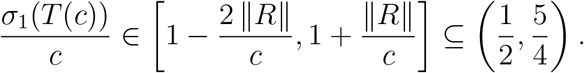

*Using Weyl’s inequalities* (31), *the second largest eigenvalue of T* (*c*) *satisfies σ*_2_(*T* (*c*)) *≤* 2 ∥*R*∥. *The proposition then follows from* (37). ▪

*It is well known [14, Chapter 1] that the projections defined in* (44) *satisfy the matrix representations:*

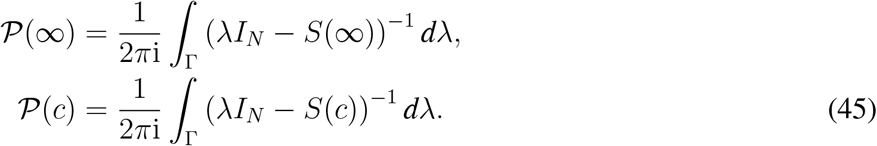

*We require the following auxiliary result that provides a Neumann series representation for the difference between these two projections. The proof is provided for completeness. Let*

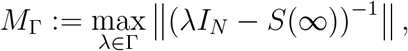

*i*.*e*., *the maximal norm of the resolvent of S*(*∞*) *over* Γ. *This maximum exists since* (*λI*_*N*_ *− S*(*∞*))^*−*1^ *is continuous in a neighborhood of* Γ.

### Lemma 1.

*Fix λ* ∈ Γ. *For any* 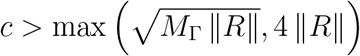, *we have*

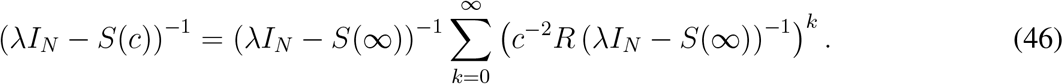

*In particular, the series on the right hand side of* (46) *converges absolutely and uniformly on* Γ, *and*

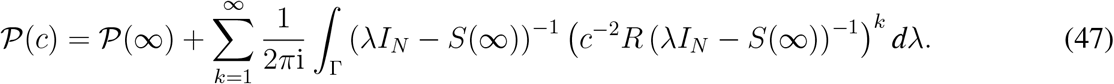

**Proof**. *Fix λ* ∈ Γ. *The definition of R gives* (*λI_N_ − S*(*c*))^*−*1^ = (*λI_N_ − S*(*∞*) *− c*^*−*2^*R*)^*−*1^, *so*

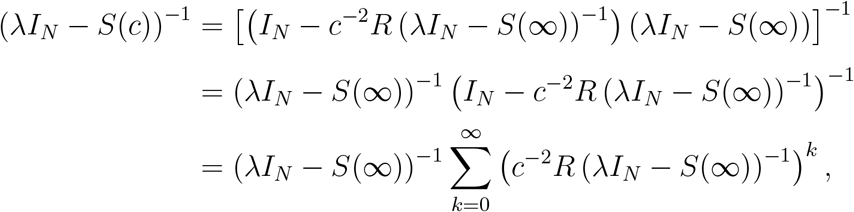

*and the series converges if ||c*^*−*2^*R* (*λI*_*N*_ *− S*(*∞*))^*−*1^|| *<* 1. *We have*

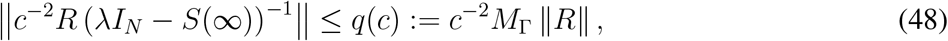

*and q*(*c*) *<* 1 *for any* 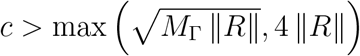. *Moreover for any such c, we have*

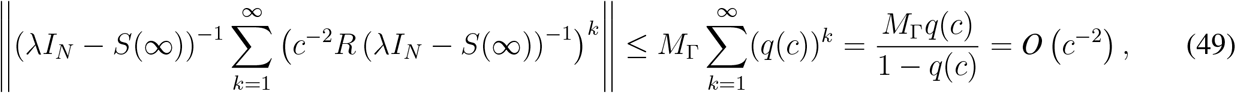

*so the Neumann series converges absolutely and uniformly on* Γ. *This proves* (46). *Combining* (45) *and* (46) *proves* (47), *and this completes the proof of Lemma 1*. ▪

*We can now prove Theorem 2. Define ε*(*c*) := (*𝒫*(*c*) *− 𝒫*(*∞*))*ν*(*∞*). *Then*

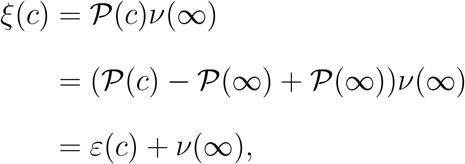

*and using Lemma 1 gives*

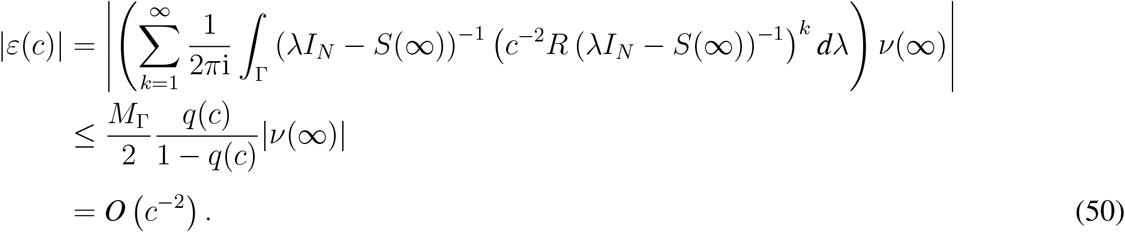

*Now, η*(*c*) = *D*^*−*1^*ξ*(*c*) *gives*

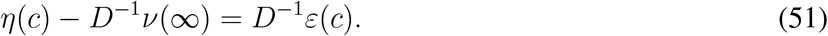

*We conclude that for any* 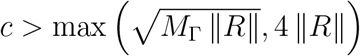, *we have*

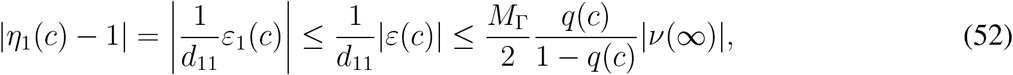

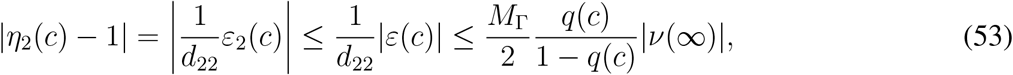

*and for any i* ∈ {3, …, *N*}, *we have*

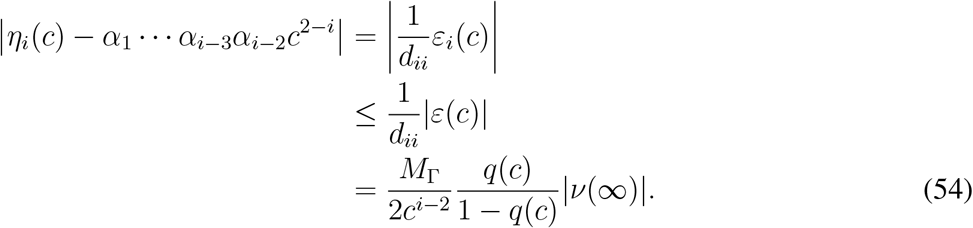

*Note that given* 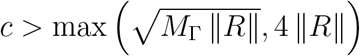, *the upper bounds in* (52), (53) *and* (54) *can be computed* explicitly *using the expressions for the vector ν*(*∞*) *in* (39), *the norm* ∥*R*∥ *in* (43), *and the expression for q*_*c*_ *in* (48). *Combining these upper bounds with* (49) *yields* (41), *and this completes the proof of Theorem 2*. ▪

### Remark 1.

*Note that approximations of the coordinates of η*(*c*) *up to arbitrarily small error can be obtained by computing higher order terms in the Neumann series* (47). *As an example, consider for simplicity the case α*_*i*_ = 1 *for i* = 1, …, *N −* 2, *which implies* ∥*R*∥ = 1. *Assume that* 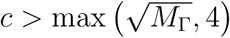 *and let δ >* 0 *be the desired error bound. Let L >* 0 *be a sufficiently large integer such that*

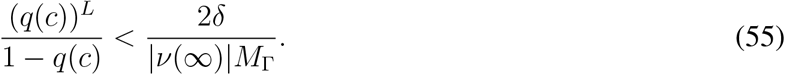

*Then, arguments similar to* (49) *imply that*

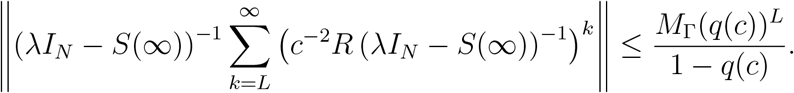

*Hence*, 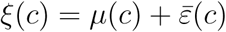, *where*

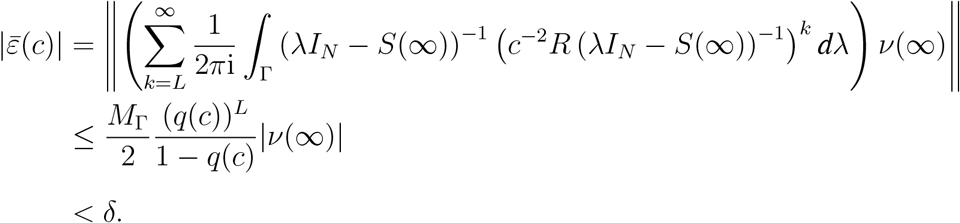

*The vector*

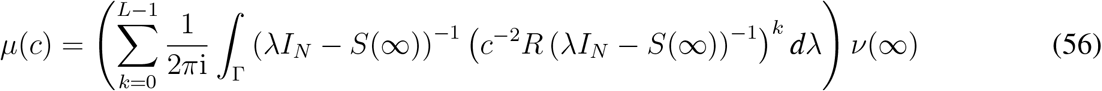

*can be computed explicitly by the Cauchy residue theorem [42]. Using the fact that η*(*c*) = *D*^*−*1^*ξ*(*c*) *yields*

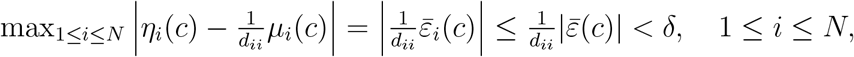

*so the entries of D*^*−*1^*μ*(*c*) *approximate those of η*(*c*) *with an error smaller than δ*.

The next result provides an explicit expression for (*λI*_*N*_ *− S*(*∞*))^*−*1^ with *λ* ∈ Γ. This can be used to derive an upper bound on the constant *M*_Γ_, given in (48), thereby leading to an explicit estimate of the errors in (54). Moreover, (*λI*_*N*_ *− S*(*∞*))^*−*1^ can be substituted into (56) to obtain high order approximations of the Perron eigenvector.

### Proposition 6.

*Let* Γ *be as in Figure 7. Then, for any λ* ∈ Γ

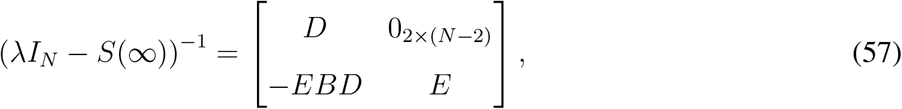

*where*

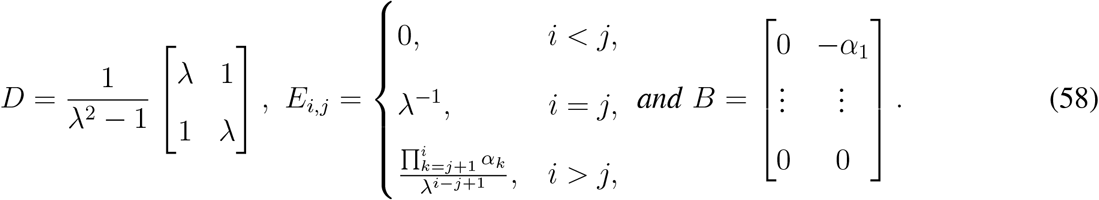

*Proof*. We can divide *λI*_*n*_ *− S*(*∞*) into blocks as

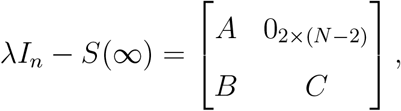

with

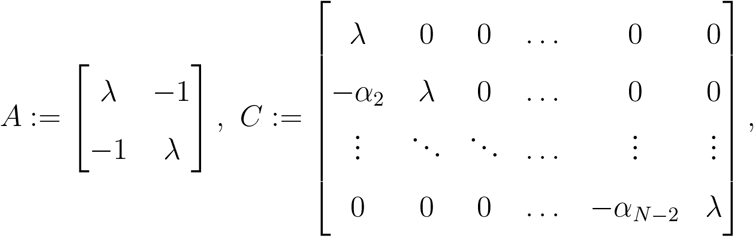

and *B* defined in (58). Since *λ* ≠0, it can be easily verified that *C*^*−*1^ = *E* and *A*^*−*1^ = *D*. Hence,

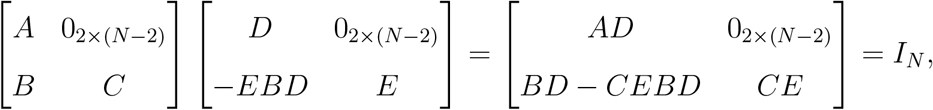

and this completes the proof. □

We can now prove the results in Section IV.

### A. Proof of Proposition 1

It was shown in [35] that for any *s >* 0 the network satisfies the following persistence property for any initial condition. After an arbitrarily short time the pool density is positive, and all the densities in the sites are larger than zero and smaller than one. In particular, this implies that

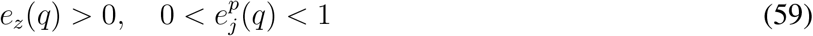

for any integer *q >* 0, any *p* ∈ {1, …, *q*}, and any *j* ∈ {1, …, *n*_*p*_}.

Seeking a contradiction, assume that the claim in Proposition 1 is not true. Then there exists an integer *m >* 0 such that

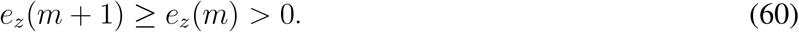

Let

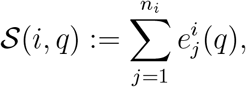

i.e., the total steady-state density of ribosomes in RFMIO #*i* when the network includes *q* RFMIOs. By (59), we have *S*(*m* + 1, *m* + 1) *>* 0. Taking into account (60) and the fact that the total number of ribosomes is fixed at *s*, we conclude that there exists some 1 *≤ p ≤ m* such that

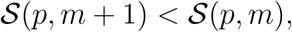

i.e., the total steady-state density of ribosomes along RFMIO #*p* has decreased due to the addition of RFMIO #(*m* + 1) to the network. In particular, for at least one site *j* ∈ {1, …, *n*_*p*_}, we have

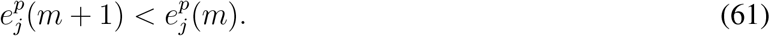

Consider the steady state equations of RFMIO #*p* when the network has *q* RFMIOs, namely,

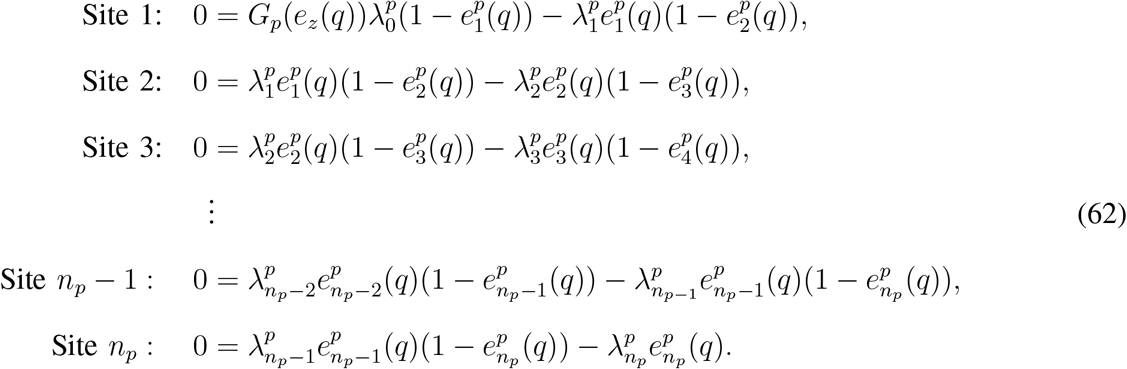

Define a function *ψ* : (0, 1)^2^ → ℝ by 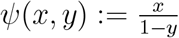, and note that

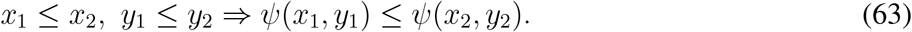

Eq. (62) with *q* = *m* + 1 yields

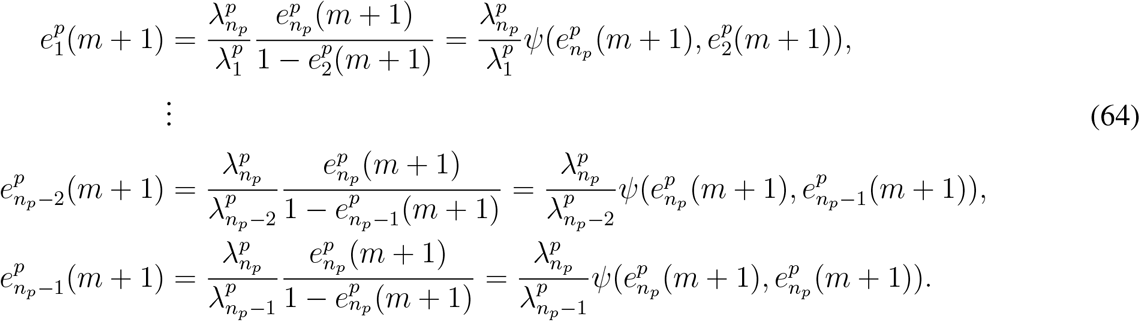

All these expressions are well-defined by (59). We now consider two cases.

*Case 1*. Suppose that

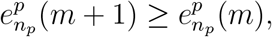

i.e., the density in the last site of RFMIO #*p* did not decrease due to the addition of RFMIO #(*m* + 1).

By backward induction in (64) and using (63), we immediately obtain that

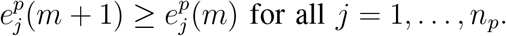

This contradicts (61).

*Case 2*. Suppose that

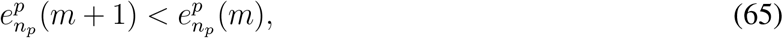

i.e., the density in the last site of RFMIO #*p* decreased due to the addition of RFMIO #(*m* + 1). By backward induction and (63), we find that

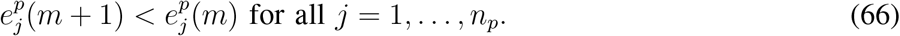

Now using (62) with *q* = *m* and *q* = *m* + 1 gives

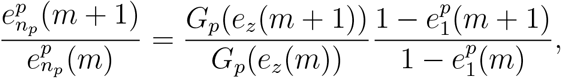

and using (60), the monotonicity of *G*_p_, and (66) yields 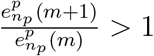. This contradicts (65).

Summarizing, we see that (60) cannot hold, and this completes the proof.

### B. Proof of Proposition 2

Seeking a contradiction, assume that the claim is not true. Then there exists *β >* 0 and a sequence *m*_1_ *< m*_2_ *<* … such that *e*_*z*_(*m*_*k*_) *≥ β* for all *k*. For any *k* and any *i* = 1, …, *m*_*k*_, let

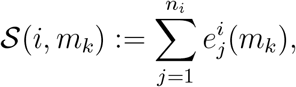

i.e., the total density of ribosomes along RFMIO #*i* at steady-state when the network includes *m*_*k*_ RFMIOs. Since the total number of ribosomes in the network is *s* (and this is independent of *m*_*k*_), we have

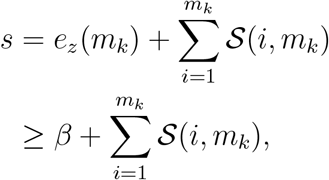

so

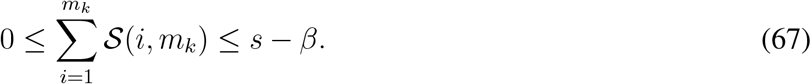

Note that the bound on the right-hand side of (67) does not depend on *m*_*k*_. Let

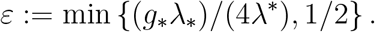

Note that *ε* ∈ (0, 1*/*2]. If for any *m*_*k*_ and any *i* ∈ {1, …, *m*_*k*_} we have that *S*(*i, m*_*k*_) *> ε* then for a large enough *k* this contradicts (67). Therefore, there exist *k* and *p* ∈ {1, …, *m*_*k*_} such that *S*(*p, m*_*k*_) *≤ ε*. In particular,

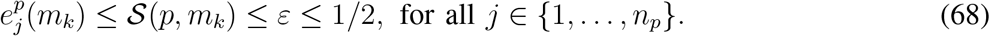

Consider the network with *m*_*k*_ RFMIOs initialized at the equilibrium *e*(*m*_*k*_) at time *t* = 0. Then for any time *t ≥* 0, we have

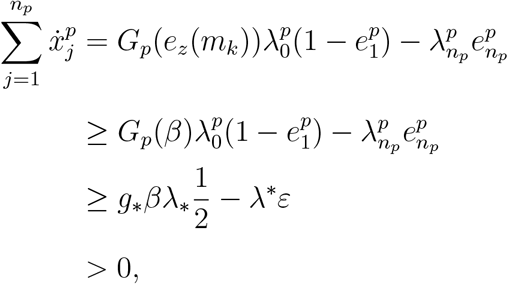

where the second line follows from the monotonicity of the pool output functions, the third from the assumptions in the statement of the proposition and (68), and the fourth from the definition of *ε*. However, since the system is initialized at the steady state, 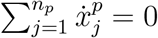 for all *t ≥* 0. This contradiction completes the proof.

### C. Proof of Proposition 3

Fix *i* ∈ {1, …, *m*}, and consider RFMIO #*i*. Setting

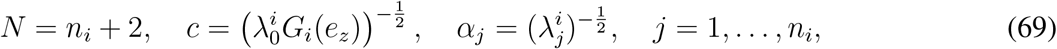

in (26), we have that *T* [*c, α*] is the matrix whose Perron eigenvalue *σ*^*i*^ and eigenvector *ζ*^*i*^ are used to compute the steady state in RFMIO #*i* via (4), that is,

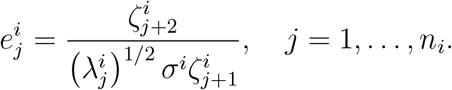

When *e*_*z*_ → 0, *c* → *∞*, and applying (28) and (30) yields (16). If the pool output function *G*_*i*_ is differentiable at zero, then

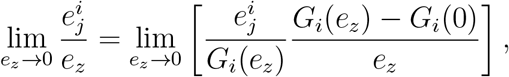

where we used the fact that *G*_*i*_(0) = 0. Equation (17) now follows from (16). This completes the proof of Proposition 3.

### D. Proof of Corollary 1

Assume a sequence of networks with *m*_*p*_ RFMIOs, *p* = 1, 2, …, with lim_*p*→*∞*_ *m*_*p*_ = *∞*. For any *p*, let *e*_*z*_(*p*) be the corresponding steady-state pool density. By Proposition 2, lim_*p*→*∞*_ *e*_*z*_(*p*) = 0. For any *p*, let 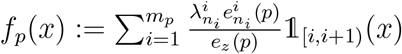. Note that *f*_*p*_(*x*) *≥* 0 for all *x*, and that 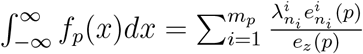 is the ratio between the total production rate and the pool density for the case of *m*_*p*_ RFMIOs in the network. By proposition 3, the functions 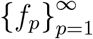 converge pointwise to 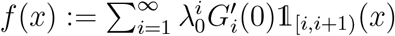. By Fatou’s lemma [7, Theorem 2.8.3]

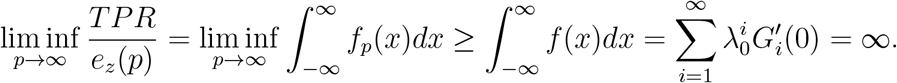

The last equality follows from Assumption 1, which implies that 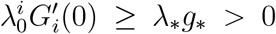. In the case where the number of RFMIOs is fixed at *m* as *e*_*z*_ → 0, by replacing Fatou’s lemma with the dominated convergence theorem [7, Theorem 2.8.1], the limit-inferior can be replaced by a limit, and the inequality can be replaced by equality. Note that in this case 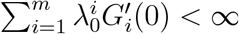. This proves (20).

To prove (21), consider first the case where the number of RFMIOs *m* is fixed and the total density in the network satisfies *s* → 0. Recall that *q* = *e*_*z*_*/s*, and (19) gives 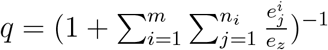. Using (17) yields

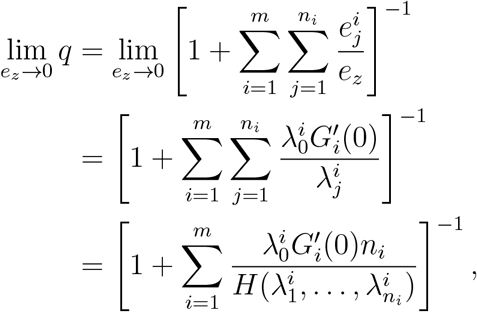

and this completes the proof. We note in passing that in the case where the total ribosome density in the network *s* is fixed and the number of RFMIOs satisfies *m* → *∞*, Assumption 1 gives

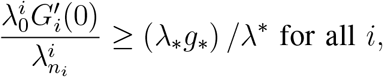

so in this case we have

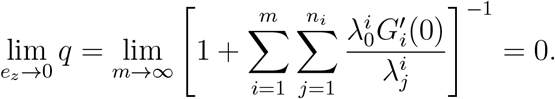

### E. Proof of Proposition 4

Using (20) gives 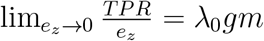 and 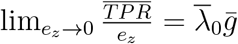, and combining this with (22) gives (24). Similarly, Eq. (21) gives

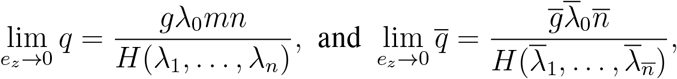

and using (22) and (23) implies that these expressions are equal. This completes the proof of Proposition 4.

## Footnotes

1 The final result of this process are the rates: 10.01, 10.33, 10.25, 9.97, 10.67, 10.59, 9.40, 9.97, 9.75, 10.67, 10.23, 10.13, 8.03.

