## Supplementary material for "Translation in the cell under fierce competition for shared resources: a mathematical model": Latex source files: rfm_near_eq_bib.bib

%%%%%%%%%%%%%%%%   bib macros

@string{CDC19="Proc.\ 19th IEEE Conf. on Decision and Control"}
@string{CDC26="Proc.\ 26th IEEE Conf. on Decision and Control"}
@string{CDC30="Proc.\ 30th IEEE Conf. on Decision and Control"}
@string{CDC32="Proc.\ 32nd IEEE Conf. on Decision and Control"}
@string{CDC33="Proc.\ 33rd IEEE Conf. on Decision and Control"}
@string{CDC34="Proc.\ 34th IEEE Conf. on Decision and Control"}
@string{CDC35="Proc.\ 35th IEEE Conf. on Decision and Control"}
@string{CDC36="Proc.\ 36th IEEE Conf. on Decision and Control"}
@string{CDC37="Proc.\ 37th IEEE Conf. on Decision and Control"}
@string{CDC38="Proc.\ 38th IEEE Conf. on Decision and Control"}
@string{CDC39="Proc.\ 39th IEEE Conf. on Decision and Control"}
@string{CDC40="Proc.\ 40th IEEE Conf. on Decision and Control"}
@string{CDC41="Proc.\ 41st IEEE Conf. on Decision and Control"}
@string{CDC42="Proc.\ 42nd IEEE Conf. on Decision and Control"}
@string{CDC43="Proc.\ 43rd IEEE Conf. on Decision and Control"}
@string{CDC44="Proc.\ 44th IEEE Conf. on Decision and Control"}
@string{CDC45="Proc.\ 45th IEEE Conf. on Decision and Control"}
@string{CDC46="Proc.\ 46th IEEE Conf. on Decision and Control"}
@string{CDC53 = "{Proc.\ 53rd IEEE Conf. on Decision and Control}"}
@string{CDC57 = "{Proc.\ 57th IEEE Conf. on Decision and Control}"}
@string{CDC60 = "{Proc.\ 60th IEEE Conf. on Decision and Control}"}


@string{ACC91="Proc.\ 1991 American Control Conf."}
@string{ACC95="Proc.\ 1995 American Control Conf."}
@string{ACC96="Proc.\ 1996 American Control Conf."}
@string{ACC97="Proc.\ 1997 American Control Conf."}
@string{ACC98="Proc.\ 1998 American Control Conf."}
@string{ACC99="Proc.\ 1999 American Control Conf."}
@string{ACC00="Proc.\ 2000 American Control Conf."}
@string{ACC01="Proc.\ 2001 American Control Conf."}
@string{ACC02="Proc.\ 2002 American Control Conf."}
@string{ACC03="Proc.\ 2003 American Control Conf."}
@string{ACC04="Proc.\ 2004 American Control Conf."}

@string{ECC2="Proc.\ 2nd European Control Conf."}
@string{ECC5="Proc.\ 5th European Control Conf."}
@string{ECC6="Proc.\ 6th European Control Conf."}
@string{ECC07="Proc.\ 9th  European Control Conf. (ECC'07)"}

@string{IFAC96="Proc.\ 13th IFAC World Congress"}
@string{IFAC99="Proc.\ 14th IFAC World Congress"}

@string{TAC="IEEE Trans.\ Automat.\ Control"}

@string{SCL="Systems Control Lett."}

@string{IJC="Int. J. Control"}

@string{IJRNC="Int. J. Robust Nonlinear Control"}

@string{IJACSP="Int. J. Adaptive Control Signal Proc."}


@string{MCSS="Math. Control Signals Systems"}

@string{EJC="European J. Control"}

@string{SIAM="SIAM J.\ Control Optim."}

@string{CSM="IEEE Control Systems Magazine"}

@string{LAA="Linear Algebra Appl."}

@string{ARC="Automat. Remote Control"}

@string{DFE="Diff. Eqns."}

@string{JDE="J. Diff. Eqns."}


@string{IEEE="Proc. IEEE"}

@string{PNAS="Proceedings of the National Academy of Sciences"}

@string{JSTATM="J. Stat. Mech."},

@string{PLOS1="PLOS ONE"}

@string{PLOSCB="PLOS Computational Biology"}

@string{PLOSB="PLOS Biology"}

@string{PLOSG="PLOS Genetics"}

@string{JTB="J. Theoretical Biology"}

@string{MBAE="Molecular Biology and Evolution"}

@string{TCBB="IEEE/ACM Trans. Comput. Biol. Bioinf."} 

@string{J_PHYS_A="J. Phys. A: Math. Gen."}

@string{p="Preprint"}


@string{in_pre="in preparation"}

@string{ta="To appear"}

@string{s="Submitted"}

@string{j="{\bf Journal version available}"}
%%%%%%%%%%%%%%%%%%%%%%%%%%%%%%%%%%%%%%%%%%%%%%%%%%%%%%%%%%
@article{MULD_SYSTEMS_WITH_INPUT,
title="DIMENSION PROBLEMS FOR
NONHOMOGENEOUS DIFFERENTIAL
EQUATIONS",
author="J.   Muldowney  and  E. Samuylova",
journal={ Canadian Applied Mathematics Quarterly},volume={18},number={1},year={2010},pages={93-105},
}
@article{sgt_tell,
author={Jiang, Z. -P. and Teel, A. R. and Praly, L.}, 
year={1994},
title={Small-gain theorem for {ISS} systems and applications}, journal=MCSS,pages={95-120},
volume={7},
number={2},
}

 

@book{bogachev2007measure,
  title={Measure Theory},
  author={ Vladimir Igorevich Bogachev},
  year={2007},
  publisher={Springer}, address={Berlin, Heidelberg},
}

@book{stein2010complex,
  title={Complex \uppercase{a}nalysis},
  author={Stein, Elias M and Shakarchi, Rami},
  volume={2},
  year={2010},
  publisher={Princeton University Press}
}


@article{Yueh2005EIGENVALUESOS,
  title={EIGENVALUES OF SEVERAL TRIDIAGONAL MATRICES},
  author={Wen-Chyuan Yueh},
  year={2005},journal={Applied Mathematics E-Notes},
	volume={5}, pages={66-74}
}


@article{tuller_traffic_jams2018,
    author = {Diament, Alon AND Feldman, Anna AND Schochet, Elisheva AND Kupiec, Martin AND Arava, Yoav AND Tuller, Tamir},
    journal = PLOSCB,
    title = {The extent of ribosome queuing in budding yeast},
    year = {2018},
    volume = {14},
    pages = {1-21},
  }

@article{neurojams,
volume={40},
title={Endosomal Traffic Jams Represent a Pathogenic Hub and Therapeutic Target in  {A}lzheimer's Disease},
year={2017},
pages={592-602},
author={S. A. Small and  S. Simoes-Spassov and R. Mayeux and 
G. A. Petsko},
journal={Trends Neurosci.},
}
 
@article{ants_jams,
author={A. Dussutour  and   J.-L. Deneubourg  and V. Fourcassie}, 
year={2005}, 
title={Temporal organization of bi-directional traffic in the ant {Lasius} niger {(L.)}},
journal={J. Exp. Biol.}, volume={208}, pages={2903-2912},
}


@article{HAN2020107610,
title = {Genome-wide Survey of Ribosome Collision},
journal = {Cell Reports},
volume = {31},
number = {5},
pages = {107610},
year = {2020},
author = {Peixun Han and Yuichi Shichino and Tilman Schneider-Poetsch and Mari Mito and Satoshi Hashimoto and Tsuyoshi Udagawa and Kenji Kohno and Minoru Yoshida and Yuichiro Mishima and Toshifumi Inada and Shintaro Iwasaki},
}

@article{traffic_jams_in_neuro,
author={S.  Subramaniam},
title={Ribosome traffic jam in neurodegeneration: decoding hurdles in {Huntington} disease},
journal={Cell Stress},
year={2021},
volume={5}, number={6}, pages={86-88},
}


@article{ROSS5911,
title = "The impacts of molecular motor traffic jams",
journal = PNAS,
volume = "109",
number = "16",
pages = "5911-5912",
year = "2012",
author = "Jennifer L. Ross",
}

 @article{Walter1997,
	author = {Walter, Wolfgang},
	journal = {Annales Polonici Mathematici},
	number = {1},
	pages = {269-274},
	title = {On strongly monotone flows},
	volume = {66},
	year = {1997},
}


@article{hastings1977existence,
  title={Existence of periodic solutions for negative feedback cellular control systems}, 
 author={Hastings, Stuart and Tyson, John and Webster, Dallas},
  journal=JDE,
  volume={25},
number={1},
  pages={39-64},
  year={1977},
}


@article{MALLICK201517,
title = "The exclusion process: A paradigm for non-equilibrium behaviour",
journal = "Physica A: Statistical Mechanics and its Applications",
volume = "418",
pages = "17-48",
year = "2015",
author = "Kirone Mallick",
}

@book{cfsystems,
title={Cyclic Feedback Systems},volume={134},issue={637},
author={T. Gedeon},address={Boston, MA}, 
year={1998}, series={Memoirs of the American Mathematical Society},
publisher={American Mathematical Society},
}

 
@article{smale_coop,
author={S. Smale}, title={On the differential equations of species in competition},
journal={J. Math. Bio.}, volume={3}, year={1976}, pages={5-7},
}

@article{CTPDS_old,
title={Dynamical Systems with  a Cyclic Sign Variation Diminishing Property},
author={Tsuff Ben-Avraham and  Guy Sharon and    Yoram Zarai and  Michael Margaliot},
note=s,url={http://arxiv.org/abs/1807.02779},
year={2018},
}

@article{pnas_sh,
author={E. Korkmazhan and H.  Teimouri and N. Peterman and E. Levine},
title={Dynamics of translation can determine the spatial organization of membrane-bound proteins and their {mRNA}},
journal=PNAS,
year={2017}, volume={114}, 
pages={13424-13429},
}


@article{natrevtrans,
author={E. Zlotorynski},year={2016},
title={Live stream: translation at single-{mRNA} resolution},
journal={Nature Reviews Molecular Cell Biology},
volume={17}, pages={395},
}

@article{INGOLIA201622,
title = "Ribosome Footprint Profiling of Translation throughout the Genome",
journal = "Cell",
volume = "165",
number = "1",
pages = "22-33",
year = "2016",
issn = "0092-8674",
author = "Nicholas T. Ingolia"
}

@article {Millseaan2755,
	author = {Mills, Eric W. and Green, Rachel},
	title = {Ribosomopathies: There{\textquoteright}s strength in numbers},
	volume = {358},
	number = {6363},
	year = {2017},
	journal = {Science}
}

@article{slow_rib,
author={A. Bastide  and A.  David},
year={2018},
title={The ribosome, (slow) beating heart of cancer (stem) cell},
volume={7}, issue={4},
journal={Oncogenesis},
}


@article{wang_trans_2015,
title = "Transversality for time-periodic competitive-cooperative tridiagonal systems",
journal = "Discrete \& Continuous Dynamical Systems - B",
volume = "20",
pages = "1821-1830",
year = "2015",
 author = "Yi Wang and Dun Zhou",
}


@article{hirs2,
author = {Morris W. Hirsch},
title = {Systems of Differential Equations that are Competitive or Cooperative {II}: 
Convergence Almost Everywhere},
journal = {SIAM J. Math. Anal.},
volume = {16},
number = {3},
pages = {423-439},
year = {1985},
}

@article{RUFFER2013277,
title = {Convergent systems vs. incremental stability},
journal = SCL,
volume = {62},
number = {3},
pages = {277-285},
year = {2013},
author = {Björn S. Rüffer and Nathan {van de Wouw} and Markus Mueller},
}


@article{cramer_non_sing,
volume={XX},
title={Cramer's rule for nonsingular $m \times n$ matrices},
journal={The Teaching of Mathematics},
author={Azamat Akhtyamov and Meirav Amram and Miriam Dagan and Artour Mouftahkov},
number={1},year={2017},
}


@article{hirs2,
title="Systems of differential equations which are competitive or cooperative~II: convergence almost
everywhere",
journal="",
number="",
author="",
}


@ARTICLE{DBLP:journals/corr/MargaliotC17,
author={S. Coogan and M. Margaliot},
journal=TAC,
title={Approximating the Steady-State Periodic Solutions of Contractive Systems},
year={2018},
note=ta,
}
 

@article{Takac1992,     
author={P. Tak\'a{\v c}},
journal={Proc. AMS},
title={Linearly Stable Subharmonic Orbits in Strongly Monotone Time-Periodic Dynamical Systems},
volume={115}, number={3}, year={1992}, pages={691-698},
}


@article{Nikolaev145201,
title = "Subharmonics and Chaos in Simple Periodically Forced Biomolecular Models ",
journal = "Biophysical J.",
volume = "114",
number = "5",
pages = "1232-1240",
year = "2018",
author = "Evgeni V. Nikolaev and Sahand Jamal Rahi and Eduardo D. Sontag"
}


@article{observ_time_varying,
author = {L. M. Silverman and H. E. Meadows},
title = {Controllability and Observability in Time-Variable Linear Systems},
journal = {SIAM J. Control},
volume = {5},
number = {1},
pages = {64-73},
year = {1967},
}


@article{BRUALDI1983769,
title = "Determinantal identities: {G}auss, {S}chur, {C}auchy, {S}ylvester, {K}ronecker, {J}acobi, {B}inet, {L}aplace, {M}uir, and {C}ayley",
journal = LAA,
volume = "52-53",
pages = "769-791",
year = "1983",
author = "Richard A. Brualdi and Hans Schneider"
}

@ARTICLE{forni2014,
author={F. Forni and R. Sepulchre},
journal=TAC,
title={A Differential {L}yapunov Framework for Contraction Analysis},
year={2014},
volume={59},
number={3},
pages={614-628},
}


@ARTICLE{Desoer1972,
  author={Desoer, C. and Haneda, H.},
  journal={IEEE Trans. Circuit Theory}, 
  title={The measure of a matrix as a tool to analyze computer algorithms for circuit analysis}, 
  year={1972},
  volume={19},
  number={5},
  pages={480-486},
  doi={10.1109/TCT.1972.1083507}}


@article{fulltppaper,
title={Revisiting Totally Positive Differential Systems: A Tutorial and New Results},
author={M. Margaliot and E. D. Sontag},
year = {2019}, 
journal={Automatica}, 
volume={101},pages={1-14},
}

@article{Kriecherbauer_2010,
	year = 2010,
	volume = {43},
	number = {40},
	pages = {403001},
	author = {Thomas Kriecherbauer and Joachim Krug},
	title = {A pedestrian`s view on interacting particle systems, {KPZ} universality and random matrices},
	journal = {Journal of Physics A: Mathematical and Theoretical} 
	}
	
@ARTICLE{2017arXiv171007321M,
   author = {M. Margaliot   and L. Gr{\"u}ne and T.  Kriecherbauer},
    title = {Entrainment in the Master Equation},
  journal = {Royal Society Open Science},
	volume = {5},pages={172157},
	number = {4},
    year = 2018,
}
 
 
 @article{SEIR_LI_MULD1995,
title = "Global stability for the {SEIR} model in epidemiology",
journal = "Math. Biosciences",
volume = "125",
number = "2",
pages = "155-164",
year = "1995",
author = "Michael Y. Li and James S. Muldowney"
}


@article{mosquito_2017,
author = {Ling Xue and Carrie A. Manore and Panpim Thongsripong and James M. Hyman},
title = {Two-sex mosquito model for the persistence of {Wolbachia}},
journal = {J. Biological Dynamics},
volume = {11},
pages = {216-237},
year  = {2017},    
}


@article{muldo1990,
author={Muldowney, J. S.},
title={Compound Matrices and Ordinary Differential Equations},
volume={20}, number={4}, year={1990}, pages={857-872},
journal={The Rocky Mountain J.  Math.},
}


@article{ando_survey,
author={T. Ando}, title={Totally positive matrices}, journal=LAA,
volume={90}, year={1987}, pages={165-219},
}
 


@book{fiedler_book,
title={Special Matrices and Their Applications in Numerical Mathematics},
author={Miroslav Fiedler},edition={2},
year={2008},
publisher={Dover Publications},
address={Mineola, NY},
}


@article{schwarz1970,
author = "Schwarz, Binyamin",
journal = "Pacific J. Math.",
number = "1",
pages = "203-229",
title = "Totally positive differential systems",
volume = "32",
year = "1970",
}


@article{SMITH199141,
title = "Convergent and oscillatory activation dynamics for cascades of neural nets with nearest 
neighbor competitive or cooperative interactions",
journal = "Neural Networks",
volume = "4",
number = "1",
pages = "41-46",
year = "1991",
author = "H. L. Smith",
}


@Incollection{Pinkus1996,
author="Pinkus, Allan",
editor="Gasca, Mariano
and Micchelli, Charles A.",
title="Spectral Properties of Totally Positive Kernels and Matrices",
bookTitle="Total Positivity and its Applications",
year="1996",
publisher="Springer Netherlands",
address="Dordrecht",
pages="477-511",
}


@Article{Fusco1990,
author="Fusco, G.
and Oliva, W. M.",
title="Transversality between invariant manifolds of periodic orbits for a class of monotone dynamical systems",
journal="J. Dyn. Differ. Equ.", 
year="1990",
volume="2",
number="1",
pages="1-17",
}


@article{barabanov:hal-01461606,
  TITLE = {Conditions for Almost Global Attractivity of a Synchronous Generator Connected to an Infinite Bus},
  AUTHOR = {Barabanov, Nikita and Schiffer, Johannes and Ortega, Romeo and Efimov, Denis},
  JOURNAL = TAC,
  YEAR = {2017},
  }

@book{linear_st,
title={Linear System Theory},
author={W. J.  Rugh}, 
publisher={Prentice-Hall},
year={1996},edition={2},
address={Upper Saddle River, NJ},
}

@article{green_wave,
author={Boris S. Kerner},
title={The physics of green-wave breakdown in a city},
journal={Europhysics Letters},  pages={28010},
volume={112}, year={2013}, number={2},
}

@article{epidemics_2006,
author={N. C. Grassly and C. Fraser},
title={Seasonal Infectious Disease Epidemiology},
Journal={Proc. Royal Society B: Biological Sciences},
volume={273},
pages={2541-2550}, year={2006},
}

@book{chicone_2006,
title={Ordinary Differential Equations with Applications},
author={C. Chicone}, 
publisher={Springer},
year={2006},edition={2}, 
address={New York},
}
 

@article{periodic_tridi_smith,
author = {H. L. Smith},
title = {Periodic Tridiagonal Competitive and Cooperative Systems of Differential Equations},
journal = {SIAM J. Math. Anal.},
volume = {22},
number = {4},
pages = {1102-1109},
year = {1991},
}


@article{poin_cyclic,
title = "The {Poincare-Bendixson} theorem for monotone cyclic feedback systems",
author = "J. Mallet-Paret and H. L. Smith",
year = "1990",
volume = "2",
pages = "367-421",
journal = "J. Dyn. Differ. Equ.",
number = "4",
}


@ARTICLE{chua_roska_1990,
author={L. O. Chua and T. Roska},
journal={IEEE Trans. Circuits and Systems},
title={Stability of a class of nonreciprocal cellular neural networks},
year={1990},
volume={37},
number={12},
pages={1520-1527},
}


@book{diff_eqn_limit_sets,
publisher={American Mathematical Society},
series={Graduate Studies in Mathematics},
volume={140},
year={2012},
author={Gerald Teschl},
title={Ordinary Differential Equations and Dynamical Systems},
}


 

@article{Donnell2009120,
title = "Monotone dynamics of two cells dynamically coupled by a voltage-dependent gap junction ",
journal = JTB,
volume = "261",
number = "1",
pages = "120-125",
year = "2009",
    author = "Peter Donnell and Stephen A. Baigent and Murad Banaji",
 }

 
@book{gk_book,
title={Oscillation Matrices and Kernels and Small Vibrations of Mechanical Systems},
author={F. R. Gantmacher and M. G. Krein},
publisher={American Mathematical Society},
year={2002},address={Providence, RI}, 
note={Translation based on the~1941 {Russian} original},
}


@article{intssr,
author = {M. Adm and J. Garloff},
title = {Intervals of special sign regular matrices},
journal = {Linear and Multilinear Algebra},
volume = {64},
number = {7},
pages = {1424-1444},
year  = {2016},
 }


@Article{Loewner1955,
author="Loewner, Charles",
title="On totally positive matrices",
journal="Mathematische Zeitschrift",
year="1955",
volume="63",
number="1",
pages="338-340",
}

 


@book{pinkus,
author={A. Pinkus},
title={Totally Positive Matrices},
publisher={Cambridge University Press}, address={Cambridge, UK},
  year={2010}, 
} 

@book{total_book,
title={Totally Nonnegative Matrices},
author={S. M. Fallat and C. R. Johnson},
year={2011}, 
publisher={Princeton University Press},
address={Princeton, NJ},
}


@ARTICLE{smith_sign_changes,
author={H. L. Smith},
title={A discrete {Lyapunov} function for a class of linear differential equations},
journal={Pacific J. Math.},
volume={144},  year={1990}, number={2}, pages={345-360},
} 


@ARTICLE{netwok_contractive,
author={Russo, G. and di Bernardo, M. and Sontag, E. D.},
journal=TAC,
title={A Contraction Approach to the Hierarchical Analysis and Design of Networked Systems},
year={2013},
volume={58},
number={5},
pages={1328-1331},
}


@article{Dyson_chain,
  title = {The Dynamics of a Disordered Linear Chain},
  author = {Dyson, Freeman J.},
  journal = {Phys. Rev.},
  volume = {92},
  pages = {1331-1338},
   year = {1953},
}


@book{random_matrices,
author={Madan Lal Mehta},
publisher={Academic Press},
title={Random Matrices},edition={3},
year={2004},
}


@article{Stochasticity_2005,
authoe={Kaern, Mads
and Elston, Timothy C.
and Blake, William J.
and Collins, James J.},
title={Stochasticity in gene expression: from theories to phenotypes},
journal={Nat. Rev. Genet.},
year={2005},
volume={6},
pages={451-464},
}

@article{Stoch_13,
    author = {Singh,   A. AND Soltani,   M.},
    journal = PLOS1,
    title = {Quantifying Intrinsic and Extrinsic Variability
    in Stochastic Gene Expression Models},
    year = {2013},
    volume = {8},
    pages = {e84301},
}

@article{gain_entr_RFM,
author={Ali Al-Radhawi and M. Margaliot
and E. D. Sontag}, year={2021},
title={Maximizing average throughput in oscillatory biochemical synthesis systems: an optimal control approach}, journal={R. Soc. open sci.},
number={8210878210878}, url={http://doi.org/10.1098/rsos.210878}
}

@ARTICLE{10.3389/fbioe.2019.00248,
AUTHOR={Khambhati, Khushal and Bhattacharjee, Gargi and Gohil, Nisarg and Braddick, Darren and Kulkarni, Vishwesh and Singh, Vijai},   
TITLE={Exploring the Potential of Cell-Free Protein Synthesis for Extending the Abilities of Biological Systems},      
JOURNAL={Frontiers in Bioengineering and Biotechnology},      
VOLUME={7},           
YEAR={2019},      
URL={https://www.frontiersin.org/articles/10.3389/fbioe.2019.00248},       
	}
   

@article{Elowitz16082002,
author = {Elowitz, Michael B. and Levine, Arnold J. and Siggia, Eric D. and Swain, Peter S.},
title = {Stochastic Gene Expression in a Single Cell},
volume = {297},
pages = {1183-1186},
year = {2002},
journal = {Science}
}

@article{RANDOM_ATTRACTION_TASEP,
author = {Gruene, Lars and Kriecherbauer, Thomas and Margaliot, Michael},
title = {Random Attraction in the {TASEP} Model},
journal = {SIAM J. on Applied Dynamical Systems},
volume = {20},
number = {1},
pages = {65-93},
year = {2021},
}


@article{edge_tasep_2009,
author={J. J. Dong and  R. K. P. Zia and B. Schmittmann},
title={Understanding the edge effect in {TASEP} with mean-field theoretic approaches},
journal=J_PHYS_A,
volume={42},number={1}, pages={015002}, year={2009},
}

@ARTICLE{Ben-Yehezkel2014,
  author =       {T. Ben-Yehezkel and S. Atar and T. Marx and R. Cohen and A. Diament and A. Dana and A. Feldman and E. Shapiro and T. Tuller},
  title =        {Systematic study of synthetic transcript features in {S.} cerevisiae exposes gene-expression determinants},
 note=s,
  year =         {2014},
}

@article{PhysRevE.71.011103,
  title = {Steady-state properties of a totally asymmetric exclusion process with periodic structure},
  author = {Lakatos, Greg and Chou, Tom and Kolomeisky, Anatoly},
  journal = {Phys. Rev. E},
  volume = {71},
  issue = {1},
  pages = {011103},
   year = {2005},
 }


@ARTICLE{Tuller2014,
  author =       {T. Tuller and H. Zur},
  title =        {Multiple Roles of the Coding Sequence 5 End in Gene Expression Regulation},
  note=s,
  year =         {2014},
}

@article{sturm_tridiagnal,
year={2009},
journal={Found.  Comput.  Math.},
volume={9},
number={4},
title={Sturm Sequences and Random Eigenvalue Distributions},
author={Albrecht, James T. and Chan, Cy P. and Edelman, Alan},
pages={461-483},
}


@ARTICLE{Malys2011,
  author =       {N. Malys and J.E. McCarthy},
  title =        {Translation initiation: variations in the mechanism can be anticipated},
  journal =      {Cell Mol Life Sci.},
  year =         {2011},
  volume =       {68},
  number =       {6},
  pages =        {991-1003},
}


@article{PhysRevLett.93.198101,
  title = {Clustered Bottlenecks in {mRNA} Translation and Protein Synthesis},
  author = {T. Chou and G. Lakatos},
  journal = {Phys. Rev. Lett.},
  volume = {93},
  issue = {19},
  pages = {198101},
  year = {2004},
}

@article{PhysRevE.74.031906,
  title = {Bottleneck-induced transitions in a minimal model for intracellular transport},
  author = {Pierobon, Paolo and Mobilia, Mauro and Kouyos, Roger and Frey, Erwin},
  journal = {Phys. Rev. E},
  volume = {74},
  issue = {3},
  pages = {031906},
   year = {2006},
 }

 @article{PhysRevE.76.051113,
  title = {Inhomogeneous exclusion processes with extended objects: The effect of defect locations},
  author = {Dong, J. J. and Schmittmann, B. and Zia, R. K. P.},
  journal = {Phys. Rev. E},
  volume = {76},
   pages = {051113},
   year = {2007},
 }


@article{zarai_infi,
author = {Y. Zarai and M. Margaliot and T. Tuller},
title = {Explicit expression for the steady-state translation rate in the infinite-dimensional
homogeneous ribosome flow model}, year={2013},
volume={10}, isuue={5}, pages={1322-1328},
journal = TCBB,
}


@article{toeplitz_2013,
journal={Numer. Linear Algebra Appl.},
 year={2013},
  volume={20}, pages={302-326},
title={Tridiagonal {T}oeplitz matrices: properties and novel applications},
author={S. Noschese  and L. Pasquini   and L. Reichel},
}

@ARTICLE{Ren2006,
  author =       {X.Y. Ren and O. Vorst and M.W. Fiers and W.J. Stiekema and J.P. Nap},
  title =        {In plants, highly expressed genes are the least compact},
  journal =      {Trends Genet.},
  year =         {2006},
  volume =       {22},
  number =       {10},
  pages =        {528-32},
}


@ARTICLE{Shaham2014,
  author =       {G. Shaham and T. Tuller},
  title =        {Most associations between transcript features and gene expression are monotonic},
  journal =      {Mol Biosyst.},
  year =         {2014},
  volume =       {10},
  number =       {6},
  pages =        {1426-40},
}


@ARTICLE{Na2010,
  author =       {D. Na and S. Lee and D. Lee},
  title =        {Mathematical modeling of translation initiation for the estimation of its efficiency to computationally design mRNA sequences with desired expression levels in prokaryotes},
  journal =      {BMC Syst Biol.},
  year =         {2010},
  volume =       {4},
  pages =        {71},
}


@article{Vasquez17012014,
author = {Vasquez, Juan-Jose  and Hon, Chung-Chau and Vanselow, Jens T. and Schlosser, Andreas and Siegel, T. Nicolai},
title = {Comparative ribosome profiling reveals extensive
translational complexity in different {T}rypanosoma brucei life cycle stages},
year = {2014},volume={42}, pages={3623-3637},
number={6},
journal = {Nucleic Acids Res.},
}


@ARTICLE{Bloom-Ackermann2014,
  author =       {Z. Bloom-Ackermann and S. Navon and H. Gingold and R. Towers and Y. Pilpel and O. Dahan},
  title =        {A Comprehensive tRNA Deletion Library Unravels the Genetic Architecture of the tRNA Pool},
  journal =      PLOSG,
  year =         {2014},
  volume =       {10},
  number =       {1},
  pages =        {e1004084},
}


@ARTICLE{Charneski2013,
  author =       {C.A. Charneski and L.D. Hurst},
  title =        {Positively charged residues are the major determinants of ribosomal velocity},
  journal =      PLOSB,
  year =         {2013},
  volume =       {11},
  number =       {3},
  pages =        {e1001508},
}


@article{kolo_sense_1998,
  author={A. B. Kolomeisky},
  title={Asymmetric simple exclusion model with local inhomogeneity},
  journal=J_PHYS_A,
  volume={31},
  number={4},
  pages={1153-1164},
   year={1998},
 }
	


  

@ARTICLE{Racle2013,
  author =       {J. Racle and F. Picard and L. Girbal and M. Cocaign-Bousquet and V. Hatzimanikatis},
  title =        {A genome-scale integration and analysis of {Lactococcus} lactis translation data},
  journal =      PLOSCB,
  year =         {2013},
  volume =       {9},
  number =       {10},
  pages =        {e1003240},
}


@ARTICLE{Deneke2013,
  author =       {C. Deneke and R. Lipowsky and A. Valleriani},
  title =        {Effect of ribosome shielding on {mRNA} stability},
  journal =      {Phys. Biol.},
  year =         {2013},
  volume =       {10},
  number =       {4},
  pages =        {046008},
}


@ARTICLE{Kopito2000,
  author =       {R.R. Kopito},
  title =        {Aggresomes, inclusion bodies and protein aggregation},
  journal =      {Trends Cell Biol.},
  year =         {2000},
  volume =       {10},
  number =       {12},
  pages =        {524-30},
}


@ARTICLE{Park2008,
  author =       {J.S. Park and K.Y. Han and J.H. Lee and J.A. Song and K.Y. Ahn and H.S. Seo and S.J. Sim
  and S.W. Kim and J. Lee},
  title =        {Solubility enhancement of aggregation-prone heterologous proteins by
   fusion expression using stress-responsive Escherichia coli protein, {RpoS}},
  journal =      {BMC Biotechnol.},
  year =         {2008},
  volume =       {8},
  pages =        {15},
}


@ARTICLE{Mittler2010,
  author =       {R. Mittler and E. Blumwald},
  title =        {Genetic engineering for modern agriculture: challenges and perspectives},
  journal =      {Annu Rev Plant Biol.},
  year =         {2010},
  volume =       {61},
  pages =        {443-62},
}


@ARTICLE{Goeddel1979,
  author =       {D.V. Goeddel and D.G. Kleid and F. Bolivar and H.L. Heyneker and D.G. Yansura and R. Crea and T. Hirose and A.  Kraszewski and K. Itakura and A.D. Riggs},
  title =        {Expression in {E}scherichia coli of chemically synthesized genes for human insulin},
  journal =      PNAS,
  year =         {1979},
  volume =       {76},
  number =       {1},
  pages =        {106-10},
}

@article{Alon26092003,
author = {U. Alon},
title = {Biological Networks: The Tinkerer as an Engineer},
volume = {301},
number = {5641},
pages = {1866-1867},
year = {2003},
journal = {Science},
}


@book{opt_in_bio_book,
author={R. Rosen},
title={Optimality Principles in Biology},
publisher={Butterworths},
address={London},
year={1967},
}

 
@ARTICLE{Dana2014,
  author =       {A. Dana and T. Tuller},
  title =        {The effect of {tRNA} levels on decoding times of {mRNA} codons},pages={9171-9181},volume={42},number={14},
  year =         {2014},
journal={Nucleic Acids Res.},
}


@ARTICLE{Binnie1997,
  author =       {C. Binnie and J. D. Cossar and D. I. Stewart},
  title =        {Heterologous biopharmaceutical protein expression in {Streptomyces}},
  journal =      {Trends Biotechnol.},
  year =         {1997},
  volume =       {15},
  number =       {8},
  pages =        {315-20},
}

@Article{RFM_RANDOM,
title={Variability in {mRNA} Translation: A Random Matrix Theory Approach},
author={Michael Margaliot and 
 Wasim Huleihel 
and Tamir Tuller},
journal={Scientific Reports},
year={2020},
note=ta,
}


@ARTICLE{Moks1987,
  author =       {T. Moks and L. Abrahmsen and E. Holmgren and M. Bilich and A. Olsson and G. Pohl and C. Sterky and H. Hultberg and S. A. Josephson},
  title =        {Expression of human insulin-like growth factor {I} in bacteria: use of optimized gene fusion vectors to facilitate protein purification},
  journal =      {Biochemistry},
  year =         {1987},
  volume =       {26},
  number =       {17},
  pages =        {5239-44},
}


@ARTICLE{Romanos1992,
  author =       {M.A. Romanos and C.A. Scorer and J.J Clare},
  title =        {Foreign gene expression in yeast: a review},
  journal =      {Yeast},
  year =         {1992},
  volume =       {8},
  number =       {6},
  pages =        {423-88},
}


@article{solvers_guide,
  author={R. A. Blythe and M. R. Evans},
  title={Nonequilibrium steady states of matrix-product form: a solver's guide},
  journal=J_PHYS_A,
  volume={40},
  number={46},
  pages={R333-R441},
  year={2007},
}
	

@ARTICLE{Lee2012,
  author =       {S. Lee  and B. Liu and S. Lee and S.X. Huang and B. Shen and S.B. Qian},
  title =        {Global mapping of translation initiation sites in mammalian cells at single-nucleotide resolution},
  journal =      PNAS,
  year =         {2012},
  volume =       {109},
  number =       {37},
  pages =        {E2424-32}
}

@ARTICLE{Zur2013,
  author =       {H. Zur and T. Tuller},
  title =        {New universal rules of eukaryotic translation initiation fidelity},
  journal =      PLOSCB,
  year =         {2013},
  volume =       {9},
  number =       {7},
  pages =        {e1003136},
}


@ARTICLE{dos-Reis2009,
  author =       {M. dos Reis and L. Wernisch},
  title =        {Estimating translational selection in eukaryotic genomes},
  journal =      MBAE,
  year =         {2009},
  volume =       {26},
  number =       {2},
  pages =        {451-61},
}

 


@ARTICLE{Firczuk2013,
  author =       {H. Firczuk and S. Kannambath and J. Pahle and A. Claydon and R. Beynon and J. Duncan and H. Westerhoff and P. Mendes and J.E. McCarthy},
  title =        {An in vivo control map for the eukaryotic {mRNA} translation machinery},
  journal =      {Mol Syst Biol.},
  year =         {2013},
  volume =       {9},
  pages =        {635},
}


@ARTICLE{Potapov2012,
  author =       {I. Potapov and J. Makela and O. Yli-Harja and A. S. Ribeiro},
  title =        {Effects of codon sequence on the dynamics of genetic networks},
  journal =      JTB,
  year =         {2012},
  volume =       {315},
  pages =        {17-25},
}


@ARTICLE{Shah2013,
  author =       {P. Shah and Y. Ding and M. Niemczyk and G. Kudla and J.M. Plotkin},
  title =        {Rate-limiting steps in yeast protein translation},
  journal =      {Cell},
  year =         {2013},
  volume =       {153},
  number =       {7},
  pages =        {1589-601},
}


@ARTICLE{Sauna2011,
  author =       {Z. E. Sauna and C. Kimchi-Sarfaty },
  title =        {Understanding the contribution of synonymous mutations to human disease},
  journal =      {Nature Reviews Genetics},
  year =         {2011}, volume={12}, pages={683–691},
}


@ARTICLE{Kimchi-Sarfaty2013,
  author =       {C. Kimchi-Sarfaty and T. Schiller and N. Hamasaki-Katagiri and M.A. Khan and C. Yanover and Z.E. Sauna},
  title =        {Building better drugs: developing and regulating engineered therapeutic proteins},
  journal =      {Trends Pharmacol. Sci.},
  year =         {2013}, volume={34}, number={10},pages={534-548},
}


@ARTICLE{Plotkin2011,
  author =       {J.B. Plotkin and G. Kudla},
  title =        {Synonymous but not the same: the causes and consequences of codon bias},
  journal =      {Nat. Rev. Genet.},
  year =         {2011},
  volume =       {12},
  pages =        {32-42},
}


@ARTICLE{Tuller2010,
  author =       {T. Tuller and A. Carmi and K. Vestsigian and S. Navon and Y. Dorfan and J. Zaborske and T. Pan and
                  O. Dahan and I. Furman and Y. Pilpel},
  title =        {An evolutionarily conserved mechanism for controlling the efficiency of protein translation},
  journal =      {Cell},
  year =         {2010},
  volume =       {141},
  number =       {2},
  pages =        {344-54},
}


@article{one_100_fold,
journal={RNA},
year={2012}, volume={18},
pages={2299-2305},
title={Alternative transcription start site selection leads to large differences in translation activity in yeast},
author={Maria F. Rojas-Duran and Wendy V. Gilbert},
}


@BOOK{Bonnans2006,
  author =       {J. F. Bonnans and J. C. Gilbert and C. Lemarechal and C. A. Sagastizabal },
  title =        {Numerical Optimization: Theoretical and Practical Aspects},
  publisher =    {Springer-Verlag},
  year =         {2006},
  address =      {Berlin},
}

@BOOK{Kiwiel1985,
  author =       {K. Kiwiel},
  title =        {Methods of Descent for Nondifferentiable Optimization},
  year =         {1985},
  address =      {Berlin},
}


@BOOK{Dimitri1999,
  author =       {D. P. Dimitri },
  title =        {Nonlinear Programming},
  publisher =    {Athena Scientific},
  year =         {1999},
  address =      {Cambridge, MA.},
}


@ARTICLE{Edri2013,
 author =       {S. Edri and  E. Gazit and E. Cohen and T. Tuller},
 title =        {The {RNA} Polymerase Flow Model of Gene Transcription},
 journal={IEEE Trans. Biomed. Circuits Syst.},
 volume={8}, number={1},pages={54-64},
 year =         {2014},
}


@ARTICLE{Zur2012,
  author =       {H. Zur and T. Tuller},
  title =        {RFMapp: ribosome flow model application},
  journal =      {Bioinformatics},
  year =         {2012},
  volume =       {28},
  number =       {12},
  pages =        {1663-4},
}


@ARTICLE{Chu2012,
  author =       {D. Chu and N. Zabet and T. von der Haar},
  title =        {A novel and versatile computational tool to model translation},
  journal =      {Bioinformatics},
  year =         {2012},
  volume =       {28},
  number =       {2},
  pages =        {292-3},
}


@ARTICLE{Yau2004,
journal={Annals of Mathematics},
volume={159},year={2004}, pages={377-405},
title={$(\log t)^{2/3}$ law of the two dimensional
asymmetric simple exclusion process},
author={H-T. Yau},
}


@ARTICLE{Gilchrist2006,
  author =       {M. A. Gilchrist and A. Wagner},
  title =        {A model of protein translation including codon bias, nonsense errors, and ribosome recycling},
  journal =      JTB,
  year =         {2006},
  volume =       {239},
  number =       {4},
  pages =        {417-34},
}


@BOOK{Morgan2007,
  author =       {D. O. Morgan},
  title =        {The cell cycle, principles of control},
  publisher =    {New Science Press},
  year =         {2007},
  address =      {London},
}


@ARTICLE{Talia2007,
  author =       {S. Di Talia and J. M. Skotheim and J. M. Bean and E. D. Siggia and F. R. Cross},
  title =        {The effects of molecular noise and size control on variability in the budding yeast cell cycle},
  journal =      {Nature},
  year =         {2007},
  volume =       {448},
  number =       {7156},
  pages =        {947-51},
}


@ARTICLE{Aitken2010,
  author =       {C. E. Aitken and A. Petrov and J. D. Puglisi},
  title =        {Single ribosome dynamics and the mechanism of translation},
  journal =      {Annu. Rev. Biophys.},
  year =         {2010},
  volume =       {39},
  number =       {491-513},
}


@ARTICLE{Petrov2011,
  author =       {A. Petrov and G. Kornberg and S. O'Leary and A. Tsai and S. Uemura and J.D. Puglisi},
  title =        {Dynamics of the translational machinery},
  journal =      {Curr. Opin. Struct. Biol.},
  year =         {2011},
  volume =       {21},
  number =       {1},
  pages =        {137-45},
}


@ARTICLE{Uemura2010,
  author =       {S. Uemura and C. E. Aitken and J. Korlach and B. A. Flusberg and S. W. Turner and J. D. Puglisi},
  title =        {Real-time {tRNA} transit on single translating ribosomes at codon resolution},
  journal =      {Nature},
  year =         {2010},
  volume =       {464},
  pages =        {1012-1017},
}


@ARTICLE{Culver2000,
  author =       {G. M. Culver and H. F. Noller},
  title =        {In vitro reconstitution of {30S} ribosomal subunits using complete set of recombinant proteins},
  journal =      {Methods Enzymol.},
  year =         {2000},
  volume =       {318},
  pages =        {446-60},
}


@ARTICLE{Wen2008,
  author =       {J. D. Wen and L. Lancaster and C. Hodges A. C. Zeri and S. H. Yoshimura and H. F. Noller and C. Bustamante and I. Tinoco},
  title =        {Following translation by single ribosomes one codon at a time},
  journal =      {Nature},
  year =         {2008},
  volume =       {452},
  number =       {7187},
  pages =        {598-603},
}


@ARTICLE{Kruse2005,
  author =       {K. Kruse and F. Julicher},
  title =        {Oscillations in cell biology},
  journal =      {Curr. Opin. Cell Biol.},
  year =         {2005},
  volume =       {17},
  number =       {1},
  pages =        {20-6},
}


@ARTICLE{Ledda2005,
  author =       {M. Ledda and M. Di Croce and B. Bedini and F. Wannenes and
   M. Corvaro and P. P. Boyl and S. Caldarola and F. Loreni and F. Amaldi},
  title =        {Effect of {3'UTR}
   length on the translational regulation of 5'-terminal oligopyrimidine {mRNAs}},
  journal =      {Gene},
  year =         {2005},
  volume =       {344},
  pages =        {213-20},
}


@BOOK{jonesandthron,
author={W. B. Jones and W. J. Thron},
title={Continued Fractions: Analytic Theory and Applications},
year={1980},
publisher={Addison-Wesley},
address={Reading, MA},
}


@ARTICLE{Gu2010,
  author =       {W. Gu and T. Zhou and C. O. Wilke},
  title =        {A universal trend of reduced {mRNA}
   stability near the translation-initiation site in prokaryotes and eukaryotes},
  journal =      PLOSCB,
  pages={e1000664},
  year =         {2010},
  volume =       {6},
}

@ARTICLE{infi_HRFM,
  author =       {Y. Zarai and M. Margaliot and T. Tuller},
  title =        {Explicit expression for the  steady-state translation rate  in the infinite-dimensional homogeneous ribosome flow model},
  journal =       TCBB,
  year =         {2013},
  volume=10, number={5}, pages={1322-1328},
}


@ARTICLE{Dana2012,
  author =       {A. Dana and T. Tuller},
  title =        {Determinants of translation elongation speed and ribosomal profiling biases in mouse embryonic stem cells},
  journal =      PLOSCB,
  year =         {2012},
  volume =       {8},
  number =       {12},
  pages =        {e1002755},
}


@ARTICLE{Ciandrini2013,
  author =       {L. Ciandrini and I. Stansfield and M.C. Romano},
  title =        {Ribosome Traffic on {mRNA}s Maps to Gene Ontology:
  Genome-wide Quantification of Translation Initiation Rates and Polysome Size Regulation},
  journal =      PLOSCB,
  year =         {2013},
  volume =       {9},   pages = {e1002866},
  number =       {1},
}


@ARTICLE{Kondrashov2011,
  author =       {N. Kondrashov and A. Pusic and C.R. Stumpf and K. Shimizu and A.C. Hsieh and S. Xue and J. Ishijima and T. Shiroishi and M. Barna},
  title =        {Ribosome-mediated specificity in {Hox} {mRNA} translation and vertebrate tissue patterning},
  journal =      {Cell},
  year =         {2011},
  volume =       {145},
  number =       {3},
  pages =        {383-97},
}


@ARTICLE{Hunt2008,
  author =       {A.G. Hunt and R. Xu and B. Addepalli and S. Rao and K.P. Forbes and L.R. Meeks and D. Xing and M. Mo and H. Zhao and A. Bandyopadhyay and L. Dampanaboina and A. Marion and C. Von Lanken and Q.Q. Li},
  title =        {Arabidopsis {mRNA} polyadenylation machinery:
   comprehensive analysis of protein-protein interactions and gene expression profiling},
  journal =      {BMC Genomics},
  year =         {2008},
  volume =       {9},
  pages =        {220},
}


@ARTICLE{Guhaniyogi2001,
  author =       {J. Guhaniyogi and G. Brewer},
  title =        {Regulation of {mRNA} stability in mammalian cells},
  journal =      {Gene},
  year =         {2001},
  volume =       {265},
  number =       {1-2},
  pages =        {11-23},
}


@article{Alon_gExp,
author={  Jean Hausser and Avi Mayo and Leeat Keren and Uri Alon},
title={Central dogma rates and the trade-off between precision and economy in gene expression},
journal={Nat. Commun.},
volume={10}, number={68}, year={2019},
}


@article{dekel_alon_2005,
author={E. Dekel and U. Alon},
title={Optimality and evolutionary tuning of the expression level of a protein},
journal={Nature},
year={2005},
volume={436},
pages={588-592},
}


@article{may_smith_opt,
author={G. A. Parker and J. {Maynard Smith}},
title={Optimality theory in evolutionary biology},
journal={Nature},
volume={348},
pages={27-33},
year={1990},
}

@article{ecoli_optimal,
author={R. U. Ibarra and J. S. Edwards  and B. O. Palsson},
title={Escherichia coli {K-12} undergoes
adaptive evolution to achieve in silico predicted optimal growth},
journal={Nature},
volume={420},
pages={186-189},
year={2002},
}


@article{recycle2013,
title={Tying up loose ends: ribosome recycling in 
eukaryotes and archaea},
journal={Trends Biochem Sci.},
year={2013},
volume={38},
number={2},
pages={64-74},
author={E. Nurenberg and R.  Tampe},
}


@ARTICLE{Howard1970,
  author =       {G. A. Howard and S. D. Adamson and E. Herbert },
  title =        {Subunit recycling during translation in a reticulocyte cell-free system},
  journal =      {J. Biol. Chem.},
  year =         {1970},
  volume =       {245},
  pages =        {6237-6239},
}


@ARTICLE{Tarun1997,
  author =       {S. Z. Tarun and A. B. Sachs},
  title =        {Binding of eukaryotic translation initiation factor {4E} {(eIF4E)}
   to {eIF4G} represses translation of uncapped {mRNA}},
  journal =      {Mol. Cell. Biol.},
  year =         {1997},
  volume =       {17},
  pages =        {6876-6886},
}


@ARTICLE{Craig1998,
  author =       {A. W. Craig and A. Haghighat and A. T. Yu and N. Sonenberg},
  title =        {Interaction of polyadenylate-binding protein with
  the {eIF4G} homologue {PAIP} enhances translation},
  journal =      {Nature},
  year =         {1998},
  volume =       {392},
  number =       {6675},
  pages =        {520-3},
}


@INPROCEEDINGS{AngeliS2004,
  author =      { D. Angeli and E. D. Sontag.},
  title =        { Interconnections of monotone systems with
steady-state characteristics},
  booktitle =    {Optimal control, stabilization and nonsmooth analysis},
  year =        {2004},
  editors=      {M. S. \mbox{de Queiroz} and M. A. Malisoff and P.
R. Wolenski},
  volume =      {301},
  series =      {Lecture Notes in Control and Inform. Sci.},
  pages =        {135-154},
  address =      {Berlin},
  publisher =    {Springer},
}


@article{HIRSCH1989331,
title = "Convergent activation dynamics in continuous time networks",
journal = "Neural Networks",
volume = "2",
number = "5",
pages = "331-349",
year = "1989",
 author = "Morris W. Hirsch",
 }

@book{waad,
title={Continued Fractions: Convergence Theory},
author={L. Lorentzen and H. Waadeland},
year={2008}, volume={1},edition={2},
publisher={Atlantis Press}, address={Paris},
}


@ARTICLE{Xu2013,
  author =       {Y. Xu and P. Ma and P. Shah and A. Rokas and Y. Liu and C. Hirschie Johnson},
  title =        {Non-optimal codon usage is a mechanism to achieve circadian clock conditionality},
  journal =      {Nature},
  year =         {2013},
  volume =       {doi:10.1038/nature11942},
}

@ARTICLE{Zhou2013,
  author =       {M. Zhou and J. Guo and J. Cha and M. Chae and S. Chen and J. M. Barral and M. S. Sachs and Y. Liu},
  title =        {Non-optimal codon usage affects expression, structure and function of clock protein {FRQ}},
  journal =      {Nature},
  year =         {2013},
  volume =       {doi: 10.1038/nature11833},
}


@ARTICLE{Collins1997,
  author =       {K. Collins and T. Jacks and N. P. Pavletich},
  title =        {The cell cycle and cancer},
  journal =      PNAS,
  year =         {1997},
  volume =       {94},
  number =       {7},
  pages =        {2776-2778},

}


@ARTICLE{Akashi2002,
  author =       {H. Akashi and T. Gojobori},
  title =        {Metabolic efficiency and amino acid composition in the proteomes
   of {E}scherichia coli and {B}acillus subtilis},
  journal =      PNAS,
  year =         {2002},
  volume =       {99},
  number =       {6},
  pages =        {3695-700},
}


@article{phase_lock,
    author = {Jovic, Andreja AND Howell, Bryan AND Cote, Michelle AND Wade, Susan M. AND Mehta, Khamir AND Miyawaki, Atsushi AND Neubig, Richard R. AND Linderman, Jennifer J. AND Takayama, Shuichi},
    journal = PLOSCB,
    title = {Phase-Locked Signals Elucidate Circuit Architecture of an Oscillatory Pathway},
    year = {2010},
    volume = {6},
    pages = {e1001040},
}

@article{tasep_ad_hoc_nets,
 author = {S. Srinivasa and M. Haenggi},
  title = {A Statistical Mechanics-Based Framework
   to Analyze Ad Hoc Networks with Random Access},
    journal ={IEEE Trans.  Mobile Computing},
    volume = {11},
     year = {2012},
     pages = {618-630},
}

@article{RFM_feedback,
title = {Ribosome Flow Model with Positive Feedback},
journal={J. Royal Society Interface},
year = {2013},
author = {M. Margaliot and T. Tuller},
volume={10},    pages={20130267},
}


@article{FerrellJr2011874,
title = "Modeling the Cell Cycle: Why Do Certain Circuits Oscillate?",
journal = "Cell",
volume = "144",
number = "6",
pages = "874-885",
year = "2011",
author = "J. E. {Ferrell Jr.} and T. Yu-Chen Tsai and Q. Yang",
}


@article{angeli_sontag_pnas_2004,
author={D. Angeli and
J. E. Ferrell and E. D. Sontag},
title={Detection of multistability, bifurcations, and hysteresis
in a large class of biological positive-feedback systems},
journal=PNAS,
volume={101}, pages={1822-1827},
year={2004},
}


@article{Vidyasagar197890,
title = "On matrix measures and convex {L}iapunov functions",
journal = "J. Mathematical Analysis   Applications",
volume = "62",
pages = "90-103",
year = "1978",
author = "M. Vidyasagar"
}


@ARTICLE{Frenkel2012,
  author =       {M. Frenkel-Morgenstern and T. Danon and T. Christian and T. Igarashi and L. Cohen and Y. M. Hou and L. J. Jensen},
  title =        {Genes adopt non-optimal codon usage to generate cell cycle-dependent oscillations in protein levels},
  journal =      {Mol. Syst. Biol.},
  year =         {2012},
  volume =       {8},
  pages =        {572},
}


@ARTICLE{Wells1998,
  author =       {S. E. Wells and P. E. Hillner and R.D. Vale A. B. Sachs},
  title =        {Circularization of {mRNA} by Eukaryotic Translation Initiation Factors},
  journal =      {Molecular Cell},
  year =         {1998},
  volume =       {2},
  number =       {1},
  pages =        {135-140},
}


@book{num_the_rosen,
title={Elementary Number Theory and its Applications},
author={K. H. Rosen},
publisher={Addison-Wesley},
year={2000},
edition={4},
}


@ARTICLE{LopezLastra2005,
  author =       {M. LÃ³pez-Lastra and A. Rivas and M.I. BarrÃ­a},
  title =        {Protein synthesis in eukaryotes: the growing biological relevance of cap-independent translation initiation},
  journal =      {Biol Res.},
  year =         {2005},
  volume =       {38},
  number =       {2-3},
  pages =        {121-46},
}


@ARTICLE{Belle2006,
  author =       {A. Belle and A. Tanay and L. Bitincka and R. Shamir and E.K. O'Shea},
  title =        {Quantification of protein half-lives in the budding yeast proteome.},
  journal =      PNAS,
  year =         {2006},
  volume =       {103},
  number =       {35},
  pages =        {13004-13009},
}


@article{mehra2003,
author={A. Mehra  and K. H.  Lee and V. Hatzimanikatis},
title={Insights into the relation between {mRNA} and protein expression patterns: {I.} Theoretical considerations},
journal={Biotechnol Bioeng.},
year={2003},
volume={84}, number={7},  pages={822-33},
}


@article{Zur2020,
author={Hadas Zur and Rachel Cohen-Kupiec and  Sophie Vinokour and Tamir Tuller},
title={Algorithms for ribosome traffic engineering and their potential in improving host cells' titer and growth rate},
journal={Sci. Rep.}, volume={10},pages={21202}, year={2020},
}


@incollection{cast_book,
	author="Margaliot, Michael and Tuller, Tamir and Sontag, Eduardo D.",
	editor="Petit, Nicolas",
	title="Checkable Conditions for Contraction After Small Transients in Time and Amplitude",
	booktitle="Feedback Stabilization of Controlled Dynamical Systems: In Honor of {Laurent} {Praly}",
	year="2017",
	publisher="Springer International Publishing",
	address="Cham, Switzerland",
	pages="279-305",
}
@ARTICLE{Shalem2008,
  author =       {O. Shalem and O. Dahan and M. Levo and M.R. Martinez and I. Furman and E. Segal and Y. Pilpel},
  title =        {Transient transcriptional responses to stress are generated by
                         opposing effects of {mRNA} production and degradation},
  journal =      {Mol. Syst. Biol.},
  year =         {2008},
  volume =       {4},
  number =       {223},
}


@ARTICLE{ Kawaguchi ,
  author =       {R. Kawaguchi and T.  Girke  and   E. A.  Bray   and   J.  Bailey-Serres},
  title =        {Differential m{RNA}
   translation contributes to gene regulation under
   non-stress and dehydration stress conditions in {A}rabidopsis thaliana},
  journal =      {Plant J.},
  year =         {2004},
  volume =       {38},
  number =       {5},
  pages =        {823-839},
}


@ARTICLE{Lu2007,
  author =       {P. Lu and C. Vogel and R. Wang and X. Yao and E.M. Marcotte},
  title =        {Absolute protein expression profiling estimates the relative contributions of transcriptional and translational regulation},
  journal =      {Nat. Biotechnol.},
  year =         {2007},
  volume =       {25},
  number =       {1},
  pages =        {117-124},

}


@ARTICLE{Weissman2006,
  author =       {J..R.	Newman and S. Ghaemmaghami and J. Ihmels and D.K. Breslow and M. Noble and J.L. DeRisi and J.S. Weissman},
  title =        {Single-cell proteomic analysis of S. cerevisiae reveals the architecture of biological noise},
  journal =      {Nature},
  year =         {2006},
  volume =       {441},
  number =       {7095},
  pages =        {840-846},
}


@ARTICLE{Ghaemmaghami2003,
  author =       {S. Ghaemmaghami and W.K. Huh and K. Bower and R.W. Howson and A. Belle and N. Dephoure and E.K. O'Shea and J.S. Weissman},
  title =        {Global analysis of protein expression in yeast},
  journal =      {Nature},
  year =         {2003},
  volume =       {425},
  number =       {6959},
  pages =        {737-741},
}


@article{ribosome_profiling_rev,
author={N. T. Ingolia},
title={Ribosome profiling: new views of translation, from single codons to genome scale},
journal={Nat. Rev. Genet.},
volume={15}, number={3}, pages={205-213},year={2014},
}


@article{Wilson01052012,
author = {D. N. Wilson  and J. H.  {Doudna Cate}},
title = {The Structure and Function of the Eukaryotic Ribosome},
volume = {4},
number = {5},
year = {2012},
journal = {Cold Spring Harbor Perspectives in Biology}
}


@ARTICLE{Ferrari2007,
  AUTHOR =       {P. A. Ferrari   and J. B. Martin},
  TITLE =        {Stationary distributions of multi-type totally asymmetric exclusion processes},
  JOURNAL =      {Annals of Probability},
  YEAR =         {2007},
  volume =       {35},
  skipnumber =       {3},
  pages =        {807-832},
}


@article{TASEP_feed2009,
title={Feedback and fluctuations in a totally asymmetric simple exclusion process with finite resources},
author={L. J. Cook and R. K. P. Zia},
journal=JSTATM,
year={2009},
pages={P02012},
}


@ARTICLE{Cannarozzi2010,
  AUTHOR =       {G. Cannarozzi and N. N. Schraudolph and M. Faty and P. von Rohr and M.
                         T. Friberg and A. C. Roth and P. Gonnet and G. Gonnet and Y. Barral},

  TITLE =        {A role for codon order in translation dynamics},
  JOURNAL =      {Cell},
  YEAR =         {2010},
  volume =       {141},
  number =       {2},
  pages =        {355-367},
}


@article{mans_2000,
title="Restricted permutations, continued fractions, and {C}hebyshev polynomials",
author="T. Mansour and A. Vainshtein",
journal="Electronic J. Combinatorics",
volume="7",
year="2000",
pages="R17",
}


@article{coppel74,
title={A Generalization of an Inequality of {C}oppel},
author={G. A. Hewer},
journal={Proc.  American Mathematical Society},
volume={44}, year={1974},pages={151-156},
}


@ARTICLE{mono_chem_2007,
  AUTHOR =       {P. De Leenheer and D. Angeli and E. D. Sontag},
  TITLE =        {Monotone Chemical Reaction Networks},
  JOURNAL =      {J. Mathematical Chemistry},
  YEAR =         {2007},
  volume =       {41},
  pages =        {295-314},
}


@ARTICLE{Schwanhausser2011,
  author =       {B. Schwanhausser and D. Busse and N. Li and G. Dittmar
   and J. Schuchhardt and J. Wolf and W. Chen and M. Selbach},
  title =        {Global quantification of mammalian gene expression control},
  journal =      {Nature},
  year =         {2011},
  volume =       {473},
  number =       {7347},
  pages =        {337-42},
}


 


@ARTICLE{Arava2003,
  AUTHOR =       {Arava, Y.  and Wang, Y. and Storey, J. D. and Liu, C. L. and Brown, P. O. and Herschlag, D.},
  TITLE =        {Genome-wide analysis of {mRNA} translation profiles in
   {S}accharomyces cerevisiae},
  JOURNAL =      PNAS,
  YEAR =         {2003},
  volume =       {100},
  number =       {7},
  pages =        {3889-3894},
}


@BOOK{Cooper2000,
  AUTHOR =       {Cooper, G.M.},
  TITLE =        {The Cell -- A Molecular Approach},
  YEAR =         {2000},
  address =      {Boston University Sunderland (MA): Sinauer Associates},
  edition =      {2nd edition},
}


@ARTICLE{Cramer2001,
  AUTHOR =       {Cramer, P. and Bushnell, DA. and Kornberg, RD.},
  TITLE =        {Structural basis of transcription: {RNA} polymerase {II} at 2.8 angstrom resolution},
  JOURNAL =      {Science},
  YEAR =         {2001},
  volume =       {292},
  number =       {5523},
  pages =        {1863-76},
}


@ARTICLE{Dittmar2006,
  AUTHOR =       {Dittmar, KA. and Goodenbour, JM. and Pan T.},
  TITLE =        {Tissue-specific differences in human transfer {RNA} expression},
  JOURNAL =      {PLOS Genet.},
  YEAR =         {2006},
  volume =       {2},
  number =       {12},
}


@ARTICLE{Welch2009,
  AUTHOR =       {Welch, M. and Govindarajan, S. and Ness, J.E. and
   Villalobos, A. and Gurney, A. and Minshull, J. and Gustafsson C.},
  TITLE =        {Design parameters to control synthetic gene expression in {Escherichia} coli},
  JOURNAL =      PLOS1,
  YEAR =         {2009},
  volume =       {4},pages={e7002},

  number =       {9},
}


@ARTICLE{Chu2011,
  AUTHOR =       {Chu, D. and Barnes, D. J. and von der Haar, T.},
  TITLE =        {The role of {tRNA} and ribosome competition in coupling the expression of different {mRNA}s in Saccharomyces cerevisiae},
  JOURNAL =      {Nucleic Acids Res.},
  YEAR =         {2011},
  volume =       {39},
  number =       {15},
  pages =        {6705-6714},
}


@ARTICLE{Qian2012,
  AUTHOR =       {Qian, W and Yang, JR. and Pearson, NM. and Maclean, C. and Zhang, J.},
  TITLE =        {Balanced codon usage optimizes eukaryotic translational efficiency},
  JOURNAL =      {PLOS Genet.},
  YEAR =         {2012},
  volume =       {8}, pages={e1002603},
  number =       {3},
}


@ARTICLE{Tuller2010b,
  AUTHOR =       {Tuller, T. and Waldman, Y. Y. and Kupiec, M. and Ruppin, E.},
  TITLE =        {Translation efficiency is determined by both codon bias and folding energy},
  JOURNAL =      PNAS,
  YEAR =         {2010},
  volume =       {107},
  number =       {8},
  pages =        {3645-50},
}


 @article{PhysRevLett.102.198104,
  title = {Queueing Phase Transition: Theory of Translation},
  author = {Romano, M. Carmen and Thiel, Marco and Stansfield, Ian and
Grebogi, Celso},
  journal = {Phys. Rev. Lett.},
  volume = {102},
  issue = {19},
  pages = {198104},
  year = {2009},
}


@article{mcs_angeli_2003,
title={Monotone control systems},
journal=TAC,
year={2003},
author={D. Angeli and E. D. Sontag},
volume={48},
pages={1684-1698},
}

@ARTICLE{Chowdhury2005,
  AUTHOR =       {D. Chowdhury and A. Schadschneider and K. Nishinari},
  TITLE =        {Physics of transport and traffic phenomena in biology: from molecular motors and cells to organisms},
  JOURNAL =      {Physics of Life Reviews},
  YEAR =         {2005},
  pages =        {318-352},
}

@article{traffic_TASEP_99,
title={Vehicular Traffic: A System of Interacting Particles Driven Far From Equilibrium},
author={D. Chowdhury and L. Santen and A. Schadschneider},
journal={Curr. Sci.},
volume={77},
pages={411-419},
year={1999},
}


@article{Mole_motors_2003,
year={2003},
journal={J. Statistical Physics},
volume={113},
title={Traffic of Molecular Motors Through Tube-Like Compartments},
author={S. Klumpp  and R.  Lipowsky},
pages={233-268},
}


@article{surf_gro_tasep,
year={2011},
journal={J. Statistical Physics},
volume={144},
title={Finite Time Corrections in {KPZ} Growth Models},
author={P. L. Ferrari  and R. Frings},
pages={1123-1150},
}


@article{biomotor_2012,
journal={Phys. Rev. Lett.},
year={2012}, volume={20},
title={Distinct transport regimes for two elastically coupled molecular motors},
author={F. Berger and C.  Keller and S.  Klumpp and R. Lipowsky},
pages={208101-208105},
}

@article{traffic_asep_2009,
  title = {Traffic by multiple species of molecular motors},
  author = {Y. Chai  and S. Klumpp and M. J. I.  Muller  and R. Lipowsky},
  journal = {Phys. Rev. E},
  volume = {80},
  pages = {041928},
  year = {2009},
 }

@article{pop_traffic_2001,
  author={V. Popkov and L. Santen and A. Schadschneider and G. M. Schutz},
  title={Empirical evidence for a boundary-induced nonequilibrium phase transition},
  journal=J_PHYS_A,
  volume={34},
  skipnumber={6},
  pages={L45},
    year={2001},
    }
	
@article{rock_1993,

title={Lagrange Multipliers and Optimality},
author={R. T. Rockafellar},
journal={SIAM Review}, volume={35},
number={2},
year={1993},
pages={183-238},
}

@article{TASEP_motors,
title={Stepping and Crowding of Molecular Motors:

Statistical Kinetics from an Exclusion Process Perspective},
author={L. Ciandrini and
M. C.  Romano and A. Parmeggiani},
journal={Biophysical J.},
pages={1176-1184},
volume={107},
year={2014},
}


@book{Ingraham1996,
 editor =       {F. C. Neidhardt and
  J. L. Ingraham and K. B. Low and
   B. Magasanik and
    M. Schaechter and H. E. Umbarger},
 title =        {Escherichia coli and \mbox{Salmonella} typhimurium -
   Cellular and Molecular Biology},
  publisher =      {ASM Press},
  year =         {1996},
  volume =       {1},
}


@ARTICLE{Zenklusen2008,
  author =       {D. Zenklusen and D. R. Larson and R. H. Singer},
  title =        {Single-{RNA} counting reveals alternative modes of gene expression in yeast},
  journal =      {Nat. Struct. Mol. Biol.},
  year =         {2008},
  volume =       {15},
  pages =        {1263-1271},
}


@article {TASEP_tutorial_2011,
   author = {Zia, R.  K. P.  and Dong, J.. and Schmittmann, B.},
     title = {Modeling Translation in Protein Synthesis with \mbox{TASEP}:
      A Tutorial and Recent Developments},
   journal = {J. Statistical Physics},
   pages = {405-428},
   volume = {144},
   year = {2011}
}


@ARTICLE{Li2007,
  AUTHOR =       {S.W. Li and L. Feng and D.K. Niu},
  TITLE =        {Selection for the miniaturization of highly expressed genes},
  JOURNAL =      {Biochem Biophys Res Commun.},
  YEAR =         {2007},
  volume =       {360},
  number =       {3},
  pages =        {586-92},
}


 @ARTICLE{Eisenberg2003,
  AUTHOR =       {E. Eisenberg and E. Y. Levanon },
  TITLE =        {Human housekeeping genes are compact},
  JOURNAL =      {Trends Genet.},
  YEAR =         {2003},
  volume =       {19},
  number =       {7},
  pages =        {362-5},
}


@ARTICLE{Ingolia2009,
  AUTHOR =       {N. T. Ingolia and S. Ghaemmaghami and J. R. Newman and J. S. Weissman},
  TITLE =        {Genome-wide analysis in vivo of translation with
                  nucleotide resolution using ribosome profiling},
  JOURNAL =      {Science},
  YEAR =         {2009},
  volume =       {324},
  number =       {5924},
  pages =        {218-23},
}


@ARTICLE{Ingolia2011,
  AUTHOR =       {N. T. Ingolia and L.F. Lareau and J.S. Weissman},
  TITLE =        {Ribosome profiling of mouse embryonic stem cells reveals the complexity and dynamics of mammalian proteomes},
  JOURNAL =      {Cell},
  YEAR =         {2011},
  volume =       {147},
  number =       {4},
  pages =        {789-802},
}


@article{ElMikkawy2004669,
title = "On the inverse of a general tridiagonal matrix",
journal = "Applied Mathematics and Computation",
volume = "150",
pages = "669-679",
year = "2004",
author = "Moawwad E.A. El-Mikkawy",
}


@book{ogata_dc,
title={Discrete-Time Control Systems}, author={K. Ogata},
publisher={Pearson Education},year={1994},
}

@book{wall_contin_frac,
publisher={Chelsea Publishing Company},
 year={1973},
title={Analytic Theory of Continued Fractions},
author={H. S. Wall},address={Bronx, NY},
}


@article{RFM_stability,
title = "Stability analysis of the ribosome flow model",
year = "2012",
author = "M. Margaliot and T. Tuller",
volume="9",pages="1545-1552",
journal=TCBB,
}


@article{HRFM_steady_state,
title = "On the Steady-State Distribution in the Homogeneous  Ribosome Flow Model",
year = "2012",
author = "M. Margaliot and T. Tuller",
volume="9",pages="1724-1736",
journal=TCBB,
}


@article{LOHMILLER1998683,
title = "On Contraction Analysis for Non-linear Systems",
journal = "Automatica",
volume = "34",
pages = "683-696",
year = "1998",
author = "W. Lohmiller and J.-J. E. Slotine",
}


@article{Spitzer1970246,
title = "Interaction of {M}arkov processes",
journal = "Adv. Math.",
volume = "5",
skipnumber = "2",
pages = "246-290",
year = "1970",
author = "F. Spitzer"
}


  	
@article{const_tasep_08,
  author={D. A. Adams and B. Schmittmann and R. K. P. Zia},
  title={Far-from-equilibrium transport with constrained resources},
  journal=JSTATM,
  volume={2008},
  skipnumber={06},
  pages={P06009},
  year={2008},
 }


@ARTICLE{Lakatos2003,
  AUTHOR =       {G. Lakatos and T. Chou},
  TITLE =        {Totally asymmetric exclusion processes with particles of arbitrary size},
  JOURNAL =      J_PHYS_A,
  YEAR =         {2003},
  volume =       {36},
  pages =        {20272041},
}


@article{RFM_entrain,
author={M.. Margaliot and E. D. Sontag and T. Tuller},
title={Entrainment to Periodic Initiation and Transition Rates in a Computational Model for Gene Translation},
journal=PLOS1,volume={9},number={5},pages={e96039},
 year={2014},
}


@article{entrain2011,
author={G. Russo and  M. di Bernardo  and E. D. Sontag},
title={Global entrainment of transcriptional systems to periodic inputs},
journal=PLOSCB,
volume={6}, pages={e1000739}, year={2010},
}


@ARTICLE{Baranov2002,

  AUTHOR =       {P. V. Baranov and R. F. Gesteland and J. F. Atkins},
  TITLE =        {Recoding: translational bifurcations in gene expression},
  JOURNAL =      {Gene},
  YEAR =         {2002},
  volume =       {286},
  number =       {5},
  pages =        {187-201},
}


@ARTICLE{Farabaugh1996,
  AUTHOR =       {P. J. Farabaugh},
  TITLE =        {Programmed translational frameshifting},
  JOURNAL =      {Annu. Rev. Genet.},
  YEAR =         {1996},
  volume =       {30},
  pages =        {50728},
}


@incollection {comp_res_baranov_2010,

   author = {A. E. Firth and M. Bekaert and P. V. Baranov},

   title = {Computational Resources for Studying Recoding},

   booktitle = {Recoding: Expansion of Decoding Rules Enriches Gene Expression},

   editor = {Atkins, John F. and Gesteland, Raymond F.},

   publisher = {Springer}, address={New York},

   pages = {435-461},

   volume = {24},

   series = {Nucleic Acids and Molecular Biology},

   year = {2010}

}


@ARTICLE{Churchman2011,
  AUTHOR =       {L. S. Churchman and J. S. Weissman},
  TITLE =        {Nascent transcript sequencing visualizes transcription at nucleotide resolution},
  JOURNAL =      {Nature},
  YEAR =         {2011},
  volume =       {469},
  number =       {7330},
  pages =        {368-73},
}


@article{tri_feed,
journal=SCL,year={2010},
volume={59}, pages={130-138},
title={Conditions for global stability of monotone tridiagonal systems with negative feedback},
author={L. Wang and  P. de Leenheer  and E. D. Sontag},
}


@article{mono_feed,
journal=TAC,year={2008},
volume={53}, pages={166-176},
 title={Oscillations in {I/O} monotone systems under negative feedback},
author={D. Angeli and E. D. Sontag},
}


@article{sontag_near_2007,
journal={Systems and Synthetic Biology},year={2007},
volume={1}, pages={59-87},
title={Monotone and near-monotone biochemical networks},
author={E. D. Sontag},
}


@book{vec_calc,
publisher={W. H. Freeman},
edition={5},
 year={2003},
title={Vector Calculus},
author={J. E. Marsden and A. Tromba},
}


@book{slotine,
title={Applied Nonlinear Control},
author={J. J. Slotine and W. Li},
year={1991},
publisher={Prentice Hall}, 
address={Englewood Cliffs, NJ}, 
}

@book{cheb_poly,
title={Chebyshev Polynomials},
author={J. C. Mason and D. C. Handscomb},
year={2003},
publisher={Chapman \& Hall},
address={Boca Raton, FL}, 
}

@article{fusco_oliva_1988, 
title={Jacobi matrices and transversality}, 
volume={109},   
number={3-4}, journal={Proc. Royal Society of Edinburgh: Section A Math.}, 
 author={Fusco, Giorgio and Oliva, Waldyr Muniz},
 year={1988},
 pages={231â243}
}


@book{hlsmith,
title={Monotone Dynamical Systems: An Introduction to the Theory of Competitive and
Cooperative Systems}, author={H. L. Smith}, series={Mathematical Surveys and Monographs},
volume={41}, publisher={Amer. Math. Soc.}, address={Providence, RI}, year={1995},
}


@article{diag_sca_smith,
title={Is My system of {ODEs} cooperative?},
author={H. L. Smith},
url={https://math.la.asu.edu/~halsmith/identifyMDS.pdf},year={2012},
}


@article{smillie,
 author={J. Smillie}, year={1984},
title={Competitive and cooperative tridiagonal systems of differential equations},
journal={SIAM J. Math. Anal.},
volume={15},  pages={530-534},
}


@article{Fiedler1999,
 author = {B. Fiedler and T.  Gedeon},
 title = {A {L}yapunov function for tridiagonal competitive-cooperative systems},
 journal = {SIAM J. Math. Anal.},
 volume = {30},
 issue = {3},
 sjipmonth = {March},
 year = {1999},
 pages = {469-478},
 }


@BOOK{Alberts2002,
  AUTHOR =       {B. Alberts AND A. Johnson AND J. Lewis AND M. Raff AND K. Roberts AND P. Walter},
  TITLE =        {Molecular Biology of the Cell},
  YEAR =         {2008},
  publisher={Garland Science},
  address =      {New York},
}


@article{reuveni,
    author = {S. Reuveni  AND I.  Meilijson  AND M.  Kupiec AND E. Ruppin  AND T. Tuller},
    journal = PLOSCB,
    title = {Genome-Scale Analysis of Translation Elongation with a Ribosome Flow Model},
    year = {2011},
    volume = {7},
    pages = {e1002127},
}


@ARTICLE{KimchiSarfaty2007,
  AUTHOR =       {C. Kimchi-Sarfaty AND J.M. Oh AND I.W. Kim AND Z.E. Sauna AND A.M. Calcagno AND S.V. Ambudkar AND M.M. Gottesman},
  TITLE =        {A "Silent" Polymorphism in the MDR1 Gene Changes Substrate Specificity.},
  JOURNAL =      {Science},
  YEAR =         {2007},
  volume =       {315},
  pages =        {525-528},
}


@ARTICLE{Coleman2008,
  AUTHOR =       {J.R. Coleman AND D. Papamichail AND S. Skiena AND B. Futcher AND E. Wimmer AND S. Mueller},
  TITLE =        {Virus attenuation by genome-scale changes in codon pair bias},
  JOURNAL =      {Science},
  YEAR =         {2008},
  volume =       {320},
  pages =        {1784-1787},
}


@ARTICLE{Bahir2009,
  AUTHOR =       {I. Bahir  and M. Fromer   and  Y. Prat   and M. Linial},
  TITLE =        {Viral adaptation to host: a proteome-based analysis
                   of codon usage and amino acid preferences},
  JOURNAL =      {Mol. Syst. Biol.},
  YEAR =         {2009},
  volume =       {5},
  pages =        {1-14},
}


@ARTICLE{Weringh2011,
  AUTHOR =       {A. van Weringh AND M. Ragonnet-Cronin AND E. Pranckeviciene AND M. Pavon-Eternod AND L. Kleiman AND X. Xia},
  TITLE =        {HIV-1 modulates the {tRNA} pool to improve translation efficiency},
  JOURNAL =      {Mol. Biol. Evol.},
  YEAR =         {2011},
  volume =       {28},
  number =       {6},
  pages =        {1827-34},
}


@ARTICLE{Vogel2010,
  AUTHOR =       {C. Vogel AND S. Abreu Rde AND D. Ko AND S.Y. Le AND B.A. Shapiro AND S.C. Burns AND D. Sandhu AND D.R. Boutz AND E.M. Marcotte AND L.O. Penalva},
  TITLE =        {Sequence signatures and {mRNA} concentration can explain two-thirds of protein abundance variation in a human cell line},
  JOURNAL =      {Mol. Syst. Biol.},
  YEAR =         {2010},
  volume =       {6},
  pages =        {1-9},
}


@ARTICLE{Pearson2011,
  AUTHOR =       {C.E. Pearson},
  TITLE =        {Repeat associated non-{ATG} translation initiation: one DNA, two transcripts, seven reading frames, potentially nine toxic entities},
  JOURNAL =      {PLOS Genet.},
  YEAR =         {2011},
  volume =       {7},
  pages =        {e1002018},
}


@ARTICLE{Comeron2006,
  AUTHOR =       {J.M. Comeron},
  TITLE =        {Weak selection and recent mutational changes influence polymorphic synonymous mutations in humans},
  JOURNAL =       PNAS,

  YEAR =         {2006},
  volume =       {103},
  pages =        {6940-6945},
}

%%%%%%%%%%%%%%


@ARTICLE{Gustafsson2004,
  AUTHOR =       {C. Gustafsson AND S.. Govindarajan AND J. Minshull},
  TITLE =        {Codon bias and heterologous protein expression},
  JOURNAL =      {Trends Biotechnol.},
  YEAR =         {2004},
  volume =       {22},
  pages =        {346-353},
}


@ARTICLE{Kudla2009,
  AUTHOR =       {G. Kudla AND A. W. Murray and D. Tollervey and J. B. Plotkin},
  TITLE =        {Coding-sequence determinants of gene expression in {E}scherichia coli},
  JOURNAL =      {Science},
  YEAR =         {2009},
  volume =       {324},
  pages =        {255-258},
}


@ARTICLE{Plotkin2010,
  AUTHOR =       {J. B. Plotkin AND G. Kudla},
  TITLE =        {Synonymous but not the same: the causes and consequences of codon bias},
  JOURNAL =      {Nat. Rev. Genet.},
  YEAR =         {2010},
  volume =       {12},
  pages =        {32-42},
}


@ARTICLE{Burgess-Brown2008,
  AUTHOR =       {N.A. Burgess-Brown AND S. Sharma AND F. Sobott AND C. Loenarz AND U. Oppermann AND O. Gileadi},
  TITLE =        {Codon optimization can improve expression of human genes in Escherichia coli: A multi-gene study},
  JOURNAL =      {Protein Expr. Purif.},
  YEAR =         {2008},
  volume =       {59},
  pages =        {94-102},
}


@ARTICLE{Supek2010,
  AUTHOR =       {F. Supek AND T. Smuc},
  TITLE =        { On relevance of codon usage to expression of synthetic and natural genes in Escherichia coli},
  JOURNAL =      {Genetics},
  YEAR =         {2010},
  volume =       {185},
  pages =        {1129-1134},
}


@ARTICLE{Kozak1986,
  AUTHOR =       {M. Kozak},
  TITLE =        {Point mutations define a sequence flanking the {AUG} initiator codon that modulates translation by eukaryotic ribosomes},
  JOURNAL =      {Cell},
  YEAR =         {1986},
  volume =       {44},
  number =       {2},
  pages =        {283-92},
}


%%%%%%%%%%%%%%%%%%%


@ARTICLE{Drummond2008,
  AUTHOR =       {D. A. Drummond and  C. O. Wilke},
  TITLE =        {Mistranslation-induced protein misfolding as a dominant constraint on coding-sequence evolution},
  JOURNAL =      {Cell},
  YEAR =         {2008},

  volume =       {134},
  pages =        {341-352},
}


@ARTICLE{Drummond2009,
  AUTHOR =       {D. A. Drummond and  C. O. Wilke},
  TITLE =        {The evolutionary consequences of erroneous protein synthesis},
  JOURNAL =      {Nat. Rev. Genet.},
  YEAR =         {2009},
  volume =       {10},
  pages =        {715-724},
}


@ARTICLE{Shah2010a,
  AUTHOR =       {P. Shah and M. A. Gilchrist},
  TITLE =        {Explaining complex codon usage patterns with selection for translational efficiency, mutation bias, and genetic drift},
  JOURNAL =      PNAS,
  YEAR =         {2010},
  volume =       {108},
  pages =        {10231-10236},
}


@ARTICLE{Shah2010b,
  AUTHOR =       {P. Shah AND M. A. Gilchrist},
  TITLE =        {Effect of correlated {tRNA} abundances on translation errors and evolution of codon usage bias},
  JOURNAL =      {PLOS Genet.},
  YEAR =         {2010},
  volume =       {6},  pages = {e1001128},
}


@ARTICLE{Plata2010,
  AUTHOR =       {G. Plata AND M. E. Gottesman AND D. Vitkup},
  TITLE =        {The rate of the molecular clock and the cost of gratuitous protein synthesis},
  JOURNAL =      {Genome. Biol.},
  YEAR =         {2010},
  volume =       {11},
  pages =        {R98},
}


@ARTICLE{Bulmer1991,
  AUTHOR =       {M. Bulmer},
  TITLE =        {The selection-mutation-drift theory of synonymous codon usage},
  JOURNAL =      {Genetics},
  YEAR =         {1991},
  volume =       {129},
  pages =        {897-907},
}


@ARTICLE{Sharp1987,
  AUTHOR =       {P.M. Sharp AND W.H. Li},
  TITLE =        {The rate of synonymous substitution in enterobacterial genes is inversely related to codon usage bias},
  JOURNAL =      {Mol. Biol. Evol.},
  YEAR =         {1987},
  volume =       {4},
  pages =        {222-230},
}


%%%%%%%%%%%%%%%%%%%%%%%


@ARTICLE{Danpure1995,
  AUTHOR =       {C.J. Danpure},
  TITLE =        {How can the products of a single gene be localized to more than one intracellular compartment},
  JOURNAL =      {Trends Cell Biol. },
  YEAR =         {1995},
  volume =       {5},
  pages =        {230-238},
}


@ARTICLE{Kochetov2006,
  AUTHOR =       {A.V. Kochetov},
  TITLE =        {Alternative translation start sites and their significance for eukaryotic proteomes},
  JOURNAL =      {Molecular Biology},
  YEAR =         {2006},
  volume =       {40},
  pages =        {705-712},
}


@ARTICLE{Schmeing2011,
  AUTHOR =       {T.M. Schmeing  AND R.M. Voorhees AND A.C. Kelley AND V. Ramakrishnan},
  TITLE =        {How mutations in {tRNA} distant from the anticodon affect the fidelity of decoding},
  JOURNAL =      {Nat. Struct. Mol. Biol.},
  YEAR =         {2011},
  volume =       {18},
  pages =        {432-436},
}


@ARTICLE{Warnecke2010,
  AUTHOR =       {T. Warnecke AND L.D. Hurst},
  TITLE =        {GroEL dependency affects codon usage--support for a critical role of misfolding in gene evolution},
  JOURNAL =      {Mol. Syst. Biol.},
  YEAR =         {2010},
  volume =       {6},
  pages =        {1-11},
}


@ARTICLE{Zhou2009,
  AUTHOR =       {T Zhou AND M. Weems AND C.O. Wilke},
  TITLE =        {Translationally optimal codons associate with structurally sensitive sites in proteins},
  JOURNAL =      {Mol. Biol. Evol.},
  YEAR =         {2009},
  volume =       {26},
  pages =        {1571-1580},
}


@ARTICLE{Zhang2010,
  AUTHOR =       {F. Zhang AND S. Saha AND S.A. Shabalina AND A. Kashina},
  TITLE =        {Differential Arginylation of Actin Isoforms Is Regulated by Coding Sequence-Dependent Degradation},
  JOURNAL =      {Science},
  YEAR =         {2010},
  volume =       {329},
  pages =        {1534-1537},
}


@ARTICLE{Fredrick2010,
  AUTHOR =       {K. Fredrick AND M. Ibba},
  TITLE =        {How the sequence of a gene can tune its translation},
  JOURNAL =      {Cell},
  YEAR =         {2010},
  volume =       {141},
  pages =        {227-229},
}


%%%%%%%%%%%%%%%%%%%%


@ARTICLE{Zhang2010b,
  AUTHOR =       {Z. Zhang AND L. Zhou AND L. Hu AND Y. Zhu AND H. Xu AND Y. Liu AND X. Chen AND X. Yi AND X. Kong AND L.D. Hurst},
  TITLE =        {Nonsense-mediated decay targets have multiple sequence-related features that can inhibit translation},
  JOURNAL =      {Mol. Syst. Biol.},
  YEAR =         {2010},
  volume =       {6},
  pages =        {1-9},
}


@ARTICLE{Man2007,
  AUTHOR =       {O. Man AND Y. Pilpel},
  TITLE =        {Differential translation efficiency of orthologous genes is involved in phenotypic divergence of yeast species},
  JOURNAL =      {Nat. Genet.},
  YEAR =         {2007},
  volume =       {39},
  pages =        {415-421},
}


@ARTICLE{Schmidt2007,
  AUTHOR =       {M.W. Schmidt AND A. Houseman AND A.R. Ivanov AND D.A. Wolf},
  TITLE =        {Comparative proteomic and transcriptomic profiling of the fission yeast Schizosaccharomyces pombe},
  JOURNAL =      {Mol. Syst. Biol.},
  YEAR =         {2007},
  volume =       {3},
  pages =        {79},
}


@ARTICLE{Elf2003,
  AUTHOR =       {J. Elf AND D. Nilsson AND T. Tenson AND M. Ehrenberg},
  TITLE =        {Selective charging of {tRNA} isoacceptors explains patterns of codon usage},
  JOURNAL =      {Science},
  YEAR =         {2003},
  volume =       {300},
  pages =        {1718-1722},
}


%%%%%%%%%%%%%%%


@ARTICLE{Chamary2006,
  AUTHOR =       {J.V. Chamary AND J.L. Parmley L.D. Hurst},


  TITLE =        {Hearing silence: non-neutral evolution at synonymous sites in mammals},
  JOURNAL =      {Nat. Rev. Genet.},
  YEAR =         {2006},
  volume =       {7},
  pages =        {98-108},
}

%%%%%%%%%%%

@article{parking_garage_2002,
  title = {Macroscopic car condensation in a parking garage},
  author = {M. Ha  and M. {den Nijs}},
  journal = {Phys. Rev. E},
  volume = {66},
  skipissue = {3},
  pages = {036118},
  year = {2002},
}


@ARTICLE{Shaw2003,
  AUTHOR =       {L. B. Shaw AND R. K. P.  Zia AND K. H. Lee},
  TITLE =        {Totally asymmetric exclusion process with extended objects:
   a model for protein synthesis},
  JOURNAL =      {Phys. Rev. E},
  YEAR =         {2003},
  volume =       {68},
  pages =        {021910},
}


@ARTICLE{Tuller2007,
  AUTHOR =       {T. Tuller AND M. Kupiec AND E. Ruppin},
  TITLE =        {Determinants of protein abundance and translation efficiency in S. cerevisiae. },
  JOURNAL =      PLOSCB,
  YEAR =         {2007},
  volume =       {3},
  pages =        {2510-2519},
}


@article{biorythm2011,
    author = {J. Tabak AND J. Rinzel AND R. Bertram},
    journal = PLOSCB,
    title = {Quantifying the Relative Contributions of Divisive and Subtractive Feedback to Rhythm Generation},
    year = {2011},
    volume = {7},
    pages = {e1001124},
}


@ARTICLE{TullerGB2011,
  AUTHOR =       {T. Tuller AND I. Veksler AND N. Gazit AND M. Kupiec AND E. Ruppin AND M. Ziv},
  TITLE =        {Composite Effects of Gene Determinants on the Translation Speed and Density of Ribosomes},
  JOURNAL =      {Genome Biol.},
  YEAR =         {2011},
  volume =       {12},
  number =       {11},
  pages =        {R110},
}


@ARTICLE{MacDonald1968,
  AUTHOR =       {C. T. MacDonald and  J. H.. Gibbs and A. C. Pipkin},
  TITLE =        {Kinetics of biopolymerization on nucleic acid templates},
  JOURNAL =      {Biopolymers},
  YEAR =         {1968},
  volume =       {6},
  pages =        {1-25},
}


@ARTICLE{Heinrich1980,
  AUTHOR =       {R. Heinrich AND T.A. Rapoport},
  TITLE =        {Mathematical modelling of translation of {mRNA} in eucaryotes; steady state, time-dependent processes and application to reticulocytes},
  JOURNAL =      JTB,
  YEAR =         {1980},
  volume =       {86},
  pages =        {279-313},
}


@ARTICLE{Dana2011,
  AUTHOR =       {A. Dana AND T. Tuller},
  TITLE =        { Efficient Manipulations of Synonymous Mutations for Controlling Translation
                   Rate--an Analytical Approach},
  JOURNAL =      {J. Comput. Biol.},

  YEAR =         {2012},
  volume =       {19},
  pages =        {200-231},
}


@BOOK{Spivak1994,
  AUTHOR =       {M. Spivak},
  TITLE =        {Calculus},
  PUBLISHER =    {Cambridge University Press, cop. },
  YEAR =         {1994},
  address =      {Houston, Texas},
}


@ARTICLE{Zhang1994,
  AUTHOR =       {S. Zhang AND E. Goldman AND G. Zubay},

  TITLE =        {Clustering of low usage codons and ribosome movement},
  JOURNAL =      JTB,
  YEAR =         {1994},
  volume =       {170},
  pages =        {339-54},
}


%%%%%%%%%%%%%%%%%%%%%%%%%%%%%%%


@MISC{morris11,
title={An explicit counterexample to the   {L}agarias-{W}ang
finiteness conjecture},
author={K. G. Hare and I.  D. Morris and  N.  Sidorov  and J. Theys},
year={2011},
url={http://www.mat.uniroma2.it/~morris/hmst21.pdf},
}


@article{Lagarias1995finiteness,
title = "The finiteness conjecture for the generalized spectral radius of a set of matrices",
author = "Jeffrey C. Lagarias and Yang Wang",
journal = LAA,
volume = "214",
year = "1995",
pages = "17-42",
}

@article{heaps,
journal={J. of the American Mathematical Society},
volume={15},pages={77-111},
title={ASYMPTOTIC HEIGHT OPTIMIZATION FOR TOPICAL {IFS},
{T}etris HEAPS, AND THE FINITENESS CONJECTURE},
author={T. Bousch  and J. Mairesse},year={2001},
}


@book{berk74,
title="Optimal Control Theory",
author="L. D. Berkovitz",
publisher="Springer-Verlag",
 year="1974",
}


@book{berman87,
title="Nonnegative Matrices in the Mathematical Sciences",
author="A. Berman and R. J. Plemmons",
publisher="SIAM",
 year=1987,

}

@book{farina2000,
title="Positive Linear Systems: Theory and Applications",
author="L. Farina and S. Rinaldi",
publisher="John Wiley",
 year=2000,
}


@book{berndt2000,
title="An Introduction to Symplectic Geometry",
author="R. Berndt",
publisher="American Mathematical Society",
 year=2000,
}

@book{matrix_math,
author={D. S. Bernstein},
title={Matrix Mathematics},
publisher={Princeton University Press},
year={2005},
skipaddress={Princeton, NJ},
}


@inproceedings{weiss-cdc57,
author = {E. Weiss and M. Margaliot},
title = {A Generalization of {Smillie's} Theorem on Strongly Cooperative Tridiagonal Systems},
booktitle = CDC57,
year = {2018},
address = {Miami Beach, FL},note=ta,
}


@inproceedings{gurvits-cdc03,
author = {L. Gurvits},
title = {What is the finiteness conjecture for linear continuous time
inclusions?},
booktitle = CDC42,
year = {2003},
pages = {1165--1169},
address = {Maui, HI},
}


@inproceedings{tpds_cdc,
  title={Analysis of nonlinear tridiagonal cooperative systems using totally positive linear differential systems},
  author={Margaliot, Michael and Sontag, Eduardo D},
  booktitle=CDC57,
  pages={3104--3109},
  year={2018},
}

@article{EYAL_RFMD1,
title = "Ribosome flow model with different site sizes",
 journal="{SIAM} J. Applied Dynamical Systems",
year = "2020",  pages="541-576", volume="19",
author = "Eyal Bar-Shalom and  Alexander Ovseevich and  Michael Margaliot",
}


@article{rfM_density_max,
    author = {Zarai, Yoram AND Margaliot, Michael AND Tuller, Tamir},
    journal = {PLOS ONE},
    title = {On the Ribosomal Density that Maximizes Protein Translation Rate},
    year = {2016},
    month = {11},
    volume = {11},
    pages = {1-26},
}


@inproceedings{bar-cdc,
author = {N. E. Barabanov},
title = {Lyapunov exponent and joint spectral radius: Some known
and new results},
booktitle = CDC44, address = {Seville, Spain},
year = {2005},
pages = {2332-2337},
}
@inproceedings{suss-nice,
author = {H. J. Sussmann},
title = {Reachability by means of nice controls},
booktitle = CDC26,
year = {1987},
pages = {1368-1373},
address = {Los Angeles, CA},
}

@Article{sussmann72,
title={The bang-bang problem for certain control systems in \textit{GL}$(n,\mathbb{R})$},
author={Sussmann, H. J.},
journal={SIAM J. Control},
volume=10,
year=1972,
pages={470-476}
}


@inproceedings{mason-shorten-cdc04,
author = {O. Mason and R. Shorten},
title = {Some results on the stability of positive switched linear systems},
booktitle = CDC43,
year = {2004},
pages = {4601-4606},
skipaddress = {},
}


@inproceedings{mason-shorten03,
author = {O. Mason and R. Shorten},
title = {A conjecture on the
 existence of common quadratic {L}yapunov functions for positive linear systems},
booktitle = ACC03,
year = {2003},
pages = {4469-4470},
address = {Denver, CO},
}


@inproceedings{shorten03,
author={R. Shorten and  D. Leith and  J. Foy and R. Kilduff},
title={Towards an analysis
and design framework for congestion control in communication
networks},
booktitle={Proc.  $12$th Yale Workshop on Adaptive
and Learning Systems},
year={2003}
}


@article{mordu2002,
title={Discrete Maximum Principle for Nonsmooth Optimal Control Problems with Delays},
journal={Cybernetics and Systems Analysis},
volume={38}, year={2002},
pages={255-264},
author={B. S. Mordukhovich and I. Shvartsman},
}

author={Koivo, A. J.},
title={A second--order variational method for discrete--time
optimal control problems},
journal={J.  Franklin Inst.},
volume={286}, year={1968},
}

@article{koivo,
author={Koivo, A. J.},
title={A second--order variational method for discrete--time
optimal control problems},
journal={J.  Franklin Inst.},
volume={286}, year={1968},
}

@article{morse03,
author={A. Jadbabaie and  J. Lin and A. S. Morse},
title={Coordination of groups of mobile autonomous agents using nearest neighbour rules},
journal=TAC, volume={48}, pages={988-1001}, year={2003},
}


@article{bou2002,
author={O. Bournez and M. S. Branicky},
title={The mortality problem for matrices of low dimensions},
journal={Theory Comput. Systems}, volume={35}, pages={433-448}, year={2002},
}


@article{tal1,
author={T. Monovich and M. Margaliot },
title={Analysis of  Discrete--Time Linear  Switched Systems: A Variational Approach},
note=s, year={2009},
}


@inproceedings{zho-hill05-l2,
author = {J. Zhao and D. J. Hill},
title = {On stability and ${L}_2$-gain for switched systems},
booktitle = CDC44, address = {Seville, Spain},
year = {2005},
pages = {3279-3284},
}


@inproceedings{zho-hill05-dis,
author = {J. Zhao and D. J. Hill},
title = {Dissipativity theory for switched systems},
booktitle = CDC44, address = {Seville, Spain},
year = {2005},
pages = {7003-7008},
}


@article{zhai05,
title={${L}_2$ gain analysis for switched systems with continuous-time and discrete-time subsystems},
author={G. Zhai and     H.  Lin and Y. Kim and  J. Imae and T. Kobayashi},
journal=IJC,
volume={78}, skipnumber={15}, pages={1198-1205},
year={2005}
}


@article{lin-zhai-ant,
title={Optimal Persistent Disturbance Attenuation Control for Linear Hybrid Systems},
author={Hai Lin  and  Guisheng Zhai  and  Panos J. Antsaklis},
journal={Nonlinear Analysis: Theory, Methods and Applications},
volume={65}, skipnumber={6}, pages={1231-1250},
year={2006}
}


@inproceedings{hespanha-morse99,
author = {J. P. Hespanha and A. S. Morse},
title = {Stability of switched systems with average dwell-time},
booktitle = CDC38,
year = {1999},
pages = {2655-2660},
}


@book{smirnov,
title={Introduction to the Theory of Differential Inclusions},
author={G. V. Smirnov},
publisher="American Mathematical Society",
year= "2002",
}

@article{bara93,
title="On the {A}izerman problem for third-order nonstationary systems",
author="N. E. Barabanov",
year="1993", journal=DFE,
volume="29", pages="1439-1448",
skipnumber="10",
}

@article{branicky1998,
title={A unified framework for hybrid control: Model
and optimal control theory},
author={M. S. Branicky and V. S.  Borkar and S. K.  Mitter},
 journal=TAC,
year={1998}, volume={43}, pages={31-45},
skipnumber={1},
}


@article{van_der1992,
title={${L}_2$-gain analysis of nonlinear systems and
nonlinear state feedback ${H}_\infty$ control},
author={A. J. van der Schaft},
journal=TAC,
year={1992}, volume={37}, pages={770-784},
skipnumber={6},
}


@article{xu_ant04,
title={ Optimal control of switched systems based on parameterization of the switching instants},
author={X. Xu and P. J. Antsaklis},
journal=TAC,
year={2004}, volume={49}, pages={2-16},
skipnumber={1},
}


@article{decarloetal,
title={Perspectives and results on the stability and stabilizability
of hybrid systems},
author={R. DeCarlo and M.S. Branicky and S. Pettersson and B. Lennartson},
journal=IEEE,
volume={88}, skipnumber={2}, pages={1069--1082},
year={2000}
}


@article{bengea,
title={Optimal control of switching systems},
author={S. C. Bengea and  R. A. DeCarlo},
journal={Automatica},
volume={41}, pages={11-27},
year={2005}
}


@article{branicky_98, author={M. S. Branicky},
title={Multiple {L}yapunov functions
and other analysis tools for switched and hybrid systems},
journal=TAC, volume={43}, pages={475-482}, year={1998}
}


@article{egerstedt06,
title={Transition-time optimization for switched-mode dynamical systems},
author={M. Egerstedt  and  Y. Wardi  and H. Axelsson},
journal=TAC,
year={2006}, volume={51}, pages={110-115},
skipnumber={1},
}


@article{puk1998,
title="Optimal bilinear systems on the plane",
author="A. V. Pukhlikov",
year="1998", journal=DFE,
skipnumber="11", volume="34", pages="1516-1521",
}


@article{bara88,
title="Absolute characteristic exponent of a class of linear nonstationary systems of differential equations",
author="N. E. Barabanov",
year="1988", journal="Siberian Mathematical J.",

volume="29", pages="521-530",
skipnumber="4",
}

@article{bara89,
title="Method for the computation of
the {L}yapunov exponent of a differential inclusion",
author="N. E. Barabanov",
year="1989", journal=ARC,
volume="50", pages="475-479",
skipnumber="4",
}


@article{Tretyak,
title="Chronological Calculus, High-Order Necessary Conditions for Optimality, and Perturbation Methods",
author="A. I. Tretyak",
year="1998", journal="J. Dynamical and Control Systems",
volume="4", pages="77-126",
}


@article{andru85,
title="A class of absolutely stable nonlinear nonstationary systems",
author="V. V. Andrusevich",
year="1985", journal=ARC,
volume="11", pages="26-33",

}


@article{kozy07,
title="Structure of extremal trajectories of discrete linear systems
and the finiteness conjecture",
author="V. Kozyakin",
year="2007", journal=ARC,
volume="68", pages="174-209",
}

@inproceedings{kozy05,
author = {V. Kozyakin},
title = {A Dynamical Systems Construction of a Counterexample to the Finiteness Conjecture},
booktitle = CDC44,
year = {2005},

pages = {2338-2343},
address = {Seville, Spain},
}


@article{bara_dis_88_part1,
title="The {L}yapunov indicator  of discrete inclusions: Part {I}",
author="N. E. Barabanov",
year="1988", journal=ARC,
volume="2", pages="40-46",
}


@article{bara_dis_88_part2,
title="The {L}yapunov indicator  of discrete inclusions: Part {II}",
author="N. E. Barabanov",
year="1988", journal=ARC,
volume="3", pages="24-29",
}


@article{molchanov79,
title="Absolute stability of sampled-data
systems with several nonstationary nonlinear elements",
author="A.  P. Molchanov",
year="1979", journal=ARC,
volume="3", pages="43-51",
}

@article{pyat_dis_70,
title="Absolute stability of nonlinear pulse systems
 with nonstationary nonlinearity",
author="E. S. Pyatnitskii",
year="1970", journal=ARC,
volume="8", pages="50-58",
}


@article{bara88kal,
title="On the {K}alman problem",
author="N. E. Barabanov",
year="1988", journal="Siberian Mathematical J.",
volume="29", pages="333-341",
skipnumber="3",
}


@article{levin62,
title="On the stability of solutions of equations of second order",
author="A. Yu. Levin",
year="1961", journal="Soviet Mathematics",
volume="2", pages="1642-1646",
skipnote="Translation of Doklady, Academy of Sciences of the USSR"
}


@article{short-mason05,
author="R. Shorten and D. Leith and J. Foy and R. Kilduff",
title="Analysis and design of {AIMD} congestion control algorithms
in communication networks",
journal="Automatica",
volume="41",
year="2005",pages={725-730},
}
@misc{angeli-switched-note-scl,
author="D. Angeli",
title="A note on stability of arbitrarily switched homogeneous systems",
note="Preprint",
year=1999
}


@article{jong04,
title="Qualitative simulation of genetic regulatory
networks using piecewise-linear models",
journal="Bull.       Math.        Biol.",
volume="66",
skipnumber="2",
year="2004",pages={301-340}, author="H. de Jong and
 J.-L. Gouze and C. Hernandez and  M. Page and  T. Sari and J. Geiselmann",
}

@incollection{john04,
title="Modeling of hybrid systems",
author="K. H. Johansson and  J. Lygeros and S. Sastry",
booktitle="Encyclopedia of Life Support Systems {(EOLSS)}", year="2004",

url="www.s3.kth.se/~kallej/papers/eolss_6.43.28.1.pdf ",
editor="H. Unbehauen",
}

@article{zhai01,
author="G. Zhai and B. Hu and K. Yasuda and A. N. Michel",
title="Disturbance attenuation properties of time-controlled switched systems",
journal="J.  Franklin Inst.",
volume="338", year="2001",pages="765-779", skipnumber="7",
}


@article{chase93,
author="C. Chase  and J. Serrano and P. Ramadge",
title="Periodicity and chaos from switched flow systems:
Contrasting examples of discretely
controlled continuous systems",
journal=TAC,
volume="38", year="1993",pages="70-83"
}

@article{counter,
author="M. Margaliot",
title="A Counterexample to a Conjecture of {G}urvits on Switched Systems",
journal=TAC,
volume="52", year="2007",pages="1123-1126",
}


This paper appears in: Automatic Control, IEEE Transactions on
Publication Date: June 2007
Volume: 52,  Issue: 6
On page(s): 1123-1126


@techreport{hespanha_rms_tech_rep,
author = {J. P.  Hespanha},
title = {Root-Mean-Square Gains of Switched Linear Systems},
institution = {Dept.~of Electrical and Computer Eng., University of California},
address = {Santa Barbara},
year = 2002,
month = sep,
url = "http://www.ece.ucsb.edu/~hespanha/published",
}


@ARTICLE{olshevsky_minimal_control,

author={Olshevsky, A.},
journal={IEEE Trans. Control of Network Systems},
title={Minimal Controllability Problems},
year={2014},
volume={1},
number={3},
pages={249-258},
}


@article{barabashi_nature,
author={ Yang-Yu
Liu and
   Jean-Jacques Slotine and
 Albert-Laszlo Barabasi},
 title={Controllability of complex networks},
journal={Nature},year={2011},
volume={473},
issue={7346},
pages={167-173},
}

@article{hespanha_rms_2003,
author="J. P. Hespanha",
title="Root-mean-square gains of switched linear systems",
journal=TAC,
volume="48", year="2003",pages="2040-2045"
}


@inproceedings{mar-hespanha-rmsCDC,
author="M. Margaliot and J. P. Hespanha",
title="Root-mean-square gains of switched linear systems:
 A variational  approach",
booktitle = CDC46,
year = {2007},
pages = {3221-3226},
address = {New Orleans, LA},
}

@article{mar-hespanha-rms,
author="M. Margaliot and J. P. Hespanha",
title="Root-mean-square gains of switched linear systems:
 A variational  approach", journal="Automatica",
 year="2008", note=ta, url="www.eng.tau.ac.il/~michaelm",
}

@article{mar-bra-full,
author="M. Margaliot and M. S. Branicky",
title="Nice Reachability
for Planar Bilinear Control Systems
with Applications to Planar Linear Switched Systems",
 journal=TAC,
 year="2009", volume="54", pages="1430-1435",
}

@article{lior,
author="L.  Fainshil and  M. Margaliot
  and  P. Chigansky",
title="On the Stability of Positive Linear Switched Systems
Under Arbitrary Switching Laws",
 journal=TAC,
 year="2009", volume="54", pages="897-899",
}


@article{hesp2004,
author="J. P. Hespanha",
title="Uniform stability of switched linear systems: Extensions of {L}a{S}alle's invariance principle",
journal=TAC, skinpnumber="4",
volume="49", year="2004",pages="470-482",
}


@article{walther01,
author="U. Walther and T. T.  Georgiou and A. Tannenbaum",
title="On the computation of switching surfaces
 in optimal control: A {G}robner basis approach",
journal=TAC,
volume="46", year="2001",pages="534-540"
}


@article{shorten,
title="Stability criteria for switched and hybrid systems",
author="R. Shorten and F. Wirth and O. Mason and K. Wulff and C.. King",
year="2007",
journal="SIAM Review",
volume="49",
skipnumber="4",
pages = "545-592",
}


@Book{colonius,

   author ={F. Colonius and W. Kliemann and L. Grune},
   title =   {The Dynamics of Control},
   year =   {2000}, publisher ={Birkh{\"a}user},

}

@Book{cormen,
   author =   {T. H. Cormen and C. E. Leiserson and
R. L. Rivest and C. Stein},
   title =   {Introduction to Algorithms},
   publisher = {MIT Press},
   year =   {2001}, edition={2nd},
   skipaddress = {Cambridge, Massachusetts},
}

@Book{young2000,
   author =   {L. C. Young},
   title =   {Lectures on the Calculus
 of Variations and Optimal Control Theory},
   publisher = {American  Mathematical Society},
   year =   {2000},
   address = {Providence, Rhode Island},
}

@Article{strichartz87,
title={The {C}ampbell-{B}aker-{H}ausdorff-{D}ynkin formula and solutions of differential
equations},
author={R. S. Strichartz},
journal={J. Functional Analysis},
volume={72},
year={1987},
pages={320-345},
}

@Article{pol,
title={On absolute stability analysis by polyhedral {L}yapunov functions},
author={Polanski, A.},
journal={Automatica},
volume={36},
year={2000},

pages={573-578},
}

%%%%%%%%%%%%%%%%%%%%%%%%%%%%%%%%%%%%%%%%%%%%%%%%%%%%%%%%%%


@book{olver86,
title=" Applications of Lie Groups to Differential Equations",
author="P. J. Olver",
publisher="Springer-Verlag", address="New York",
year="1986",
series="Graduate Texts in Mathematics", volume="107",
}

@book{jack90,
title="Perspectives of Nonlinear Dynamics", volume="2",
author="E. A. Jackson",
publisher="Cambridge University Press",
year="1990"
}

@book{bardi,
title="Optimal Control and Viscosity Solutions of
Hamilton-Jacobi-Bellman Equations",
author="M.  Bardi and I. Capuzzo-Dolcetta",
publisher="Birkh{\"a}user",
year="1997"
}

@incollection{mayne73,
author="R. W. Brockett",
title="Lie algebras and {L}ie groups in control theory",
booktitle="Geometric Methods in System Theory",
editor="D. Q. Mayne and R. W. Brockett",
publisher="D. Reidel Publishing Company",
year=1973,
pages="43-82",
}


@book{agrachev-book,
title="Control Theory From The Geometric Viewpoint",
author="A. A. Agrachev and Y. L. Sachkov",
series="Encyclopedia of Mathematical Sciences",
volume="87",
publisher="Springer-Verlag",
year="2004"
}

@Book{gol,
   author =   "Goldstein, H.",
   title =   "Classical Mechanics",
   publisher =   "Addison-Wesley",
   year =   "1980",
   skipaddress = "",
}
@Book{vid,
   author =   "Vidyasagar, M.",
   title =   "Nonlinear Systems Analysis",
   publisher =   "Prentice Hall",
   year =   "1978",
  address = "Englewood Cliffs, NJ",
}

@book{matveev-savkin-book,
author="A. S. Matveev and A. V. Savkin",
title="Qualitative Theory of Hybrid Dynamical Systems",
publisher="Birkh{\"a}user",
year=2000
}

@Book{hall,
   author =   "Hall, B. C.",
   title =   "Lie Groups, Lie Algebras, and Representations:
 An Elementary Introduction",
   publisher =   "Springer",
   year =   "2003",
   skipddress = "",
}
@Book{erickson,
   author =   "Erickson, R. W. and Maksimovic, D.",
   title =   "Fundamentals of Power Electronics",
   publisher =   "Kluwer",
   year =   "2001",
   skipaddress = "",
}


@book{boyd,
author="S. Boyd and L. {El Ghaoui} and E. Feron and V. Balakrishnan",
year=1994,
title="Linear Matrix Inequalities in System and Control Theory",
volume=15,
series="SIAM Studies in Applied Mathematics",
publisher="SIAM",
address="Philadelphia"
}

@article{cics2006,
journal=SCL,
year={2006}, 
volume={55},
pages={707-710},
title={Well-defined steady-state response does not imply {CICS}},
author={Eugene P. Ryan and Eduardo D. Sontag},
}


@article{Fang2015,
title={Non-hyperbolic minimal sets for tridiagonal competitive-cooperative systems},
author={Chun Fang and Mats Gyllenberg and Yi Wang},
journal={Proc. Amer. Math. Soc.},
volume={143}, year={2015}, pages={3063-3074},
}


@book{karlin_tp,
title={Total Positivity},volume={1},
author={Samuel Karlin},
publisher={Stanford University Press}, address={Stanford, CA},
year={1968},
}

@article{fgwang2013,
journal={SIAM J. Math. Anal.},
year={2013}, pages={2477-2498},volume={45},number={4},
title={FLOQUET BUNDLES FOR TRIDIAGONAL
COMPETITIVE-COOPERATIVE SYSTEMS AND THE DYNAMICS
OF TIME-RECURRENT SYSTEMS},
author={C. Fang and M. Gyllenberg and Y. Wang},
}

@book{liberzon_book,
author="D. Liberzon",
title="Switching in Systems and Control",
publisher="Birkh{\"a}user",
year=2003,
address="Basel, Switzerland", 
}

@book{aubin-viability,
author="J.-P. Aubin",
title=" Viability Theory",
 publisher="Birkh{\"a}user",
year="1991",
skipaddress="",
}

@Article{hermes78,
   author =   {Hermes, H.},
   title =   {Lie Algebras of Vector Fields
and Local Approximation of Attainable Sets},
   journal = SIAM,
   year =   {1978},
   volume = {16},
   issue={5},
   pages =  "715-727",
}


@Article{bianchini_90,
   author =   {R. M. Bianchini and G. Stefani},
   title =   {Graded approximations and controllability along a trajectory},
   journal = SIAM,
   year =   {1990},
   volume = {28},
   pages =  {903-924},
}


@Article{hermes82,
   author =   "Hermes, H.",
   title =   "Control systems which generate decomposable {L}ie algebras",
   journal = JDE,
   year =   "1982",
   volume = "44",
   pages =  "166-187",
}
@incollection{suss-levinson,
author="Sussmann, H. J. and Nohel, J. A.",
title="Commentary on {N}orman {L}evinson's paper `{M}inimax,
 {L}iapunov and {B}ang-{B}ang'",
booktitle="Selected Papers of Norman Levinson",
volume="2",
editor="Nohel, J. A. and Sattinger, D. H.",
publisher="Birkh{\"a}user",
year=1998,
pages="463-475",
}

@article{arts80,
author="Z. Artstein",
title=" Discrete and Continuous
Bang-Bang and Facial Spaces or: Look for the Extreme Points",
journal="SIAM Review",
volume="22",
pages="172-185",
year="1980",
}

@article{hermes91,
author="H. Hermes",
title="Nilpotent and high-order approximations of vector field systems",
journal="SIAM Review",
volume="33",
pages="238-264",
year="1991"
}

@book{elliott,
title={Bilinear Control Systems: Matrices in Action},
author={D. L. Elliott},
year={2009},
publisher={Springer},
}


@Article{mar-open2001,
title={Some nonlinear optimal control
  problems with closed-form solutions},
author={M. Margaliot and G. Langholz},
year={2001}, journal={Int. J. Robust Nonlinear Control},
volume={11},pages={1365-1374},
}


@article{yoav,
title={Third-order nilpotency,
finite switchings and asymptotic stability},
author={Y. Sharon and M. Margaliot},
year={2007}, journal=JDE, volume={233},
pages={136-150},
}


@Article{yfou04,
title={A numerical technique
for the stability analysis of linear switched systems},
author={Yfoulis, C. A.  and Shorten, R.},
journal=IJC,
volume={77},
year={2004},
pages={1019-1039},
}
@article{gurvits-shorten-mason07,
   author =   "L. Gurvits and R. Shorten and O. Mason",
   title =   "On the stability of switched positive linear systems",
journal=TAC,   year="2007",
pages="1099-1103", volume="52",
skipissue="6",
}


@article{yfou,
   author =   "Margaliot, M. and  Yfoulis, C.",
   title =   "Absolute Stability
of Third-Order Systems:   A Numerical Algorithm",
journal="Automatica",
volume="42",
skipnumber="10",year="2006", pages={1705-1711},
}

@book{yfbook,
author="C. A. Yfoulis",
title="Stabilisation of Nonlinear Systems: The Piecewise Linear Approach",
publisher="Research Studies Press",
year=2002,
}

@Article{bellman,
   author =   "Bellman, R. and Bucy, R.",
   title =   "Asymptotic control theory",
   journal = SIAM,
   number = "1",
   year =   "1964",
   volume = "2",
   pages =  "11-18",
}


@book{aubin-cellina-inclusions-book,
author="J.-P. Aubin and A. Cellina",
title="Differential Inclusions: Set-Valued Maps and
Viability Theory",
 publisher="Springer",
address="Berlin", year=1984

}


@article{lin-sontag-wang-converse,
author="Y. Lin and E. D. Sontag and Y. Wang",
title="A smooth converse {L}yapunov theorem for robust stability",
journal=SIAM,
volume=34,
pages="124-160",
year=1996
}


@article{narendra-balakrishnan-commuting,
author="K. S. Narendra and J. Balakrishnan",
title="A common {L}yapunov function for stable {LTI} systems
with commuting {$A$}-matrices",
journal=TAC, volume=39, year=1994,
 pages="2469-2471"
 }

@Article{alur,
title={Modeling and analyzing
    biomolecular networks},
author={R. Alur et al.},
journal={Computing in Science and Engineering},
volume={4},
number={1},
year={2002},
pages={20-31},
}

@Article{mol,
title={Criteria of asymptotic stability of differential
 and difference inclusions encountered in control theory},
          author={Molchanov, A. P. and Pyatnitskiy, Ye. S.},
year={1989},
journal=SCL,
volume={13},
pages={59-64},
}


@Article{soko82,
title={Numerical methods of {L}yapunov function
construction and their application to the absolute stability problem},
          author={Pyatnitskii, E. S. and Sokorodinskii, V. I.},
year={1982},
journal=SCL,
volume={2},skipnunber={2},
pages={130-135},
}

@inproceedings{gurvits07,
author = {L. Gurvits and A. Olshevsky},
title = {Stability testing of matrix polytopes},
booktitle = ECC7,
year = {2007},
pages = {3919-3925},
}
@inproceedings{gurvits_quantom,
author = {L. Gurvits},
title = {Controllabilities and stabilities of switched systems (with applications to the quantum systems)},
booktitle = MTNS02,
year = {2002},
skippages = {},
}


@Article{ratmansky,
title={A simplification of the {A}grachev--{G}amkrelidze second-order variation for
 bang--bang controls},
author={I. Ratmansky and M. Margaliot},
journal=SCL,
 year={2010}, volume={59}, pages={25-32},
}


@article{gurvits-nilpotent-laa95,
author="L. Gurvits",
title="Stability of discrete linear inclusion",
journal=LAA,
volume="231", year="1995", pages="47-85"
}

@article{wirth-LAA,
author="F. Wirth",
title="The generalized spectral radius and extremal norms",
journal=LAA,
volume="342", year="2002", pages="17-40"
}


@article{valcher2007,
title={On the zero pattern properties and asymptotic behavior of
continuous-time positive system trajectories},
author={P.  Santesso and  M.  E.  Valcher},
journal=LAA,
volume={425}, year={2007}, pages={283-302},
}


@article{agrachev,
title="{L}ie-algebraic
stability criteria for
 switched systems",
author="A. A. Agrachev and D. Liberzon",
journal=SIAM,
volume=40,year=2001,pages="253-269"
}

@incollection{symplectic,
author="Agrachev, A. A. and Gamkrelidze, R. V.",
title="Symplectic geometry for optimal control",
booktitle="Nonlinear Controllability and Optimal Control",
editor="H. J. Sussmann",
publisher="Marcel Dekker",
address="New York",
year=1990,
pages="263-277",
}


@article{sussmann_plane_cinf,
title="The Structure of Time-Optimal
 Trajectories for Single-Input Systems in the Plane:
The ${C}^\infty$ Nonsingular Case",
author="H. J. Sussmann",
journal=SIAM,
volume=25, year=1987, pages="433-465",
}
  	
@article{nems2011,
  author={T. Chou and K. Mallick and R. K. P. Zia},
  title={Non-equilibrium statistical mechanics: from a paradigmatic model to biological transport},
  journal={Reports on Progress in Physics},
  volume={74},
  pages={116601},
  year={2011},
  }

	


@article{sussmann_plane_ana,
title="The Structure of Time-Optimal Trajectories for Single-Input Systems in the Plane: The General Real Analytic Case",
author="H. J. Sussmann",
journal=SIAM,
volume=25, year=1987, pages="868-904",
}

@article{sussmann_plane_syn,
title="Regular Synthesis for Time-Optimal Control of Single-Input
Real Analytic Systems in the Plane",
author="H. J. Sussmann",
journal=SIAM,
volume=25, year=1987, pages="1145-1162",
}

@book{TASEP_book,
title={Stochastic Transport in Complex Systems: From Molecules to Vehicles},
author={A. Schadschneider and D. Chowdhury and K. Nishinari},

year={2011},
publisher={Elsevier},
}

@article{bressan,
title="A high order test for optimality of bang-bang controls",
author="Bressan, A.",
journal=SIAM,
volume=23,year=1985,pages="38-48",
}


@article{krener,
title="The high order maximal principle and its applications to singular
extremals",
author="A. J. Krener",
journal=SIAM,
volume=15,year=1977,pages="256-293"
}


@article{mancilla-commuting-tac,
author="J. L. Mancilla-Aguilar",
title="A condition for the stability of switched nonlinear systems",
journal=TAC,
volume=45,
year=2000,
skip-month=nov,
pages="2077-2079"
}

@article{mcclamroch-kolmanovsky-ieeesurvey2,
author="N. H. McClamroch and I. Kolmanovsky",
title="Performance benefits of hybrid control
design for linear and nonlinear systems",
journal=IEEE,
volume=88,
skip-issue=7,
skip-month=jul,
year=2000,
pages="1083-1096"
}


@article{shim-...-siamcca-98,
author="H. Shim and D. J. Noh and J. H. Seo",
title="Common {L}yapunov function for exponentially stable nonlinear systems",
journal="J. Korean Institute of Electrical Engineers",
volume=11, skip-no=2, pages="108-111", year=2001,
note="(Presented at the \textit{4th SIAM Conf.\ on Control and its
Applications}, 1998)"
}

@article{linh-commuting,
author="L. Vu and D. Liberzon",
title="Common {L}yapunov functions for
families of commuting nonlinear systems",
journal=SCL, note=s, year=2003
}


@incollection{laff-suss,
author="Lafferriere, G. and Sussmann, H. J.",
title="Differential Geometric Approach to motion Planning",
booktitle="Nonholonomic Motion Planning",
editor="Li, Z. and Canny, J. F.",
publisher="Kluwer Academic Publishers",
year=1993,
pages="235-270"
}

@article{murray-sastry-tac-93,
author="R. M. Murray and S. S. Sastry",
title="Nonholonomic motion planning: Steering using sinusoids",
journal=TAC, volume=38, year=1993, pages="700-716"
}


@article{sussmann-bang-bang-siam-1979,
author="H. J. Sussmann",
title="A bang-bang theorem with bounds on the number of switchings",
journal=SIAM, volume=17, year=1979,
skip-no=5, pages="629-651"
}
@Article{xu,
title={Stabilization of second-order \mbox{LTI} switched systems},
author={Xu, X. and Antsaklis, P. J.},
year={2000},
journal=IJC,
volume={73},
number={14},
pages={1261-1279}
}

@periodical{nonlinearhyb2,
title={Nonlinear Analysis,
 Special Issue on Hybrid Systems and Applications (2)},
key={nonlinear},
year={2005},
volume={63},
number={3},
}
@periodical{nonlinearhyb1,
title={Nonlinear Analysis,
 Special Issue on Hybrid Systems and Applications},
key={nonlinear},
year={2005},
volume={62},
number={8},
}


@periodical{hsi-tcs-95,
title={Theoretical Computer Science,
 Special Issue on Hybrid Systems},
 editor={A. Pnueli and J. Sifakis},
key={Pnueli},
year={1995},
volume={138},
number={1},
}

@periodical{ant,
title={Proc. IEEE,        Special Issue on Hybrid
Systems: Theory and Applications},
 editor={Antsaklis, P. J.},
key={Antsaklis},
year={2000},
volume={88},
number={7},
}

@article{tasep_ants,
  title = {Trafficlike Collective Movement of Ants on Trails: Absence of a Jammed Phase},
  author = {A. John and  A. Schadschneider and D. Chowdhury and K. Nishinari},
  journal = {Phys. Rev. Lett.},
  volume = {102},
  pages = {108001},
  year = {2009},
}


@article{domain_wall_02,
year={2002},
 journal={J. Statistical Physics},
volume={106},
 title={The Asymmetric Exclusion Process Revisited: Fluctuations and Dynamics in the Domain Wall Picture},
  author={L. Santen  and  C. Appert},
pages={187-199},
}


@article{mixed_tasep,
  title = {Mixed population of competing totally asymmetric simple exclusion processes with a shared reservoir of particles},
  author = {P. Greulich and L. Ciandrini and  R. J. Allen and M. C. Romano},
  journal = {Phys. Rev. E},
  volume = {85},
   pages = {011142},
   year = {2012},
 }


@periodical{hsi-tac-98,
title={IEEE Trans. Automatic Control,    Special Issue on Hybrid
Control Systems},
 editor={P. J. Antsaklis and A. Nerode},
key={Antsaklis},
year={1998},
volume={43},
number={4},
}

@periodical{hsi-automatica-99,
title={Automatica,    Special Issue on Hybrid Systems},
 editor={A. S. Morse and C. C. Pantelides and
S. S. Sastry and J. M. Schumacher},
key={Morse},
year={1999},
volume={35},
number={3},
}


@Article{day,
title={A converse {L}yapunov theorem for a class of
dynamical systems which undergo switching},
author={Dayawansa, W. P. and Martin, C. F.},
year={1999},
journal=TAC,
volume={44},
number={4},
pages={751-760}
}


@Article{fil80,
title={Stability conditions in homogeneous systems
with arbitrary regime switching},
author={Filippov, A. F.},
year={1980},
journal=ARC,
volume={41},
pages={1078-1085},
number={8},
}

@article{pyat70,
title="Absolute stability of nonstationary nonlinear systems",
author="E. S. Pyatnitskii",
year="1970", journal=ARC,
volume="1", pages="5-15"
}

@article{pyat71,
title="Criterion for the absolute stability of second-order nonlinear
controlled systems with one nonlinear nonstationary element",
author="E. S. Pyatnitskii",
year="1971", journal=ARC,
volume="1", pages="5-16"
}


@article{rapo93a,
title="Existence of nonsmooth invariant functions on the
boundary of absolute stability domain of nonlinear nonstationary systems",
author="L. B. Rapoport",
year="1993", journal=ARC,
volume="54", pages="448-453"
}


@article{filippov-paper,
author="A. F. Filippov",
title="On certain questions in the theory of optimal control",
journal=SIAM, volume=1, year=1962,
skip-no=5, pages="76-84"
}


@article{zubov2001,
author="Camilli, F. and Grune, L. and Wirth, F.",
title="A generalization of {Z}ubov's method to perturbed systems",
journal=SIAM, volume=40, year=2001,
 pages="496-515"
}

@article{periodic_dis,
author="W. P. London  and J. A. Yorke",
year="1973",
title="Recurrent outbreaks of measles, chickenpox and mumps. {I.} {S}easonal variation in contact rates",
journal="Am. J. Epidemiol.", 
volume="98", 
number="6", 
pages="453–468",
}

@book{kiss_book,
author="I. Z. Kiss and J. Miller and P. L. Simon",
 title="Mathematics of Epidemics on Networks:
From Exact to Approximate Models",
 publisher="Springer",   year="2017",
}


@book{filippov-book,
author="A. F. Filippov", title="Differential Equations
 with Discontinuous Righthand Sides",
 publisher="Kluwer", address="Dordrecht", year=1988,
 skip-note="(Original Russian edition: \textit{Differentsial'nye
 Uravneniya s Razryvnoi Pravoi Chast'yu}, Nauka, Moscow, 1985)"
}

 @ARTICLE{RFMR,
author={Raveh, A. and Zarai, Y. and Margaliot, M. and Tuller, T.},
journal=TCBB,
title={Ribosome Flow Model on a Ring},
year={2015},
volume={12},
number={6},
pages={1429-1439},
}


@Article{hespan,
title={Switching between stabilizing controllers},
author={Hespanha, J. P. and Morse, A. S.},
journal={Automatica},
volume={38},
number={11},
year={2002},
pages={1905-1917},
}


@article{lie,
author = "D. Liberzon and J. P. Hespanha and A. S. Morse",
title = "Stability of switched  systems:
A {L}ie-algebraic condition",
journal = SCL,
skip-month = jun,
pages = "117-122",
year = "1999",
volume = "37"
}


@Article{blondel2003,
title={An elementary counterexample to the finiteness conjecture},
author={V. D. Blondel and J. Theys and A. A. Vladimirov},
year={2003},
journal={SIAM J. Matrix Anal. Appl.},
volume={24},
pages={963-970},
}


@Article{blondel,
title={A survey of computational complexity results in systems and control},
author={Blondel, V. D. and Tsitsiklis, J. N.},
year={2000},
journal={Automatica},
volume={36},
pages={1249-1274},
}

@misc{jaku,
author="B. Jakubczyk",
title="Introduction to Geometric Nonlinear Control;
Controllability and {L}ie Bracket",
year="2001",
url="http://www.impan.gov.pl/User/jakubczy/",
}


@incollection{sus97,
author="H. J. Sussmann",
title="Some recent results on the Maximum Principle of optimal
control theory",
booktitle=" Systems and Control in the Twenty-First
 Century",
editor="C. I. Byrnes and  B. N. Datta and  D. S. Gilliam and
C. F. Martin",
publisher="Birkh{\"a}user",
 year="1997", pages="351-372"
}

@incollection{lie-open-problem,
author="D. Liberzon",
title="Lie algebras and stability of switched nonlinear systems",
booktitle="Unsolved Problems in Mathematical Systems and Control Theory",
editor="V. Blondel and A. Megretski", publisher="Princeton University
Press", year=2004, pages="203-207"
}


@incollection{hespanha-open-problem,
author={J.  P. Hespanha},
title={${L}_2$-induced gains of switched linear systems},
booktitle="Unsolved Problems in Mathematical Systems and Control Theory",
editor="V. Blondel and A. Megretski", publisher="Princeton University
Press", year=2004, pages="131-133"
}


@book{fixedpoint,
title={Fixed Point Theorems with Applications to Economics and Game Theory},
author={K. C. Border},
publisher={Cambridge University Press},
year={1989},
}


@incollection{hes-morse-open-problem99,
author="J. P. Hespanha and A. S. Morse",
title="Input-output gains of switched linear systems",
booktitle="Open Problems in Mathematical Systems Theory
 and Control",
editor="V. D. Blondel and E. D. Sontag and
M. Vidyasagar and J. C. Willems",
publisher="Springer-Verlag",
year="1999", pages="121-125",
}


@incollection{kawski-open-problem,
author="M. Kawski",
title="Bases for {L}ie algebras and a continuous
{CBH} formula",
booktitle="Unsolved Problems in Mathematical Systems and Control Theory",
editor="V. Blondel and A. Megretski", publisher="Princeton University
Press", year=2004, pages="97-102",
}


@incollection{rasvan,
author="Rasvan, V.",
title="Delay independent and delay dependent {A}izerman problem",
booktitle="Unsolved Problems
 in Mathematical Systems and Control Theory",
editor="V. Blondel and A. Megretski",
publisher="Princeton University Press",
year=2004,
pages="212-220"
}

@Article{megretski-rantzer,
title={System analysis via integral quadratic constraints},
author={Megretski, A. and Rantzer, A.},
year={1997},
journal=TAC,
volume={42},
number={6},
pages={819-830}
}


%%%%%%%%%%%%%%%%%% new refs.
@book{isidori-book,
author="A. Isidori",
title="Nonlinear Control Systems",
edition="3rd", publisher="Springer",
address="Berlin", year=1995
}


@Book{vinter,
   author =   {Vinter, R.},
   title =   {Optimal Control},
  publisher="Birkh{\"a}user",
 year =   {2000},
}
@Article{xie,
title={Piecewise {L}yapunov
 functions for robust stability of linear time-varying systems},
author={Xie, L. and Shishkin, S. and Fu, M.},
year={1997},
journal={Systems and Control Letters},
volume={31},
pages={165-171},
number={3},
}


@Article{holcman,
title={Stability analysis of switched homogeneous  systems in the plane},
author={Holcman, D.   and Margaliot, M.},
journal=SIAM,
volume={41},
number={5},
year={2003},

pages={1609-1625},
}

@article{mar-lib,
author="M. Margaliot and D. Liberzon",
title="Lie--algebraic
stability conditions for nonlinear switched systems and
differential inclusions",
journal=SCL,  year=2006, pages={8-16},
volume={55}, number={1}, skipmonth={jan},
}

@article{margaliot_hall_sussmann,
author="M. Margaliot",
title="On the Analysis of Nonlinear Nilpotent Switched Systems using the
{H}all-{S}ussmann System",
journal=SCL,  volume={58}, pages={766-772},  year="2009",
}

@Article{mar-lang,
title={Necessary and Sufficient Conditions for Absolute Stability:
        The Case of Second-Order Systems},
author={Margaliot, M. and Langholz, G.},
journal={IEEE Trans. Circuits Syst.-I},
volume={50},
skip-number={2},
year={2003},
pages={227-234},
}

@incollection{sussmann1986,
author="Sussmann, H. J.",
title="A product expansion for the {C}hen series",
booktitle="Theory and Applications of Nonlinear Control Systems",
editor="Byrnes, C. I. and Lindquist, A.",
year="1986", pages="323-335",
publisher="Elsevier Science Publishers"
}


@Article{xinghuo,
title={Generating chaotic attractors with multiple merged
basins of attraction: A switching piecewise-linear control approach},
author={Xinghuo, J. L. and Chen, G.},
journal={IEEE Trans. Circuits Syst.-I},
volume={50},
skip-number={2},
year={2003},
pages={198-207},
}
@Article{zel,
title={Nonquadratic {L}yapunov functions
for robust stability analysis of linear uncertain systems},
author={Zelentsovsky, A. L.},
year={1994},
journal=TAC,
volume={39},
number={1},
pages={135-138}
}


@Article{pyarap,
title={Criteria of asymptotic stability of differential inclusions and
periodic motions of time-varying nonlinear control systems},
author={Pyatnitskiy, E. S. and Rapoport, L. B.},
year={1996},
journal={IEEE Trans.  Circuits  Syst.-I},
volume={43},
skip-number={3},
pages={219-229}
}

@Incollection{kawski,
   author = {Kawski, M.},
   title = {High-order maximal principles},
  pages = {313-326},
   booktitle = {New Trends in Nonlinear Dynamics and Control},
   series={Lecture Notes in Control and Information Sciences},
    volume={295},
   publisher = {Springer-Verlag},
   year = {2003},
   editor ={Kang, W. and   Xiao, M. and Borges, C.}
}

@book{gil,
   author = {M. I. Gil},
   title ={Explicit Stability Conditions for Continuous Systems:
   A Functional Analytic Approach},
   series={Lecture Notes in Control and Information Sciences},
    volume={314},
   publisher = {Springer-Verlag},
   year = {2005},
}


@book{Switched-and-impulsive-systems,
   author = {Zhengguo, L.  and  Yengchai, S. and Changyun, W.},
   title = {Switched and Impulsive Systems:
Analysis, Design and Applications},
   series={Lecture Notes in Control and Information Sciences},
    volume={313},
   publisher = {Springer-Verlag},
   year = {2005},
}


@book{borrelli,
   author = {F. Borrelli},
   title = {Constrained Optimal Control of Linear and
Hybrid Systems},
   series={Lecture Notes
in Control and Information Sciences},
    volume={290},
   publisher = {Springer-Verlag},
   year = {2003},
}


@book{johansson,
   author = {M. Johansson},
   title = {Piecewise Linear Control Systems},
   series={Lecture Notes
in Control and Information Sciences},
    volume={284},
   publisher = {Springer-Verlag},
   year = {2003},
}


@Incollection{rap,
   author = {Rapoport, L. B.},
   title = {Asymptotic stability and periodic motions of
            selector-linear differential inclusions},
  pages = {269-285},
   booktitle = {Robust Control via Variable Structure and
                Lyapunov Techniques},
 series={Lecture Notes in Control and Information Sciences},
    volume={217},
   publisher = {Springer},
   year = {1996},
   editor ={Garofalo, F. and Glielmo, L.},
}


@Article{mar-gitizadeh,
title={The problem of absolute stability:
 A~dynamic programming approach},
author={Margaliot, M. and Gitizadeh, R.},
journal={Automatica},
pages={1247-1252},
volume={40},
number={7},
year=2004,
}

@incollection{Sontag2010,
author="Sontag, Eduardo D.",
editor="Willems, Jan C.
and Hara, Shinji
and Ohta, Yoshito
and Fujioka, Hisaya",
title="Contractive Systems with Inputs",
bookTitle="Perspectives in Mathematical System Theory, Control, and Signal Processing: A Festschrift in Honor of Yutaka Yamamoto on the Occasion of his 60th Birthday",
year="2010",
publisher="Springer",
address="Berlin, Heidelberg",
pages="217-228",
}


@Incollection{Sontag2008,
author="Sontag, Eduardo D.",
editor="Nistri, Paolo and Stefani, Gianna",
title="Input to State Stability: Basic Concepts and Results",
bookTitle="Nonlinear and Optimal Control Theory: Lectures given at the C.I.M.E. Summer School held in Cetraro, Italy, June 19-29, 2004",
year="2008",
publisher="Springer Berlin Heidelberg",
address="Berlin, Heidelberg",
pages="163-220",
}


%%%%%%%%%%%%%%%%%%%%%%%%%%still more refs

@article{ugo-switched-siam,
author="U. Boscain",
title="Stability of planar switched systems:
The linear single input case",
journal=SIAM,
volume=41, skip-no=1, year=2002, pages="89-112"
}

@book{jurdjevic-book,
author="V. Jurdjevic",
title="Geometric Control Theory",
publisher="Cambridge University Press",
year=1996
}

@article{palais,
author="R. S. Palais",
title="A global formulation of the {L}ie theory of transformation groups",
journal="Mem. Amer. Math. Soc.",
volume=22, year=1957, pages="1-123"
}

@article{vakhrameev-1997,
author="S. A. Vakhrameev",
title="A bang-bang theorem with a finite number of switchings for nonlinear
smooth control systems",
journal="J.\ Math.\ Sci.",
volume=85, skip-no=3, year=1997, pages="2002-2016"
}

@article{agrachev-vakhrameev-doklady-1984,
author="A. A. Agrachev and S. A. Vakhrameev",
title="Nonlinear control systems of constant rank and bang-bang conditions
for extremal controls",
journal="Soviet Math.\ Dokl.",
volume=30,skip-no=3,year=1984,pages="620-624",
skip-note="(Translated from \textit{Dokl.\ Akad.\ Nauk SSSR},
279:265--269, 1984)"
}

@incollection{kupka90,
author="I. A. K. Kupka",
title="The ubiquity of {F}uller's phenomenon",
booktitle="Nonlinear Controllability and Optimal Control",
editor="H. J. Sussmann",
publisher="Marcel Dekker",
address="New York",
year=1990,
pages="313-350",
}


@incollection{grasse_and_sussmann1990,
author="K. A. Grasse and  H. J. Sussmann",
title="Global Controllability by Nice Controls",
booktitle="Nonlinear Controllability and Optimal Control",
editor="H. J. Sussmann",
publisher="Marcel Dekker",
address="New York",
year=1990,
pages="33-79",
}


@article{sigalotti05,
author="Sigalotti, M.",
title="Local regularity of optimal trajectories for control problems
with general boundary conditions",
journal="J. Dynam. Control Systems",
volume="11",
pages="91-123",
year=2005
}


@article{agrachev-sigalotti,
author="A. A. Agrachev and M. Sigalotti",
title="On the local structure of optimal trajectories
 in {R}$^3$",
journal=SIAM,
volume=42,
pages="513-531",
year=2003
}


 @article{magnus85,
author="J. R. Magnus",
title="On Differentiating Eigenvalues and Eigenvectors",
 journal="Econometric Theory",
 year="1985", volume="1", pages="179-191",
}


@incollection{sussmann-survey-83,
author="H. J. Sussmann",
title="Lie brackets, real analyticity and geometric control",
booktitle="Differential Geometric Control Theory",
editor="R. W. Brockett and R. S. Millman and H. J. Sussmann",
publisher="Birkh{\"a}user",
year=1983,
pages="1-116"
}


@incollection{schattler-1998,
author="Schattler, H.",
title="Time optimal feedback control for nonlinear systems: A geometric
approach",
booktitle="Geometry of Feedback and Optimal Control",
editor="Jakubczyk, B. and Respondek, W.",
publisher="Marcel Dekker",
address="New York",
year="1998",
pages="383-421",
}


@incollection{schattler90,
author="Schattler, H.",
title="Regularity properties of optimal trajectories:
 Recently developed techniques",
booktitle="Nonlinear Controllability and Optimal Control",
editor="H. J. Sussmann",
publisher="Marcel Dekker",
address="New York",
year=1990,
pages="263-277",
}


@Book{sch,
   author =   "van der Schaft, A. J.  and Schumacher, H.",
   title =   "An Introduction to Hybrid Dynamical Systems",
 series={Lecture Notes in Control and Information Sciences},
    volume={251},
  publisher = "Springer",
   year =   "2000",
   skipaddress ="" ,
}


@article{Ji-Fa01091994,
author = {J.  Ji-Fa},
title = {On the Global Stability of Cooperative Systems},
volume = {26},
skipnumber = {5},
pages = {455-458},
year = {1994},
journal = {Bulletin of the London Mathematical Society}
}


@article{seah-di-complete-1982,
author="S. W. Seah",
title="Existence of solutions and asymptotic equilibrium of multivalued
differential systems",
journal="J. Math. Anal. Appl.",
volume=89, year=1982, pages="648-663"
}

@article{taniguchi-di-complete-1992,
author="T. Taniguchi",
title="Global existence of solutions of
differential inclusions",
journal="J. Math. Anal. Appl.",
volume=166, year=1992, pages="41-51"
}

@article{howe-1983,
author="R. Howe",
title="Very basic {L}ie theory",
journal="American Mathematical Monthly",
volume=90, year=1983, pages="600-623"
}

@Incollection{arts,
   author = {Artstein, Z.},
   title = {Examples of stabilization with hybrid feedback},
  pages = {173-185},
   booktitle = {Hybrid Systems III: Verification and Control},
 series={Lecture Notes in Computer Science},
    volume={1066},

   publisher = {Springer},
   year = {1996},
   editor ={Alur, R. and Henzinger, T. A. and  Sontag, E. D. },
}


@article{sontag_gt_2010,
author={D. Angeli and P. {de Leenheer} and
   E. D. Sontag},
 title={Graph-theoretic characterizations
  of monotonicity of chemical networks in reaction coordinates},
  journal={J. Mathematical Biology},
  volume={61}, pages={581-616}, year={2010},
  }


@article{mono_posi_2005,
title={Monotone systems under positive feedback: multistability and a reduction theorem},
journal=SCL, author={G. Enciso and  E. D. Sontag},
volume={54},
pages={159-168},
year={2005},
}


@book{sontag_book,
author={E. D. Sontag},
 title={Mathematical Control Theory: Deterministic Finite Dimensional Systems},
edition={2}, publisher={Springer}, address={New York}, year={1998},
}


@Incollection{hes01,
   author = {J.  Hespanha and S. Bohacek and K.  Obraczka
and J.  Lee},
   title = {Hybrid Modeling of {TCP} Congestion Control},
  pages = {291-304},
   booktitle = {Hybrid Systems: Computation and Control},
 series={Lecture Notes in Computer Science},
    volume={2034},
   publisher = {Springer},
   year = {2001},
   editor ={M. D. {Di Benedetto} and A. Sangiovanni-Vincentelli},
}


@Article{bla,
title={Set invariance in control},
author={Blanchini, F.},
year={1999},
journal={Automatica},
volume={35},
pages={1747-1767},
number={11},
}

@Incollection{cuk,
   author = {Cuk, S.},
   title = {Basics of switched-mode power conversion:

Topologies, magnetics, and control},
  pages = {265-296},
   booktitle = {Modern Power Electronics:
         Evolution, Technology, and Applications},
   publisher = {IEEE Press},
   year = {1992},
   editor ={Bose, B. K.},
}

@book{liberzon_opt_cont,
title={Calculus of Variations and Optimal Control Theory: A Concise Introduction},
author={Daniel Liberzon},
year={2012},publisher={Princeton University Press}, address={Princeton, NJ},
}

@article{libsur99,
title="Basic problems in stability and design of switched systems",
journal=CSM,
author="D. Liberzon and A. S. Morse",
volume="19",year="1999",pages="59-70",
}

@Article{mar-simple,
title={Stability Analysis of
Switched Systems
using  Variational  Principles: An Introduction},
 year={2006},
author={Margaliot, M.},
 journal={Automatica},
volume={42}, pages={2059-2077}, skipnumber={12},
}

@ARTICLE{Dalal2000,
title = {Strict concavity with homogeneity and decreasing returns to scale},
author = {Dalal, Ardeshir},
year = {2000},
journal = {Atlantic Economic Journal},
volume = {28},
number = {3},
pages = {381-382},
}

@article{ss_taep_ps,
	Author = {Lakatos,G and Chou,T and Kolomeisky,A},
	Date = {2005/01/},
	Date-Added = {2013-12-15 11:35:56 +0000},
	Date-Modified = {2013-12-15 11:35:56 +0000},
	Journal = {Physical review. E, Statistical, nonlinear, and soft matter physics},	
	Month = {01},
	Number = {1 Pt 1},
	Pages = {011103},
	Title = {Steady-state properties of a totally asymmetric exclusion process with periodic structure.},
	Url = {http://europepmc.org/abstract/MED/15697576},
	Volume = {71},
	Year = {2005},
}

@PhDThesis{hespanha_phd,
  AUTHOR     = {J. P. Hespanha},
  YEAR       = {1998},
  TITLE      = {Logic-based switching algorithms in control},
  SCHOOL     = {Dept. Elec. Eng, Yale University},
  address    = {New Haven, CT},
  ID         = {PhD}
}

% RESPONSE, MSB 04/04/2009: added
@book{sun_ge_book,
title="Switched Linear Systems: Control and Design",
author="Z. Sun and S. S. Ge",
publisher="Springer",
year="2005",
}


@book{sun_ge_book_new,
title="Stability Theory of Switched Dynamical Systems",
author="Z. Sun and S. S. Ge",
publisher="Springer",
year="2011",
}

% for subadditive
@book{fund_convx_ana,

title="Fundamentals of Convex Analysis",
author="J. Baptiste and H. Urruty and C. Lemarechal",
publisher="Springer",
year="2001",
}

% convex optimization
@book{convex_boyd,
title="Convex Optimization",
author="S. Boyd and L. Vandenberghe",
publisher="Cambridge University Press",
year="2004",
}

% semidefinite matrix
@book{pract_opt,
title="Practical Optimization: Algorithms and Engineering Applications",
author="A. Antoniou and W. Lu",
publisher="Springer",
year="2010",
}


@book{gohberg2013classes,
  title={Classes of Linear Operators},
  author={Gohberg, Israel and Goldberg, Seymour and Kaashoek, Marius A.},
  volume={1},
  year={2013},
  publisher={Birkh{\"a}user}
}

@book{matrx_ana,
title="Matrix Analysis",
author="R. A. Horn and C. R. Johnson",
publisher="Cambridge University Press",
year="2013", edition="2",
}

@article{HRFM_concave,
author={Y. Zarai and M. Margaliot and T. Tuller},
title={Maximizing    Protein Translation Rate in the Ribosome Flow Model: the Homogeneous Case},
  year={2014}, journal=TCBB, pages={1184-1195},volume={11},
}


@article{edri2014rna,
  title={The {RNA }polymerase flow model of gene transcription},
  author={Edri, Shlomit and Gazit, Eran and Cohen, Emmanuel and Tuller, Tamir},
  journal={Biomedical Circuits and Systems, IEEE Transactions on},
  volume={8},
  number={1},
  pages={54--64},
  year={2014},
  publisher={IEEE}
}

@article{RFMEO,
author={Yoram Zarai and  Michael Margaliot and Tamir Tuller},
title={The Ribosome Flow Model with Extended Objects},
year={2017},journal={J. R. Soc. Interface},pages={1420170128},
url={http://doi.org/10.1098/rsif.2017.0128},
}

@inproceedings{sontag_cotraction_tutorial,
title={Contraction Methods for Nonlinear Systems: A Brief Introduction and Some Open Problems},
author={Z. Aminzare and E. D. Sontag},
booktitle=CDC53,year={2014},
address={Los Angeles, CA},pages={3835-3847},
}

@inproceedings{sontag2014three,
  title={On three generalizations of contraction},
  author={E. D. Sontag and M. Margaliot and T. Tuller},
  booktitle=CDC53,
  pages={1539-1544},
  year={2014},
  organization={IEEE}
}


@article{mexdenrfm,
    author = {Zarai, Yoram AND Margaliot, Michael AND Tuller, Tamir},
    journal = {PLOS ONE},
    title = {On the Ribosomal Density that Maximizes Protein Translation Rate},
    year = {2016},
    volume = {11},
    url = {https://doi.org/10.1371/journal.pone.0166481},
    pages = {1-26},
    number = {11},
}

@article{rfm_concave,
	author = "Poker, G. and Zarai, Y. and Margaliot, M. and Tuller, T.",
	journal = "J. Royal Society Interface",
	volume="11",number="100",
	title = "Maximizing protein translation rate in the
	nonhomogeneous ribosome flow model: a convex optimization approach",
	year = "2014",
}


@article{RFM_sense,
	author = "Poker, Gilad and Margaliot, Michael and Tuller, Tamir",
	title = "Sensitivity of {mRNA} translation",
	volume="5",year={2015}, journal={Scientific Reports},pages = "12795",
}


@article{PhysRevE.70.046101,
  title = {Totally asymmetric simple exclusion process with {Langmuir} kinetics},
  author = {Parmeggiani, A. and Franosch, T. and Frey, E.},
  journal = {Phys. Rev.. E},
  volume = {70},
  issue = {4},
  pages = {046101},
  numpages = {20},
  year = {2004},
}

@article{1751-8121-45-18-185002,
  author={Takahiro Ezaki and Katsuhiro Nishinari},
  title={Exact stationary distribution of an asymmetric simple exclusion process with {Langmuir} kinetics and memory reservoirs},
  journal={Journal of Physics A: Mathematical and Theoretical},
  volume={45},
  number={18},
  pages={185002},
   year={2012},
  abstract={We present exact stationary distribution of the asymmetric simple exclusion process with {Langmuir} kinetics and its related model on a ring. The distribution is expressed in simple factorized forms, that is supported by detailed balance conditions satisfied in the attachment and detachment transitions. Using the equations, we derive the density of particles, the current and the variance of particle numbers in each lane. The expressions for the density are well associated with the ones found in equilibrium theories. Moreover, a simple relation between the variance of particle number and the current is proved}
}


@article{willmann2002exact,
  title={Exact {Hurst} exponent and crossover behavior in a limit order market model},
  author={Willmann, RD and Sch{\"u}tz, GM and Challet, Damien},
  journal={Physica A: Statistical Mechanics and its Applications},
  volume={316},
  number={1},
  pages={430--440},
  year={2002},
  publisher={Elsevier}
}


@article{PhysRevLett.90.086601,
  title = {Phase Coexistence in Driven One-Dimensional Transport},
  author = {Parmeggiani, A. and Franosch, T. and Frey, E.},
  journal = {Phys. Rev. Lett.},
  volume = {90},
  issue = {8},
  pages = {086601},
  numpages = {4},
  year = {2003},
  }
	
	
@article{GARLOFF2003103,
title = "Intervals of almost totally positive matrices",
journal = LAA,
volume = "363",
pages = "103-108",
year = "2003",
author = "J. Garloff",
}


 
 @Article{Whitney1952,
author="Whitney, A. M.",
title="A reduction theorem for totally positive matrices",
journal="J. d'Analyse Math{\'e}matique",
year="1952",
volume="2",
number="1",
pages="88-92",
}


@ARTICLE{rola,
author={R. Alseidi and    M. Margaliot  and  J. Garloff},
title={On the spectral properties of  nonsingular
matrices that are strictly sign-regular for some order with applications 
to    totally positive discrete-time systems},
year={2019},
journal={J. Math. Anal. Appl.},
volume={474}, pages={524-543}, 
}


@ARTICLE{sandberg78,
author={Sandberg, Irwin W.},
journal={IEEE Trans. Circuits and Systems},
title={On the mathematical foundations of compartmental analysis in biology, medicine, and ecology},
year={1978},
volume={25},
number={5},
pages={273-279},
}

@article{Arcak20111219,
title = "Certifying spatially uniform behavior in reaction-diffusion  {PDE} and compartmental {ODE} systems",
journal = "Automatica",
volume = "47",
number = "6",
pages = "1219-1229",
year = "2011",
 author = "Murat Arcak",
}


 @article{Lipowsky200553,
title = "Life is motion: multiscale motility of molecular motors ",
journal = "Physica A: Statistical Mechanics and its Applications ",
volume = "352",
number = "1",
pages = "53-112",
year = "2005",
issn = "0378-4371",
doi = "http://dx.doi.org/10.1016/j.physa.2004.12.034",
author = "Reinhard Lipowsky and Stefan Klumpp"
}


@article{lipowsky2001random,
  title={Random walks of cytoskeletal motors in open and closed compartments},
  author={Lipowsky, Reinhard and Klumpp, Stefan and Nieuwenhuizen, Theo M},
  journal={Phys. Rev. Lett.},
  volume={87},
  number={10},
  pages={108101},
  year={2001},
  publisher={APS}
}


@article{Chowdhury2005318,
title = "Physics of transport and traffic phenomena in biology: from molecular motors and cells to organisms ",
journal = "Physics of Life Reviews ",
volume = "2",
number = "4",
pages = "318 - 352",
year = "2005",
note = "",
issn = "1571-0645",
doi = "http://dx.doi.org/10.1016/j.plrev.2005.09.001",
author = "Debashish Chowdhury and Andreas Schadschneider and Katsuhiro Nishinari"
}


@article{backtracking_rnap,
	Annote = {10.1038/nature02191},
	Author = {Shaevitz, Joshua W. and Abbondanzieri, Elio A. and Landick, Robert and Block, Steven M.},
	Date = {2003/12/11/print},
	Date-Added = {2015-11-06 10:41:07 +0000},
	Date-Modified = {2015-11-06 10:41:07 +0000},
	Day = {11},
	Isbn = {0028-0836},
	Journal = {Nature},
	L3 = {http://www.nature.com/nature/journal/v426/n6967/suppinfo/nature02191_S1.html},
	M3 = {10.1038/nature02191},
	Month = {12},
	Number = {6967},
	Pages = {684--687},
	Title = {Backtracking by single {RNA} polymerase molecules observed at near-base-pair resolution},
	Ty = {JOUR},
	Volume = {426},
	Year = {2003},
	Bdsk-Url-1 = {http://dx.doi.org/10.1038/nature02191}
}


@article{rnap_backtrack,
	Author = {Nudler, Evgeny},
	Journal = {Cell},
year={2015},
	Number = {7},
	Pages = {1438--1445},
	Title = {{RNA} Polymerase Backtracking in Gene Regulation and Genome Instability},
	Volume = {149},
}


@article{structual_rnap,
	Annote = {10.1038/nature09785},
	Author = {Cheung, Alan C. M. and Cramer, Patrick},
	Date = {2011/03/10/print},
	Date-Added = {2015-11-06 10:49:57 +0000},
	Date-Modified = {2015-11-06 10:49:57 +0000},
	Day = {10},
	Isbn = {0028-0836},
	Journal = {Nature},
	L3 = {http://www.nature.com/nature/journal/v471/n7337/abs/10.1038-nature09785-unlocked.html#supplementary-information},
	M3 = {10.1038/nature09785},
	Month = {03},
	Number = {7337},
	Pages = {249--253},
	Publisher = {Nature Publishing Group, a division of Macmillan Publishers Limited. All Rights Reserved.},
	Title = {Structural basis of {RNA} polymerase {II} backtracking, arrest and reactivation},
	Ty = {JOUR},
	Volume = {471},
	Year = {2011},
	Bdsk-Url-1 = {http://dx.doi.org/10.1038/nature09785}
}


@article{nascent_tran_vis,
	Annote = {10.1038/nature09652},
	Author = {Churchman, L. Stirling and Weissman, Jonathan S.},
	Date = {2011/01/20/print},
	Date-Added = {2015-11-06 10:53:25 +0000},
	Date-Modified = {2015-11-06 10:53:25 +0000},
	Day = {20},
	Isbn = {0028-0836},
	Journal = {Nature},
	L3 = {http://www.nature.com/nature/journal/v469/n7330/abs/10.1038-nature09652-unlocked.html#supplementary-information},
	M3 = {10.1038/nature09652},
	Month = {01},
	Number = {7330},
	Pages = {368--373},
	Publisher = {Nature Publishing Group, a division of Macmillan Publishers Limited. All Rights Reserved.},
	Title = {Nascent transcript sequencing visualizes transcription at nucleotide resolution},
	Ty = {JOUR},
	Volume = {469},
	Year = {2011},
	Bdsk-Url-1 = {http://dx.doi.org/10.1038/nature09652}
}

@article{zhang2010global,
  title={Global and local depletion of ternary complex limits translational elongation},
  author={Zhang, Gong and Fedyunin, Ivan and Miekley, Oskar and Valleriani, Angelo and Moura, Alessandro and Ignatova, Zoya},
  journal={Nucleic Acids Res.},
  volume={38},
  number={14},
  pages={4778--4787},
  year={2010},
  publisher={Oxford Univ Press}
}

@article{kurland1992translational,
  title={Translational Accuracy and the Fitness of Bacteria},
  author={Kurland, CG},
  journal =      {Annu. Rev. Genet.},
  volume={26},
  number={1},
  pages={29--50},
  year={1992},
  publisher={Annual Reviews 4139 El Camino Way, PO Box 10139, Palo Alto, CA 94303-0139, USA}
}

@article{schoenberg1953smoothing,
  title={On smoothing operations and their generating functions},
  author={Schoenberg, Isaac Jacob and others},
  journal={Bulletin of the American Mathematical Society},
  volume={59},
  number={3},
  pages={199--230},
  year={1953},
  publisher={American Mathematical Society}
}


@article{mairhuber1959variation,
  title={On variation diminishing transformations on the circle},
  author={Mairhuber, J. C. and Schoenberg, I. J. and Williamson, R. E.},
  journal={Rendiconti del Circolo matematico di Palermo},
  volume={8},
  number={3},
  pages={241--270},
  year={1959},
  publisher={Springer}
}

@INPROCEEDINGS{rfm_chap,
  author =      {Y. Zarai and M. Margaliot and T. Tuller},
  title =        {Modeling and Analyzing the Flow of Molecular Machines  in Gene Expression},
  booktitle =    {Systems Biology},
  editor=      {N. Rajewsky and  S. Jurga and J. Barciszewski},
  publisher =    {Springer},year={2018}, pages={275-300},
}


@article {schoenberg51,
    AUTHOR = {Schoenberg, I. J. and Whitney, Anne},
     TITLE = {A theorem on polygons in {$n$} dimensions with applications to
              variation-diminishing and cyclic variation-diminishing linear
              transformations},
   JOURNAL = {Compositio Math.},
     VOLUME = {9},
      YEAR = {1951},
     PAGES = {141--160},
 }

@article{rfmnets,
title={Networks of ribosome flow models for modeling and analyzing intracellular traffic},
author={Itzik Nanikashvili and Yoram Zarai and Alexander Ovseevich and  Tamir Tuller and  Michael Margaliot},
journal={Scientific Reports},
volume={9}, pages={1703}, 
year={2019}}


@article{exp_osci,
title={On the Exponent of Several Classes of Oscillatory Matrices},
author={Y. Zarai and  M. Margaliot},
journal=LAA,
volume={608}, pages={363-386},
 year={2021},
}


@article{rfmdr,
title={Optimal down regulation of {mRNA} translation},
author={Y. Zarai and  M. Margaliot and T. Tuller},
journal={Scientific Reports},
volume={7}, pages={41243}, year={2017},
}


@article{RFM_model_compete_J,
title = "A model for competition for ribosomes in the cell",
author="Alon Raveh and  Michael Margaliot and  Eduardo D. Sontag and Tamir Tuller",
year = "2016",
journal="J. Royal Society Interface",
volume="13",
	number = "116",
}


@article{fallat2007class,
  title={A class of oscillatory matrices with exponent~$n-1$},
  author={Fallat, Shaun and Liu, Xiao Ping},
  journal=LAA,
  volume={424},
  number={2-3},
  pages={466--479},
  year={2007},
  publisher={Elsevier}
}

@article{fallat2004remark,
  title={A remark on oscillatory matrices},
  author={Fallat, Shaun M},
  journal=LAA,
  volume={393},
  pages={139--147},
  year={2004},
  publisher={Elsevier}
}

@article{gasca1992total,
  title={Total positivity and {Neville} elimination},
  author={Gasca, Mariano and Pena, Juan Manuel},
  journal=LAA,
  volume={165},
  pages={25--44},
  year={1992},
  publisher={North-Holland}
}

@article{whitney1952reduction,
  title={A reduction theorem for totally positive matrices},
  author={Whitney, Anne M},
  journal={Journal d'Analyse Math{\'e}matique},
  volume={2},
  number={1},
  pages={88--92},
  year={1952},
  publisher={Springer}
}


@article{price1968monotone,
  title={Monotone and oscillation matrices applied to finite difference approximations},
  author={Price, Harvey S},
  journal={Mathematics of Computation},
  volume={22},
  number={103},
  pages={489--516},
  year={1968},
  publisher={JSTOR}
}

@MastersThesis{kardell2010total,
    author     =     {Marcus Kardell},
    title     =     {Total positivity and oscillatory kernels:
                                An overview, and applications to the spectral theory of the cubic string},
    school     =     {Link\"{o}ping University},
    address     =     {Sweden},
    year     =     {2010},
    }


@article{banaji_angeli,
author = {Banaji, Murad and Angeli, David},
title = {Convergence in Strongly Monotone Systems with an Increasing First Integral},
journal = {SIAM J. Mathematical Analysis},
volume = {42},
number = {1},
pages = {334-353},
year = {2010},
}

@article{fomin2000total,
  title={Total positivity: tests and parametrizations},
  author={Fomin, Sergey and Zelevinsky, Andrei},
  journal={The Mathematical Intelligencer},
  volume={22},
  number={1},
  pages={23--33},
  year={2000},
  publisher={Springer}
}


@article{finkel2021,
author={Yaara Finkel and  Avi Gluck and  Aharon Nachshon and  Roni Winkler and  Tal Fisher and  Batsheva Rozman and  Orel Mizrahi and  Yoav Lubelsky and  Binyamin Zuckerman and  Boris Slobodin and  Yfat Yahalom-Ronen and  Hadas Tamir and  Igor Ulitsky and  Tomer Israely and  Nir Paran and  Michal Schwartz and  Noam Stern-Ginossar},
title={ {SARS-CoV-2} uses a multipronged strategy to impede host protein synthesis},
journal={Nature},volume={594}, pages={240–245}, year={2021},
}


@article{Aditi_competition_2021,
	author = {Jain, Aditi and Margaliot, Michael and Gupta, Arvind Kumar},
	title = {Large-scale {mRNA} translation and the intricate effects of competition for the finite pool of ribosomes},
	year = {2022},volume={19},pages={20220033}, 
	journal={J. Royal Society Interface},
 }

 


@article{Mier991,
author = {Mierczynski, Janusz},
journal = {Colloquium Mathematicae},
number = {1},
pages = {43-47},
title = {A class of strongly cooperative systems without compactness},
volume = {62},
year = {1991},
}

@article{bartroff2010optimal,
  title={On optimal allocation of a continuous resource using an iterative approach and total positivity},
  author={Bartroff, Jay and Goldstein, Larry and Rinott, Yosef and Samuel-Cahn, Ester},
  journal={Advances in Applied Probability},
  volume={42},
  number={3},
  pages={795--815},
  year={2010},
  publisher={Cambridge University Press}
}

@article{curtis1998circular,
  title={Circular planar graphs and resistor networks},
  author={Curtis, Edward B and Ingerman, David and Morrow, James A},
  journal=LAA,
  volume={283},
  number={1-3},
  pages={115--150},
  year={1998},
  publisher={Elsevier}
}

@article{Hausdorff_contract,
title="Generalization of the multiplicative and additive compounds of square matrices and contraction in the {Hausdorff} dimension",
author="C. Wu and R. Pines and
M. Margaliot and J.J. Slotine",
year="2021",note=s,
url="https://arxiv.org/abs/2008.10321",
}

@inproceedings{comp_barshalom,
title={Compound matrices in systems and control theory},
author={E. Bar-Shalom and M. Margaliot},
 booktitle = CDC60,
 note=ta,
year={2021},
url={https://arxiv.org/abs/2103.15097}
}
@article{higueras2002logarithmic,
  title={Logarithmic norms and nonlinear {DAE} stability},
  author={Higueras, Inmaculada and S{\"o}derlind, Gustaf},
  journal={BIT Numerical Mathematics},
  volume={42},
  number={4},
  pages={823--841},
  year={2002},
  publisher={Springer}
}
@article{mono_norms,
author={F. L. Bauer and J. Stoer and C. Witzgall},
title={Absolute and monotone norms},
journal={Numer. Math.}, volume={3}, pages={257-264},year={1961},
}

@article{contract_DAES,
author = {Higueras, Inmaculada and Garcia-Celayeta, Berta},
title = {Logarithmic Norms for Matrix Pencils},
journal = {SIAM J. on Matrix Analysis and Applications},
volume = {20},
number = {3},
pages = {646-666},
year = {1999},
}

@ARTICLE{SLOTINE_RANDOM,
  author={Pham, Quang-Cuong and Tabareau, Nicolas and Slotine, Jean-Jacques},
  journal=TAC, 
  title={A Contraction Theory Approach to Stochastic Incremental Stability}, 
  year={2009},
  volume={54},
  number={4},
  pages={816-820},
}
@article{kordercont,
	title={$k$-contraction: theory and applications},
	author={Chengshuai Wu and  Ilya Kanevskiy  and Michael Margaliot},
	year={2020}, 
	url={https://arxiv.org/abs/2008.10321}
}

@article{kordercont,
	title={$k$-contraction: theory and applications},
	author={Chengshuai Wu and  Ilya Kanevskiy  and Michael Margaliot},
	year={2020}, 
	url={https://arxiv.org/abs/2008.10321}
}


@ARTICLE{Insberghe2021,
  author={Michael Van Insberghe and  Jeroen van den Berg and Amanda Andersson-Rolf and Hans Clevers and Alexander van Oudenaarden },
  journal={Nature}, 
  title={Single-cell Ribo-seq reveals cell cycle-dependent translational pausing}, 
  year={2021},
  volume={597},
  pages={561–565},
}


 @misc{Secher2009,
 title={INSULIN PRECURSORS AND A PROCESS FOR THEIR PREPARATION},
 year={16.09.2009}, url={https://patents.google.com/patent/EP1377608A1/en}, author={Thomas Boerglum Kjeldsen},
 }
 %author={Secher Anne et al},
 
 
 
 @ARTICLE{Dana2014B,
  author =       {A. Dana and T. Tuller},
  title =        {Mean of the typical decoding rates: a new translation efficiency index based on the analysis of ribosome profiling data
},
  year =         {2014},
journal={G3 (Bethesda)},
volume={5},
  number={1},
  pages={73-80},
}


 @ARTICLE{Bonven1979,
  author =       {B Bonven and K Gulløv},
  title =        {Peptide chain elongation rate and ribosomal activity in Saccharomyces cerevisiae as a function of the growth rate 
},
  year =         {1979},
journal={Mol Gen Genet},
volume={170},
  number={2},
  pages={225-30},
}

@ARTICLE{Haar2008,
  author =       {Tobias von der Haar },
  title =        {A quantitative estimation of the global translational activity in logarithmically growing yeast cells},
  year =         {2008},
journal={BMC Syst Biol},
volume={2},
  pages={87},
}


@ARTICLE{Lackner,
  author =       {Daniel H Lackner and Traude H Beilharz and Samuel Marguerat and Juan Mata and Stephen Watt and Falk Schubert and Thomas Preiss and Jürg Bähler},
  title =        {A network of multiple regulatory layers shapes gene expression in fission yeast},
  year =         {2007},
journal={Mol Cell},
volume={26},
  number={1},
  pages={145-55},
}


@ARTICLE{Sabi2019,
  author =       {R. Sabi  and T. Tuller},
  title =        {Modelling and measuring intracellular competition for finite resources during gene expression.},
  year =         {2019},
journal={J R Soc Interface},
volume={16},
  number={154},
}


  


@ARTICLE{Ceroni2015,
  author =       { Francesca Ceroni and
  Rhys Algar and  Guy-Bart Stan and
  Tom Ellis},
  title =        {Quantifying cellular capacity identifies gene expression designs with reduced burden},
  year =         {2015},
journal={Nat. Methods},
volume={12},
  pages={415–418},
}


@ARTICLE{Zur2016,
  author =       {H. Zur  and T. Tuller},
  title =        {Predictive biophysical modeling and understanding of the dynamics of mRNA translation and its evolution},
  year =         {2016},
journal={Nucleic Acids Res.},
volume={44},
  pages={9031-9049},
}

@ARTICLE{Swartz2006,
  author =       {J. Swartz},
  title =        {Developing cell-free biology for industrial applications},
  year =         {2006},
journal={J. Industrial Microbiology and Biotechnology},
volume={33},
  pages={476–485},
}
