## Supplementary material for "Translation in the cell under fierce competition for shared resources: a mathematical model": Latex source files: VIRAL_PEN.tex

%\documentclass[letterpaper, 12 pt, onecolumn ]{ieeeconf}  % Comment this line out
                                                          % if you need a4paper
%\documentclass[a4paper, 10pt, conference]{ieeeconf}      % Use this line for a4
                                                          % paper

%\IEEEoverridecommandlockouts                              % This command is only
                                                          % needed if you want to


%\documentclass[a4paper, final]{ieeeconf}
  % Use this line for a4 paper
%\IEEEoverridecommandlockouts
  % Needed if you want to use the \thanks command
%\overrideIEEEmargins
  % Needed to meet printer requirements.

% If IEEEtran.cls has not been installed into the LaTeX system files,
% manually specify the path to it like:
\documentclass[12pt,journal,onecolumn]{IEEEtran}

%\usepackage{subfig}
\usepackage{graphicx}

%\usepackage[utf8]{inputenc}
%\usepackage[T1]{fontenc}
\usepackage{url}
%%%\usepackage{kbordermatrix}
%\usepackage{natbib}
\usepackage{amsmath,amssymb,mathtools}
 \usepackage{amsthm,thmtools}
%\usepackage{ thmtools}
\usepackage{amsfonts} 

%%%\usepackage{theorem}
%\usepackage[dvips]{epsfig}      % ******* for figures ***************
%\usepackage{graphicx}
%%\usepackage{auto-pst-pdf}
\usepackage{enumerate}
\usepackage{color}                  %******** for colored remarks********

\usepackage{verbatim}
%\usepackage{biblatex}

%%\usepackage{refcheck} 

%\usepackage{multicol}

\usepackage{amssymb}
\usepackage{amsbsy}
\usepackage{amsmath}
\usepackage{setspace}
\usepackage{url}

\usepackage{subcaption}
\usepackage{float}
 \usepackage{dsfont}
 
\usepackage{tikz}
\usetikzlibrary{positioning}

%\usepackage{pdfpages}

%\usepackage{enumitem}  % for using different labels in enumerate

%\theoremstyle{plain}
%\newtheorem{Theorem}{Theorem}
%%\newtheorem{Lemma}{Lemma}
%\newtheorem{Proposition}{Proposition}
%\newtheorem{Corollary}{Corollary}
%\newtheorem{Conjecture}{Conjecture}
%\newtheorem{Problem}{Problem}
%\newtheorem{Fact}{Fact}


\newcommand{\imagi}{{\mathrm{i}}}

\newcommand{\jjs}[1]{\textcolor{blue}{jjs -- #1}}

 \newcommand{\Ave}{\operatorname{Ave}}
 \newcommand{\dist}{\operatorname{dist}}
 \renewcommand{\div}{\operatorname{div}}
 \newcommand{\Int}{\operatorname{int}}
 \newcommand{\Clos}{\operatorname{clos}}
 \newcommand{\real}{\operatorname{Re}}
 \newcommand{\imag}{\operatorname{Im}}
 \newcommand{\diag}{\operatorname{diag}}
 \newcommand{\sign}{\operatorname{sgn}}
 \newcommand{\tr}{\operatorname{tr}}
 \newcommand{\adju}{\operatorname{adj}}
 
\newcommand{\st}{\, | \,}

\newcommand\mycom[2]{\genfrac{}{}{0pt}{}{#1}{#2}}

\newcommand{\spanop}{\operatorname{span}}
\newcommand{\vecop}{\operatorname{vec}}

\newcommand{\updt}[1]{{\color{blue}#1}}


%%\newcommand{\trace}{\operatorname{trace}}

\newcommand{\inter}{\operatorname{int}}

\newcommand*\diff{\mathop{}\!\mathrm{d}}


\newcommand{\trace}{\operatorname{trace}}

 \declaretheorem[name={Proof},    numbered=no,qed=$\blacksquare$ ]{Proof}


\declaretheorem[name={Definition}  ] {Definition}
\declaretheorem[name={Example}  ] {Example}
\declaretheorem[name={Theorem}  ] {Theorem}
\declaretheorem[name={Lemma}  ] {Lemma}
\declaretheorem[name={Remark}  ] {Remark}
\declaretheorem[name={Corollary}  ] {Corollary}
\declaretheorem[name={Assumption}  ] {Assumption}
\declaretheorem[name={Proposition}  ] {Proposition}
\declaretheorem[name={Fact}  ] {Fact}
%\newcommand*\diff{\mathop{}\!\mathrm{d}}

 %\newcommand{\real}{\operatorname{Re}}

\newcommand{\beq}       {\begin{equation}}
\newcommand{\eeq}       {\end{equation}}
\newcommand{\bery}      {\begin{array}}
\newcommand{\eery}      {\end{array}}
\newcommand{\berys}     {\begin{array*}}
\newcommand{\eerys}     {\end{array*}}
\newcommand{\beqry}     {\begin{eqnarray}}
\newcommand{\eeqry}     {\end{eqnarray}}
\newcommand{\beqrys}    {\begin{eqnarray*}}
\newcommand{\eeqrys}    {\end{eqnarray*}}
\newcommand{\dis}   {\displaystyle}
\newcommand{\noi}{\noindent}
\def\bbr{{\mathbb R}}
\def \R {{\mathbb R}}
\newcommand{\E}{{\mathbf E}}
\newcommand{\M}{{\mathbf M}}
\newcommand{\Prob}{{\rm Prob}}
\newcommand{\Z}{{\mathbf Z}}
 
 
 %\renewcommand{\theequation}{\thesection.\arabic{equation}}
\def\myref#1{(\ref{#1})}
\def\mysection#1{\section{#1}\setcounter{equation}{0}}
\def\sh#1{{\rm Sh}(#1)}
\def\Sh{\mathop{\rm Sh}\nolimits}
\def\mod{\mathop{\rm mod}\nolimits}
\def\sign{\mathop{\rm sign}\nolimits}
\def\Var{\mathop{\rm Var}\nolimits}
\def\Spec{\mathop{\rm Spec}\nolimits}
\def\Sh{\mathop{\rm Sh}\nolimits}
\def\ext{\mathop{\rm ext}\nolimits}
\def\Re{\mathop{\rm Re}\nolimits}
\def\diag{\mathop{\rm diag}\nolimits}
\def\GL{\mathop{\rm GL}\nolimits}
 \def\C{{\mathbf C}}
\def\Z{{\mathbf Z}}
\def\T{{\mathcal T}}
\def\A{{\mathcal A}}
\def\t{{\mathbf t}}
\def\suppt{\mathop{\rm supp}\nolimits}
\def\interior{\mathop{\rm int}\nolimits}

\newcommand{\be}{\begin{equation}}
\newcommand{\ee}{\end{equation}}

%\usepackage{xcolor}
%\def\LG{\textcolor{blue}}
\doublespace 
 
\begin{document}
%%%%%%%%%%%%%%%%%%%%%%%%%%%%%%%%%%%%%%%%%%%%%%%%%%%%%%%
\title{\LARGE     Translation in the cell under fierce  competition for shared resources:  a mathematical~model   }

\author{Rami Katz, Elad Attias, Tamir Tuller, and Michael Margaliot\thanks{RK, EA, and~MM  are with the School of Electrical Engineering, Tel Aviv University, Israel.
TT is with the department of Biomedical Engineering, Tel Aviv University, Israel.  The research of~MM is partially supported by a research grants from  the ISF and the DFG. Corresponding author: \texttt{} }}

\maketitle
 
 \begin{abstract}
%%%%
During the process of    translation   the mRNAs   in the cell ``compete'' for shared resources like tRNA molecules and ribosomes. This creates an indirect and intricate coupling between the  mRNAs.
For example, if ribosomal ``traffic jams'' 
evolve on some mRNA then the abundance of free ribosomes may decrease
leading to lower  initiation  rates in the other mRNAs.   
When the shared resources are abundant the coupling  between mRNAs due to this  competition is weak. However, when the resources are scarce, e.g., when the pool of free ribosomes is starved, the competition may have a dramatic   effect on the dynamics of translation in the cell.  This scenario may be relevant  for example under stress conditions or during a high yield viral infection, where  the viral mRNAs ``hijack'' components of the translation machinery. Fierce competition for shared resources   may also take place in synthetic
or engineered systems such as cell free systems or in the case of high-throughput
heteroglougs gene expression. 

We study this scenario 
using a mathematical model that includes a network of~$m$ ribosome flow models~(RFMs)  interconnected via a pool of free ribosomes. Each RFM is a non-linear dynamical model for ribosome flow along a single mRNA molecule, and the interconnection via the pool encapsulates the  competition for shared resources. 
We   analyze the case where~$m$ is large, i.e., a there is a large number of mRNAs. This  implies that many ribosomes are attached to the   mRNAs and  thus the pool is starved. 

Our model allows quantitative and qualitative analysis of the   network steady state  when the pool is starved. 
%%%%%%%%%%%%%%%%%%%%%%%%%%%%%%
Our analysis results show  
that adding an mRNA
to the network 
always decreases the steady state pool density. This makes sense, as every new mRNA
``consumes'' ribosomes.
  We also show that adding an mRNA   
 has an intricate effect on the total protein production in the network: 
 on the one-hand, the new mRNA  produces new proteins. On the other-hand, the other mRNAs produce less proteins, as the pool that feeds these mRNAs now has a smaller abundance of ribosomes.
 Our analysis  yields  an explicit bound for the total production rate of the network when  the number  of RFMIOs is very large. In particular, we analyze 
 how     the total density of ribosomes in the network  bounds the total production rate. This bound demonstrates that when the  number of mRNAs  increases, the marginal utility  of adding another mRNA  diminishes, and the total protein production  rate  saturates to a limiting value. 
We demonstrate   our
analysis  approach using an example of producing insulin in a  
 cell free system. 
%%%%
 \end{abstract}
 
\begin{IEEEkeywords}
mRNA translation,
competition for shared resources,
cell free systems,
Perron-Frobenius theory,
spectral  analysis of
 tri-diagonal matrices, perturbations of eigenvalues and eigenvectors.
\end{IEEEkeywords}
 
 %%%%%%%%%%%%%%%%%%%%%%%%%%%%%%%%%%%%%%%%%%%%
 \section{Introduction}
 %%%%%%%%%%%%%%%%%%%%%%%%%%%%%%%%%%%%%%%%%
 mRNA translation is a fundamental process in gene  expression, i.e., the transformation of  genetic information into a functional  protein~\cite{Alberts2002}.
 During translation, 
 the ribosomes (complex macro-molecules) scan the mRNA in a sequential manner, ``read'' it codon by codon, and build a corresponding chain of amino-acids. When the ribosome reaches a stop codon, it detaches from the mRNA and releases the chain that, after additional  processing, becomes a functional protein. 
 
 Many ribosomes may attach to the same mRNA and 
 translate it in parallel. This    pipelining 
 increases the protein production rate.  The speed   of the ribosome movement along the  mRNA  is determined by the mRNA sequence and structure, 
 pauses due to collisions with other ribosomes, 
 and by various translation factors,
 e.g., the abundance of cognate tRNA molecules in the vicinity of the mRNA.
 Understanding the dynamics of translation is of considerable importance,
 as it plays a major role in determining the protein production rate~\cite{Alon_gExp,TullerGB2011}. 
Furthermore, the dynamics of
mRNA translation and specifically the evolution of 
 ribosomal ``traffic jams''~\cite{tuller_traffic_jams2018,HAN2020107610} along an mRNA have been implicated to  various diseases~\cite{traffic_jams_in_neuro,Sauna2011}. 
 
 
Computational models for ribosome
 flow along an mRNA and the resulting protein production rate  can be used to integrate and explain the growing number of experimental findings (e.g., via methods like ribosome profiling~\cite{ribosome_profiling_rev} that can be performed for single cells~\cite{Insberghe2021},
 and methods for  
 imaging the translation of a single mRNA molecule~\cite{natrevtrans}). 
 Such models can also 
  predict the effect of various manipulations or regulation of the genetic machinery on the protein production rate. Such manipulations are common both in biotechnology and also during  viral infection, where the virus 
   ``hijacks'' and potentially shuts down parts of the host cell translation  machinery. For example, SARS-CoV-2 uses a multipronged strategy to impede host protein synthesis and affect translation in a global manner~\cite{finkel2021}. In addition, heterologous genes tend to overload the   
   translation machinery and also have a  global effect on translation~\cite{Sabi2019,Ceroni2015}.
 
 
 A popular model for ribosome flow, and many other natural and artificial systems and processes, is the totally asymmetric simple  exclusion process~(TASEP) (see, e.g.,~\cite{TASEP_book,solvers_guide}). This is a stochastic discrete-time process that includes a  one dimensional chain of sites and particles that hop stochastically along  the chain
 in a unidirectional manner. Simple exclusion refers to the fact that two particles cannot be in the same site at the same time, i.e., a particle can only hop to an empty site. In the context of translation, the particles are ribosomes  and the sites are consecutive  (groups of) codons along the mRNA~\cite{TASEP_tutorial_2011,Zur2016,RANDOM_ATTRACTION_TASEP}. Simple exclusion corresponds to the fact that a ribosome  cannot ``overtake'' a ribosome  in front  of it, as the information on the mRNA must be decoded in a sequential manner. TASEP has become a phenomenological model in statistical mechanics, yet rigorous analysis of this model is difficult, except for some very special
 cases, e.g., when all the internal hopping rates are assumed to be equal~\cite{solvers_guide}.  
 
 
 
 The ribosome  flow model~(RFM) is the dynamic mean-field approximation of TASEP.
 This yields a continuous-time, deterministic, non-linear model for the flow of ribosomes along the mRNA~\cite{reuveni}. The~RFM is highly amenable to analysis using tools from systems and control theory, and it has been used to model and analyze many 
 important aspects of translation including: entrainment 
 of the protein production rate to periodic initiation 
 and elongation rates with a common period~\cite{RFM_entrain},
 sensitivity analysis of the steady-state production rate~\cite{RFM_sense},
 optimizing the protein production rate subject to convex constraints  on
 the rates~\cite{rfm_concave}, the effect of ribosome 
 recycling~\cite{RFM_feedback,RFMR}, determining the ribosome 
 density that maximizes protein production~\cite{rfM_density_max},
 stochastic variability  in translation~\cite{RFM_RANDOM}, maximizing the average protein production~\cite{gain_entr_RFM},
 and more~\cite{EYAL_RFMD1}.
 
 
 
 The  cell is a factory for producing proteins that 
 includes a large number of mRNA molecules and ribosomes. 
 For example, an    \emph{S. cerevisiae} cell includes  about  60,000 mRNA molecules
and~200,000 ribosomes. About 85\%
of the   ribosomes are associated with mRNAs~\cite{mehra2003,Zenklusen2008,Zur2020}, and the rest form the  ``pool of free ribosomes''.  The mRNAs thus indirectly ``compete''  for the available  ribosomes.  
  This generates  a network with   intricate indirect coupling between the  mRNAs.
For example,   ribosomal ``traffic jams'' 
on    mRNAs may deplete the pool leading to
 lower  initiation  rates in the other mRNAs.   
When the shared resources are abundant the coupling  between mRNAs due to this  competition is weak, and the network can potentially  be analyzed using models of translation along a single, isolated  RNA. However, when the resources are scarce, e.g., when the pool of free ribosomes is starved, the competition may have a strong effect on the global 
dynamics of translation in the cell. 

The latter  scenario is relevant both in synthetic systems, where the goal is to optimize the production rate, and 
under physiological conditions.
For example,   under stress conditions or during a high yield viral infection, where  the viral mRNAs ``hijack'' the translation machinery, and consume many of the shared resources to produce viral proteins. It is also   relevant in     heterologous gene expression (e.g., when the heterologous gene is overexpressed and consumes most of the ribosomes in the cell) and
in  cell free systems where  the number of mRNA molecules may be  relatively large in comparison to the number of ribosomes~\cite{10.3389/fbioe.2019.00248}.

We study this scenario 
using     a mathematical model that includes a network of~$m$ ribosome flow models~(RFMs)  interconnected via a pool of free ribosomes. 
The RFM belongs to the class of
compartmental models that have been used in various domains of systems biology including genetics, physiology, and pharmacology~\cite{sandberg78}. 
Each RFM describes 
the dynamics of translation 
along one  mRNA molecule, and the interconnection via the pool encapsulates the  competition for shared resources. This model was first suggested  in~\cite{RFM_model_compete_J}. Mathematically, it is a cooperative dynamical system~\cite{hlsmith,mcs_angeli_2003} that admits a first integral:
the total density of ribosomes in the network is conserved. 
Such systems have a well-ordered asymptotic behaviour~\cite{Mier991,banaji_angeli}.
It was shown in~\cite{RFM_model_compete_J} that any solution of the  network  converges to a steady state, where the flow of ribosomes into and out of any site along   any  mRNA  is equal.
Also, the flows in and out of the pool are equal. 
%%
Sensitivity analysis of this steady state~\cite{RFM_model_compete_J} with respect to modifying one of the translation rates in a specific  mRNA 
demonstrated
 that  the steady-state production rates in all the other mRNAs either all increase or all decrease. 

 A generalization of this network, that includes the possibility  of ribosome
 drop-off and attachment at each site along the~RFM,
 was  recently suggested in~\cite{Aditi_competition_2021}. Simulations of this model showed that ribosome drop-off from an isolated  mRNA always decreases the protein production rate, yet in the network ribosome drop-off from a jammed mRNA may increase the total production rate of all the mRNAs, as the drop-off frees
 ribosomes that enter  the pool,
 and this improves
 the production rate in the other mRNAs. This illustrates how the network perspective provides  new biological insights.

In this paper, we analyze this  network of~$m$  RFMs interconnected via a pool from a new, structural  perspective, that is, we study how adding new RFMs to the network affects the dynamics.  
The main contributions of this paper include the following:
 \begin{enumerate}
 %%
 \item We prove  that adding an RFM to the network always decreases the steady state pool density. This makes sense, as every new RFM ``consumes'' ribosomes; 
  \item We show that adding an RFM 
 has an intricate effect on the network steady state: 
 on the one-hand, the additional  RFM produces new proteins. On the other-hand, the other RFMs produce less proteins, as the pool that feeds these RFMs now has a smaller abundance of ribosomes; 
 %%%%%%%
 \item We provide a detailed asymptotic  analysis of the network steady  state when the pool is starved. In particular, we show that in this case the initiation rates in every RFM become the bottleneck rates, and provide a closed-form expression for the total steady state density  and total production rate on any subset of RFMs in the network, relative to the steady state pool density.
  \item We provide an explicit bound for the total production rate of the network when  the number  of RFMs is very large. In particular, we show how  the total density of ribosomes in the networks bounds the total production rate. This bound demonstrates that  when the number of RFMs increases the marginal utility  of adding another RFM diminishes, and the total protein production   saturates to a limiting value. 
  %
  \item We demonstrate our analysis approach for the case of producing insulin proteins in a  cell free system, and show how it can provide useful guidelines for setting the parameters in such a system. 
\end{enumerate}

 
The remainder of this paper is organized as follows. The next section reviews the network model. Before going into the mathematical analysis, Section~\ref{sec:simulation}
 illustrates several  structural questions that can be studied  using simulations of the model. Section~\ref{sec:main} details the main mathematical results. Section ~\ref{sec:Biology} reports  analyses of a biological system (gene expression in a cell free system) with our models.  For the sake of readability, all the proofs are placed in the Appendix. Throughout, we try to describe the  biological implications of 
 every  analysis result. 

 
 
 
 
 %%%%%%%%%%%%%%%%%%%%%%
 \section{The mathematical model}
 %%%%%%%%%%%%%%%%%%%%%%%%%
We use a model that includes~$m$ RFMs interconnected via a pool of free ribosomes. Each RFM models the dynamics of ribosome flow along one  mRNA molecule and, in particular, 
each RFM may have a different length and different parameters (i.e., different codon decoding rates and initiation rates). 
The pool of free ribosomes represents 
ribosomes in the cell that are not attached  to any mRNA. 
 We begin by  reviewing  
 the various 
 components  of this networks. 
 
\subsubsection{Ribosome flow model}
%%%%%%%%%%%%%%%%%%%%%%%%%%%%%%%%%%%
The RFM includes $n$ state-variables~$x_1,\dots,x_n$ representing the normalized ribosome  density in~$n$ sites along the mRNA, where each site corresponds to a group of consecutive codons. The density is normalized such that~$x_i(t)\in[0,1]$ for all~$t$, where~$x_i(t)=0$ represents that the site is empty, and~$x_i(t)=1$
represents that the site is completely full. Thus,~$x_i(t)$ may also be interpreted as the probability that site~$i$ is occupied at time~$t$.
%%%%%%%%%%%%%%%
The RFM also includes~$n+1$ positive parameters~$\lambda_0,\dots,\lambda_n$, where~$\lambda_i$ controls  
the transition rate from site~$i$ to site~$i+1$. In particular~$\lambda_0$ controls  the initiation rate,  and~$\lambda_n$ controls the exit rate. 

The dynamics of the RFM is described by~$n$ balance equations:
\begin{align}\label{eq:rfm}
%%%%%%%
\dot x_1 & =\lambda_0 (1-x_1)-\lambda_1 x_1(1-x_2),\nonumber \\
%%%%%%%%%%%%%%%%%%%%%%
\dot x_2 & =\lambda_1x_1 (1-x_2)-\lambda_2 x_2(1-x_3),\nonumber\\
%%%%%%%%%%%%%%%%
&\vdots\\
%%%%%%%%%%%%%%%%%%%%%%
\dot x_n & =\lambda_{n-1}x_{n-1} (1-x_n)-\lambda_n x_n.\nonumber
%%%%%%%%%%%%%%%%
%%%%%%%%%%%%%
\end{align}
To explain this, consider the equation for the  change in density in the second site, namely, 
\be\label{eq:x2eq}
\dot x_2  =\lambda_1x_1 (1-x_2)-\lambda_2 x_2(1-x_3).
\ee
The term~$\lambda_1x_1 (1-x_2)$ represents the flow of ribosomes  from site~$1$ to site~$2$. This is proportional to the transition rate~$\lambda_1$, to the density of ribosomes~$x_1$ in site~$1$ and to the ``free space'' $(1-x_2)$ in site~$2$.
In particular, if site~$2$ fills up, i.e.,~$x_2$ is close to one, then the flow into site~$2$ decreases to zero. This is a ``soft'' version of the simple exclusion principle, i.e., the notion  that two particles cannot be in the same place at the same time. Similarly, the second term on the right-hand side of~\eqref{eq:x2eq} is the flow from site~$2$ to site~$3$. Thus, Eq.~\eqref{eq:x2eq} states that the change in density in site~$2$ is the flow from site~$1$ to site~$2$ minus the flow from  site~$2$ to site~$3$. 
%%%%%%%%%%%%%%%%%%%%%%%%%
The exit rate from the last site is~$R(t):=\lambda_n x_n(t)$, and this is also the protein production rate at time~$t$ (see Fig.~\ref{fig:rfm_flow}). 
%%%%%%%%%%%%%%%%%%%%
 Note that~$x_i$ is dimensionless, and that~$\lambda_i$ has units  of~$1/\text{time}$. In all the biological simulations below, $\lambda_i$ is in units of~$1/\text{sec}$.
 
\begin{figure}[t]
\centering
\includegraphics[width=1\linewidth]{diagram_1_eps_type.eps}  
\caption{Ribosome flow model.}
\label{fig:rfm_flow}
\end{figure} 

%\begin{figure}[t]
%\centering
%\includegraphics[width=1\linewidth]{diagram_2_for_eps_option_2.eps}  
%\caption{RFMNP model - Option 2.}
%\label{fig:rfm_flow}
%\end{figure} 

The state space of the RFM  is the~$n$-dimensional cube~$[0,1]^n$. Since this invariant set is compact and convex, the RFM admits an  equilibrium point~$e\in[0,1]^n$. At an equilibrium, the flows  into each site and out of each site are equal, so all the site densities are constant.  Analysis of the equations describing the equilibrium point shows that there is a single equilibrium point~$e\in [0,1]^n$~\cite{RFM_stability}. 
 

The RFM has been extensively used for studying  the translation of a single, isolated mRNA. The model is highly amenable  to analysis using tools from systems and control theory. It was shown in~\cite{RFM_stability} that the~RFM is a totally positive differential system~\cite{fulltppaper} and this implies that any solution of the~RFM converges to the unique equilibrium~$e$. In particular, the protein production rate~$R(t)=\lambda_n x_n(t)$ converges
to the steady-state production rate~$R:=\lambda_n e_n$.
In other words, the positive transition rates~$\lambda_0,\dots,\lambda_n$ determine a unique  steady-state density $x_1 = e_1,\dots, x_n=e_n$ along the mRNA, and 
for any initial density the dynamics converges
to this profile.  


%Ref.~\cite{RFM_entrain}  showed that the RFM is an (almost) contractive system. This implies that if we assume that the rates~$\lambda_i$ are not constant, but time-varying, positive functions of time and all have a common period~$T$ then the model admits a unique~$T$ periodic solution~$\gamma:[0,T)\to [0,1]^n$ and any solution of this periodic RFM converges to~$\gamma$. In other words, if we view the time varying~$\lambda_i$s as a periodic excitation then the dynamics \emph{entrains} to this excitation. 

Ref.~\cite{rfm_concave} derived a useful 
\emph{spectral representation} for the mapping from the rates~$\lambda_0,\dots,\lambda_n$ to the steady state~$e$. Given the RFM, consider the~$(n+2)\times(n+2)$ tri-diagonal matrix
\begin{equation}\label{eq:spetral_matrix}
A:=  \begin{bmatrix}
 0& \lambda_0^{-1/2}& 0 &0 &\dots &0 &0 &0\\
 %%%%%%%%%%%%%
\lambda_0^{-1/2}& 0&\lambda_1^{-1/2} & 0&\dots &0 &0 &0\\
%%%%%%%%%%%%%%%%%
 0& \lambda_1^{-1/2}& 0 &\lambda_2^{-1/2} &\dots &0 &0 &0\\
%
 &&\vdots\\
 %%%%%%%%%%%%%%%%%%%%%
 0& 0& 0 &0 &\dots &\lambda_{n-1}^{-1/2} & 0&\lambda_n^{-1/2}\\ 
 0& 0& 0 &0 &\dots&0 &\lambda_n^{-1/2} &0  
%%%%%%%%%%% 
\end{bmatrix}.
\end{equation}
Since~$A$ is symmetric, all its eigenvalues are real. Since~$A$ is an irreducible matrix, with all entries   non-negative, the Perron-Frobenius theorem~\cite{matrx_ana} implies that~$A$  admits a simple maximal eigenvalue~$\sigma>0$, and the 
corresponding eigenvector~$\zeta\in\R^{n+2}$ is unique (up to scaling) 
and
satisfies~$\zeta_i>0$ for all~$i\in\{1,\dots,n+2\}$. Then, the entries of~$e$ satisfy~\cite{EYAL_RFMD1}:
%%%%%%%%%%%%%%%%%%%%%%%%%%%%%%
\be\label{eq:ei_spectral}
e_i =  \frac{\zeta_{i+2}}{ \lambda_i^{1/2}\sigma \zeta_{i+1}},\quad i=1,\dots,n,
\ee
and   
the steady-state production rate satisfies 
\be\label{eq:rsigma}
R=\sigma^{-2}. 
\ee
In other words, the Perron eigenvalue and eigenvector of~$A$ provide all
the information needed to determine the steady state profile~$e$, and the  steady state production rate~$R$ in the~RFM. 


This spectral representation has several  implications. For example, 
it implies that it is possible to numerically calculate efficiently the steady state even for very large RFMs
using algorithms for computing the Perron eigenvalue and eigenvector of tri-diagonal matrices. 
Also, it follows from~\eqref{eq:rsigma}  that  the function~$R=R(\lambda_0,\dots,\lambda_n)$ is strictly concave~\cite{rfm_concave}, thus allowing to show that  general steady-state protein production optimization problems are  convex optimization problems~\cite{rfm_concave}. It also reduces the sensitivity analysis of~$R$ with respect to any rate~$\lambda_i$ to an eigenvalue sensitivity problem for the matrix~$A$~\cite{RFM_sense}. 


\subsubsection{Ribosome flow model with input and output~(RFMIO)}
%%%%%%%%%%%%%%%%%%%%%%%%%%%%%%%%%%%
As noted above, a cell typically includes a large number of mRNA molecules and ribosomes, that compete for shared resources, 
and studying translation on a single, isolated mRNA  may thus provide limited insight on large-sale  translation in the cell. To model  translation in the cell requires a network of interconnected~RFMs. 
The first step in building such a network is adding an input and output to the~RFM. 
This yields the \emph{ribosome flow model with input and output~(RFMIO)}~\cite{RFM_model_compete_J}.
The   RFMIO  dynamics is 
 \begin{align}\label{eq:rfmio}
%%%%%%%
\dot x_1 &  = u \lambda_0 (1-x_1)-\lambda_1 x_1(1-x_2),\nonumber \\
%%%%%%%%%%%%%%%%%%%%%%
\dot x_2 & =\lambda_1x_1 (1-x_2)-\lambda_2 x_2(1-x_3),\nonumber\\
%%%%%%%%%%%%%%%%
&\vdots\\
%%%%%%%%%%%%%%%%%%%%%%
\dot x_n & =\lambda_{n-1}x_{n-1} (1-x_n)-\lambda_n x_n,
\nonumber\\
y&=\lambda_n x_n.\nonumber 
%%%%%%%%%%%%%%%%
%%%%%%%%%%%%%
\end{align}
%%%%%%%%%%%%%%%%%%%%%%
The scalar input~$u:\R_+\to\R_+$ represents the density of ribosomes in the vicinity of the initiation site.
Thus, if~$u(t)$ is large then the effective  initiation rate at time~$t$, given by~$ u (t) \lambda_0  $, increases.
The scalar output~$y(t)=\lambda_nx_n(t)$ is the rate of ribosomes exiting the mRNA at time~$t$. 
The additional input and output allow to connect RFMIOs in a network. 
Note that for~$u(t)\equiv 1$,
Eq.~\eqref{eq:rfmio} reduces to the~RFM.

\begin{figure}[t]
\centering
\includegraphics[width=1\linewidth]{diagram_2_for_eps_option_1.eps}  
\caption{The network includes $m$ RFMIOs interconnected via  a pool of free ribosomes. For illustration only, we assume that RFMIO~$\#1$ has length~$n_1=2$ and write its equations explicitly.  }
\label{fig:network}
\end{figure} 

\subsubsection{The network}
%%%%%%%%%%%%%%%%%%%%%%%%%%%%%%%%%%%
Ref.~\cite{RFM_model_compete_J} introduced  a model composed of~$m$ RFMIOs interconnected via  a pool of free ribosomes  (see Fig.~\ref{fig:network}). Let~$n_i$, $i=1,\dots,m$, denote the length of RFMIO~$\#i$.
The state-variables in  RFMIO~$\#i$  are denoted by~$x^i_1, \dots,x^i_{n_i}$.
The density in the pool at time~$t$ is modeled by the scalar function~$z(t)$.
 The ribosomes that 
 initiate translation in  RFMIO $\#i$ are supplied from  the
 pool through the pool output  function~$G_i(z(t))$.
 Thus, the input to 
 RFMIO~$\#i$ is~$u_i(t)=G_i(z(t))$, so the 
effective  initiation rate  in RFMIO~$\#i$   is~$\lambda_0^i G_i(z(t))  $.
 
 The functions~$G_i:\R_+\to \R_+$ satisfy~$G_i(0)=0$ (i.e., when the pool is empty the initiation rate in the RFMIO is zero), and~$G_i(z) $ is continuous and strictly   increasing 
 in~$z$ (i.e., an increase  in the pool density yields an increase  in the initiation rates). Many possible functions satisfy these constraints, e.g.,~$G_i(z)= cz$, with~$c>0$, and 
 the uniformly bounded function~$G_i(z)=\alpha \tanh(\beta z)$, with~$\alpha,\beta>0$. 
  
 The pool feeds all the RFMIOs, and  is fed by the ribosomes exiting all the RFMIOs, so the balance  equation for the change  in~$z(t)$ is 
 %%%%%%%
\be\label{eq:zpool_eq}
%%
\dot z=\sum_{i=1}^m y^i -\sum_{i=1}^m  \lambda_0^i G_i(z) (1-x_1^i), %%%%
\ee
where~$y^i$ is the ribosome  exit rate from RFMIO~$\#i$. 

Let
\be\label{eq:def_st}
s(t):= z(t)+\sum_{i=1}^m \sum_{j=1}^{n_i} x^i_j(t)
\ee
denote the  total density of ribosomes in the network at time~$t$.
Since ribosomes cannot leave nor enter the network, 
\be\label{eq:stfirstint}
s(t)=s(0) \text{ for all } t\geq 0. 
\ee
In other words,~$s(t)$ is a first integral of the dynamics. 

Ref.~\cite{RFM_model_compete_J} used results on cooperative systems with a first  integral (see. e.g.,~\cite{Mier991,banaji_angeli})
to prove that any solution of the network converges to a steady-state. More precisely, for any~$p\geq 0$, let~$L_p$ denote the~$p$ level set of the first integral, that is,
$L_p$ includes all the initial conditions of pool and RFMIO densities with total density~$s(0)=p$. 
Then~$L_p$ includes a single equilibrium point 
and any trajectory emanating from an initial condition in~$L_p$ converges to this equilibrium point. In other words, any two initial conditions of the network with the same total ribosome density will converge to the same equilibrium state. At this state, the total   density   is distributed along the mRNAs and the pool such that the flow into and out of each site are equal. 

%%%%%%%%%%

Let~$e_z\in [0,s(0)]$ denote 
the  steady-state pool density,  % I am not sure that "density" is the right word here ?
and let~$e^i_j$ denote the steady-state density in site~$j$ in RFMIO~$\#i$. 
Also, let
\begin{equation}\label{eq:SSVect}
e:=\begin{bmatrix} 
e_z& e^1_1&\dots&e^1_{n_1} &\dots&
 e^m_1&\dots&e^m_{n_m} 
\end{bmatrix}^T, 
\end{equation}
i.e.,~$e$ collects all the steady state values in the network. 

\section{Simulation Results}\label{sec:simulation}
%%%%%%%%%%%
We begin with several synthetic simulation  results that demonstrate 
the wealth of biological questions that can be addressed using the network model when allowing the number of RFMIOs~$m$ to vary, that is, when mRNAs are added or removed from the network. This also illustrates our general analysis approach that combines the spectral   representation of the steady state in every RFMIO with the equation describing the first integral of the network. 

%%%%%%%%%%%%%%%%%%%%
We begin by considering  a network that includes~$m$ identical RFMIOs, where each RFMIO has length~$n_i=2$, $i=1,\dots,m$.
We also assume that every        RFMIO is homogeneous,  with $\lambda_0=\lambda_1=\lambda_2=1$. 
(Note, however, that all the theoretical results in Section~\ref{sec:main} below hold for general lengths and rates.)
We also assume   that~$G_i(z)=z$ for all~$i$ (i.e., the effective  initiation rate is proportional to the number of free ribosomes in the pool). 

To apply the  spectral approach to each RFMIO in the network,  let 
\begin{equation}\label{eq:spetral_matrix_exa_1}
A(c):=   \begin{bmatrix}
 0& c& 0 &0  \\
 %%%%%%%%%%%%%
c& 0& 1 & 0 \\
%%%%%%%%%%%%%%%%%
 0& 1& 0 &1  \\
%
 0& 0& 1 &0  
%%%%%%%%%%% 
\end{bmatrix},
\end{equation}
where~$c:=e_z^{-1/2}$.
Note that this is exactly the matrix~\eqref{eq:spetral_matrix} with~$n=2$, $\lambda_0 G(e_z)=e_z$, and~$\lambda_i=1$ for~$i=1,2$.
The Perron root of~$A(c)$   is
\[
\sigma(c)= \frac{\sqrt{\sqrt{c^4+4}+c^2+2}}{\sqrt{2}},
\]
and the corresponding  Perron eigenvector is
\[
\zeta(c) = \begin{bmatrix}
\frac{\left(\sqrt{c^4+4}+c^2-2\right)
   \sqrt{\sqrt{c^4+4}+c^2+2}}{2 \sqrt{2} c} & \frac{1}{2}
   \left(\sqrt{c^4+4}+c^2\right) & \frac{\sqrt{\sqrt{c^4+4}+c^2+2}
   }{\sqrt{2}} & 1 \end{bmatrix}^T.
\]
It follows from~\eqref{eq:ei_spectral} that the steady-state densities in every RFMIO
are
\begin{align}\label{eq:e1e2c}
e_1(c)&=  \frac{2}{\sqrt{c^4+4}+c^2},\nonumber\\
e_2(c)&=  \frac{2}{\sqrt{c^4+4}+c^2+2}  , 
\end{align}
%%
and  since~$\lambda_2=1$,   the steady-state production rate of each RFMIO is  
\[
R(c)= e_2(c). 
\]
The equation for the total density of ribosomes~$s$ in the network is
\begin{align}\label{eq:seq}
s&=e_z +m(e_1(c)+e_2(c)) \\
&=  c^{-2} +m(e_1(c)+e_2(c)).\nonumber
%%%  
\end{align}
%%%%%%%%%%%%%%%
Combining this with~\eqref{eq:e1e2c} provides an explicit expression  for~$s$ as a function of~$c$. 
This can be inverted (at least numerically) to conclude for every total density~$s$ the
corresponding~$c$ (and thus~$e_z$), and then  the spectral approach allows to  obtain  all the steady state profiles in all the~RFMIOs.


The network allows to study how important steady state quantities depend on
the number of RFMIOs in the network. We first define several such quantities. 
The ratio between the density of ribosomes in the pool and the total density of ribosomes in the network   is
\[
q:=  {e_z}/{s}. 
\]
The total protein production at steady state, denoted~$\text{TPR}$,  is the production rate of all the RFMIOs in the network. 
Since there are~$m$ identical RFMIOs,  
$$\text{TPR}=m e_2(c).$$
%%
As~$m$ is increased, more ribosomes are attached to mRNAs
and thus we can expect the steady state pool density~$e_z$ to go to zero. Then the initiation rate $\lambda_0G(e_z)=e_z$ in each RFMIO becomes the bottleneck rate, and thus~$e_i \approx e_z$ for~$i=1,2$, in every RFMIO. Substituting this in~\eqref{eq:seq} gives~$e_z \approx s/(1+2m)$, and  the total production rate is then
\[ \text{TPR} =me_2 \approx m e_z \approx ms/(1+2m).
\]
Thus, for a large~$m$ we expect the total steady state production rate in the network
to converge to~$s/2$.

Figure~\ref{fig:hopping_2}
depicts   the exact network steady state values for~$s=50$ as a function of the number of RFMIOs~$m$.  
It may be seen that: (1)~the steady state densities~$e_1$ and~$e_2$ in every RFMIO decrease monotonically   with~$m$. The same holds for~$q=e_z/s$; and 
%Note that~$q$ reaches the value~$0.1$ at approximately for~$m=30$ (but of course this may be different for other values of~$s$). 
(2)~the  total production rate~$me_2$ increases monotonically with~$m$, and converges to a saturation value of~$s/2=25$. 

Summarizing, the addition  of mRNAs  increases the total  protein production, as ribosomes now also translate the new mRNAs. However, it decreases the translation rate of 
the other  mRNAs by depleting the pool of free ribosomes. As the number of mRNAs increases, the marginal utility in adding another mRNA decreases to zero. The total production rate in the network is bounded, and an important factor in this bound is the total density of ribosomes in the network. 

These simulation results suggest that 
when optimizing the production rate of a synthetic system (e.g., in  high-throughput  heterologous gene expression)  it nay be useful
to use mRNA levels   below
the regime where  the marginal utility in adding another mRNA become negligible.

The next section provides a rigorous treatment of   
these topics.


\begin{figure}[t]
\begin{subfigure}{.5\textwidth}
  \centering
  \includegraphics[width=1\linewidth]{all_eis_as_func_m_s50.eps}  
  \caption{}
  \label{subfig1_e1e2}
\end{subfigure}
\begin{subfigure}{.5\textwidth}
  \centering
  \includegraphics[width=1\linewidth]{tot_prate_as_func_m_s50.eps}  
  \caption{}
  \label{subfig_total}
\end{subfigure}
\caption{ (a) steady-states~$e_1$, $e_2$ and~$e_z/s$ as a function of the number of RFMIOs~$m$ when~$s=50$; (b)~total production rate~$m e_2$  in the network as a function of the number of RFMIOs~$m$.}
\label{fig:hopping_2}
\end{figure}

 

%%%%%%%%%%%%%%%%%
\section{Main results}\label{sec:main}
%%%%%%%%%%%%%%%


We begin by considering the effect of adding an RFMIO to the network. We introduce some notation. 
Recall that RFMIO~$\#i$ is  
characterized by the tuple 
\begin{equation}\label{eq:quadruple}
%%%%%%%%%%%%%%%
\mathcal{T}_i:=\left \{ 
n_i,G_i, \lambda^i_0,\dots,\lambda^i_{n_i} \right\}, 
\end{equation}
where $n_i$ is the length of   RFMIO~$\#i$,  $G_i:\R_+\to \R_+$ is the $ i$th pool output function, 
and $ \lambda^i_j  $ are the rates along  RFMIO~$\#i$.


 Consider a network with~$m+1$ RFMIOs obtained by adding an RFMIO to a network of $m$ RFMIOs. Let~$e_z(m)$ [$e_z(m+1)$] denote the pool density in the network with~$m$ [$m+1$] RFMIOs. 
What is the relation between the steady state pool densities before and after adding   RFMIO~$\#(m+1)$?
The next result shows 
that~$e_z(m+1)$ is always smaller than~$e_z(m)$. 

%%%%%%%%%%%%%%%%%%%%%%%%%%%
\begin{Proposition}\label{Prop:Monotonicity}
Fix~$s>0$, and a collection of RFMIOs~$\mathcal{T}_1,\mathcal{T}_2,\dots$. For any $m$, consider a network of $m$ RFMIOs $\mathcal{T}_1,\dots, \mathcal{T}_m$ interconnected via a pool of free ribosomes,   with   total ribosome  density~$s  $.
%%
Let~$e(m)$ denote  the corresponding network steady-state  (see~\eqref{eq:SSVect}), where the  coordinates    depend  on~$m$. Then 
\[
0<e_z(m+1)<e_z(m).
\]
\end{Proposition}
The proof is placed in the Appendix.   

In other words, adding an RFMIO to the network, while keeping the total ribosome  density constant,  always decreases the steady  state pool density. This makes sense, as the new mRNA ``consumes'' ribosomes from the pool. 

Proposition~\ref{Prop:Monotonicity} implies in  particular  that
in  a network built by repeatedly adding new RFMIOs   the sequence of  steady-state pool densities $e_z(1), e_z(2),\dots$ is monotonically decreasing.
  Since~$e_z(m)\geq 0$ for all~$m$,  this implies that  the limit
 \[
 \lim_{m\to \infty}e_z(m)
\]
 exists.
The next result shows that  this limit is   zero. Since we take~$m\to\infty$, we need to impose some technical conditions on the RFMIOs. 
\begin{Assumption}\label{assume:lam}
%%%%%%%%%%%%%%%%%%%%
From here on we always  assume that the following properties hold. %%%%
\begin{enumerate}
%%%%%%%%%%%%%%%
    % \item There exists some $n^*\in \mathbb{N}$ such that
    % \[
    % n_i\leq n^* \text{ for all } i;
    % \]
    \item 
    There exists~$\lambda_*>0 $ such that
    \[
    \lambda_0^i\geq \lambda_* \text{ for all } i;
    \]
    i.e., 
    all the initiation rates are bounded from below by~$\lambda_*$;
    %%%
    \item 
    There exists~$ \lambda^*>0$ such that
    \[
    \lambda^i_{n_i} \leq \lambda^* 
    \text{ for all } i,
    \]
    i.e., all the exit rates are bounded from above by~$\lambda^*$; 
    %%%%%%%%%
    %%%%
    \item
    There exist~$ p>0$ and~$ g_*>0$ such that for 
      any~$z \in [0,p]$, we have
      \[
      G_i(z) \geq g_* z  \text{ for all }i,
      \]
      i.e.,  all the pool output  functions~$G_i(z)$ are bounded from below by the linear function~$g_* z$ on the interval~$[0,p]$. %%%%%%%%%%%%%%%
\end{enumerate}
%%%%%%%%%%%%%%%%%
\end{Assumption}
 These three conditions  are clearly reasonable. 


  
\begin{Proposition}\label{Prop:Depletion}
Fix~$s>0$, and a collection of RFMIOs~$\mathcal{T}_1,\mathcal{T}_2,\dots$.
 %%%%%%%%%%%
 For any~$m$, consider a network of $m$ RFMIOs $\mathcal{T}_1,\dots, \mathcal{T}_m$ interconnected via a pool of free ribosomes,   with   total ribosome  density~$s$. Let~$e(m)$ denote  the corresponding network steady-state  (see~\eqref{eq:SSVect}), with coordinates  
 that depend  on~$m$. Then  
 \be \label{eq:ezgotozero}
 \lim_{m\to \infty}e_z(m) = 0.
 \ee
\end{Proposition}
%%%%%%%%%%%%%%%%%
The proof is placed in the Appendix. 
 

From a biological point of view, the case~$m\to \infty$  may seem unreasonable. However, 
Proposition~\ref{Prop:Depletion}
 implies that for a large~$m$, $e_z(m)$ will be small. 
 This scenario is the    focus of   this paper. Indeed, when the pool includes  a large number of free ribosomes 
there is little  competition, and thus the indirect coupling between the mRNAs is weak. The interesting case is thus
when  the pool is close to  being  depleted, that is, when~$e_z$ is small. In the biological context, this
may represent a cell under stress conditions, 
or under a  high yield viral infection, or when there are many ribosomal ``traffic jams'' along the mRNAs, so the pool is depleted. 
 
  The next result uses the spectral representation to analyze the steady  state densities and production rate in every RFMIO when the pool is starved. 
%%%%%%%%%%%%%%%%%%%%%%%%%%%%%%%
  \begin{Proposition}\label{Prop:Depletion_now_eis}
%%%%%%%%%%%%%%%%%%%%
Consider a network of  RFMIOs~$\mathcal{T}_1,\mathcal{T}_2,\dots$, interconnected via a pool. Suppose that either the total ribosome density~$s$ goes to zero, or that the number of RFMIOs tends to infinity, so that~$e_z\to 0$. 
 Then for any~$i$,   RFMIO~$\#i$  satisifes
%%%%%%%%%%%%%%%%%%%%%%%%%%%
    \begin{align} \label{eq:Limration}
    &\lim_{e_z\to 0}\frac{e_j^i}{G_i(e_z)}  
    =   \frac{\lambda_0^i  }{\lambda_j^i } , \quad j=1,\dots, n_i.
%%%
\end{align}
%%%%%%
% In particular, if the pool output function~$G_i$ is linear, i.e.,  
% $G_i(e_z) =  {g}_i e_z$, with $ {g}_i>0$, then 
 In particular, if the pool output function~$G_i$ is differentiable at zero, then 
%%%%%%%%%%%%
\begin{equation}\label{eq:eij_diff}
 \lim_{e_z\to 0}\frac{e_j^i}{e_z} =      \frac{\lambda_0^i   {G}_i'(0)} {\lambda_j^i}, \quad j=1,\dots, n_i.
\end{equation} 
%%%%%%%%%%%%
\end{Proposition}
%%%%%%%%%%%%%%%%%
The proof is placed in the Appendix. 

 In other words, when the pool becomes  depleted (e.g., because~$m$ is   large or the total ribosome  density~$s$ is small) 
every density along the~$i$th mRNA 
behaves asymptotically like the pool output function~$G_i(e_z)$. This makes sense, as the effective  initiation rate~$\lambda_0^i G_i(e_z) $ 
becomes the bottleneck rate in the mRNA. 
Note that~\eqref{eq:Limration}
implies that~$\frac{e^i_j}{\lambda_0^i G_i(e_z)}$ 
  is inversely proportional to~$\lambda^i_j$.
This is reasonable,  as~$\lambda^i_j$ controls the flow \emph{out} of site~$j$.  

\begin{Example}\label{exa:scer}
%%%%%%%%%%%%%%%%%%%
To illustrate Proposition~\ref{Prop:Depletion_now_eis},  we use
a network consisting of~$m$ identical RFMIOs. 
Each~RFMIO has length~$5$ and rates 
\be\label{eq:ratess}
\begin{bmatrix}0.1678 & 0.2572 & 0.2758 & 0.2514 & 0.2612 & 0.3002\end{bmatrix}.
\ee
These values are taken from~\cite{rfmdr} and correspond to the  
  S. cerevisiae gene YBL025W that encodes the protein RRN10, which is related to regulation of RNA polymerase I. This gene has 145 codons (excluding the stop codon), and was  divided into 6 consecutive groups of codons: the first group includes the first 24 codons (that are also related to later stages of initiation). 
  The other groups  include 25 non-overlapping codons each, except for the last one that includes 21 codons. This partitioning was found to optimize the correlation between the~RFM prediction and biological data measurements (see~\cite{rfmdr} for more details).
  We increased~$m$ from~$1$ to~$1000$ while keeping the total ribosome density fixed at~$s=20$, 
  thus depleting the steady-state pool density. The input functions are~$G_i(x)=x$ for all~$i$. Since all the RFMIOs are identical, it is sufficient to consider the steady state density in a single RFMIO.   
    Figure~\ref{fig:relations_for_asymptotics} 
    depicts~$e_j /e_z$, $j=1,\dots,5$,  as a function of~$(1/e_z)$. It may be seen that as~$e_z$ decreases,  
    every ratio~$e_j /e_z$ converges to the asymptotic value given  in  Proposition~\ref{Prop:Depletion_now_eis}.
%%%%%%%%%%%%%%%
\begin{comment}
Computations performence: Total calculations took~$0.84$ seconds. The simulations were implemented in MATLAB version R2021b, using MSI Stealth 15M Gaming Laptop i7-11375H, NVIDIA GeForce RTX 3060. %Tamir: maybe move this to the appendix
%%%%%%%%%%%%%%%%%%%
\end{comment}
\end{Example} 

\begin{figure}[t]
\centering
\includegraphics[width=10cm,height=10cm,keepaspectratio,width=1\linewidth]{relations_for_asymptotics_grid_3.eps}  
\caption{Ratios~$e_j/e_z$ for~$j=1,\dots,5$, as a function of~$1/e_z$.
Dashed lines  are the asymptotic values in~\eqref{eq:eij_diff}.}
%%%%%%%%%%%%%%
\label{fig:relations_for_asymptotics}
\end{figure} 

  
  
The following result  is an immediate corollary of Proposition~\ref{Prop:Depletion_now_eis}.
Recall that the constant total ribosome
density in the network is
\be\label{eq:sineq}
s=e_z+ \sum_{i=1}^m \sum_{j=1}^{n_i}  e^i_j,
\ee
and   the total steady state production rate is
\[
\text{TPR}: = \sum_{i=1}^m  \lambda^i_{n_i}e^i_{n_i}.
\]
Also, let~$q:=e_z/s$   denote the ratio between the  
free ribosomes and the total number of ribosomes in the network.  
%%%%%%%%%%%%%%%%%%%%%%%%%%%%%%%
 \begin{Corollary}\label{Corr:TotDens}
Suppose that 
the pool output functions~$G_i$ are differentiable at zero for all $i = 1,2,\dots$. Then,
\begin{enumerate}
    \item The total production rate satisfies
    \begin{align}\label{eq:rtotalatinf}
    %%%%%%%%%%%%%%%%%%%%%%%%%%%%%%
     \lim_{e_z\to 0}\frac{\text{TPR}}{e_z} = \sum_{i=1}^m
     \lambda_0^i  {G}_i'(0).
    %%%%%%%%%%%%%%
    \end{align}
    \item The ratio between the  
    free ribosomes to the total number of ribosomes in the network satisfies 
    \begin{align} \label{eq:qatinf}
    %%%%%%%%%%%%%%%%%%%%%%%%%%%
    \lim_{e_z\to 0}q = \left(1 + \sum_{i=1}^m  \frac {\lambda_0^i  {G}_i'(0)n_i} {  H (\lambda_1^i,\dots, \lambda_{n_i}^i  )} \right) ^{-1},
    \end{align}
    where~$H  (\lambda_1^i,\dots, \lambda_{n_i}^i ) := n_i \left(\sum_{j=1}^{n_i}\frac{1}{\lambda_j^i }
    \right )^{-1}$ is the harmonic mean of the rates $\lambda_1^i,\cdots, \lambda_{n_p}^i$, that is, all the rates  except for the initiaion rate.
\end{enumerate}
%%%%%%%%%%%%%%%%%%%%%%%%%%%%%%%%%%%%%%%%%
\end{Corollary} 
%%%%%%%%%%%%%%%%%%%%%%%%%%%%%%%%%%%%%%%%%%%
%%%%%%%%%%%%%%%%%%%%%%%%%%%%%%%
% \begin{Corollary}\label{Corr:TotDens}
% Suppose that 
% the pool output functions~$G_i$ are differentiable at zero for all $i = 1,2,\dots$.  
% Then
% \begin{align} \label{eq:qatinf}
% %%%%%%%%%%%%%%%%%%%%%%%%%%%
%   \lim_{e_z\to 0}q = \left(1 + \sum_{i}  \frac {\lambda_0^i  {G}_i'(0)n_i} {  H (\lambda_1^i,\dots, \lambda_{n_i}^i  )} \right) ^{-1},
% \end{align}
% %%%%
% and
% \begin{align}\label{eq:rtotalatinf}
% %%%%%%%%%%%%%%%%%%%%%%%%%%%%%%
%   \lim_{e_z\to 0}\frac{TPR}{e_z} = \sum_{i}
%   \lambda_0^i  {G}_i'(0),
% %%%%%%%%%%%%%%
% \end{align}
% where~$H  (\lambda_1^i,\dots, \lambda_{n_i}^i ) := n_i \left(\sum_{j=1}^{n_i}\frac{1}{\lambda_j^i }
% \right )^{-1}$ is the harmonic mean of the rates $\lambda_1^i,\cdots, \lambda_{n_p}^i$.
% %%%%%%%%%%%%%%%%%%%%%%%%%%%%%%%%%%%%%%%%%
% \end{Corollary}
The proof is placed in the Appendix.

These results provide   closed-form asymptotic  expressions  
for important biological 
quantities when the pool is starved. 
Note that as~$e_z\to 0$, $\frac{TPR }{e_z}$ depends   on all the initiation  rates~$\lambda_0^i$, but not on any of the other rates (since the number of ribosome along any mRNA is low, there are no traffic jams). However, the ratio between the density of ribosomes  in the pool and the total number of ribosomes in the network  does depend on the harmonic mean of all the rates in the network.


We note that a similar closed-form expression can be obtained for the total steady-state density  and total production rate on any subset of RFMIOs in the network. 

\begin{comment}
%%%%%%%%%%%%%%
 The following is an immediate corollary of Proposition~\ref{Prop:Depletion_now_eis}.
\begin{Corollary}\label{Corr:TotDens}
Under the assumptions of Proposition~\ref{Prop:Depletion_now_eis}, let the pool output functions~$G_i$ be differentiable at zero for all $i = 1,2,\dots$. Fix a subset of ribosomes $\mathcal{I}$ and denote by $\mathcal{S}(\mathcal{I}) = \sum_{p\in \mathcal{I}}\sum_{j=1}^{n_p}e_j^p$ and $R(\mathcal{I}) = \sum_{p\in \mathcal{I}}\lambda_{n_p}^pe_{n_p}^p$ the total ribosome density and the production rate of the ribosomes belonging to $\mathcal{I}$ at the steady state, respectively. Then
\begin{equation*}\label{eq:Tot}
\begin{array}{lll}
& \lim_{e_z\to 0}\frac{\mathcal{S}(\mathcal{I})}{e_z} = \sum_{p\in \mathcal{I}}\frac{\lambda_0^p  {G}_p'(0)n_p}{\text{H}\left(\lambda_1^p,\dots, \lambda_{n_p}^p \right)},\\
& \lim_{e_z\to 0}\frac{R(\mathcal{I})}{e_z} = \sum_{p\in \mathcal{I}} \frac{\lambda_0^p  {G}_p'(0)}{\lambda_{n_p}^p }.
\end{array}
\end{equation*}
Here $\text{H}\left(\lambda_1^p,\dots, \lambda_{n_p}^p \right) = n_p\left(\sum_{j=1}^{n_p}\frac{1}{\lambda_j^p }
\right)^{-1}$ is the harmonic mean of the rates $\lambda_1^p,\cdots, \lambda_{n_p}^p$.
\end{Corollary}
The proof is placed in the Appendix.
%%%%%%%
\end{comment}


  Corollary~\ref{Corr:TotDens} imlies that when the pool is starved we  
 can replace an  entire  network of~$m$ identical RFMIOs
by a much simpler network, while keeping the asymptotic steady
state properties unchanged. 
%%%%%%%%%%%%%%%%%%%
\begin{Proposition}\label{prop:Exch}
Let $s>0$ denote the  total number of ribosomes.
Consider the following two networks of RFMIOs:
\begin{enumerate}
    \item A network of $m$ identical RFMIOs, each  of length $n$, rates $\lambda_0,\cdots,\lambda_n$, and output functions~$G(z)=gz$,
    with~$g>0$. Let~$\text{TPR} $ [$q$] denote
    the total production rate [ratio between free ribosomes and $s$]  in this network.
    \item A network consisting of a single
    RFMIO of length $\overline{n}=1$, rates $\overline{\lambda}_0, \overline{\lambda}_{1}$, and $\overline{G}(z)=\overline{g}z$, with~$\overline{g}>0$. Let $\overline{TPR}$ [$\overline{q}$] denote
    the total production rate [ratio between free ribosomes and $s$]  in this  network.
\end{enumerate}
If the parameters   of the second network are chosen such that 
\begin{align}
  \overline{\lambda}_0\overline{g}  = \lambda_0 g m   ,\label{eq:ExchCond}
    \end{align}
    and
    \begin{align}
 (\overline{\lambda}_1) ^{-1} =    \sum_{j=1}^{n} (\lambda_j) ^{-1}   \label{eq:ExchCond2}
\end{align}
then, as $s\to 0$
\begin{equation}\label{eq:NetwEquiv1}
\lim_{e_z\to 0}\frac{TPR}{e_z} =    \lim_{e_z\to 0}\frac{\overline{TPR}}{e_z}, 
\end{equation}
and
\begin{equation}\label{eq:NetwEquiv2}
 \lim_{e_z\to 0}q = \lim_{e_z\to 0}\overline{q}.
\end{equation}
%%%%%%%%%%
\end{Proposition}
%%%%%%%%%%%%%%%%
In other words, we can use a network with a pool and 
a single RFMIO, with a single site,   to simulate and analyze the first network. Note that
conditions~\eqref{eq:ExchCond} and~\eqref{eq:ExchCond2} are quite intuitive. Roughly speaking,
the first implies that the initiation  rates in the two networks are equal (taking into
account that in the first network there are~$m$ RFMIOs and in the second a single RFMIO),  
whereas the second condition requires that some mean of the other rates along the RFMIO is also equal. 
%%%%%%%%%%%%%%%%%%%%%%
\begin{proof}
%%%%%%%%%%%%%%%%%%
Using \eqref{eq:rtotalatinf}, we have $\lim_{e_z\to 0}\frac{TPR}{e_z}=\lambda_0 g m$ and 
$\lim_{e_z\to 0}\frac{\overline{TPR}}{e_z}=\overline{\lambda}_0 \bar{g}$, and combining this with~\eqref{eq:ExchCond}
gives~\eqref{eq:NetwEquiv1}. Similarly, Eq.~\eqref{eq:qatinf} gives
%%%%
\begin{align*}
             \lim_{e_z\to 0}q = \frac{ g\lambda_0 m n}{H(\lambda_1,\dots,\lambda_n)} , 
             \text{ and }     \lim_{e_z\to 0}\overline{q}   = \frac{\overline{g}\overline{\lambda}_0  \overline{n}}{H(\overline{\lambda}_1,\dots, \overline{\lambda}_{\overline{n}} ) }      ,           
\end{align*}
and using \eqref{eq:ExchCond} and~\eqref{eq:ExchCond2} implies that these  expressions are  equal. 
%%%%%%%%%%%%%%
\end{proof}


\begin{Example}
Consider again the network in Example~\ref{exa:scer}. Recall that this has~$m$ identical RFMIOs of length~$n=5$ 
and the rates given in~\eqref{eq:ratess}. In this case, $g=1$, $\lambda_0=0.1678$, and
 \[
 \sum_{j=1}^{5} (\lambda_j) ^{-1} =18.6512 . 
\]
%
Proposition~\ref{prop:Exch} implies that we can replace this network of~$m$ RFMIOs by a network consisting of a single RFMIO of length one, with~$\overline{G}(x)=x$, $\overline{\lambda}_0=0.1678 m$, and~$\overline{\lambda}_1=1/18.6512$, and the asymptotic behaviour of the two networks when the pool is starved will be identical. 
%%%%%%%%%%%%%%%%%%%%%%%%%%%%%%%%%%%%%%
\end{Example} 

%   We provide two examples of this property:
% \begin{enumerate}
%     \item Consider the case of a single RFMIO, $\mathcal{I} = \left\{p\right\}$. Then, as $e_z\to 0$, $\frac{\mathcal{S}(\left\{p\right\})}{e_z}$ and $\frac{R(\left\{p\right\})}{e_z}$ tend to $\frac{\lambda_0^p  {G}_p'(0)n_p}{\text{H}\left(\lambda_1^p,\cdots, \lambda_{n_p}^p \right)}$ and $\lambda_0^p  {G}_p'(0)$, respectively.   Exchanging the single RFMIO~$\#p$ for another RFMIO that preserves the latter quantities will result in a new network, with the same total steady state ribosome density and production rate at every RFMIO, relative to the state pool density. An example of such an replacement is exchanging RFMIO~$\#p$ with a new RFMIO of length~$2n_p$ with $\bar{\lambda}_0 = \frac{\lambda_0^p}{2}$, $\bar{G}_p = 2G_p$ and any rates $\bar{\lambda}_1,\dots, \bar{\lambda}_{2n_p}$ satisfying $\text{H}\left(\bar{\lambda}_1,\dots,\bar{\lambda}_{2n_p}\right) = 2\text{H}\left(\lambda_1^p,\dots, \lambda_{n_p}^p \right)$.
%     \item As another example, recall that in Section \ref{sec:simulation} we considered a network of~$m$ identical RFMIOs of length~$n_i=2$, $i=1,\dots,m$.
%     We also assumed that~$\lambda_0=\lambda_1=\lambda_2=1$ and~$G_i(z)=z$ for all~$i$. For this network, we found that as $e_z\to 0$, $\frac{\mathcal{S}_{\text{total}}}{e_z}$ and $\frac{R_{\text{total}}}{e_z}$ tend to $2m$ and $m$, respectively. Assume that $m>1$ is a composite number with $m = a b, \quad a,b>1$. Consider a single network with a single RFMIO of length $n=2$, $G(z)=bz$, $\lambda_0 = a$, $\lambda_1 = 2$ and $\lambda_2 = \frac{2}{3}$. In this case we have
%     \begin{equation*}
%          H\left(\lambda_1,\lambda_2\right) = \frac{2}{\frac{1}{\lambda_1}+\frac{1}{\lambda_{2}}} = \frac{2}{ \frac{1}{2}+\frac{3}{2}} = 1.
%     \end{equation*}
%     Hence, in the new network, we will have
%     \begin{equation*}
%         \begin{array}{lll}
%         &\lim_{e_z\to 0} \frac{\mathcal{S}_{\text{total}}}{e_z} = \frac{\lambda_0G'(0)n}{H\left(\lambda_1,\lambda_2\right)} = \frac{2a b}{1} = 2m,\\
%         &\lim_{e_z\to 0} \frac{R_{\text{total}}}{e_z} = \lambda_0G'(0) = ab = m.
%         \end{array}
%     \end{equation*}
% \end{enumerate}

 
  
%%%%%%%%%%%%%%%%%%%%%%%%%%%


 %%%%%%%%%%%%%%%%%%%%%%%%%%%%%%%%%%%%%%%%%%%%%%%
 
\begin{comment}


max location for r=1: (0.13,0.46,0.41), val: 0.1889
max location for r=2: (0.25,0.95,0.8), val: 0.379
max location for r=4: (0.5,1.9,1.6), val: 0.7592

\end{comment}

 %%%%%%%%%%%%%%%%%%%%%%%%%%%%%%%%%%%%%
 
 
 
 %%%%%%%%%%%%%%%%%%%%%%%%%%%%%%%%%%%%%%%%%%%%%%%
\section{A biological example: production of  insulin  in a cell free system}\label{sec:Biology}
%%%%%%%%%%%%%%%%%%
In this section, we  apply our model to
analyze 
the production of 
insulin protein in a cell free system.
A cell free system is an in vitro based approach 
to study and/or generate biological reactions that take place
within cells via the isolation of relevant cellular components (e.g. ribosomes, RNA  polymerases, tRNA molecules). This approach 
 reduces the complex interactions typically found when working  with
 whole cells.  Cell free systems are often used in biotechnology for heterologous gene expression~\cite{Swartz2006}.
 

Data on the  coding region of insulin fitted to {\it S. cerevisiae} was taken from~\cite{Secher2009}. The codon decoding rates,
that are based on   the analysis of typical decoding rates in-vivo from ribo-seq data~\cite{Dana2014}, 
were taken from~\cite{Dana2014B}. Groups of 10 consecutive codons were coarse grained into one RFMIO site, as was done in previous studies (see, e.g.,~\cite{Zur2020}). This yields 12 sites. 
The transition rate~$\lambda_i$ of  site~$i$ is 
the inverse of   the sum of the decoding times along the 10 related codons.  These rates were then normalized such that 
their average value is~$10$ codons per second (the typical decoding rate in eukaryotes~\cite{Bonven1979}).\footnote{The final result of this process are the rates:
$
10.01 ,\; 10.33, \; 10.25, \; 9.97 ,\; 10.67, \; 10.59, \; 9.40, \; 9.97, \; 9.75, \; 10.67, \; 10.23, \; 10.13, \; 8.03 .
$}

%[10.01,	10.33, 10.25, 9.97, 10.67, 10.59, 9.40, 9.97, 9.75, 10.67, %10.23, 10.13, 8.03]

Since we study the translation of insulin in a cell free system, 
we assume that all the mRNAs in the system encode the protein insulin. 
 Thus, the network  includes $m$ 
 identical RFMIOs interconnected via  a pool.    

 %%%%%%%%%%%%%%%%%%%%%%%%%%%%%%%%%%%%%%%%%%%%
We set~$s=250,000$, $m=39,500$ (see the data in~\cite{Haar2008,Lackner,Zur2020}),
and assume that all the pool output functions are identical:~$G_i(x)=cx$ for all~$i=1,\dots,m$. 
To calibrate the  constant $c$, we assume
that~$c$ maximizes the effective  initiation rate while keeping it lower than all the other transition rates. Mathematically, this yields the equation
\[
\lambda_0G(e_z )=\lambda_0c e_z =\min \{\lambda_1,\dots,\lambda_n\}
\]
and this gives~$c=7.4758\cdot10^{-5}$.


 %%%%%%%%%%%%%%%%%%%%%%%%%%%%%%%%%%%%%%%%%%%%
To study the scenario
where the pool is starved, we consider
two cases: in the first   we fix~$m$ and decrease~$s$, and in the second we fix~$s$ and increase~$m$.
%%%
From a biological perspective, both cases  correspond to the fact that ribosomes may be  ``more expensive'' then mRNAs, and the goal is to optimize production while using a minimal number of ribosomes. 

\subsection{Varying the total density
of ribosomes}
%%%%%%%%%%%%%%
Consider the case where~$m=39,500$ is fixed, and~$s$ varies. 
Figure~\ref{fig:TPR/ez vs m/s0} depicts~$\text{TPR}/e_z$ (that is, the ratio between the steady-state 
total production rate and the steady-state pool density) as a function of~$m/s$ (that is, the number of mRNA molecules divided by the total number of ribosomes in the network). As expected,  $\text{TPR}/e_z$
increases with~$m/s$, and converges, as~$s\to 0$,  to the  asymptotic value~$\sum_{i=1}^m \lambda_0^i{G}_i'(0)=c  m \lambda_0  =29.56$.  
 
 
\begin{figure}[t]
\centering
\includegraphics[width=8cm,height=8cm,keepaspectratio,width=1\linewidth]{TPR_vs_ratio_more_updated.eps}  
\caption{The ratio~$\text{TPR}/e_z$ as a function of~$m/s $ for~$m=39,500$,
and~$s $ varies in the range~$[250, 12,500]$ (blue dots).
The predicted asymptotic value~$\sum_{i=1}^m \lambda_0^i{G}_i'(0)$  is marked with a dashed red line.} 
\label{fig:TPR/ez vs m/s0}
\end{figure} 

 %%%%%%%%%%%%%%%%%%%%%%%%%%%%%%%%%%%%%%%%%%%%
 \subsection{Varying the number of mRNAs}
%%%%%%%%%%%%
Consider the case where~$s=250,000$ is fixed, and~$m$ is varied.
 Figure~\ref{fig:ez vs m/s0} 
 depicts  the number of  free ribosomes~$e_z$ as a function of~$m/s$. As~$m$ increases,~$e_z$ sharply decreases, as more ribosomes attach to the additional 
 mRNAs, and thus the pool is starved. In particular,~$e_z$ goes to zero as~$m\to\infty$ (see Proposition~\ref{Prop:Depletion}). 
 In practice, when~$m/s =80$ we already get a very low value of~$e_z$,   and then the asymptotic results described in our  analysis can already   be used. 
 
 
  Note that these results  can provide important guidelines for  setting the   cell free system parameters. For example, Figure~\ref{fig:TPR/ez vs m/s0}
  shows that to achieve a steady  state production rate that is~$99\%$ of the 
maximal possible  production rate we should set $m/s = 10$, that is,
10 mRNA molecules for each ribosome in the system. 


 %%%%%%%%%
 \begin{figure}[t]
\centering
\includegraphics[width=8cm,height=8cm,keepaspectratio,width=1\linewidth]{ez_vs_ratio_s_constant_m_changes.eps}  
\caption{~$e_z$ as a function of~$m/s$ for~$s=250,000$,
and~$m$   in the  range~$[2.5, 750]\cdot10^5$.}
\label{fig:ez vs m/s0}
\end{figure} 


 %%%%%%%%%%%%%%%%%%%%%%%%%%%%%%%%%%
 \section{Discussion}\label{sec:Discussion}
 %%%%%%%%%%%%%%%%%%%%%%%%%%%%%%%%%%
 We considered a network of mRNAs fed by a pool of free ribosomes  in the scenario when the pool is starved.
 This scenario is relevant for example in biological networks   under stress conditions 
 or viral attack, and for synthetic networks where the goal is to optimize the total
 production rate. Furthermore, we believe that  studying systems under extreme
 conditions may   provide intuition 
 for how the system was designed and how it will operate
  even under normal conditions.  
 
 We used a mathematical model of a network  of RFMIOs connected via a pool of free ribosomes. 
 Using the 
 spectral representation of the RFMIO steady
 state we derived closed-form expressions for several relevant biological quantities in the regime when the pool is starved.  These include the total protein
 production rate in the network, and the ratio between the number of ribosomes in the pool and the total number of ribosomes  in the network. 
 
 
 
   We demonstrated the analytical  results using both
   synthetic examples and an example based on biological data. The results reported here can be used both in systems biology studies of natural systems and   for the     design of synthetic systems. 
 We believe that an interesting direction for further research is to use our results 
 in the biological context of a cell attacked by viruses.  
 This will allow to study both qualitatively and quantitatively important questions. For example,
 can the virus effectively shut down the host protein production (and in particular immune-related proteins) 
 by depleting the pool of free ribosomes or are other mechanisms needed? 
 
 
\section*{Appendix: Proofs}
%%%%%%%%%%%%%%%%%%%%%%%%%%%%%%%%%%%%%%%%%%%%%%%%%%%%%%%%%%%%%
For the sake of readability, we begin with a short and general description 
of our analysis approach.    Consider without loss of generality the steady-state densities~$e^1=\begin{bmatrix} e^1_1&\dots& e^1_{n_1}\end{bmatrix}^T$ along RFMIO~$\#1$ in the network.
The spectral representation  implies that we can retrieve~$e^1$ from the Perron root and Perron eigenvector of the matrix
%%%%%%%%%%%%%%%
\[
A_1:=  \begin{bmatrix}
 0& (\lambda_0^ 1 G_1(e_z))^{-1/2}& 0 &0 &\dots &0 &0 &0\\
 %%%%%%%%%%%%%
(\lambda_0^1 G_1(e_z))^{-1/2}& 0&  (\lambda_1^ 1 )^{-1/2}  & 0&\dots &0 &0 &0\\
%%%%%%%%%%%%%%%%%
 0& (\lambda_1^1)^{-1/2}& 0 &(\lambda_2^1)^{-1/2} &\dots &0 &0 &0\\
%
 &&\vdots\\
 %%%%%%%%%%%%%%%%%%%%%
 0& 0& 0 &0 &\dots &(\lambda_{n_1-1}^1)^{-1/2} & 0&(\lambda_{n_1}^1)^{-1/2}\\ 
 0& 0& 0 &0 &\dots&0 &(\lambda_{n_1}^1)^{-1/2} &0  
%%%%%%%%%%% 
\end{bmatrix}.
\]
%%
Indeed, at   steady state the pool density is~$z(t)=e_z$, so the initiation rate at RFMIO~$\#1$ is~$\lambda_0^1 G_1(z(t))=\lambda_0^1 G_1(e_z)$.

When~$e_z$ is close to zero, 
so is~$G_1(e_z)$ and this implies that entries~$(1,2)$ and~$(2,1)$ in~$A_1$ are very large. We use an asymptotic
analysis of  the spectral properties of~$A_1$ to derive an approximate expression for~$e^1$ and, similarly, for any~$e^i$, $i=1,\dots, m$. Now by~\eqref{eq:def_st} and~\eqref{eq:stfirstint},
\[
e_z=s  - \sum_{i=1}^m\sum_{j=1}^{n_i} e^i_j,
\]
and thus we obtain  the  entire steady  state of the network. 


We begin with several auxiliary  results that describe the asymptotic spectral properties of a specific tri-diagonal matrix. We use~$\R^n_+ :=\{x\in\R^n: x_i\geq 0 ,\; i=1,2,\dots,n\}$ to denote  the non-negative orthant in~$\R^n$, and
$\R^n_{++} :=\{x\in\R^n: x_i> 0 ,\; i=1,2,\dots,n\}$ to denote  the positive orthant in~$\R^n$. For a Hermitian matrix~$S\in\C^{N\times N }$ we denote its eigenvalues by
\[
\sigma_1(S)\geq\dots\geq\sigma_N(S).
\]


 

%For~$x\in\R$, let~$[x]$ denote the greatest integer less than or equal to~$x$. 
%%%%%%%%%%%%%%%
 
%%%%
\begin{Theorem}\label{thm:asymnp}
%%%%%
Given~$c>0$ and
a vector~$\alpha=\begin{bmatrix} \alpha_1,\dots,\alpha_{N-2} \end{bmatrix}^T$, with~$\alpha_i>0$, 
    consider the~$N\times N$ tri-diagonal and symmetric matrix  
\begin{equation}\label{eq:T_NSeq}
T\left[c,\alpha  \right]: = \begin{bmatrix}
 0& c& 0 &0 &\dots &0 &0 &0\\
 %%%%%%%%%%%%%
 c& 0& \alpha_1 & 0&\dots &0 &0 &0\\
%%%%%%%%%%%%%%%%%
 0& \alpha_1& 0 &\alpha_2 &\dots &0 &0 &0\\
%
 &&\vdots\\
 %%%%%%%%%%%%%%%%%%%%%
 0& 0& 0 &0 &\dots &\alpha_{N-3} & 0&\alpha_{N-2}\\ 
 0& 0& 0 &0 &\dots&0 &\alpha_{N-2} &0  
%%%%%%%%%%% 
\end{bmatrix} .
\end{equation}
Let $M:=\displaystyle \max_{1\leq j\leq N-2}\alpha_j  $. 
Then for any~$c\geq 2M$, we have 
\begin{equation}\label{eq:gmax2bnd}
    \sigma_{1}\left(T\left[c,\alpha  \right]\right) \in [c-2M,c+M], 
\end{equation}
and
\begin{equation}\label{eq:ctoinf}
 \lim_{c\to \infty}   \frac{1}{c}    \sigma_{1}\left(T\left[c,\alpha  \right]\right)  = 1.
\end{equation}
%%%%%%%%%%
Furthermore, let $\zeta(c)\in\R^N_{++}$ be the eigenvector of $T\left[c,\alpha  \right]$
corresponding to $\sigma_{1} \left(T \left[c,\alpha \right ] \right )  $, 
normalized such that its first entry is~$ \zeta_1(c)=1$.    Then
%%%%%%%%%%%%%%%%%%%%%%%%%%%%%%%%%%%%%%%%%%%
\begin{equation}\label{eq:veczeta}
\lim_{c\to \infty}\zeta(c) =  \begin{bmatrix}
1
&
1&0 &\dots&0 
\end{bmatrix}^T.
\end{equation}
%%%%%%%%%%%%%%%
More precisely,
the entries of this vector satisfy
\begin{equation}\label{eq:ZetaCoords0}
%%
\lim_{c\to \infty}\frac{c\zeta_{i+2}(c)}{\zeta_{i+1}(c)} = {\alpha_i}, 
\text{ for all } i \in \{1,\dots, N-2\}.
\end{equation}
% where we define~$\alpha_0:=1$. 
\end{Theorem}
%%%%%%%%%%%%%%%%%
 

%%%%%%%%%%%%%%%%%%%%%%%%%%
\begin{Proof}
%%%%%%%%%%%%%%%%%%%%%%%%%
Recall that a theorem of Weyl~\cite[Section~4.3]{matrx_ana} asserts that if~$A,B \in\C^{N\times N }$ are Hermitian then for any~$i,j\in \left\{1,\dots,N \right\}$, we have
 %%%%%%%%%%
\begin{align}\label{Eq:weyl}
    &\sigma_{i+j-1}(A+B)\leq \sigma_i(A)+\sigma_j(B), \quad i+j\leq N+1,\nonumber \\
    & \sigma_i(A)+\sigma_j(B)\leq \sigma_{i+j-N}(A+B), \quad i+j\geq N+1.
\end{align}
Recall also that if~$|\cdot|:\C^{N }\to\R_+$ is a vector norm, and~$||\cdot||:\C^{N\times N}\to\R_+$ is the induced matrix norm, then~$|\sigma_i(S)|\leq ||S||$ for any~$i$.


 We can now prove Theorem~\ref{thm:asymnp}.
 First note that for any~$c\geq M$, we have 
 \begin{align*}
     \sigma_{1}\left(T\left[c,\alpha  \right]\right)
     &
     \leq ||T\left[c,\alpha  \right] ||_\infty\\
     & \leq 
     M+c,
 %%%
 \end{align*}
 and this proves the upper bound in~\eqref{eq:gmax2bnd}.
 
 To prove the lower bound, fix~$c>0$. 
 Define
$A := T\left[c,0,\dots, 0\right]$, $ B: = T\left[0, \alpha \right]$.
Note that~$A+B = T\left[c, \alpha\right]$.
Applying \eqref{Eq:weyl}  with~$i=1$ and~$j=N$ gives
%
\begin{align*}
%%%%%%%%
\sigma_{1}\left(T\left[c, \alpha  \right]\right) &\geq \sigma_{1}\left(T\left[c,0,\dots, 0\right]\right)+\sigma_{N}\left(T\left[0, \alpha  \right]\right)\nonumber\\
&=   c+ \sigma_{N}\left(T\left[0, \alpha  \right]\right)\\
&\geq c-2M,
\end{align*}
%%%%%%
where the last inequality follows from the fact that
\[ 
\left|\sigma_{N}\left(T\left[0, \alpha  \right]\right)\right|\leq \left\|T\left[0, \alpha  \right] \right\|_{\infty}\leq \max_{1\leq j \leq N-3}\left\{ {\alpha_j} +  {\alpha_{j+1}} \right\}\leq 2M. 
\]
This completes the proof of~\eqref{eq:gmax2bnd}. Taking~$c\to \infty$ in~\eqref{eq:gmax2bnd} proves~\eqref{eq:ctoinf}.

To prove~\eqref{eq:veczeta}, 
let $0<c_1<c_2<\dots$ be 
such that $\lim_{k\to \infty}c_k = \infty$. To simplify the notation, let~$\zeta^k:=\zeta(c_k)\in \R^N_{++}$.   We may assume that
  every~$\zeta^k $ has norm one, and thus we can  extract a subsequence $\zeta^{k_j}$, $j=1,2,\dots$,
that converges to  a limit vector~$\zeta \in \R^N_{++}$,  that also has norm one. Then 
\begin{equation}\label{eq:EigVLim}
\zeta  = \lim_{j\to \infty}\zeta^{k_j} = \lim_{j\to \infty}\frac{T\left[c_{k_j},{\alpha}  \right] \zeta^{k_j}    }{\sigma_{1}\left(T\left[c_{k_j}, \alpha  \right]\right)}  = T\left[1,0,\dots, 0\right]\zeta .
\end{equation}
Thus, $\zeta $ is a normalized  eigenvector
of $T\left[1,0,\dots, 0\right]$
corresponding to $\sigma_{1}\left(T\left[1,0,\dots, 0\right]\right)=1$, and it is straightforward to verify that  \[\zeta =\frac{1}{\sqrt{2}}\begin{bmatrix}
1& 1&0&\dots&0
\end{bmatrix} ^T.
\]
%%%%%%%%%%%%%%%%%%%
We next claim that $\lim_{k\to \infty}\zeta^k = \zeta $. Indeed, assume the contrary. Then there is a  subsequence $\zeta^{k_\ell} \in\R^N_{++}$, $\ell=1,2,\dots$,  and   $\varepsilon>0$ such that $\left\|\zeta^{k_\ell}-\zeta \right\|_2\geq \epsilon$ for all $\ell$. Extracting a  convergent subsequence yields  a normalized  eigenvector $  \xi\in\R^N_{++}$ of $T\left[1,0,\dots, 0\right]$ corresponding to $\sigma_{1}\left(T\left[1,0,\dots, 0\right]\right)=1$, and satisfying~$\left\|  \xi -\zeta \right\|_2\geq\epsilon$. Hence, $\sigma_{1}\left(T\left[1,0,\dots, 0\right]\right)$ is not a simple eigenvalue of $T\left[1,0,\dots, 0\right]$. This contradiction completes the proof of~\eqref{eq:veczeta}.


%%%%%%%%%%%%%%%%%%%%%%%%%%%%%%%%%%%%%%%%%%%%
We now analyze  the ratios between consecutive  entries of the  Perron eigenvector of~$T[c,\alpha]$. Fix $c>0$. For simplicity, we will denote $T\left[c,\alpha   \right]$ and $\sigma_{1}\left(T\left[c,\alpha  \right] \right)$ by $T(c)$, and~$\sigma(c)$, respectively. Define the~$N\times N$   diagonal scaling matrix
\begin{equation}\label{eq:diagscale}
\begin{array}{lll}
    D: = \operatorname{diag}\left(1,1,{c,\dots, c^{N-2}} \right) ,
\end{array}
\end{equation}
  A direct calculation  gives 
\be \label{eq:MatEquiv0}
DT(c)D^{-1} = c S(c),
\ee
where
\[
S(c):= 
  \begin{bmatrix}
 0& 1& 0 &0 &\dots &0 &0 &0\\
 %%%%%%%%%%%%%
 1& 0& {\frac{\alpha_1}{c^2}} & 0&\dots &0 &0 &0\\
%%%%%%%%%%%%%%%%%
 0& \alpha_1& 0 &{\frac{\alpha_2}{c^2}} &\dots &0 &0 &0\\
%
 &&\vdots\\
 %%%%%%%%%%%%%%%%%%%%%
 0& 0& 0 &0 &\dots &\alpha_{N-3} & 0&{\frac{\alpha_{N-2}}{c^2}}\\ 
 0& 0& 0 &0 &\dots&0 &\alpha_{N-2} &0  
%%%%%%%%%%% 
\end{bmatrix}.
\]
Let 
\begin{equation}\label{eq:Rescale}
\begin{array}{lll}
        \nu(c): = D\zeta(c) = \begin{bmatrix}  \zeta_1(c)&\zeta_2(c)&{c\zeta_3(c)}&\dots& {c^{N-2}\zeta_N(c)} \end{bmatrix}^T.
\end{array}
\end{equation}
Then 
\begin{equation}\label{eq:ScalingEig}
   T(c)\zeta(c) = \sigma(c) \zeta(c) \iff  S(c) \nu(c)=\frac{\sigma(c) }{c} \nu(c).
\end{equation}
In particular
\begin{equation}\label{eq:SpecT}
\operatorname{spec}\left(S(c) \right) = \frac{1}{c}\operatorname{spec}\left(T(c) \right),
\end{equation}
where $\operatorname{spec}(A)$ is the spectrum of $A$. Define
\begin{equation}\label{eq:MatEquiv1}
S(\infty):=    \lim_{c\to \infty}S(c) = \begin{bmatrix}
 0& 1& 0 &0 &\dots &0 &0 &0\\
 %%%%%%%%%%%%%
 1& 0& 0 & 0&\dots &0 &0 &0\\
%%%%%%%%%%%%%%%%%
 0& \alpha_1& 0 &0 &\dots &0 &0 &0\\
%
 &&\vdots\\
 %%%%%%%%%%%%%%%%%%%%%
 0& 0& 0 &0 &\dots &\alpha_{N-3} & 0&0\\ 
 0& 0& 0 &0 &\dots&0 &\alpha_{N-2} &0  
%%%%%%%%%%% 
\end{bmatrix}  . 
\end{equation}
The characteristic polynomial of $S(\infty)$ is $q(z):=\operatorname{det}(zI_N-S(\infty)) = z^{N-2}(z-1)(z+1)$, so  $\sigma_{1}\left(S(\infty) \right)=1$  is a simple eigenvalue with corresponding eigenvector $\nu(\infty)$, which is given by
\begin{align}\label{eq:vinfexp}
\nu_1(\infty)& =\nu_2(\infty) = 1,\nonumber\\
%%%%%%%
\nu_j(\infty) &= \prod_{k=1}^{j-2}\alpha_k \text{ for all } j\in\{3, \dots, N\}.
\end{align}
Using arguments similar to \eqref{eq:EigVLim} and taking the limit~$c\to \infty$ in \eqref{eq:ScalingEig}, and using the fact that $\lim_{c\to \infty}\frac{\sigma(c)}{c}=1$ yields $  S(\infty) \nu(\infty)= \nu(\infty) $. By \eqref{eq:Rescale}, for any $i=1,\cdots,N-2$ we have $\frac{\nu_{i+2}(c)}{\nu_{i+1}(c)} = \frac{c\zeta_{i+2}(c)}{\zeta_{i+1}(c)}$. This 
  proves~\eqref{eq:ZetaCoords0}.
%%%%%%%%%%
%%%%%%%%%%
\end{Proof}

  


In practice~$e_z$ is positive (even if small), so it is useful to derive \emph{explicit} 
  error bounds in the asymptotic expressions. 
Note that since we are interested in ratios between entries of the Perron vector, we do not necessarily require it to be normalized. 
For two functions~$f,g:\R_+ \to \R_+$, we write~$f = \text{O}(g)$  if there exist~$c,y>0$   such that 
\be\label{eq:asympo}
\left|f(x) \right|\leq  c \left| g(x)\right|\text{ for any } x\geq y.
%%%%%%%%%%%%%%%%
\ee


The next result provides more explicit information on
the Perron eigenvector of~$T\left[c,\alpha\right]$. 

\begin{Theorem}\label{Thm:ZetaCoords}
%%
Assume that the 
  assumptions of Theorem~\ref{thm:asymnp} hold. Then as $c\to \infty$, $T\left[c,\alpha\right]$  admits
  a Perron  
  eigenvector~$\eta(c)\in\R^N_+$  satisfying 
  %%%%%%%%%%
 % for  any~$c>\sqrt{2\max_{1\leq i \leq N-2}\left(\alpha_i^2\alpha_{i-1}^{-1} \right)}$, we have 
  %%%%%%%%%%
\begin{equation}\label{eq:ZetaCoords}
\begin{array}{lll}
&\eta_1(c) =1 + \text{O}\left(c^{-2} \right),\\
&\eta_2(c) = 1 + \text{O}\left(c^{-2} \right),\\
&\eta_{i}(c) =  \alpha_1\cdots \alpha_{i-3}{\alpha_{i-2}  } 
c^{2-i}+ \text{O}\left(c^{-i} \right), \quad 3\leq i \leq N.
\end{array}
\end{equation}
 %%%%%
%%%%%%%%
\end{Theorem}
 
 In the proof of this result we actually  provide  explicit upper bounds for the  asymptotic terms in \eqref{eq:ZetaCoords}  (see Eq.~\eqref{eq:ZetaCoordsPf} below).
 
 
\begin{Proof}
 %%%%%%%%%%%%%%%
For a vector~$q\in \mathbb{R}^N$, let~$\text{sp}(q):=\{ r q\;| \; r\in\R \}$ denote the span of~$q$. Define an~$N\times N$ matrix~$R $ by
 \be\label{eq:defR}
 R:=c^2(S(c)-S(\infty)). 
 \ee
 %%%%%%%%%%%%%%%%%%%%%%
 Note that this implies that~$R$ does not depend on~$c$, and that
\be\label{eq:normofmatr}
\left\|R\right\|=\max_{1\leq i \leq N-2}{\alpha_i}  .
\ee
  %%%
  
Let~$\nu (c)$ be the Perron eigenvector of~$S(c)$ 
defined in~\eqref{eq:Rescale}, and let
%%
\begin{equation}\label{eq:Proj}
\begin{array}{lll}
    &\mathcal{P}(\infty) : = \operatorname{Proj}_{\text{sp}\left(\nu(\infty) \right)}, \qquad \mathcal{P}(c) : = \operatorname{Proj}_{\text{sp}\left(\nu(c) \right)}
\end{array}
\end{equation}
  be the projection operators on $\text{sp}\left(\nu(\infty) \right)$ and $\text{sp}\left(\nu(c) \right)$, respectively. 
  %%%%
  Let $\xi(c) := \mathcal{P}(c)\nu(\infty)$.
  Since we project on~$\text{sp}\left(\nu(c) \right)$, this implies 
  that~$\xi(c)$ is a Perron eigenvector of~$S(c)$ corresponding to~$\frac{\sigma(c)}{c}$. Let~$\eta(c) :=D^{-1}\xi(c)$. Then \eqref{eq:ScalingEig}  implies that $\eta(c)$ is a Perron  eigenvector of $T(c)$  corresponding to~$\sigma(c)$. We will show that~$\eta(c)$  satisfies~\eqref{eq:ZetaCoords}.

We begin by analyzing~$\xi(c)$. Let~$\Gamma$ be the circle in the complex plane parametrized   by  $\gamma(t) = 1+\frac{1}{2}\exp(\imagi t)$,
$ t\in [0,2\pi)$ (see Figure \ref{fig:SpectSeparateMod}). 
\begin{figure}[t]
  \centering
  \includegraphics[width=1\linewidth]{curve_in_complex_plane.eps}  
  \caption{The curve~$\Gamma$ in the complex plane. The eigenvalues of~$S(\infty)$ are~$-1,0,1$, implying that for any~$c$ large enough all   the eigenvalues of~$S(c)$, except for~$\sigma_{1} (S(c))$, are located outside of~$\Gamma$. For presentation purposes, $\sigma_{1}(S(c))$ is assumed to be on 
  the right of the point~$\begin{bmatrix}1&0\end{bmatrix}^T$. }
  \label{fig:SpectSeparateMod}
\end{figure}
\begin{Proposition}
For any $c>4\left\|R\right\|$ 
we have 
that $\sigma_{1}\left(S(c) \right)$ and $\sigma_{1} \left(S(\infty) \right)=1$ are the only eigenvalues of~$S(c)$ and $S(\infty)$, respectively, located in the interior of $\Gamma$. All  other (real) eigenvalues are to the left of the line  $\left\{s\in \mathbb{C} \ | \ \mathfrak{Re}(s) = 1/2\right\}$,
where~$\mathfrak{Re}(s)$ is the real part of~$s$.
\end{Proposition}
\begin{Proof}
%%%%%%%%%%%%%%%%%%%%%
Recall that $\operatorname{spec}\left(S(\infty) \right) = \left\{-1,0,1 \right\}$. Hence, the claim is true for $S(\infty)$. For $S(c)$, consider first the corresponding $T(c)$, satisfying \eqref{eq:MatEquiv0}. Recall that all eigenvalues of $T(c)$ are real. By Theorem \ref{thm:asymnp} and~\eqref{eq:normofmatr}, we have 
\begin{equation*}
    \frac{\sigma_{1}(T(c))}{c}\in \left[1-\frac{2\left\|R\right\|}{c},1+\frac{\left\|R\right\|}{c}\right]\subseteq \left(\frac{1}{2},\frac{5}{4} \right).
\end{equation*}
Using Weyl's inequalities \eqref{Eq:weyl}, the second largest eigenvalue of $T(c)$ satisfies
$\sigma_2(T(c))\leq 2\left\|R \right\|$. The proposition then follows from \eqref{eq:SpecT}.
\end{Proof}
It is well known~\cite[Chapter~1]{gohberg2013classes}  that the projections defined in~\eqref{eq:Proj} 
satisfy the  matrix representations:
\begin{align}\label{eq:Proj2}
%%%%%%%%%%%%%%%%%%%%%%%%%%
\mathcal{P}(\infty)& = \frac{1}{2\pi \imagi }\int_{\Gamma}\left(\lambda I_N -  S(\infty)\right)^{-1} \text{d}\lambda,\nonumber \\
\mathcal{P}(c)& = \frac{1}{2\pi \imagi }\int_{\Gamma}\left(\lambda I_N -  S(c)\right)^{-1} \text{d}\lambda.
%%%
\end{align}
We require the following auxiliary result that provides a Neumann series representation for the difference between these two projections. The proof is provided for completeness. Let
\[
M_{\Gamma}: = \max_{\lambda\in \Gamma}\left\|\left( \lambda I_N -  S(\infty)\right)^{-1} \right\|,
\]
 i.e.,  the maximal norm of the resolvent of $S(\infty)$ over $\Gamma$. This maximum exists since $\left( \lambda I_N -  S(\infty)\right)^{-1}$ is continuous in a neighborhood of $\Gamma$. 
%%%%%%%%%%%%%%%%%%%%%
\begin{Lemma}\label{lem:Aux2}
Fix~$\lambda \in \Gamma
$.
For  any $c> \max\left(\sqrt{M_{\Gamma}\left\|R \right\|},4\left\|R \right\|\right)$, we have
\begin{equation}\label{eq:Neumann}
\left( \lambda I_N -  S(c)\right)^{-1} = \left( \lambda I_N -  S(\infty)\right)^{-1}\sum_{k=0}^{\infty} \left(c^{-2}R\left( \lambda I_N -  S(\infty)\right)^{-1} \right)^k .
\end{equation}
In particular, the series on the right hand side of~\eqref{eq:Neumann}
 converges absolutely and uniformly on $\Gamma$, and
\begin{equation}\label{Neumann1}
\mathcal{P}(c) = \mathcal{P}(\infty) + \sum_{k=1}^{\infty}\frac{1}{2\pi \imagi }\int_{\Gamma}\left( \lambda I_N -  S(\infty)\right)^{-1}\left(c^{-2}R\left( \lambda I_N -  S(\infty)\right)^{-1} \right)^k\text{d}\lambda.
\end{equation}
\end{Lemma}
%%%%%%%%%%%%%%%%%%%%%
\begin{Proof}
%%%%%%%%%%%%%%%%%%%
Fix  $\lambda \in \Gamma$.
The definition of~$R$ gives
$\left( \lambda I_N -  S(c)\right)^{-1} =  \left(\lambda I_N -S(\infty) - c^{-2}R\right)^{-1} $, so 
%%%%%%%%%%%%%%%%%%%
\begin{align*} 
       \left( \lambda I_N -  S(c)\right)^{-1}   
     &= \left[\left(I_N - c^{-2}R\left( \lambda I_N -  S(\infty)\right)^{-1} \right)\left( \lambda I_N -  S(\infty)\right)\right]^{-1}\\
     & = \left( \lambda I_N -  S(\infty)\right)^{-1}\left(I_N - c^{-2}R\left( \lambda I_N -  S(\infty)\right)^{-1} \right)^{-1}\\
     & = \left( \lambda I_N -  S(\infty)\right)^{-1}\sum_{k=0}^{\infty} \left(c^{-2}R\left( \lambda I_N -  S(\infty)\right)^{-1} \right)^k,
\end{align*}
and the series  converges 
if~$\left\| c^{-2}R\left( \lambda I_N -  S(\infty)\right)^{-1}\right\|<1$. We have
%%%%%%
\begin{equation}\label{eq:qcdef}
\left\| c^{-2}R\left( \lambda I_N -  S(\infty)\right)^{-1}\right\|\leq  q(c):= c^{-2}M_{\Gamma}\left\|R \right\|,
\end{equation}
and~$q(c)<1$ for any~$c>\max\left(\sqrt{M_{\Gamma}\left\|R \right\|},4\left\|R \right\|\right)$. Moreover for any such~$c$, we have
\begin{equation}\label{eq:SeriesBound}
 \left\|\left( \lambda I_N -  S(\infty)\right)^{-1}\sum_{k=1}^{\infty} \left(c^{-2}R\left( \lambda I_N - S(\infty)\right)^{-1} \right)^k \right\|\leq  M_{\Gamma}\sum_{k=1}^{\infty}(q(c))^k = \frac{  M_{\Gamma} q(c)}{1-q(c)} = \text{O}\left(c^{-2} \right) ,
\end{equation}
so the Neumann series converges absolutely and uniformly on $ \Gamma$. This proves~\eqref{eq:Neumann}.  
%%
%%%
Combining    \eqref{eq:Proj2} and~\eqref{eq:Neumann} proves~\eqref{Neumann1}, and this completes the proof of Lemma~\ref{lem:Aux2}.
%%%%%%%%%%%%%%%%%%%%%%%
\end{Proof}

We can now prove Theorem~\ref{Thm:ZetaCoords}. Define~$\varepsilon(c):=(   \mathcal{P}(c) -  \mathcal{P}(\infty))\nu(\infty)$. Then
\begin{align*}
    \xi(c)  & =   \mathcal{P}( c )   \nu(\infty)\\
    &= (  \mathcal{P}( c ) -  \mathcal{P}( \infty )+  \mathcal{P}( \infty ))  \nu(\infty)\\
    &= \varepsilon(c)+     \nu(\infty),
    %%%
\end{align*}
and using Lemma~\ref{lem:Aux2} gives
%%%%%
\begin{align}\label{eq:XiC}
       |\varepsilon(c) | & = \left|\left(\sum_{k=1}^{\infty}\frac{1}{2\pi \imagi }\int_{\Gamma}\left( \lambda I_N -  S(\infty)\right)^{-1}\left(c^{-2}R\left( \lambda I_N -  S(\infty)\right)^{-1} \right)^k\text{d}\lambda\right)\nu(\infty)\right|\nonumber \\
    &  \leq  \frac{M_{\Gamma}}{2} \frac{q(c)}{1-q(c)}   |\nu(\infty) |\nonumber \\
    &= \text{O}\left(c^{-2} \right) . 
\end{align}
Now,~$ \eta(c) = D^{-1}\xi(c) $ gives 
\be\label{eq:Etac}
    \eta(c)-D^{-1}\nu(\infty)  =  D^{-1}\varepsilon(c). 
\ee
%%%%%%%%%%%%%%%
We conclude that for any~$c>\max\left(\sqrt{M_{\Gamma}\left\|R \right\|},4\left\|R \right\|\right)$, we have 
 \be\label{eq:bound1}
 \left|\eta_1(c)- 1\right|  = \left|\frac{1}{d_{11}}\varepsilon_1(c)\right| \leq \frac{1}{d_{11}} |\varepsilon(c) | \leq \frac{M_{\Gamma}}{2} \frac{q(c)}{1-q(c)}   |\nu(\infty)  |,
 \ee
%%%
 \be\label{eq:bound2}
 \left|\eta_2(c)- 1\right|  = \left|\frac{1}{d_{22}}\varepsilon_2(c)\right| \leq \frac{1}{d_{22}} |\varepsilon(c) | \leq \frac{M_{\Gamma}}{2} \frac{q(c)}{1-q(c)}   |\nu(\infty)  |,
 \ee
and for any~$i\in\{3,\dots,N\}$, we have 
%%%%
%%%%% 
%%%%
\begin{align}\label{eq:ZetaCoordsPf}
 \left|\eta_{i}(c) -  \alpha_1\cdots \alpha_{i-3}{\alpha_{i-2}}    c^{2-i} \right| &=  \left|\frac{1}{d_{ii}}\varepsilon_i(c)\right| \nonumber\\
 & \leq \frac{1}{d_{ii}} |\varepsilon(c) | \nonumber\\
&   = \frac{M_{\Gamma}}{2c^{i-2}}   \frac{q(c)}{1-q(c)}  |\nu(\infty)  | .
\end{align}
%%%%%%%%%%%%%%%%%%%%%%
Note that given $c>\max\left(\sqrt{M_{\Gamma}\left\|R \right\|},4\left\|R \right\|\right)$,   the upper bounds in~\eqref{eq:bound1}, \eqref{eq:bound2} and~\eqref{eq:ZetaCoordsPf}
can be computed \emph{explicitly}
using the expressions for the vector~$\nu(\infty)$ in~\eqref{eq:vinfexp},  the norm~$\left\|R\right\|$ in~\eqref{eq:normofmatr}, and the expression for~$q_c$  in \eqref{eq:qcdef}. Combining these upper bounds with~\eqref{eq:SeriesBound} yields~\eqref{eq:ZetaCoords}, and this completes the proof of Theorem~\ref{Thm:ZetaCoords}.
\end{Proof}
 
\begin{Remark}
%%%%%%%%%%%%%%%%%%%%%%%%%%%
Note that approximations of the coordinates of $\eta(c)$ up to arbitrarily small error can be obtained by computing higher order
terms in the Neumann series \eqref{Neumann1}. As an example, consider for simplicity the case $\alpha_i = 1$ for $i=1,\dots, N-2$, which implies $\left\|R \right\|=1$. Assume that $c>\max\left(\sqrt{M_{\Gamma}},4\right)$ and let~$\delta > 0$ be the desired error bound. Let $L>0$ be a sufficiently large integer such that
\begin{equation}\label{eq:SeriesBound1}
    \frac{(q(c))^L}{1-q(c)}  <\frac{2\delta}{ |\nu(\infty) |M_{\Gamma}}.
\end{equation}
Then, arguments similar to \eqref{eq:SeriesBound} imply that 
\begin{equation*}
 \left\|\left( \lambda I_N -  S(\infty)\right)^{-1}\sum_{k=L}^{\infty} \left(c^{-2}R\left( \lambda I_N - S(\infty)\right)^{-1} \right)^k \right\|\leq  \frac{M_{\Gamma}(q(c))^L}{1-q(c)}.
\end{equation*}
Hence, $\xi(c) = \mu(c)+ \bar{\varepsilon}(c)$, where
\begin{align*}
   |\bar{\varepsilon}(c)  |& = \left\|\left(\sum_{k=L}^{\infty}\frac{1}{2\pi \imagi  }\int_{\Gamma}\left( \lambda I_N -  S(\infty)\right)^{-1}\left(c^{-2}R\left( \lambda I_N -  S(\infty)\right)^{-1} \right)^k\text{d}\lambda\right)\nu(\infty)\right\|\\
    & \leq \frac{M_{\Gamma}}{2} \frac{(q(c))^L}{1-q(c)}   |\nu(\infty)  | \\&<\delta.
\end{align*}
The vector
\begin{equation}\label{eq:HighOrd}
\mu(c) = \left(\sum_{k=0}^{L-1}\frac{1}{2\pi \imagi  }\int_{\Gamma}\left( \lambda I_N -  S(\infty)\right)^{-1}\left(c^{-2}R\left( \lambda I_N -  S(\infty)\right)^{-1} \right)^k\text{d}\lambda\right)\nu(\infty)
\end{equation}
can be computed explicitly by the Cauchy residue theorem \cite{stein2010complex}. Using the fact that~$\eta(c) = D^{-1}\xi(c)$ yields 
\begin{equation*}
\begin{array}{lll}
&\max_{1\leq i \leq N}\left|\eta_i(c) - \frac{1}{d_{ii}}\mu_i(c)\right| =  \left|\frac{1}{d_{ii}}\bar{\varepsilon}_i(c)\right|\leq \frac{1}{d_{ii}} |\bar{\varepsilon}(c) |< \delta, \quad 1\leq i \leq N,
\end{array}
\end{equation*}
so  the entries of $D^{-1}\mu(c)$ approximate those of $\eta(c)$ with an error smaller than $\delta$.
\end{Remark}

 The next result  provides an explicit 
 expression for~$\left(\lambda I_N - S(\infty)\right)^{-1}$ with~$\lambda\in \Gamma$. This 
 can be used to derive an upper bound on the constant~$M_{\Gamma}$, given in \eqref{eq:qcdef}, thereby leading to an explicit estimate of the errors in \eqref{eq:ZetaCoordsPf}. Moreover, $\left(\lambda I_N - S(\infty)\right)^{-1}$ can be substituted into \eqref{eq:HighOrd} to obtain high order approximations of the Perron eigenvector.
\begin{Proposition}
  Let~$\Gamma$ be as in Figure~\ref{fig:SpectSeparateMod}. Then, for any~$\lambda\in \Gamma$
\begin{align}\label{eq:ResolventInverse}
     &\left(\lambda I_N - S(\infty)\right)^{-1} = \begin{bmatrix}
     D & 0_{2\times(N-2)} \\
     -E BD & E
     \end{bmatrix}, 
\end{align}
where
\begin{align}\label{eq:defB}
     &D = \frac{1}{\lambda^2-1}\begin{bmatrix}
 \lambda & 1\\
 1 & \lambda
\end{bmatrix}, \ E_{i,j} = \begin{cases}
    0,&  i<j,\\
     \lambda^{-1},   &i=j,\\
    \frac{\prod_{k=j+1}^{i}\alpha_k}{\lambda^{i-j+1}},&   i>j,
    \end{cases}  \text{ and } B = \begin{bmatrix}
 0 & -\alpha_1\\
 \vdots & \vdots \\
 0 & 0
\end{bmatrix}.
\end{align}
\end{Proposition}
%%%%%%%%%%%
\begin{proof}
We can divide~$\lambda I_n - S(\infty)$ into blocks as 
\begin{equation*}
\lambda I_n - S(\infty) = \begin{bmatrix}
 A & 0_{2\times(N-2)}\\
 B & C
\end{bmatrix},
\end{equation*}
with
\[
\ A := \begin{bmatrix}
 \lambda & -1\\
 -1 & \lambda
\end{bmatrix},\ C:=\begin{bmatrix}
 \lambda & 0 & 0&\dots & 0 & 0\\
 -\alpha_2 & \lambda& 0 &\dots & 0 & 0\\
 \vdots & \ddots & \ddots & \dots & \vdots & \vdots\\
 0 & 0 & 0 & \dots & -\alpha_{N-2} & \lambda
\end{bmatrix},
\]
and~$B$ defined in~\eqref{eq:defB}. 
Since $\lambda\neq 0$, it can be easily verified that~ $C^{-1} = E$ and~$A^{-1} = D$. Hence, 
\begin{equation*}
\begin{bmatrix}
 A & 0_{2\times(N-2)}\\
 B & C
\end{bmatrix} \begin{bmatrix}
     D & 0_{2\times(N-2)} \\
     -E BD & E
     \end{bmatrix} = \begin{bmatrix}
      AD & 0_{2\times(N-2)}\\
      BD-CEBD & CE 
     \end{bmatrix} = I_N,
\end{equation*}
and this completes the proof.
\end{proof}
 
 
 We can now prove the results in Section~\ref{sec:main}.
 %%%%%%%%%%%%%%%%%%%%%%%%%%%%%%%%%%%%%%%%%%%%%%%%%%%%%%%%%%%%%
 
\subsection{Proof of Proposition \ref{Prop:Monotonicity}}
%%%%%%%%%%%%%%%%%%%%%%%%%%%%%%%%%%%%%%%%%%%%%%%%%%%%%%%%%%%%%
It was shown in~\cite{RFM_model_compete_J} that for any~$s>0$ the network satisfies the following persistence property for any initial condition.
After an arbitrarily short time the pool density is   positive, and all
the densities in the sites are larger than zero and smaller than one. In particular, this implies that 
%%%%%%
\be\label{eq:persis}
e_z(q)>0,\quad   0< e_j^p(q) <1
\ee
for any integer~$q>0$, any~$p\in\{1,\dots,q\}$, and any~$j\in\{1,\dots,n_p\}$.

Seeking a contradiction, assume that the claim in Proposition \ref{Prop:Monotonicity} is not true. Then  there exists an integer~$m>0$ such that 
\begin{equation}\label{eq:ContradMonot}
    e_z(m+1)\geq e_z(m)>0.
\end{equation}
 
 Let 
\[
\mathcal{S}(i,q) := \sum_{j=1}^{n_i}e^i_j(q),  
\]
i.e., the total  steady-state  density  of ribosomes in RFMIO~$\#i$    when the network includes~$q$ RFMIOs. By~\eqref{eq:persis}, we have $\mathcal{S}(m+1,m+1)>0$. Taking into account \eqref{eq:ContradMonot} and the fact that the total number of ribosomes is fixed at $s$, we conclude that there exists some $1\leq p \leq m$ such that 
\begin{equation*}
    \mathcal{S}(p,m+1) < \mathcal{S}(p,m),
\end{equation*}
i.e., the total steady-state density of ribosomes along  RFMIO~$\#p$  
has decreased due to the addition of RFMIO~$\#(m+1)$ to the network.
In particular, for at least one site~$j \in\{1,\dots, n_p\}$, we have 
\begin{equation}\label{eq:SiteDecrease}
    e_j^p(m+1) < e_j^p(m).
\end{equation}

  Consider the steady state equations of RFMIO~$\#p$ when
the network has~$q$ RFMIOs, namely, 
%%
\begin{align}\label{eq:rfmiop}
%%%%%%%
\text{Site 1:}\quad 0 &  = G_p(e_z(q)) \lambda_0^p (1-e_1^p(q))-\lambda_1^p e_1^p(q)(1-e_2^p(q)),\nonumber \\
%%%%%%%%%%%%%%%%%%%%%%
\text{Site 2:}\quad 0 & =\lambda_1^p e_1^p(q)(1-e_2^p(q))-\lambda_2^p e_2^p(q)(1-e_3^p(q)),\nonumber\\
%%%%%%%%%%%%%%%%
\text{Site 3:}\quad 0 & =\lambda_2^p e_2^p(q)(1-e_3^p(q))-\lambda_3^p e_3^p(q)(1-e_4^p(q)),\nonumber\\
%%%%%%%%%%%%%%%%
&\vdots\\
%%%%%%%%%%%%%%%%%%%%%%
\text{Site} \  n_{p}-1:\quad 0 & =\lambda_{n_p-2}^p e_{n_p-2}^p(q)(1-e_{n_p-1}^p(q))-\lambda_{n_{p-1}}^p e_{n_p-1}^p(q)(1-e_{n_p}^p(q)),\nonumber\\
%%%%%%%%%%%%%%%%
\text{Site} \  n_{p}:\quad 0 & =\lambda_{n_{p}-1}^p e_{n_p-1}^p(q)(1-e_{n_p}^p(q))-\lambda_{n_p}^p e_{n_p}^p(q).\nonumber 
%%%%%%%%%%%%%%%%
%%%%%%%%%%%%%
\end{align}
Define a function~$\psi:(0,1)^2\to \mathbb{R}$ 
by $\psi(x,y) := \frac{x}{1-y}$, and note that  
\begin{equation}\label{eq:PsiMon}
    x_1\leq x_2, \ y_1\leq y_2\Rightarrow \psi(x_1,y_1)\leq \psi(x_2,y_2).
\end{equation}
Eq.~\eqref{eq:rfmiop} with $q=m+1$ yields  
%%%%%%%%%%%%%%%%%%%%%%%%%%%%%%%%%%%%%%
\begin{align}\label{eq:rfmiop2}
e_1^p(m+1) & = \frac{\lambda_{n_p}^p}{\lambda_{1}^p}
\frac{e_{n_p}^p(m+1)}{1 - e_2^p(m+1)} = \frac{\lambda_{n_p}^p}{\lambda_{1}^p}  \psi(e_{n_p}^p(m+1), e_2^p(m+1)),\nonumber\\
%%%%%%%%%%%%%%%%
&\vdots\\
%%%%%%%%%%%%%%%%%%%%%%
e_{n_p-2}^p(m+1) & =\frac{\lambda_{n_p}^p}{\lambda_{n_p-2}^p}   \frac{e_{n_p}^p(m+1)}{1 - e_{n_p-1}^p(m+1)} = \frac{\lambda_{n_p}^p}{\lambda_{n_p-2}^p}  \psi(e_{n_p}^p(m+1), e_{n_p-1}^p(m+1)),\nonumber  \\
%%%%%%%%%%%%%%%%
e_{n_p-1}^p(m+1) & =\frac{\lambda_{n_p}^p}{\lambda_{n_p-1}^p}   \frac{e_{n_p}^p(m+1)}{1 - e_{n_p}^p(m+1)} = \frac{\lambda_{n_p}^p}{\lambda_{n_p-1}^p}  \psi(e_{n_p}^p(m+1), e_{n_p}^p(m+1)).\nonumber 
%%%%%%%%%%%%%%%%
\end{align}
All these  expressions are well-defined by~\eqref{eq:persis}. 
We now consider  two cases.

\emph{Case 1.} Suppose that
\[
e_{n_p}^p(m+1) \geq e_{n_p}^p(m)  ,
\]
 i.e.,  the density in the last site of RFMIO~$\#p$ did not decrease due to the addition of RFMIO~$\#(m+1)$. 
  %%%
  By backward induction in \eqref{eq:rfmiop2} and using~\eqref{eq:PsiMon},
  we immediately obtain that 
  \[
  e^p_j(m+1)\geq e^p_j(m) \text{ for all } j=1,\dots, n_p.  
  \]
  This contradicts~\eqref{eq:SiteDecrease}. 
  
  \emph{Case 2.} Suppose that
  \be\label{eq:case2t}
  e_{n_p}^p(m+1) < e_{n_p}^p(m),
  \ee
   i.e.,  the density in the last site of RFMIO~$\#p$   decreased 
   due to the addition of RFMIO~$\#(m+1)$. 
  By backward induction and \eqref{eq:PsiMon}, we find that  
  \be\label{eq:np1}
  e^p_j(m+1)< e^p_j(m) \text{ for all }  j =1,\dots, n_p.
  \ee
Now using~\eqref{eq:rfmiop} with~$q=m$ and~$q=m+1$ gives
\begin{align*}
           \frac{e_{n_p}^p(m+1)}{e_{n_p}^p(m)}&= \frac{G_p(e_z(m+1))}{G_p(e_z(m))}  \frac{1-e_1^p(m+1)}{1-e_1^p(m)},        
%%%
\end{align*}    
and using~\eqref{eq:ContradMonot},
the monotonicity of~$G_p$, and~\eqref{eq:np1} yields~$  \frac{e_{n_p}^p(m+1)}{e_{n_p}^p(m)} > 1$. This contradicts~\eqref{eq:case2t}.

Summarizing, we see that~\eqref{eq:ContradMonot} cannot hold, 
and this  completes the proof.

%%%%%%%%%%%%%%%%%%%%%%%%%%%%%%%%%%%%%%
\subsection{Proof of Proposition \ref{Prop:Depletion}}
%%%%%%%%%%%%%%%%%%%%%%%%%%%%%%%%%%%%%%%%
Seeking a contradiction, assume  that the claim is not true.   Then  there exists~$\beta>0$ and a sequence~$m_1<m_2<\dots $ such that $e_z(m_k)\geq \beta$ for all~$k$. For any $k$ and any~$i=1,\dots, m_k $, let 
\[
\mathcal{S}(i,m_k) := \sum_{j=1}^{n_i}e^i_j(m_k),  
\]
i.e., the total density  of ribosomes along
RFMIO~$\#i$  at steady-state when the network includes~$m_k$ RFMIOs. Since the total number of ribosomes in the network is~$s$ (and this is independent of~$m_k$), we have 
\begin{align*}
    s &= e_z(m_k)+\sum_{i=1}^{m_k}\mathcal{S}(i,m_k)\\&\geq \beta +\sum_{i=1}^{m_k}\mathcal{S}(i,m_k),
\end{align*}
so  
\begin{equation}\label{eq:sobeta}
     0 \leq \sum_{i=1}^{m_k}\mathcal{S}(i,m_k)\leq s-\beta.
\end{equation}
Note that the bound on the right-hand side of~\eqref{eq:sobeta} does not depend on~$m_k$.
Let
\[
\varepsilon:=   \min\left\{
(  g_*  \lambda_*) / ( 4 \lambda^* ), 
1/2  \right\}  .
\]
Note that~$\varepsilon\in(0,1/2]$.
If for any~$m_k$ and any~$i\in\{1,\dots,m_k\}$ we have that~$S(i,m_k)>\varepsilon$ then for a large enough~$k$ this contradicts~\eqref{eq:sobeta}. Therefore, there exist~$k$ and~$p \in\{1,\dots,m_{k}\}$ 
such that~$\mathcal{S}(p,m_{k})\leq \varepsilon$. In particular,  
\be\label{eq:beei}
%%%%%%%%%
e_j^{p}(m_{k})\leq \mathcal{S}(p,m_{k})\leq \varepsilon\leq 1/2, \text{ for all }  j\in\{1,\dots,  n_{p}\}.
\ee
Consider the network with~$m_k$ RFMIOs    initialized at the equilibrium~$e(m_{k})$ at time~$t = 0$. 
Then  for any time~$t\geq 0$, we have 
\begin{align*}
%%%%
      \sum_{j=1}^{n_{p}}\dot x_j^{p}    & = G_{p}(e_z(m_{k})) \lambda_0^{p} (1-e_1^{p})-\lambda_{n_{p}}^{p} e_{n_{p}}^{p}\\
    & \geq G_{p}(\beta) \lambda_0^{p} (1-e_1^{p})-\lambda_{n_{p}}^{p} e_{n_{p}}^{p} \\
    & \geq  g_*\beta  \lambda_* \frac{1}{2}- \lambda^*\varepsilon\\&>0,
%%
\end{align*}
where the second line follows from the monotonicity of the 
pool output functions,
the third from the assumptions in the statement of the proposition and~\eqref{eq:beei}, and the fourth from the definition of~$\varepsilon$.
However, since the system is initialized at the steady state, $\sum_{j=1}^{n_{p}}\dot x_j^{p} =0$ for all~$t\geq 0$. 
This contradiction completes the proof.  
 
 
 
 
%%%%%%%%%%%%%%%%%%%%%%%%%
\subsection{Proof of Proposition~\ref{Prop:Depletion_now_eis}}
%%%%%%%%%%%%%%%%%%%%%%%%%%%%%%%%%%%%
Fix~$i\in\{1,\dots,m\}$, and 
consider RFMIO~$\#i$.   Setting 
\begin{equation}\label{eq:NotChange}
    N = n_i+2, \quad c = \left(\lambda_0^i G_i(e_z)\right)^{-\frac{1}{2}},\quad \alpha_j = (\lambda_j^i)^{-\frac{1}{2}}, \quad j = 1,\dots, n_i,
\end{equation}
in \eqref{eq:T_NSeq}, we have that $T\left[c,\alpha\right]$ is the matrix whose Perron eigenvalue $\sigma^i$ and eigenvector $\zeta^i$ are used to compute the steady state in RFMIO~$\#i$  via~\eqref{eq:ei_spectral}, that is, 
\begin{equation*}
    e_j^i =  \frac{\zeta_{j+2}^i}{ \left(\lambda_j^i\right)^{1/2}\sigma^i \zeta_{j+1}^i},\quad j=1,\dots,n_i.
\end{equation*}
%%%%%%%%%%%%%%%%
When~$e_z\to 0$, $c\to \infty$, and applying  \eqref{eq:ctoinf} and \eqref{eq:ZetaCoords0}
yields~\eqref{eq:Limration}. { If the pool output function~$G_i$ is differentiable at zero, then
\begin{equation*}
\lim_{e_z\to 0}\frac{e_j^i}{e_z} = \lim_{e_z\to 0}\left[\frac{e_j^i}{G_i(e_z)} \frac{G_i(e_z)-G_i(0)}{e_z} \right],
\end{equation*}
where we used the fact  that~$G_i(0) = 0$.}  Equation~\eqref{eq:eij_diff} now follows from~\eqref{eq:Limration}.
This completes the proof of Proposition~\ref{Prop:Depletion_now_eis}. 
%%%%%%%%%%%%%%%%%%%%

%%%%%%%%%%%%%%%%%%%%%%%%%
{
\subsection{Proof of Corollary~\ref{Corr:TotDens}}
%%%%%%%
Assume a sequence of 
networks with~$m_p$ RFMIOs, $p=1,2,\dots$, with~$\lim_{p\to \infty}m_p = \infty$. For any~$p$, let~$e_z(p)$ be the corresponding steady-state pool density. By Proposition \ref{Prop:Depletion},  $\lim_{p\to \infty}e_z(p)=0$. For any $p$, let $f_p(x) := \sum_{i=1}^{m_p}\frac{\lambda_{n_i}^ie_{n_i}^i(p)}{e_z(p)}\mathds{1}_{[i,i+1)}(x)$. Note that   $f_p(x)\geq 0$ for all $x$, and that  $\int_{-\infty}^{\infty}f_p(x)dx=\sum_{i=1}^{m_p}\frac{\lambda_{n_i}^ie_{n_i}^i(p)}{e_z(p)}$ is the ratio between the total production rate and the pool density for the case of $m_p$ RFMIOs in the network. By proposition \ref{Prop:Depletion_now_eis}, the functions $\left\{f_p \right\}_{p=1}^{\infty}$ converge pointwise to $f(x) := \sum_{i=1}^{\infty}\lambda_0^iG_i'(0)\mathds{1}_{[i,i+1)}(x)$. By Fatou's lemma \cite[Theorem 2.8.3]{bogachev2007measure}
    \begin{align*}
        \liminf_{p\to \infty}\frac{TPR}{e_z(p)} = \liminf_{p\to \infty}\int_{-\infty}^{\infty}f_p(x)dx\geq \int_{-\infty}^{\infty}f(x)dx = \sum_{i=1}^{\infty}\lambda_0^iG_i'(0)=\infty.
    \end{align*}
    The last equality follows from Assumption 1, which implies that  $\lambda_0^iG_i'(0)\geq \lambda_*g_*>0$. In the case where the number of RFMIOs is fixed at~$m$ as $e_z \to 0$, by replacing Fatou's lemma with the dominated convergence theorem \cite[Theorem 2.8.1]{bogachev2007measure}, the limit-inferior can be replaced by a limit, and  the inequality can be replaced by equality. Note that in this case $\sum_{i=1}^{m}\lambda_0^iG_i'(0)<\infty$. This proves~\eqref{eq:rtotalatinf}. 
    
    To prove~\eqref{eq:qatinf},
      consider first the case where the number of RFMIOs $m$ is fixed and the total density in the network satisfies $s\to 0$. Recall that $q = e_z/s$, and~\eqref{eq:sineq}
gives~$q=(1+\sum_{i=1}^m\sum_{j=1}^{n_i}\frac{e^i_j}{e_z})^{-1}$.
Using~\eqref{eq:eij_diff} yields 
    \begin{align*}
     \lim_{e_z\to 0}q &= \lim_{e_z\to 0}\left[1+\sum_{i=1}^m\sum_{j=1}^{n_i}\frac{e^i_j}{e_z} \right]^{-1}\\
    &=\left[1+\sum_{i=1}^m\sum_{j=1}^{n_i} \frac{\lambda_0^i   {G}_i'(0)} {\lambda_j^i} \right]^{-1}\\&=\left[1 + \sum_{i=1}^m  \frac {\lambda_0^i  {G}_i'(0)n_i} {  H (\lambda_1^i,\dots, \lambda_{n_i}^i  )} \right] ^{-1},
\end{align*}
and this completes the proof. 
We note in passing that in   the case where the total ribosome density in the network~$s$ is fixed and the number of RFMIOs satisfies $m\to \infty$,  Assumption 1 gives 
\begin{equation*}
    \frac{\lambda_0^iG_i'(0)}{\lambda_{n_i}^i}\geq \left(\lambda_*g_*\right)/\lambda^* \text{ for all } i,
\end{equation*}
  so in this case we have 
\begin{align*}
&\lim_{e_z\to 0}q   =\lim_{m\to \infty}\left[1+\sum_{i=1}^m\sum_{j=1}^{n_i} \frac{\lambda_0^i   {G}_i'(0)} {\lambda_j^i} \right]^{-1}=0.
\end{align*}
 


\subsection{Proof of Proposition~\ref{prop:Exch}}
%%%%%%%%%%%%%%%%%%
Using \eqref{eq:rtotalatinf} gives $\lim_{e_z\to 0}\frac{TPR}{e_z}=\lambda_0 g m$ and 
$\lim_{e_z\to 0}\frac{\overline{TPR}}{e_z}=\overline{\lambda}_0 \bar{g}$, and combining this with~\eqref{eq:ExchCond}
gives~\eqref{eq:NetwEquiv1}. Similarly, Eq.~\eqref{eq:qatinf} gives
%%%%
\begin{align*}
             \lim_{e_z\to 0}q = \frac{ g\lambda_0 m n}{H(\lambda_1,\dots,\lambda_n)} , 
             \text{ and }     \lim_{e_z\to 0}\overline{q}   = \frac{\overline{g}\overline{\lambda}_0  \overline{n}}{H(\overline{\lambda}_1,\dots, \overline{\lambda}_{\overline{n}} ) }      ,           
\end{align*}
and using \eqref{eq:ExchCond} and~\eqref{eq:ExchCond2} implies that these  expressions are  equal. 
%%%%%%%%%%%%%%
This completes the proof of Proposition~\ref{prop:Exch}. 


\bibliographystyle{IEEEtranS}
\bibliography{rfm_near_eq_bib}

 \end{document}


\begin{comment}
%%%%%%%%%%%%%%%%%%%%%%
MATLAB PROGRAMS - DO NOT DELETE
%%%%%%%%%%%%%%%%%

PROGRAM 1
%%%%%%%%%%%%%%%%
figure(1);hold off;
s=50;
for m=10:500
    coef=[ 1 2-s/m -(1+(s+1)/m) (s-1)/m-2 s/m  ]; 
    res=roots(coef);
    for i=1:4
        if (real(res(i))>0 & real(res(i))<1  & imag(res(i))<10^(-14) )
            e1=res(i);plot(m,e1,'*k'); hold on; 
            e2=e1/(1+e1);  plot(m,e2,'.k'); hold on;
            % ez=e1/(1-e1^2);  plot(m,ez,'ok'); hold on;
           % plot(m,s/2/m+1/4*(s-1)*s/m/m,'go')
        end
    end
end 
xlabel('$m$','fontsize',16,'interpreter','latex');grid on;
text(50,0.7,'$e_1$','fontsize',16,'interpreter','latex');grid on;
text(50,0.25,'$e_2$','fontsize',16,'interpreter','latex');grid on;
 
%%%% plot total production
%%%%%%%%%%%%%%%%%%%%%%%%%%
figure(2);
hold off;
s=50;
for m=10:500
    coef=[ 1 2-s/m -(1+(s+1)/m) (s-1)/m-2 s/m  ]; 
    res=roots(coef);
    for i=1:4
        if (real(res(i))>0 & real(res(i))<1  & imag(res(i))<10^(-14) )
            e1=res(i); 
            e2=e1/(1+e1); 
            plot(m, m*e2,'.k'); hold on;
        end
    end
end
xlabel('$m$','fontsize',16,'interpreter','latex');grid on;
 

\end{comment}
