## Supplementary figures and images for "Translation in the cell under fierce competition for shared resources: a mathematical model"

### all_eis_as_func_m_s50-eps-converted-to.pdf

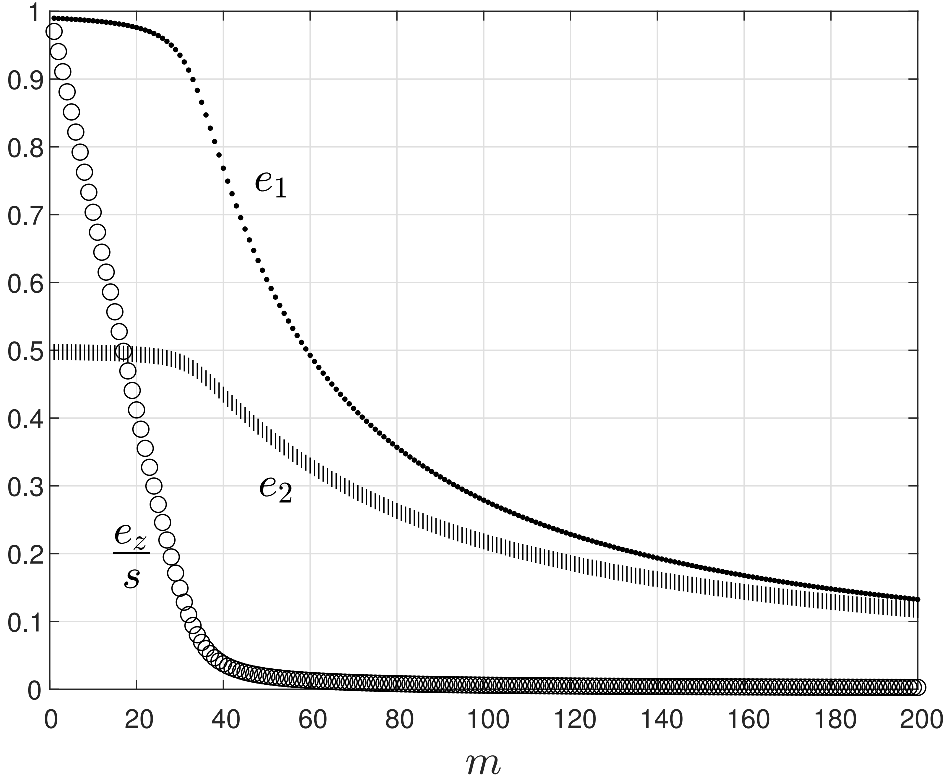

### curve_in_complex_plane-eps-converted-to.pdf

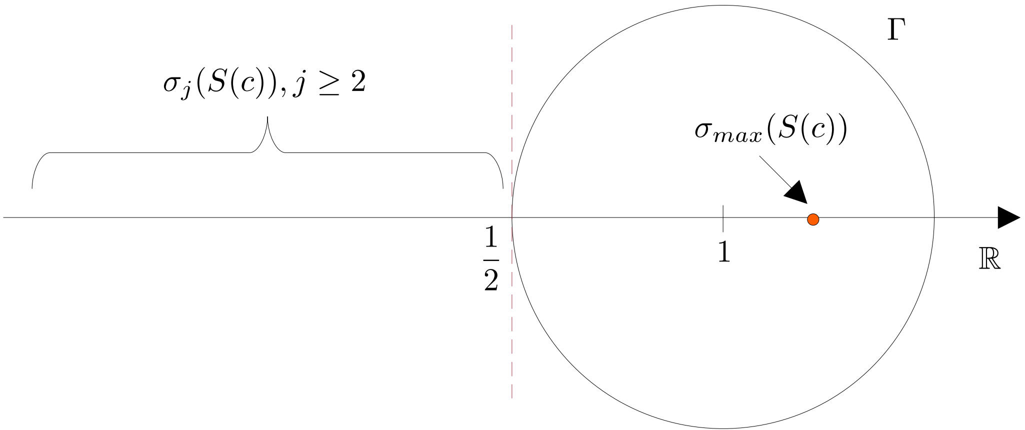

### diagram_1_eps_type-eps-converted-to.pdf

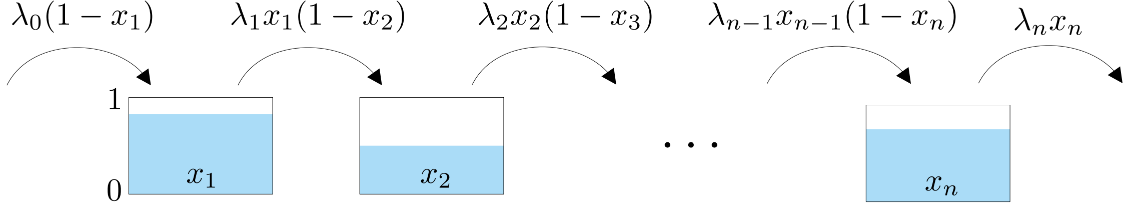

### diagram_2_for_eps_option_1-eps-converted-to.pdf

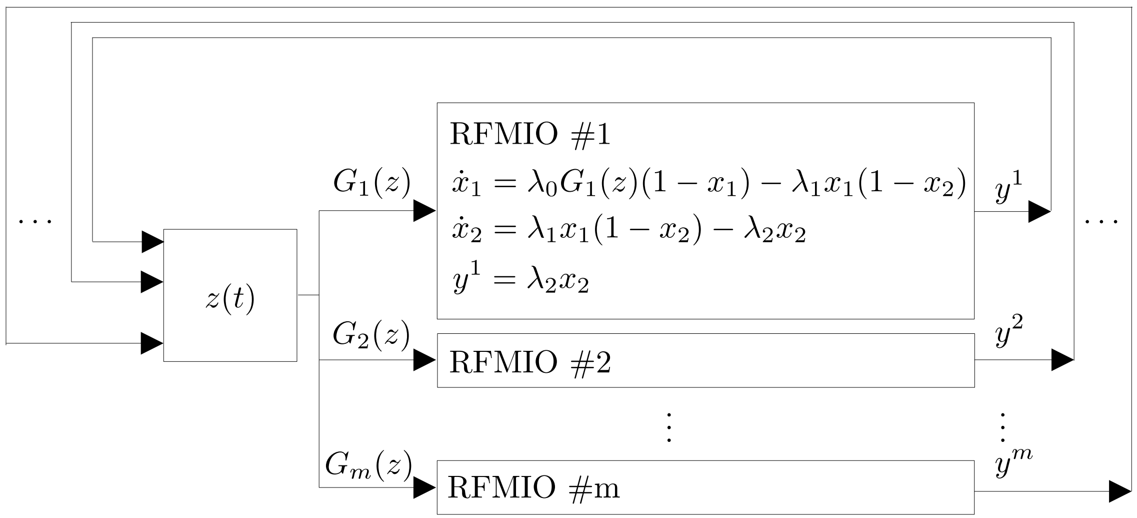

### ez_vs_ratio_s_constant_m_changes-eps-converted-to.pdf

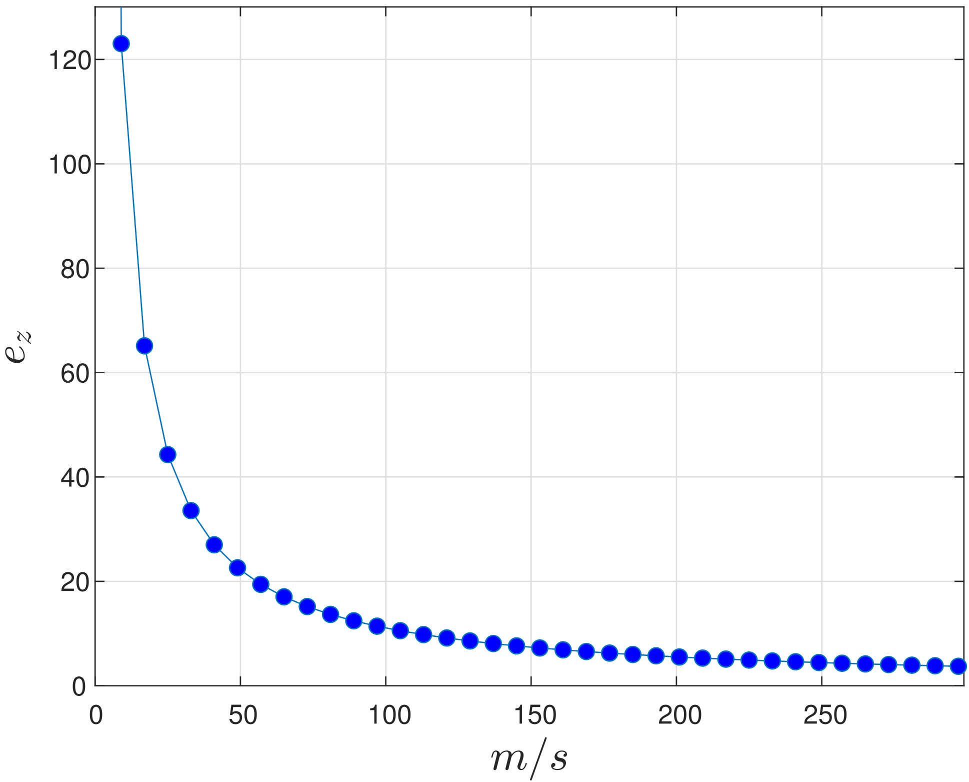

### ez_vs_ratio_updated.png

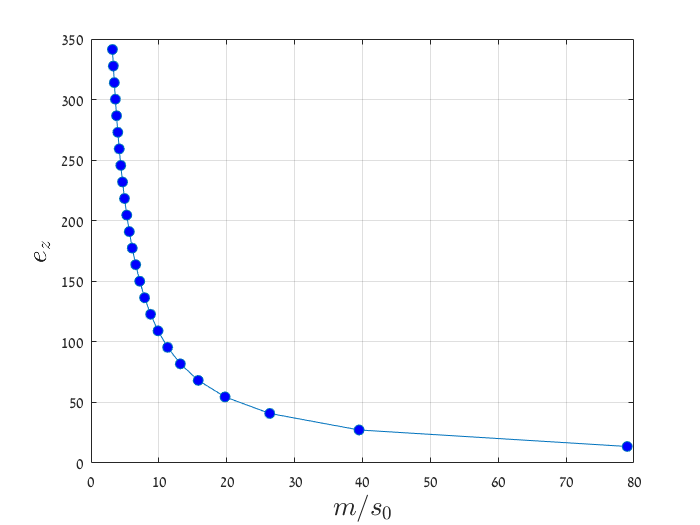

### relations_for_asymptotics_grid_3-eps-converted-to.pdf

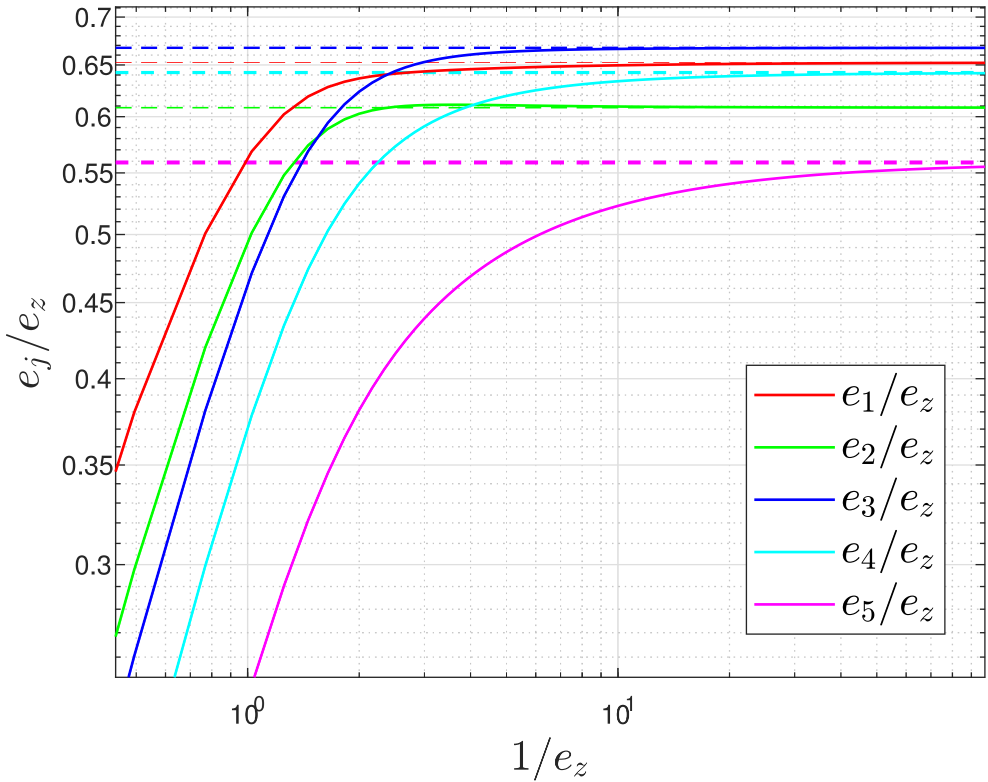

### tot_prate_as_func_m_s50-eps-converted-to.pdf

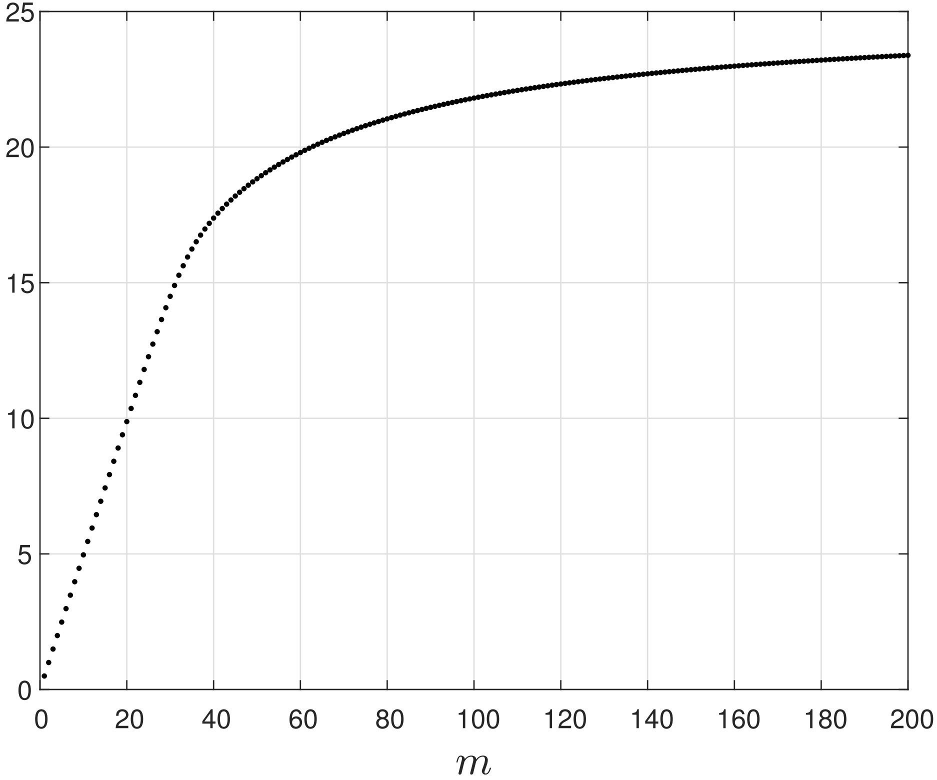

### TPR_vs_ratio_more_updated-eps-converted-to.pdf

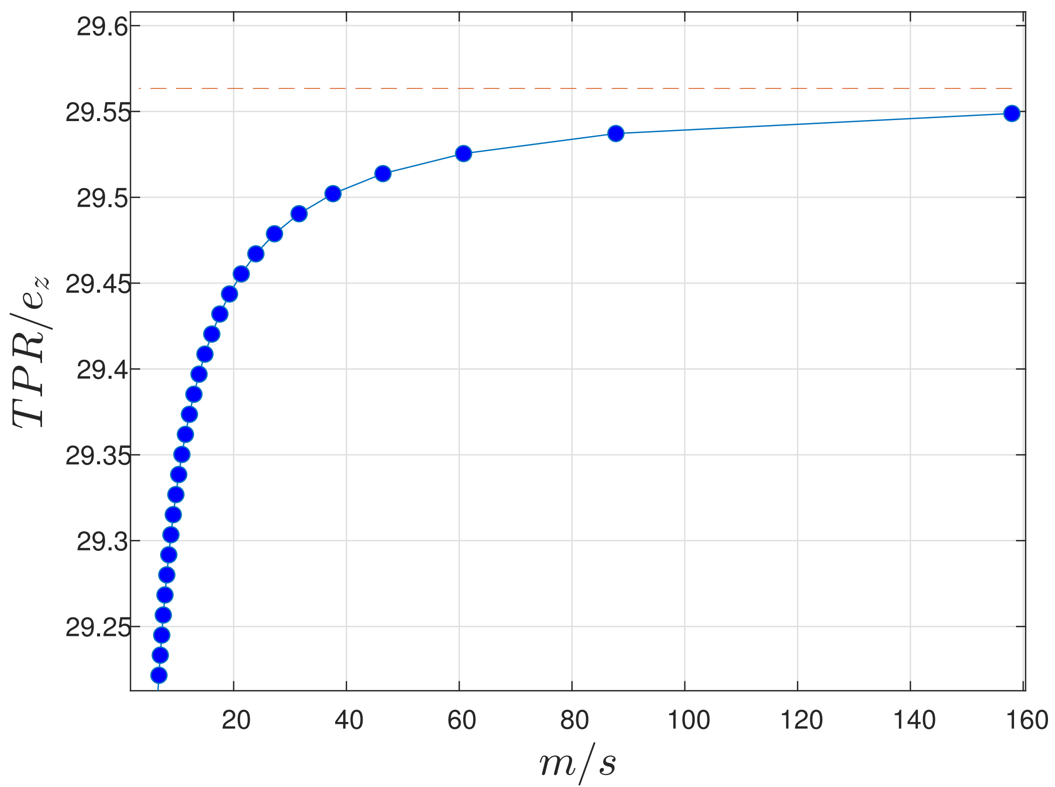
